# A mathematical framework for evo-devo dynamics

**DOI:** 10.1101/2021.05.17.444499

**Authors:** Mauricio González-Forero

## Abstract

Natural selection acts on phenotypes constructed over development, which raises the question of how development affects evolution. Classic evolutionary theory indicates that development affects evolution by modulating the genetic covariation upon which selection acts, thus affecting genetic constraints. However, whether genetic constraints are relative, thus diverting adaptation from the direction of steepest fitness ascent, or absolute, thus blocking adaptation in certain directions, remains uncertain. This limits understanding of long-term evolution of developmentally constructed phenotypes. Here we formulate a general tractable mathematical framework that integrates age progression, explicit development (i.e., the construction of the phenotype across life subject to developmental constraints), and evolutionary dynamics, thus describing the evolutionary developmental (evo-devo) dynamics. The framework yields simple equations that can be arranged in a layered structure that we call the evo-devo process, whereby five core elementary components generate all equations including those mechanistically describing genetic covariation and the evo-devo dynamics. The framework recovers evolutionary dynamic equations in gradient form and describes the evolution of genetic covariation from the evolution of genotype, phenotype, environment, and mutational covariation. This shows that genotypic and phenotypic evolution must be followed simultaneously to yield a dynamically sufficient description of long-term phenotypic evolution in gradient form, such that evolution described as the climbing of a fitness landscape occurs in “geno-phenotype” space. Genetic constraints in geno-phenotype space are necessarily absolute because the phenotype is related to the genotype by development. Thus, the long-term evolutionary dynamics of developed phenotypes is strongly non-standard: (1) evolutionary equilibria are either absent or infinite in number and depend on genetic covariation and hence on development; (2) developmental constraints determine the admissible evolutionary path and hence which evolutionary equilibria are admissible; and (3) evolutionary outcomes occur at admissible evolutionary equilibria, which do not generally occur at fitness landscape peaks in geno-phenotype space, but at peaks in the admissible evolutionary path where “total genotypic selection” vanishes if exogenous plastic response vanishes and mutational variation exists in all directions of genotype space. Hence, selection and development jointly define the evolutionary outcomes if absolute mutational constraints and exogenous plastic response are absent, rather than the outcomes being defined only by selection. Moreover, our framework provides formulas for the sensitivities of a recurrence and an alternative method to dynamic optimization (i.e., dynamic programming or optimal control) to identify evolutionary outcomes in models with developmentally dynamic traits. These results show that development has major evolutionary effects.

**Highlights:**

- We formulate a framework integrating evolutionary and developmental dynamics.
- We derive equations describing the evolutionary dynamics of traits considering their developmental process.
- This yields a description of the evo-devo process in terms of closed-form formulas that are simple and insightful, including for genetic covariance matrices.

## 1. Introduction

Development may be defined as the process that constructs the phenotype over life (Barresi and Gilbert, 2020). In particular, development includes “the process by which genotypes are transformed into phenotypes” (Wolf et al., 2001). As natural selection screens phenotypes produced by development, a fundamental evolutionary problem concerns how development affects evolution. Interest in this problem is long-standing (Baldwin 1896, Waddington 1959 p. 399, and Gould and Lewontin 1979) and has steadily increased in recent decades. It has been proposed that developmental constraints (Gould and Lewontin, 1979; Maynard Smith et al., 1985; Brakefield, 2006; Klingenberg, 2010), causal feedbacks over development occurring among genes, the organism, and environment (Lewontin, 1983; Rice, 2011; Hansen, 2013; Laland et al., 2015), and various development-mediated factors (Laland et al., 2014, 2015), namely plasticity (Pigliucci, 2001; West-Eberhard, 2003), niche construction (Odling-Smee et al., 1996, 2003), extragenetic inheritance (Baldwin, 1896; Cavalli-Sforza and Feldman, 1981; Boyd and Richerson, 1985; Jablonka and Lamb, 2014; Bonduriansky and Day, 2018), and developmental bias (Arthur, 2004; Uller et al., 2018), may all have important evolutionary roles. Understanding how development — including these elements acting individually and together — affects the evolutionary process remains an outstanding challenge (Baldwin, 1896; Waddington, 1959; Müller, 2007; Pigliucci, 2007; Laland et al., 2014, 2015; Galis et al., 2018).

Classic evolutionary theory indicates that development affects evolution by modulating the genetic covariation upon which selection acts. This can be seen as follows. In quantitative genetics, an individual’s *i*-th trait value *x*_*i*_ is written as 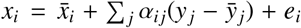, where the overbar denotes population average, *y* _*j*_ is the individual’s gene content at the *j*-th locus, α_*i j*_ is the partial regression coefficient of the *i*-th trait deviation from the average on the deviation from the average of the *j*-th locus content, and *e*_*i*_ is the residual (Fisher, 1918; Crow and Kimura, 1970; Falconer and Mackay, 1996; Lynch and Walsh, 1998; Walsh and Lynch, 2018). The quantity α_*i j*_ is Fisher’s additive effect of allelic substitution (his α; see Eq. I of Fisher 1918 and p. 72 of Lynch and Walsh 1998) and is a description of some of the linear effects of development, specifically of how genotypic change is transformed into phenotypic change. In matrix notation, the vector of an individual’s trait values is 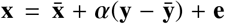, where the matrix *α* corresponds to what Wagner (1984) calls the developmental matrix (his **B**). The breeding value of the multivariate phenotype **x** is defined as 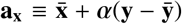, which does not consider the residual that includes non-linear effects of genes on phenotype. Breeding value thus depends on development via the developmental matrix *α*. The Lande (1979) equation describes the evolutionary change due to selection in the mean multivariate phenotype 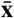 as 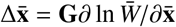, where the additive genetic covariance matrix is **G** ≡ cov[**a**_**x**_, **a**_**x**_] = *α*cov[**y, y**]*α*^⊺^ (e.g., Wagner 1984), mean absolute fitness is 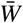, and the selection gradient is ∂ ln 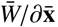, which points in the direction of steepest increase in mean fitness (here and throughout we use matrix calculus notation described in Appendix A). An important feature of the Lande equation is that it is in gradient form, so the equation shows that, within the assumptions made, phenotypic evolution by natural selection proceeds as the climbing of a fitness landscape. Moreover, the Lande equation shows that additive genetic covariation, described by **G**, may divert evolutionary change from the direction of steepest fitness ascent, and may prevent evolutionary change in some directions if genetic variation in those directions is absent (in which case **G** is singular). Since additive genetic covariation depends on development via the developmental matrix *α*, the Lande equation shows that development affects evolution by modulating genetic covariation via *α* (Charlesworth et al., 1982; Cheverud, 1984; Maynard Smith et al., 1985).

However, the Lande equation might provide a limited description of how development affects evolution for two reasons. First, the Lande equation does not explicitly consider phenotype construction, that is, development, but rather a regression-based description of development in terms of *α*. This limits the possibility of translating a mechanistic understanding of development, for instance, expressed as developmentally dynamic equations, into a mechanistic understanding of genetic covariation. Second, the Lande equation describes short-term evolution by assuming negligible allele frequency change (Walsh and Lynch, 2018, pp. 504 and 879). More specifically, the Lande equation describes the evolution of mean traits 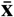 but not of mean gene content 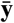, that is, it does not describe change in allele frequency but it depends on allele frequency; for instance, since *α* is a matrix of regression coefficients calculated for the current population, *α* depends on the current state of the population including allele frequency 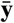. The Lande equation considers negligible genetic evolution by postulating Fisher’s (1918) infinitesimal model, whereby each trait is assumed to be controlled by an arbitrarily large number of loci such that allele frequency change per locus per generation is negligible (Bulmer, 1971, 1980; Turelli and Barton, 1994; Barton et al., 2017; Hill, 2017). Yet, over the long-term there is genetic evolution that is mapped to phenotypic evolution via development, but by assuming negligible genetic evolution the effect of this mapping might not be revealed by the Lande equation.

There is a lack of equations describing the long-term phenotypic evolutionary dynamics in gradient form while explicitly considering the developmental dynamics of phenotype construction and how development translates into genetic covariation. Various research lines have enabled the mathematical modelling of either long-term phenotypic evolutionary dynamics, age progression, development, or how development translates into genetic covariation, without including all these elements at the same time. Both the classic Lande equation (Lande, 1979) and the classic canonical equation of adaptive dynamics (Dieckmann and Law, 1996) describe the evolutionary dynamics of a multivariate trait in gradient form without an explicit account of development, by considering no explicit age progression nor development. Extensions to these equations have considered explicit age progression by implementing age structure, which allows individuals of different ages to coexist and to have age-specific survival and fertility rates (Lande, 1982; Charlesworth, 1993, 1994; Durinx et al., 2008). An important feature of age-structured models is that the forces of selection decline with age due to demography, in particular due to mortality and fewer remaining reproductive events as age advances (Medawar, 1952; Hamilton, 1966; Caswell, 1978; Caswell and Shyu, 2017). Although age progression is often taken as describing development, age structure does not explicitly describe development defined as phenotype construction.

Another research line in life-history theory has extended age-structured models to explicitly consider phenotype construction (Gadgil and Bossert, 1970; Taylor et al., 1974; León, 1976; Schaffer, 1983; Houston et al., 1988; Roff, 1992; Houston and McNamara, 1999; Sydsæter et al., 2008). This line has considered models with two types of age-specific traits: genotypic traits called control variables, which are under direct genetic control, and developed traits called state variables, which are constructed according to developmentally dynamic equations called dynamic constraints. This explicit consideration of development in an evolutionary context has mostly assumed that the population is at an evolutionary equilibrium. Thus, this approach identifies evolutionarily stable (or uninvadable) controls and associated states using techniques from dynamic optimization such as optimal control and dynamic programming (Gadgil and Bossert, 1970; Taylor et al., 1974; León, 1976; Schaffer, 1983; Houston et al., 1988; Roff, 1992; Houston and McNamara, 1999). While the assumption of evolutionary equilibrium yields great insight, it does not address the evolutionary dynamics which would provide a richer understanding. Moreover, the relationship between developmental dynamics and genetic covariation is not made evident by this approach.

Yet another research line has made it possible to mathematically model the evolutionary dynamics of developmentally constructed traits, but this has remained prohibitively challenging. A first step in this research line has been to consider function-valued or infinite-dimensional traits, which are genotypic traits indexed by a continuous variable (e.g., age) rather than a discrete variable as in the classic Lande equation. Thus, the evolutionary dynamics of univariate function-valued traits (e.g., body size across continuous age) has been described in gradient form by the Lande equation for function-valued traits (Kirkpatrick and Heckman, 1989) and the canonical equation for function-valued traits (Dieckmann et al., 2006). Although function-valued traits may depend on age, they are not constructed according to developmentally dynamic equations, so the consideration of the evolutionary dynamics of function-valued traits alone does not model the evolutionary dynamics of developmentally constructed traits. To our knowledge, Parvinen et al. (2013) were the first to mathematically model the evolutionary dynamics of developmentally constructed traits (the first to do it analytically; there have also been numerical models, for instance, integrating mathematical modeling of the developmental dynamics and individual-based modeling of the evolutionary dynamics; Salazar-Ciudad and Marín-Riera, 2013 and Watson et al., 2013). Parvinen et al. (2013) did so by considering the evolutionary dynamics of a univariate function-valued trait (control variable) that modulates the developmental construction of a multivariate developed trait (state variables; they refer to these as process-mediated models). However, the analysis of these models poses substantial technical challenges, by requiring calculation of functional derivatives and the solution of (partial) differential equations at evolutionary equilibrium with terminal age conditions in addition to the equations describing the developmental dynamics with initial age conditions (a two-point boundary value problem) (Dieckmann et al., 2006; Parvinen et al., 2013; Metz et al., 2016; Avila et al., 2021). Moreover, these models have yielded evolutionary dynamic equations in gradient form for genotypic traits, but not for developed traits (Dieckmann et al., 2006), so they have left unanswered the question of how the evolution of traits constructed via developmental dynamics proceeds in the fitness landscape. Additionally, these models have not provided a link between developmental dynamics and genetic covariation (Metz 2011; Dieckmann et al. 2006 discuss a link between constraints and genetic covariation in controls, not states; see Supplementary Information (SI) section S1.1 for further details).

Finally, a separate research line in quantitative genetics has analyzed how implicit development translates into genetic covariation. This line has considered models where a set of traits are functions of underlying traits such as gene content or environmental variables (Wagner, 1984, 1989; Hansen and Wagner, 2001; Rice, 2002; Martin, 2014; Morrissey, 2014, 2015). This dependence of traits on other traits is used by this research line to describe development and the genotype-phenotype map. However, this research line considers short-term evolution, no explicit age progression, and no explicit development (i.e., no developmentally dynamic phenotype construction). Although this line has provided an understanding of how implicit development translates into genetic covariation, it might miss evolutionary effects of the mapping of genotype to phenotype provided by development because of its assumption of negligible genetic evolution. Furthermore, this line has not provided equations describing the long-term evolution in gradient form of traits constructed according to explicit developmental dynamics.

Here we formulate a tractable mathematical framework to model the evolutionary dynamics of developmentally constructed traits. The framework provides closed-form equations to model the evolutionary dynamics of genotypic traits and the concomitant developmental dynamics of developed traits subject to developmental constraints. Moreover, the framework provides equations describing the long-term evolutionary dynamics in gradient form for developmentally constructed traits and equations that translate explicit development into genetic covariation. The framework is based on adaptive dynamics assumptions (Dieckmann and Law, 1996; Metz et al., 1996; Champagnat, 2006; Durinx et al., 2008). We consider general developmental dynamics that allow the phenotype to be “predisposed” to develop in certain ways, thus allowing for developmental bias (Arthur, 2004; Uller et al., 2018). We allow development to depend on the environment, which allows for a mechanistic description of plasticity (Pigliucci, 2001; West-Eberhard, 2003). We also allow development to depend on social interactions, which allows for a mechanistic description of extra-genetic inheritance (Boyd and Richerson, 1985; Jablonka and Lamb, 2014; Bonduriansky and Day, 2018) and indirect genetic effects (Moore et al., 1997). Social development entails that a mutant’s phenotype may change as the mutant genotype spreads, which complicates evolutionary invasion analysis. In turn, we allow the environment faced by each individual to depend on the traits of the individual and of social partners, thus allowing for individual and social niche construction although we do not consider ecological inheritance (Odling-Smee et al., 1996, 2003). We also let the environment depend on processes that are exogenous to the evolving population, such as eutrophication or climate change caused by members of other species, thus allowing for exogenous environmental change. To facilitate analysis, we let population dynamics occur over a short time scale, whereas environmental and evolutionary dynamics occur over a long time scale. Crucially, we measure age in discrete time, which simplifies the mathematics yielding closed-form formulas for otherwise implicitly defined quantities. Our methods use concepts from optimal control (Sydsæter et al., 2008) and integrate tools from adaptive dynamics (Dieckmann and Law, 1996) and matrix population models (Caswell, 2001; Otto and Day, 2007). In particular, we conceptualize the genotype as “control” variables and the phenotype as “state” variables whose developmental dynamics are modulated by controls. While we use concepts from optimal control, we do not use optimal control itself. Instead, we derive a method to model the evolutionary dynamics of controls, which yields an alternative method to optimal control that can be used to obtain optimal controls in a broad class of evolutionary models with dynamic constraints. Our use of optimal control concepts is thus useful to see how our results relate to optimal control, which has wide-ranging theory and applications.

We obtain three sets of main results. First, we derive several closed-form formulas for the total selection gradient of genotypic traits 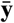 (i.e., of control variables) that affect the development of the phenotype 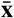(i.e., of state variables), formulas that can be easily computed with elementary operations. Second, we derive equations in gradient form describing the evolutionary dynamics of developed traits 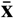 and of the niche-constructed environment. These equations depend on a mechanistic counterpart of the developmental matrix *α*, for which we obtain developmentally explicit and evolutionarily dynamic formulas for a broad class of models. Such formulas are in terms of closed-form formulas that we derive for the sensitivity of a system of recurrence equations, which are of use beyond evolutionary or biological applications. The obtained equation describing the evolutionary dynamics of the developed traits 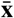 is generally dynamically insufficient because it depends on the genotypic traits 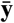, whose evolutionary dynamics are not described by the equation, a problem similarly present in the classic Lande equation. Third, we obtain synthetic equations in gradient form simultaneously describing the evolutionary dynamics of genotypic, developed, and environmental traits. These equations are in gradient form and are dynamically sufficient in that they include as many evolutionarily dynamic equations as evolutionarily dynamic variables, which enables one to describe the long-term evolution of developed multivariate phenotypes as the climbing of a fitness landscape. Such equations describe the evolutionary dynamics of the constraining matrix analogous to **G** as an emergent property, where genotypic traits 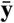 play an analogous role to that of allele frequency under quantitative genetics assumptions while linkage disequilibrium is not an issue as we assume clonal reproduction. The obtained dynamically sufficient gradient system depends on a constraining matrix that is always singular, which is mathematically trivial, but biologically crucial. The reason is that this singularity entails that genetic variation is necessarily absent in certain directions such that adaptive evolution at best converges to outcomes defined by both selection and development rather than by selection alone. Consequently, we find that development plays a major evolutionary role.

## 2. Problem statement

We begin by describing the mathematical problem we address. We consider a finite age-structured population with deterministic density-dependent population dynamics with age measured in discrete time. Each individual is described by three types of traits that we call genotypic, phenotypic (or developed), and environmental, all of which can vary with age and can evolve. We let all traits take continuous values, which allows us to take derivatives. Genotypic traits are defined by being directly specified by the genotype. Phenotypic traits are defined by being constructed over life subject to a developmental constraint: for instance, a phenotypic trait may be body size subject to the influence of genes, developmental history, environment, social interactions, and developmental processes constructing the body. Environmental traits are defined as describing the local environment of the individual subject to an environmental constraint: for instance, an environmental trait may be ambient temperature, which the individual may adjust behaviorally such as by roosting in the shade. We assume that reproduction transmits genotypic traits clonally, but developed and environmental traits need not be transmitted clonally due to social interactions. Given clonal reproduction of genotypic traits, we do not need to specify the genetic architecture (e.g., ploidy, number of loci, or linkage) and it may depend on the particular model. We assume that the genotypic traits are *developmentally independent*, whereby genotypic traits are entirely specified by the individual’s genotype and do not depend on other traits expressed over development: in particular, this means that a given genotypic trait at a given age can only be modified by mutation, but does not depend on other genotypic traits, the phenotype, or the environment. Developmental independence corresponds to the notion of “open-loop” control of optimal control theory (Sydsæter et al., 2008). Genotypic traits may still be *mutationally* correlated, whereby genotypic traits may tend to mutate together or separately. We assume that environmental traits are mutually independent, which facilitates derivations. We obtain dynamically sufficient equations in gradient form for the evolution of the phenotype by aggregating the various types of traits. We give names to such aggregates for ease of reference. We call the aggregate of the genotype and phenotype the geno-phenotype. We call the aggregate of the genotype, phenotype, and environment the geno-envo-phenotype.

The above terminology departs from standard terminology in adaptive dynamics as follows. In adaptive dynamics, our genotypic traits are referred to as the phenotype and our phenotypic traits as state variables. We depart from this terminology to follow the biologically common notion that the phenotype is constructed over development. In turn, adaptive dynamics terminology defines the environment as any quantity outside the individual, and thus refers to the global environment. In contrast, by environment we refer to the local environment of the individual. This allows us to model niche construction as the local environment of a mutant individual may differ from that of a resident.

We use the following notation (Table 1). Each individual can live from age 1 to age *N*_a_ ∈ {2, 3, …}. Each individual has a number *N*_g_ of genotypic traits at each age. A mutant’s genotypic trait *i* ∈ {1, …, *N*_g_} at age *a* ∈ {1, …, *N*_a_} is *y*_*ia*_ ∈ ℝ. For instance, *y*_*ia*_ may be the value of a life-history trait *i* at age *a* assumed to be directly under genetic control (i.e., a control variable in life-history models; Gadgil and Bossert, 1970; Taylor et al., 1974; León, 1976; Schaffer, 1983). Given our assumption of developmental independence of genotypic traits, the genotypic trait value *y*_*ia*_ for all *i* ∈ {1, …, *N*_g_} and all *a* ∈ {1, …, *N*_a_} of a given individual is exclusively controlled by her genotype but mutations can tend to change the value of *y*_*ia*_ and *y*_*k j*_ simultaneously for *k* ≠*i* and *j* ≠*a*. Additionally, each individual has a number *N*_p_ of developed traits, that is, of phenotypes at each age. A mutant’s phenotype *i* ∈ {1, …, *N*_p_} at age *a* ∈ {1, …, *N*_a_} is *x*_*ia*_ ∈ R. Moreover, each individual has a number *N*_e_ of environmental traits that describe her local environment at each age. A mutant’s environmental trait *i* ∈ {1, …, *N*_e_} at age *a* ∈ {1, …, *N*_a_} is ϵ_*ia*_ ∈ ℝ.. We do not consider the developmental or evolutionary change of the number of traits (i.e., of *N*_g_, *N*_p_, or *N*_e_), but our framework allows for the modeling of the developmental or evolutionary origin of novel traits (e.g., the origin of a sixth digit where there was five previously in development or evolution; Chan et al., 1995; Litingtung et al., 2002; Müller, 2010) by implementing a suitable codification (e.g., letting *x*_*ia*_ mean sixth-digit length, being zero in a previous age or evolutionary time).

**Table 1:**
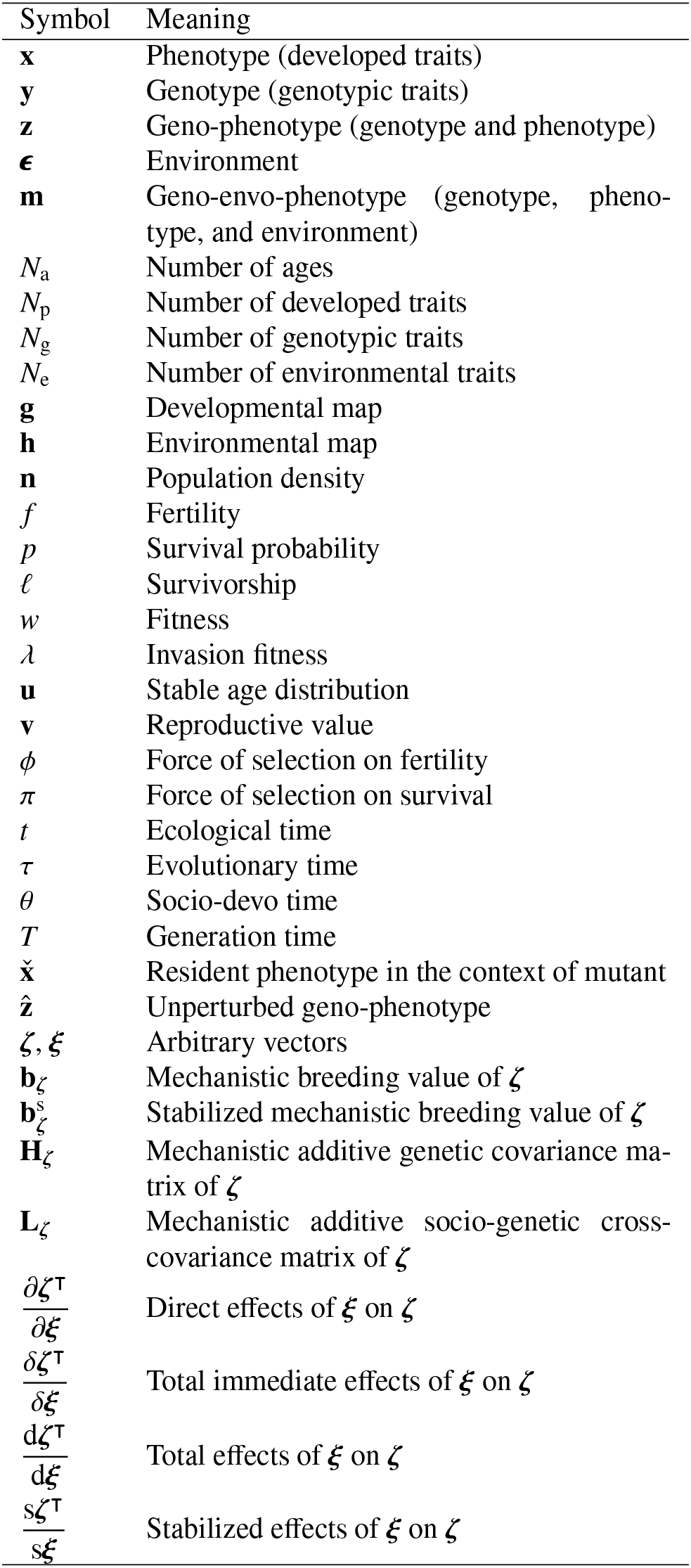
Notation summary.

We use the following notation for collections of these quantities. A mutant’s *i*-th genotypic trait across all ages is denoted by the column vector 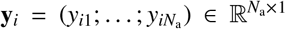, where the semicolon indicates a line break, that is, 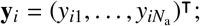 we write 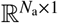 rather than simply 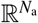 as it will be important to clearly distinguish column from row vectors. A mutant’s *i*-th phenotype across all ages is denoted by the column vector 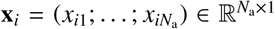. A mutant’s *i*-th environmental trait across all ages is denoted by the column vector 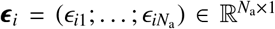. A mutant’s genotype across all genotypic traits and all ages is denoted by the block column vector 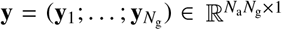. A mutant’s phenotype across all developed traits and all ages is denoted by the block column vector 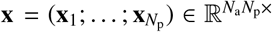. A mutant’s environment across all environmental traits and all ages is denoted by the block column vector 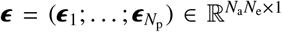. To simultaneously refer to the genotype and phenotype, we denote the geno-phenotype of the mutant individual at age *a* as 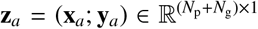, and the geno-phenotype of a mutant across all ages as 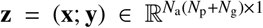. Moreover, to simultaneously refer to the genotype, phenotype, and environment, we denote the geno-envo-phenotype of a mutant individual at age *a* as 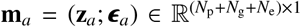, and the geno-envo-phenotype of the mutant across all ages as 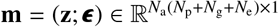. We denote resident values analogously with an overbar (e.g., 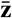 is the resident geno-phenotype).

The developmental process that constructs the phenotype is as follows, with causal dependencies described in Fig. 1. We assume that an individual’s multivariate phenotype at a given age is a function of the genotypic, phenotypic, and environmental traits that the individual had at the immediately previous age as well as of the social interactions experienced at that age. Thus, we assume that a mutant’s multivariate phenotype at age *a* + 1 is given by the developmental constraint

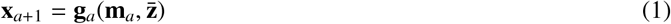

for all *a* ∈ {1, …, *N*_a_ − 1} with initial condition 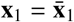. The function

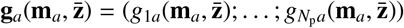

is the developmental map at age *a*, which we assume is a continuously differentiable function of the individual’s geno-envo-phenotype at that age and of the geno-phenotype of the individual’s social partners who can be of any age; thus, an individual’s development directly depends on the individual’s local environment but not directly on the local environment of social partners. For simplicity, we assume that the phenotype 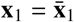 at the initial age is constant and does not evolve. This assumption corresponds to the common assumption in life-history models that state variables at the initial age are given (Gadgil and Bossert, 1970; Taylor et al., 1974; León, 1976; Schaffer, 1983; Sydsæter et al., 2008). The term developmental function can be traced back to Gimelfarb (1982) through Wagner (1984).

**Figure 1.**
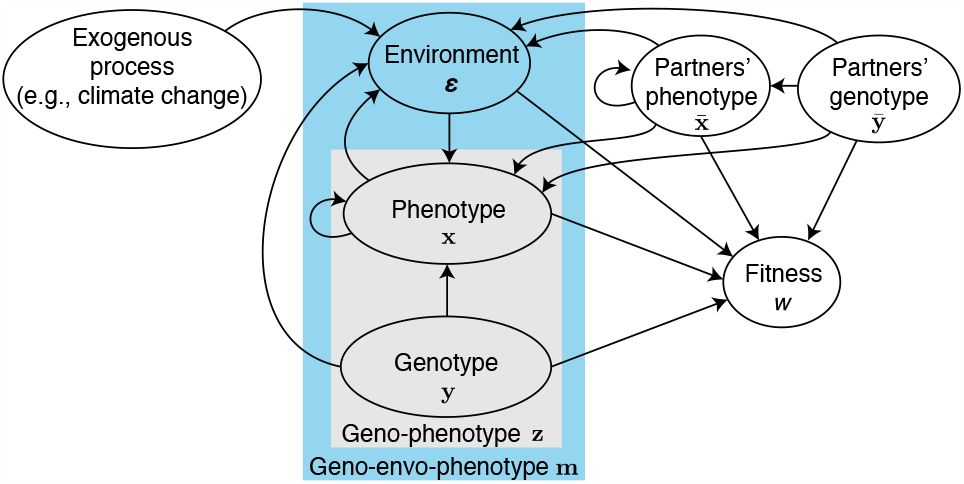
Causal diagram among the framework’s components. Variables have age-specific values that are not shown for clarity. The phenotype **x** is constructed by a developmental process. Each arrow indicates the direct effect of a variable on another one. A mutant’s genotypic traits may directly affect the phenotype (with the slope quantifying developmental bias from genotype), environment (niche construction by genotype), and fitness (direct selection on genotype). A mutant’s phenotype at a given age may directly affect her phenotype at an immediately subsequent age (quantifying developmental bias from the phenotype), thus the direct feedback loop from phenotype to itself. A mutant’s phenotype may also directly affect her environment (niche construction by the phenotype) and fitness (direct selection on the phenotype). A mutant’s environment may directly affect the phenotype (plasticity) and fitness (environmental sensitivity of selection). The social partners’ genotype may directly affect their own phenotype (quantifying developmental bias from genotype), the mutant’s phenotype (indirect genetic effects from genotypes), and the mutant’s fitness (social selection on genotype). The social partners’ phenotype at a given age may directly affect their own phenotype at an immediately subsequent age (quantifying developmental bias from phenotypes), thus the direct feedback loop. The social partners’ phenotype at a given age may also directly affect the mutant’s phenotype (quantifying indirect genetic effects from the phenotype), the mutant’s environment (social niche construction), and the mutant’s fitness (social selection on the phenotype). The environment may also be directly influenced by exogenous processes. We assume that the genotype is developmentally independent (i.e., controls **y** are open-loop), which means that there is no arrow towards the genotype.

The developmental constraint (1) allows for a wide range of models. Eq. (1) is a mathematical, deterministic description of Waddington’s (1957) “epigenetic landscape”. Eq. (1) is a constraint in that the phenotype **x**_*a*+1_ cannot take any value but only those that satisfy the equality (e.g., an individual’s body size today cannot take any value but depends on her body size, gene expression, and environment since yesterday). The developmental map in Eq. (1) is an extension of the notions of genotype-phenotype map (often a function from genotype to phenotype, without explicit developmental dynamics) and reaction norm (often a function from environment to phenotype, also without explicit developmental dynamics), as well as of early mathematical descriptions of development in an evolutionary context (Alberch et al., 1979). The developmental constraint (1) can describe gene regulatory networks (Alon, 2020), for instance, letting *x*_*ia*_ be the expression level of gene *i*; learning in deep neural networks (Russell and Norvig, 2021), for instance, letting **ϵ**_*a*_ describe the network’s input at age *a* and **x**_*a*_ describe the network’s weights at that age; and reaction-diffusion models of morphology (Murray, 2003), for instance, letting *x*_*ia*_ be a morphogen’s level at the *i*-th spatial location at age *a* (SI section S1.2). The developmental map in Eq. (1) may be non-linear and can change over development (e.g., from *g*_*ia*_ = sin *x*_*ia*_ to 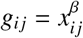 for *a* < *j* and some parameter β, for instance, due to metamorphosis) and over evolution (e.g., from a sine to a power function if 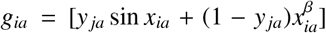as genotypic trait *y* _*ja*_ evolves from 0 to 1). The dependence of the mutant phenotype on the phenotype of social partners in (1) allows one to implement Jablonka and Lamb’s (2014) notion that extra-genetic inheritance transmits the phenotype rather than the genotype (see their p. 108), such that the mutant phenotype can be a possibly altered copy of social partners’ phenotype. Simpler forms of the developmental constraint (1) are standard in life-history models (Gadgil and Bossert, 1970; Taylor et al., 1974; León, 1976; Schaffer, 1983; Sydsæter et al., 2008) and physiologically structured models of population dynamics (de Roos, 1997, Eq. 7).

We describe the local environment as follows. We assume that an individual’s local environment at a given age is a function of the genotypic traits, phenotype, and social interactions of the individual at that age, and of processes that are not caused by the population considered. Thus, we assume that a mutant’s environment at age *a* is given by the environmental constraint

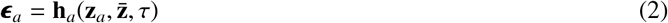

for all *a* ∈ {1, …, *N*_a_}. The function

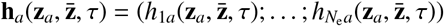

is the environmental map at age *a*, which can change over development and evolution. We assume that the environmental map is a continuously differentiable function of the individual’s geno-phenotype at that age (e.g., the individual’s behavior at a given age may expose it to a particular environment at that age), the geno-phenotype of the individual’s social partners who can be of any age (e.g., through social niche construction), and evolutionary time τ due to slow exogenous environmental change (so the exogenous process changing the environment in Fig. 1 acts as forcing, as τ appears explicitly in Eq. 2). We assume slow exogenous environmental change to enable the resident population to reach carrying capacity to be able to use relatively simple techniques of evolutionary invasion analysis to derive selection gradients. The environmental constraint (2) may also be non-linear and can change over development (i.e., over *a*) and over evolution (as the genotype or phenotype evolves or exogenously as evolutionary time advances).

The environmental constraint (2) is a minimalist description of the environment of a specific kind (akin to “feed-back functions” used in physiologically structured models to describe the influence of individuals on the environment; de Roos, 1997). We use the minimalist environmental constraint (2) as a first approximation to shorten derivations; our derivations illustrate how one could obtain equations with more complex developmental and environmental constraints. With the minimalist environmental constraint (2), the environmental traits are mutually independent in that changing one environmental trait at one age does not *directly* change any other environmental trait at any age (i.e., ∂ϵ_*k j*_ /∂ϵ_*ia*_ = 0 if *i* ≠*k* or *a* ≠*j*). We say that development is social if 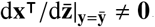.

Our aim is to obtain closed-form equations describing the evolutionary dynamics of the resident phenotype 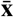 subject to the developmental constraint (1) and the environmental constraint (2). The evolutionary dynamics of the phenotype 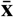 emerge as an outgrowth of the evolutionary dynamics of the genotype 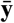and environment 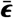. In the SI section S2.5, we show that under our assumptions the evolutionary dynamics of the resident genotype 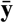 are given by the canonical equation of adaptive dynamics (Dieckmann and Law, 1996):

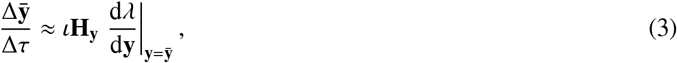

where 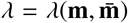 is invasion fitness, 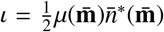 is a non-negative scalar measuring mutational input in terms of the mutation rate 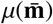 and the carrying capacity 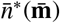, and **H**_**y**_ = cov[**y, y**] is the mutational covariance matrix (of genotypic traits). The selection gradient in Eq. (3) involves total derivatives so we call it the *total* selection gradient of the genotype, which measures the effects of genotypic traits **y** on invasion fitness λ across all the paths in Fig. 1 (λ has the same derivatives with respect to mutant trait values as fitness *w*, defined below). Total selection gradients differ from Lande’s selection gradient in that the latter is defined in terms of partial derivatives and so measures only the direct effects of traits on fitness (e.g., 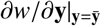 measures the effect of **y** on *w* only across the path directly connecting the two in Fig. 1). We will be concerned with describing the evolutionary dynamics to first-order of approximation, so we will treat the approximation in Eq. (3) as an equality although we keep the approximation symbols throughout to distinguish what is and what is not an approximation.

The arrangement above describes the evolutionary developmental (evo-devo) dynamics: the evolutionary dynamics of the resident genotype are given by the canonical equation (3), while the concomitant developmental dynamics of the phenotype are given by the developmental (1) and environmental (2) constraints evaluated at resident trait values. To complete the description of the evo-devo dynamics, we obtain closed-form expressions for the total selection gradient of the genotype. Moreover, to determine whether the evolution of the resident developed phenotype 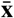 can be understood as the climbing of a fitness landscape, we derive equations in gradient form describing the evolutionary dynamics of the resident phenotype 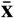, environment 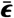, geno-phenotype 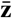, and geno-envo-phenotype 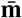. Before doing so, we first give an overview of the model underpinning the arrangement above, which describes a complication introduced by social development, how we handle it, and a fitness function that has the same gradient as invasion fitness in age-structured populations. We then use these descriptions to write our results.

## 3. Model overview

Here we give an overview of the model. We describe it formally in the SI section S2.

### 3.1. Set up

We base our framework on standard assumptions of adaptive dynamics, particularly following Dieckmann and Law (1996), but social development introduces a non-standard complication. We separate time scales, so developmental and population dynamics occur over a short discrete ecological time scale *t* and evolutionary dynamics occur over a long discrete evolutionary time scale τ. Although the population is finite, in a departure from Dieckmann and Law (1996), we let the population dynamics be deterministic rather than stochastic for simplicity, so there is no genetic drift. Thus, the only source of stochasticity in our framework is mutation. We assume that mutation is rare, weak, and unbiased. Weak mutation means that the variance of mutant genotypic traits around resident genotypic traits is marginally small (i.e., 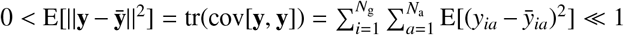). Weak mutation (Gillespie, 1983; Walsh and Lynch, 2018, p. 1003) is also called δ-weak selection (Wild and Traulsen, 2007). Unbiased mutation means that mutant genotypic traits are symmetrically distributed around the resident genotypic traits (i.e., the mutational distribution 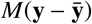 is even). Unbiased mutation in genotypic traits allows for bias in the distribution of mutant phenotypes (i.e., the distribution of 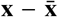 is not necessarily even). Thus, we do not make the isotropy assumption of Fisher’s (1930) geometric model (Orr, 2005), although isotropy may arise for mechanistic breeding values (defined below) with large *N*_a_*N*_g_ and additional assumptions (e.g., high pleiotropy and high developmental integration) from the central limit theorem (Martin, 2014). We assume that a monomorphic resident population having geno-envo-phenotype 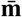 undergoes density-dependent population dynamics that bring it to carrying capacity. At this carrying capacity, rare mutant individuals arise which have a marginally different genotype **y** and that develop their phenotype in the context of the resident. If the mutant genotype increases in frequency, it increasingly faces mutant rather than resident individuals. Thus, with social development, the mutant phenotype may change as the mutant genotype spreads, which complicates invasion analysis.

### 3.2. A complication introduced by social development

With social development, the phenotype an individual develops depends on the traits of her social partners. This introduces a complication to standard evolutionary invasion analysis, for two reasons. First, the phenotype of a mutant genotype may change as the mutant genotype spreads and is more exposed to the mutant’s traits via social interactions, making the mutant phenotype frequency dependent. Thus, the phenotype developed by a rare mutant genotype in the context of a resident phenotype may be different from the phenotype developed by the same mutant genotype in the context of itself once the mutant genotype has approached fixation. Second, because of social development, a recently fixed mutant may not breed true, that is, her descendants may have a different phenotype from her own despite clonal reproduction of the genotype and despite the mutant genotype being fixed (Fig. 2; see also Kobayashi et al. 2015, Eq. 14 in their Appendix). Yet, to apply standard invasion analysis techniques, the phenotype of the fixed genotype must breed true, so that the phenotype of a mutant genotype developed in the context of individuals with the mutant genotype have the same phenotype.

**Figure 2.**
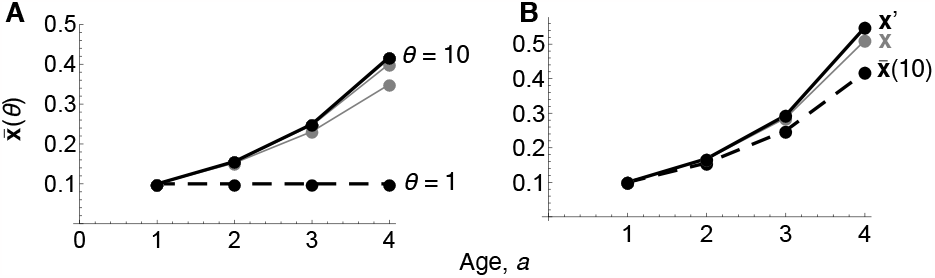
A difficulty introduced by social development. (A) Illustration of socio-devo dynamics converging to a quasi socio-devo stable equilibrium (solid, black line). The dashed line is a socio-devo initial resident phenotype 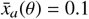 for all *a* ∈ {1, …, 4}, 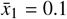, and socio-devo time θ = 1. The gray line immediately above is a phenotype developed in the context of such resident, where 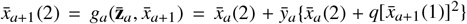, with 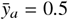 for all *a* ∈ {1, …, 4} and *q* = 0.5. Setting this phenotype 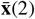 as resident and iterating up to θ = 10 yields the remaining gray lines, with iteration 10 given by the black line, where 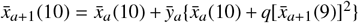 and 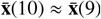 is approximately a socio-devo stable equilibrium, which breeds true. (B) Introducing in the context of such resident 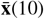(dashed line) a mutant genotype **y** yields the mutant phenotype **x** (gray line), where 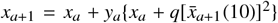 and *y*_*a*_ = 0.6 for all *a* ∈ {1, …, 4}. Such mutant does not breed true: a mutant **x**^′^ (solid black line) with the same genotype developed in the context of mutant **x** has a different phenotype, where 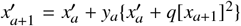. One can use socio-devo dynamics (A) to find for such mutant genotype **y** a phenotype that breeds true under social development.

Thus, to carry out invasion analysis, we proceed as follows. Ideally, one should follow explicitly the change in mutant phenotype as the mutant genotype increases in frequency and achieves fixation, and up to a point where the fixed mutant phenotype breeds true. Yet, to simplify the analysis, we separate the dynamics of phenotype convergence and the population dynamics. We thus introduce an additional phase to the standard separation of time scales in adaptive dynamics so that phenotypic convergence occurs first and then resident population dynamics follow. Such additional phase does not describe a biological process but is a mathematical technique to facilitate mathematical treatment (akin to using best-response dynamics to find Nash equilibria). However, this phase might still be biologically justified under somewhat broad conditions. In particular, Aoki et al. (2012, their Appendix A) show that such additional phase is justified in their model of social learning evolution if mutants are rare and social learning dynamics happen faster than allele frequency change; they also show that this additional phase is justified for their particular model if selection is δ-weak. As a first approximation, here we do not formally justify the separation of phenotype convergence and resident population dynamics for our model and simply assume it for simplicity.

### 3.3. Phases of an evolutionary time step

To handle the above complication introduced by social development, we partition a unit of evolutionary time in three phases: socio-developmental (socio-devo) dynamics, resident population dynamics, and resident-mutant population dynamics (Fig. 3).

**Figure 3.**
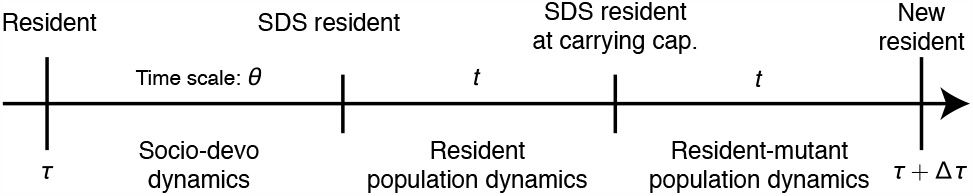
Phases of an evolutionary time step. Evolutionary time is τ. SDS means socio-devo stable. The socio-devo dynamics phase is added to the standard separation of time scales in adaptive dynamics, which only consider the other two phases. The socio-devo dynamics phase is only needed if development is social (i.e., if the developmental map **g**_*a*_ depends directly or indirectly on social partners’ geno-phenotype for some age *a*).

At the start of the socio-devo dynamics phase of a given evolutionary time τ, the population consists of individuals all having the same resident genotype, phenotype, and environment. A new individual arises which has identical genotype as the resident, but develops a phenotype that may be different from that of the original resident due to social development. This developed phenotype, its genotype, and its environment are set as the new resident. This process is repeated until convergence of the geno-envo-phenotype to what we term a “socio-devo stable” (SDS) resident equilibrium or until divergence. These socio-devo dynamics are formally described by Eq. S2.1.1 and illustrated in Fig. 2A. If development is not social, the resident is trivially SDS so the socio-devo dynamics phase is unnecessary. If an SDS resident is achieved, the population moves to the next phase; if an SDS resident is not achieved, the analysis stops. We thus study only the evolutionary dynamics of SDS resident geno-envo-phenotypes. More specifically, we say a geno-envo-phenotype 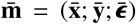 is a socio-devo equilibrium if and only if 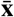 is produced by development when the individual has such genotype 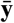 and everyone else in the population has that same genotype, phenotype, and environment (Eq. S2). A socio-devo equilibrium 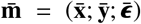 is locally stable (i.e., SDS) if and only if a marginally small deviation in the initial phenotype 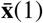 from the socio-devo equilibrium keeping the same genotype leads the socio-devo dynamics (Eq. S1) to the same equilibrium. A socio-devo equilibrium 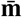 is locally stable if all the eigenvalues of the matrix

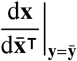

have absolute value (or modulus) strictly less than one. For instance, this is always the case if social interactions are only among peers (i.e., individuals of the same age) so the mutant phenotype at a given age depends only on the phenotype of immediately younger social partners (in which case the above matrix is block lower triangular so all its eigenvalues are zero; Eq. S5.6.9). We assume that there is a unique SDS geno-envo-phenotype for a given developmental map at every evolutionary time *τ*.

If an SDS resident is achieved in the socio-devo dynamics phase, the population moves to the resident population dynamics phase. Because the resident is SDS, an individual with resident genotype developing in the context of the resident geno-phenotype is guaranteed to develop the resident phenotype. Thus, we may proceed with the standard invasion analysis. Hence, in this phase of SDS resident population dynamics, the SDS resident undergoes density dependent population dynamics, which we assume asymptotically converges to a carrying capacity.

Once the SDS resident has achieved carrying capacity, the population moves to the resident-mutant population dynamics phase. At the start of this phase, a random mutant genotype **y** marginally different from the resident genotype 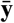 arises in a vanishingly small number of mutants. We assume that the mutant becomes either lost or fixed in the population (Geritz et al., 2002; Geritz, 2005; Priklopil and Lehmann, 2020), establishing a new resident geno-envo-phenotype.

Repeating this evolutionary time step generates long term evolutionary dynamics of an SDS geno-envo-phenotype.

### 3.4. Fitness in age structured populations

To compute the total selection gradient of the genotype, we now write a fitness function that is more tractable than invasion fitness and that has the same first-order derivatives with respect to mutant trait values as invasion fitness for age-structured populations. To do this, we first write a mutant’s survival probability and fertility at each age. At the resident population dynamics equilibrium, a rare mutant’s fertility at age *a* is

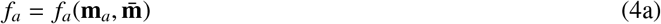

and the mutant’s survival probability from age *a* to *a* + 1 is

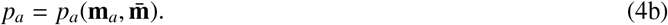

The first argument **m**_*a*_ in Eqs. (4) is the direct dependence of the mutant’s fertility and survival at a given age on her own geno-envo-phenotype at that age. The second argument 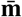 in Eqs. (4) is the direct dependence on social partners’ geno-envo-phenotype at any age (thus, fertility and survival may directly depend on the environment of social partners, specifically, as it may affect the carrying capacity, and fertility and survival are density dependent).

In the SI section S2.3, we show that the gradients of invasion fitness λ with respect to mutant trait values are equal to (not an approximation of) the corresponding gradients of the relative fitness *w* of a mutant individual per unit of generation time (Eq. S2.3.9), defined as

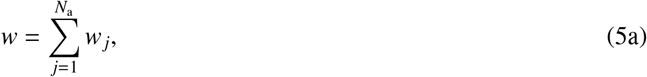

where a mutant’s relative fitness at age *j* is

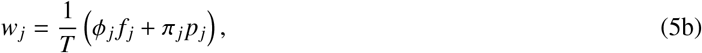

and generation time is

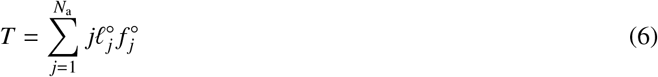

(Charlesworth 1994, Eq. 1.47c; Bulmer 1994, Eq. 25, Ch. 25; Bienvenu and Legendre 2015, Eqs. 5 and 12). The superscript ° denotes evaluation at 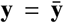 (so at 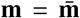 as the resident is a socio-devo equilibrium). The quantity 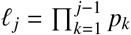 is the survivorship of mutants from age 1 to age *j*, and 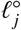 is that of neutral mutants. Thus, the generation time appearing in *w* is for a neutral mutant, or resident, rather than the mutant. This can be intuitively understood as the mutant having to invade over a time scale determined by the resident rather than the mutant. We denote the force of selection on fertility at age *j* (Hamilton 1966 and Caswell 1978, his Eqs. 11 and 12) as

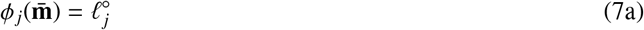

and the force of selection on survival at age *j* (Baudisch 2005, her Eq. 5a) as

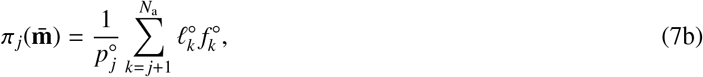

which are independent of mutant trait values because they are evaluated at the resident trait values. It is easily checked that *ϕ* _*j*_ and *π* _*j*_ decrease with *j* (respectively, if 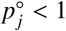 and 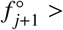 provided that 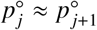).

In the SI section S2.4.2, we show that the gradients of invasion fitness λ with respect to mutant trait values are also equal to 1/*T* times the corresponding gradients of a rare mutant’s expected lifetime reproductive success *R*_0_ (Eq. S2.4.11) (Bulmer, 1994; Caswell, 2009). This occurs because of our assumption that mutants arise when residents are at carrying capacity (Mylius and Diekmann, 1995). For our life cycle, a mutant’s expected lifetime reproductive success is

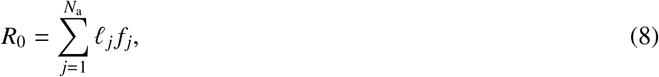

(Caswell, 2001). From the equality of the gradients of invasion fitness and fitness, it follows that invasion fitness λ for age-structured populations is approximately equal to the mutant’s relative fitness *w* to first-order of approximation around resident genotypic traits, that is, λ ≈ *w* (Eq. S2.3.11). Similarly, from the equality of the gradients of invasion fitness and a mutant’s lifetime reproductive success per generation time, it follows that invasion fitness λ for age-structured populations is given by λ ≈ 1 + (*R*_0_ − 1)/*T* to first-order of approximation around resident genotypic traits (Eq. S2.3.13). Taking derivatives of *w* with respect to mutant trait values is generally simpler than for λ or *R*_0_, so we present most results below in terms of *w*.

## 4. Summary of main results

We now give an overview of the three sets of main results. We provide a comprehensive presentation of ancillary results and further analysis in section 5 and SI section S3. The derivations of all these results are in SI section S5 and involve repeated use of the chain rule due to the recurrence and feedbacks involved in the developmental constraint (1).

### 4.1. Total selection gradient of the genotype

The total selection gradient of the genotype is

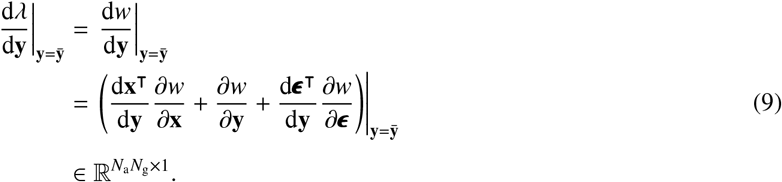

The chain rule in Eq. (9) is reversed to its standard presentation because the gradient in the canonical equation (3) is a column vector. Yet, the order in Eq. (9) gives a natural biological interpretation: a perturbation in the genotype first affects the phenotype and then the affected phenotype affects fitness, and similarly for the effect of genotypic change on the environment. The total selection gradient of the genotype depends on the block matrix of *total effects of a mutant’s genotype on her phenotype*, given by

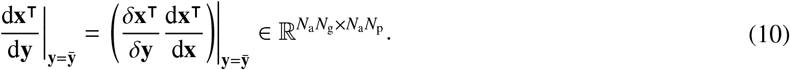

This matrix is a mechanistic counterpart of Fisher’s (1918) additive effects of allelic substitution and of Wagner’s (1984) developmental matrix. This matrix also gives the sensitivity of a recurrence of the form (1) to perturbations in parameters **y** at any time *a*. Moreover, this matrix can be interpreted as measuring the total developmental bias of the phenotype from the genotype (Uller et al., 2018).

The block matrix of *total effects of a mutant’s phenotype on her phenotype* is

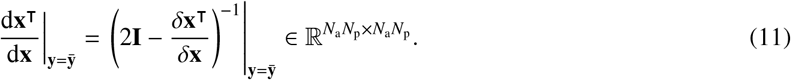

This matrix describes *developmental feedback*, is always invertible (SI section S5.1, Eq. S5.1.15), and gives the sensitivity of a recurrence of the form (1) to perturbations in state variables **x** at any time *a*. Moreover, this matrix can be interpreted as a lifetime collection of total immediate effects of the phenotype on itself. Partitioning this matrix in blocks of size *N*_p_ × *N*_p_, the *a j*-th block entry gives the total effects of the phenotype at age *a* on the phenotype at age *j*, that is,

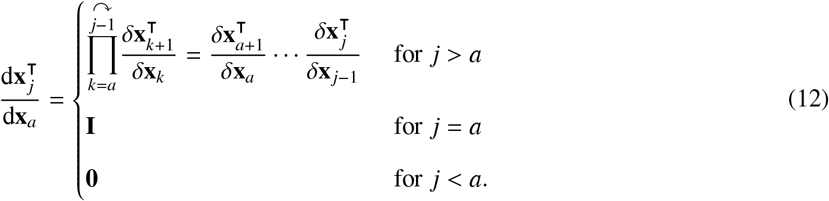

Since matrix multiplication is not commutative, the ↷ denotes right multiplication. Eq. (11) has the same form of an equation for total effects used in path analysis (Greene 1977, p. 380; see also Morrissey 2014, Eq. 2), particularly if 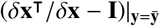 is interpreted as a matrix listing the path coefficients of “direct” effects of the phenotype on itself (if environmental traits are not explicitly considered in the analysis).

The block matrix of *total immediate effects of ζ on a mutant’s phenotype* is

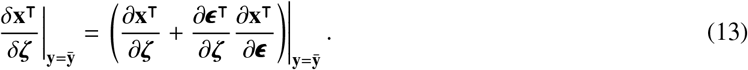

for 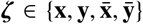. This matrix depends on direct niche construction (∂**ϵ**^⊺^/∂*ζ*) and direct plasticity (∂**x**^⊺^/∂**ϵ**). Thus, the developmental feedback of the phenotype (Eq. 11) depends on direct developmental bias from the phenotype (∂**x**^⊺^/∂**x**), direct niche-construction by the phenotype (∂**ϵ**^⊺^/∂**x**), and direct plasticity of the phenotype (∂**x**^⊺^/∂**ϵ**).

The total selection gradient of the genotype also depends on the block matrix of *total effects of a mutant’s genotype on her environment*

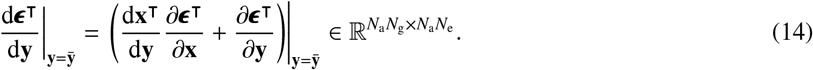

This matrix quantifies the total niche construction by the genotype and depends on direct niche construction by the phenotype and the genotype.

Additionally, the *direct selection gradient of the phenotype* is

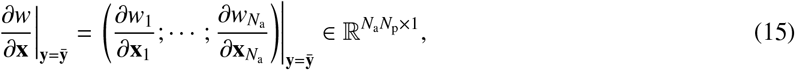

where the direct selection gradient of the phenotype at age *a* ∈ {1, …, *N*_a_} is

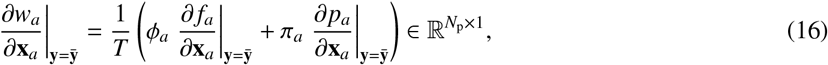

and the direct effect on fertility of the mutant phenotype at age *a* ∈ {1, …, *N*_a_} is

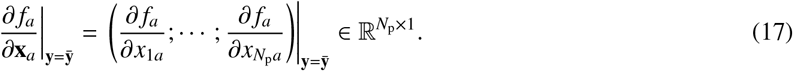

The other direct selection gradients and direct effects on fertility or survival are defined analogously.

In turn, the block matrix of *direct effects of a mutant’s phenotype on her phenotype* is

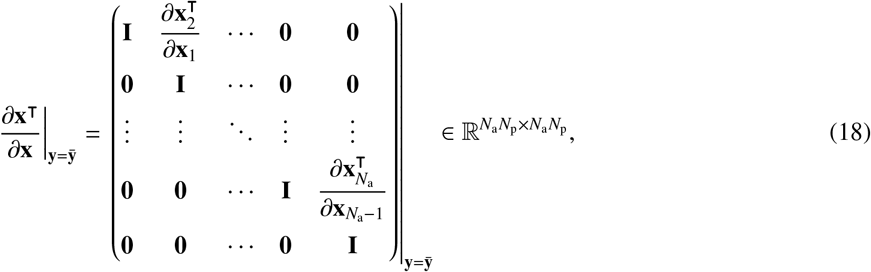

where the matrix of *direct effects of a mutant’s phenotype at age a on her phenotype at age a* + 1 is

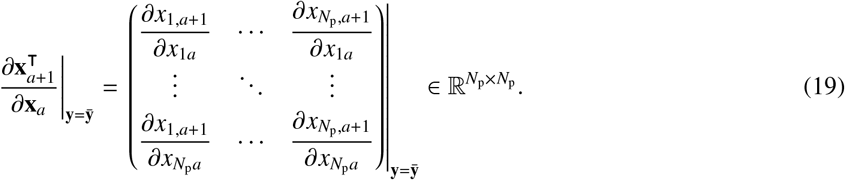

Given the interpretations of d**x**^⊺^/d**y** and d**ϵ**^⊺^/d**y**, the total selection gradient of the genotype (9) can then be interpreted as measuring total (directional) genotypic selection in a fitness landscape modified by: (1) the interaction of total developmental bias of the phenotype from the genotype and directional selection on the phenotype, and (2) the interaction of total niche construction by the genotype and environmental sensitivity of selection ∂*w*/∂**ϵ** (Chevin et al., 2010). In a standard quantitative genetics framework, the total selection gradient of the genotype would correspond to Lande’s (1979) selection gradient of the genotype if phenotypic and environmental traits are not explicitly included in the analysis.

To build a minimal evo-devo dynamics model using our approach, one first needs expressions for fertility *f*_*a*_, survival *p*_*a*_, and development **g**_*a*_. Then, one computes the partial derivatives (17), (19), and analogous partial derivatives, and finally feeds these partial derivatives to Eqs. (9)-(11), (13)-(16), and (18) to compute the evo-evo dynamics using Eqs. (1)-(3).

### 4.2. Evolutionary dynamics of the phenotype in gradient form

The evo-devo dynamics above allow for the modeling of the evolutionary dynamics of the phenotype under explicit development, but do not provide an interpretation of the evolutionary dynamics of the phenotype as the climbing of a fitness landscape. To obtain such an interpretation, we obtain equations in gradient form for the evolutionary dynamics of the phenotype.

As a first step, temporarily assume that the following four conditions hold: (I) development is non-social 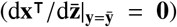, and there is (II) no exogenous plastic response of the phenotype 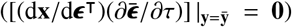, (III) no total immediate selection on the genotype 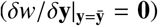, and (IV) no niche-constructed effects of the phenotype on fitness 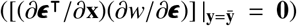. Then, in the limit as Δ*τ* → 0, the evolutionary dynamics of the resident phenotype satisfies

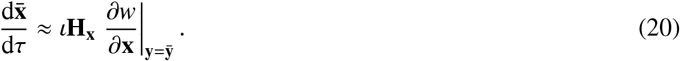

This is a mechanistic version of the Lande equation for the phenotype and is in gradient form, where the *mechanistic additive genetic covariance matrix of the phenotype* 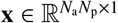 is

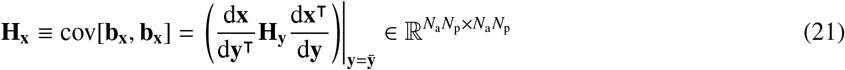

(H for heredity), which guarantees that the developmental constraint (1) is met at all times given the formulas for the total effects of the genotype on phenotype provided in section 4.1. The matrix **H**_**x**_ describes genetic covariation in the phenotype as the covariation of *mechanistic* breeding value **b**_**x**_. Mechanistic breeding value is a mechanistic counterpart of breeding value, defined not in terms of regression coefficients but in terms of total derivatives and so has different properties explained in section 5.1.

Eq. (20) describes phenotypic evolution in gradient form but is dynamically insufficient since it depends on the resident genotype 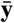 but does not describe its evolutionary dynamics. Indeed, the only reason that the resident phenotype 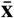 evolves under the assumptions of Eq. (20) is that the resident genotype 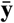 evolves, but the evolution of the resident genotype is not described by such equation. Hence, the mechanistic Lande equation (20) is not sufficient to describe phenotypic evolution as the climbing of a fitness landscape.

### 4.3. Evolutionary dynamics of the geno-envo-phenotype in gradient form

To describe phenotypic evolution as the climbing of a fitness landscape, we obtain dynamically sufficient equations in gradient form for phenotypic evolution. Dropping assumptions (I-IV) above, one such an equation describes the evolutionary dynamics of the resident geno-envo-phenotype as

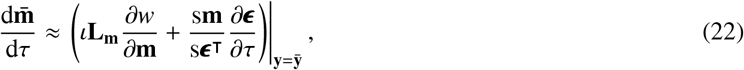

which is an extended mechanistic Lande equation. This equation describes the evolution of the geno-envo-phenotype as the climbing of a fitness landscape and is dynamically sufficient because it describes the evolution of all the variables involved, including the resident genotype 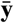. This equation depends on the *mechanistic additive socio-genetic cross-covariance matrix of the geno-envo-phenotype*

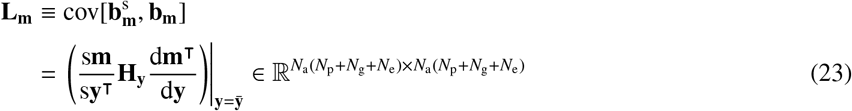

(L for legacy). We say that the matrix **L**_**m**_ describes the “socio-genetic” covariation of the geno-envo-phenotype, that is, the covariation between the stabilized mechanistic breeding value 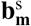 and mechanistic breeding value **b**_**m**_. Whereas mechanistic breeding value considers the total effect of mutations on the traits within the individual, stabilized mechanistic breeding value considers the stabilized effect of mutations on the traits across the population, after social development has stabilized. The matrix **L**_**m**_ may be asymmetric and its main diagonal entries may be negative (unlike variances) due to social development. Moreover, the matrix **L**_**m**_ is always singular because d**m**^⊺^/d**y** has fewer rows than columns, regardless of whether development is social. This means that there are always directions in geno-envo-phenotypic space in which there cannot be socio-genetic covariation, so there are always absolute socio-genetic constraints on adaptation of the geno-envo-phenotype (see SI section S2.2 for a definition of absolute mutational or genetic constraints). In particular, this implies that evolution does not generally stop at peaks of the fitness landscape where the direct selection gradient is zero.

Although the extended mechanistic Lande equation (22) is useful to interpret evolution as the climbing of a fitness landscape, in practice it may often be more useful to compute the evo-devo dynamics using Eqs. (1)-(3) since the extended mechanistic Lande equation may still require computing the evo-devo dynamics, particularly when development is social.

In the remainder of this section, we give the formulas for additional quantities involved in the extended mechanistic Lande equation (22). These formulas show that stabilized effects depend on social feedback (Eq. 27), joint total developmental bias (Eq. 26), and joint direct niche construction (Eqs. 29 and 32). Analysis of the obtained equations for genetic covariation and evolutionary dynamics is given in section 5. A reader interested in seeing an illustration of the method may jump to section 6.

The matrix **L**_**m**_ depends on the block matrix of *total effects of a mutant’s genotype on her geno-envo-phenotype*

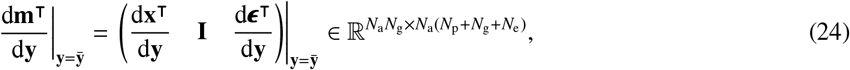

measuring total developmental bias of the geno-envo-phenotype from the genotype, which is singular because it has fewer rows than columns. The matrix **L**_**m**_ also depends on the transpose of the matrix of *stabilized effects of a focal individual’s genotype on the geno-envo-phenotype*

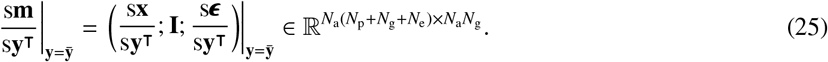

This matrix gives the total effects of the genotype on the geno-envo-phenotype after the effect of the mutation has propagated through the population through social development and such propagation has stabilized. This matrix can be interpreted as measuring stabilized developmental bias of the geno-envo-phenotype from the genotype.

Eq. (25) depends on the transpose of the matrix of *stabilized effects of a focal individual’s phenotype or genotype on her phenotype*

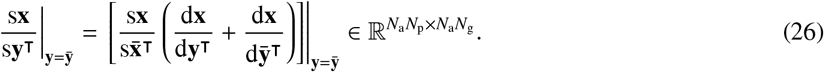

This matrix can be interpreted as measuring stabilized developmental bias of the phenotype from the genotype. It depends on the effect of the genotype of the focal individual and of social partners on the phenotype, and on the feedback of such altered phenotype through the population (social feedback described by 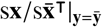). We say that stabilized developmental bias is “joint” as it includes both the effects of the focal individual and of social partners (i.e., 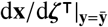 for 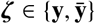). If development is not social (i.e., 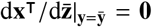), then the stabilized developmental bias matrix of the phenotype from the genotype 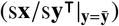 reduces to the corresponding total developmental bias matrix 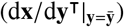.

The matrix in Eq. 26 depends on *social feedback*, given by the transpose of the matrix of *stabilized effects of social partners’ phenotypes on a focal individual’s phenotype*

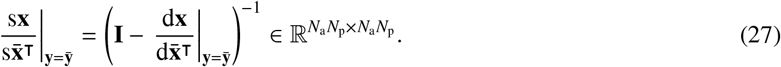

This matrix is invertible by our assumption that all the eigenvalues of 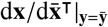 have absolute value strictly less than one, to guarantee that the resident is socio-devo stable. The matrix 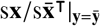 can be interpreted as the total effects of social partners’ phenotypes on a focal individual’s phenotype after socio-devo stabilization; or vice versa, of a focal individual’s phenotype on social partners’ phenotypes. This matrix corresponds to an analogous matrix found in the indirect genetic effects literature (Moore et al., 1997, Eq. 19b and subsequent text). If development is not social from the phenotype (i.e., 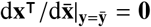), then the matrix 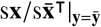 is the identity matrix. This is the only stabilized-effect matrix that does not reduce to the corresponding total-effect matrix when development is not social.

The transpose of the block matrix of *total effects of social partners’ phenotype on a mutant’s phenotype* is

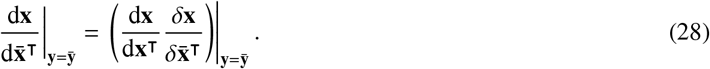

This matrix can be interpreted as measuring total social developmental bias of the phenotype from phenotype, as well as the total effects on the phenotype of extra-genetic inheritance. It is also a mechanistic version of the matrix of interaction coefficients in the indirect genetic effects literature (i.e., Ψ in Eq. 17 of Moore et al. 1997, which is defined as a matrix of regression coefficients). Moreover, the matrix 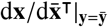can be interpreted as involving a developmentally immediate pulse of phenotype change caused by a change in social partners’ traits followed by the triggered developmental feedback of the mutant’s phenotype.

Eq. (25) also depends on the transpose of the matrix of *stabilized effects of a focal individual’s genotype on the environment*

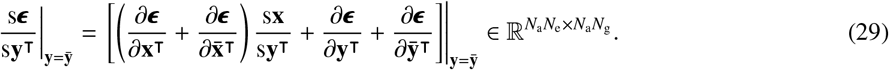

This matrix can be interpreted as measuring stabilized niche construction by the genotype. This matrix is formed by stabilized developmental bias of the geno-phenotype from genotype followed by joint direct niche construction by the geno-phenotype (Layer 5, Eq. S6a). If development is not social (i.e., 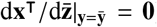), then stabilized niche construction by the genotype 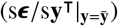 reduces to total niche construction by the genotype 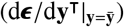.

The extended mechanistic Lande equation (22) also depends on the transpose of the matrix of *stabilized effects of a focal individual’s environment on the geno-envo-phenotype*

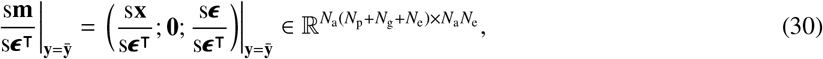

measuring stabilized plasticity of the geno-envo-phenotype.

The transpose of the matrix of *stabilized effects of a focal individual’s environment on the phenotype* is

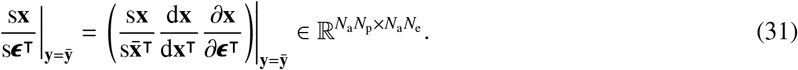

This matrix measures the stabilized plasticity of the phenotype, where the environment first causes direct plasticity in a focal individual, and then there is developmental and social feedback. In contrast to Eq. (26), stabilized plasticity does not depend on the joint effects of the environment. If development is not social (i.e., 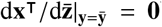, then stabilized plasticity reduces to total plasticity.

*Stabilized environmental feedback* is given by the transpose of the matrix of *stabilized effects of a focal individual’s environment on the environment*

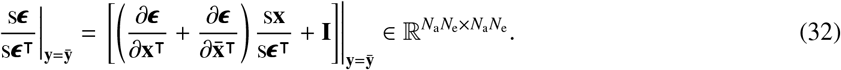

This matrix depends on stabilized plasticity of the phenotype, joint direct niche construction by the phenotype, and direct mutual environmental dependence. Such direct mutual environmental dependence is here the identity matrix because of our assumption that environmental traits are mutually independent.

## 5. Top layers of the evo-devo process

The equations listed in section 4 are part of a broader set of equations that we derive to analyze the evolutionary dynamics under explicit development. We term “evo-devo process” this set of equations. The evo-devo process can be arranged in a layered structure, where each layer is formed by components in layers below (Fig. 4). This layered structure helps see how complex interactions between variables involved in genetic covariation are formed by building blocks describing the direct interaction between variables. In this section 5, we present and analyze the top two layers of the evo-devo process, namely, the layer of genetic covariation and the layer of evolutionary dynamics (Fig. 4A-D). In SI section S3, we describe and analyze the bottom layers, starting from the layer of elementary components up to the layer of stabilized effects (Fig. 4E-I). Sections 5, S3, and S5 include multiple ancillary results. For instance, SI section S3.4 includes closed-form equations for the total selection gradients of the genotype, phenotype, environment, geno-phenotype, and geno-envo-phenotype, and alternative arrangements thereof.

**Figure 4.**
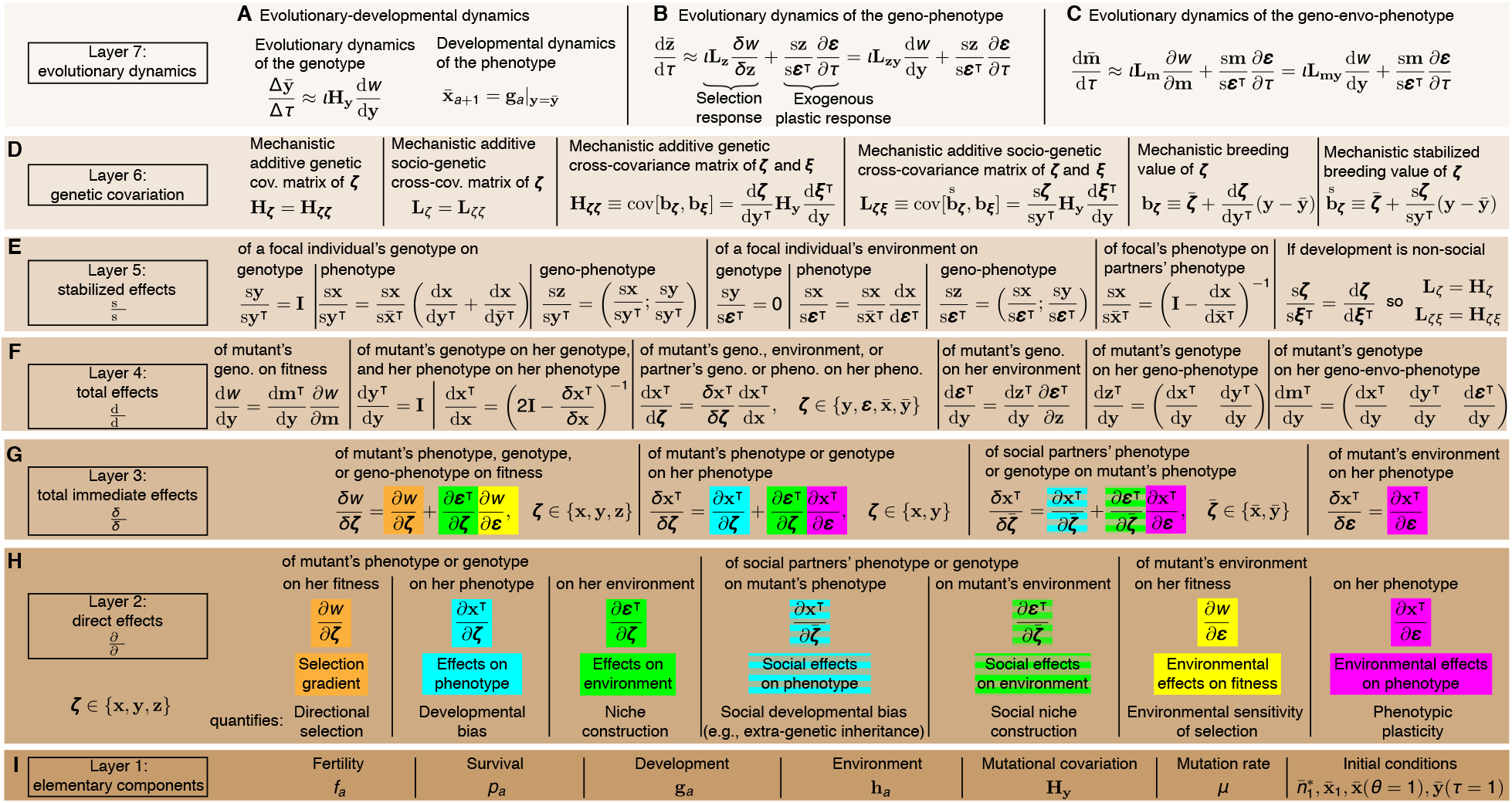
The evo-devo process and its layered structure. Here we summarize the equations composing the evo-devo process arranged in a layered structure. Each layer is formed by components in layers below. Layer 7 describes the evolutionary dynamics as (A) evo-devo dynamics, which in the limit as Δ*τ* → 0 implies (B) the evolutionary dynamics of the geno-phenotype, and (C) the evolutionary dynamics of the geno-envo-phenotype.(D) Layer 6 describes genetic covariation. (E) Layer 5 describes stabilized effects (total derivatives over life after socio-devo stabilization, denoted by s/s). (F) Layer 4 describes total effects (total derivatives over life before socio-devo stabilization, denoted by d/d). (G) Layer 3 describes total immediate effects (total derivatives at the current age, denoted by *δ/δ*). (H) Layer 2 describes direct effects (partial derivatives, denoted by ∂/∂).(I)Layer 1 comprises the elementary components of the evo-devo process that generate all layers above. All derivatives are evaluated at 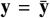. See SI section S3.2 for the equations of direct-effect matrices, which have structure due to age structure. See Fig. 1 and Table 1 for the meaning of symbols.

### 5.1. Layer 6: genetic covariation

In this section, we describe and analyze the layer of genetic covariation of the evo-devo process. We first define mechanistic breeding value under our adaptive dynamics assumptions, which allows us to define mechanistic additive genetic covariance matrices under our assumptions. Then, we define (socio-devo) stabilized mechanistic breeding value, which we use to define mechanistic additive socio-genetic cross-covariance matrices. The notions of stabilized mechanistic breeding values and mechanistic socio-genetic cross-covariance generalize the corresponding notions of mechanistic breeding value and mechanistic additive genetic covariance to consider the effects of social development.

We follow the standard definition of breeding value to define its mechanistic analogue under our assumptions. The breeding value of a trait is defined under quantitative genetics assumptions as the best linear estimate of the trait from gene content (Lynch and Walsh, 1998; Walsh and Lynch, 2018). As stated above, under quantitative genetics assumptions, the *i*-th trait value *x*_*i*_ is written as 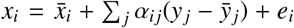 where the overbar denotes population average, *y* _*j*_ is the *j*-th predictor (gene content in *j*-th locus), α_*i j*_ is the partial least-square regression coefficient of 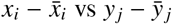, and *e*_*i*_ is the residual; the breeding value of *x*_*i*_ is 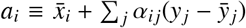. Accordingly, we define the mechanistic breeding value **b**_*ζ*_ of a vector *ζ* as its first-order estimate with respect to genotypic traits **y** around the resident genotypic traits 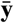:

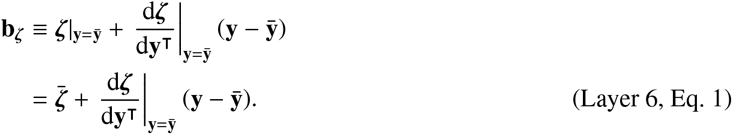

The key difference of this definition with that of breeding value is that rather than using regression coefficients, this definition uses the total derivatives of *ζ* with respect to the genotype, 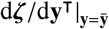, which are a mechanistic analogue of Fisher’s additive effect of allelic substitution (his α; see Eq. I of Fisher 1918 and p. 72 of Lynch and Walsh 1998).

There are material differences between breeding value and its mechanistic counterpart, as made evident with heritability. Because breeding value under quantitative genetics uses linear regression via least squares, breeding value *a*_*i*_ is guaranteed to be uncorrelated with the residual *e*_*i*_. This guarantees that heritability is between zero and one. Indeed, the (narrow sense) heritability of trait *x*_*i*_ is defined as *h*^2^ = var[*a*_*i*_]/var[*x*_*i*_], where using *x*_*i*_ = *a*_*i*_ + *e*_*i*_ we have var[*x*_*i*_] = var[*a*_*i*_] + var[*e*_*i*_] + 2cov[*a*_*i*_, *e*_*i*_]. The latter covariance is zero due to least squares, and so *h*^2^ ∈ [0, 1]. In contrast, mechanistic breeding values may be correlated with residuals. Indeed, in our framework we have that the phenotype 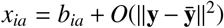, but mechanistic breeding value *b*_*ia*_ is not computed via least squares, so *b*_*ia*_ and the error 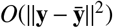 may covary, positively or negatively. Hence, the classic quantitative genetics partition of phenotypic variance into genetic and “environmental” (i.e., residual) variance does not hold with mechanistic breeding value, as there may be mechanistic genetic and “environmental” covariance. Consequently, since the covariance between two random variables is bounded from below by the negative of the product of their standard deviations, mechanistic heritability defined as the ratio between the variance of mechanistic breeding value and phenotypic variance cannot be negative but it may be greater than one.

Our definition of mechanistic breeding value recovers Fisher’s (1918) infinitesimal model under certain conditions, but we do not assume the infinitesimal model. According to Fisher’s (1918) infinitesimal model, the normalized breeding value excess is normally distributed as the number of loci approaches infinity. Using Layer 6, Eq. 1, we have that the mechanistic breeding value excess for the *i*-th entry of **b**_*ζ*_ is

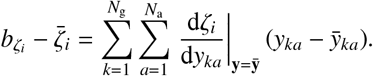

Let us denote the mutational variance for the *k*-th genotypic trait at age *a* by

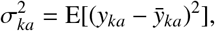

and let us denote the total mutational variance by

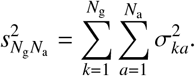

If the *y*_*ka*_ are mutually independent and Lyapunov’s condition is satisfied, from the Lyapunov central limit theorem we have that, as either the number of genotypic traits *N*_g_ or the number of ages *N*_a_ tends to infinity (e.g., by reducing the age bin size), the normalized mechanistic breeding value excess

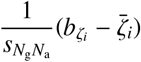

is normally distributed with mean zero and variance 1. Thus, this limit yields Fisher’s (1918) infinitesimal model, although we do not assume such limit. Our framework thus recovers the infinitesimal model as a particular case, when either *N*_g_ or *N*_a_ approaches infinity (provided that the *y*_*ka*_ are mutually independent and Lyapunov’s condition holds).

From our definition of mechanistic breeding value, we have that the mechanistic breeding value of the genotype is the genotype itself. Indeed, the mechanistic breeding value of the genotype **y** is

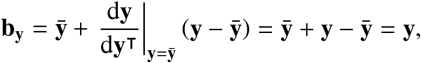

since 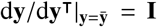 because, by assumption, the genotype does not have developmental constraints and is developmentally independent (Layer 4, Eq. S5). In turn, from Layer 6, Eq. 1, the expected mechanistic breeding value of vector *ζ* is

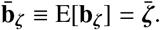

We now define mechanistic additive genetic covariance matrices under our assumptions. The additive genetic variance of a trait is defined under quantitative genetics assumptions as the variance of its breeding value, which is extended to the multivariate case so the additive genetic covariance matrix of a trait vector is the covariance matrix of the traits’ breeding values (Lynch and Walsh, 1998; Walsh and Lynch, 2018). Accordingly, we define the *mechanistic additive genetic covariance matrix* of a vector *ζ* ∈ ℝ^*m*×1^ as the covariance matrix of its mechanistic breeding value:

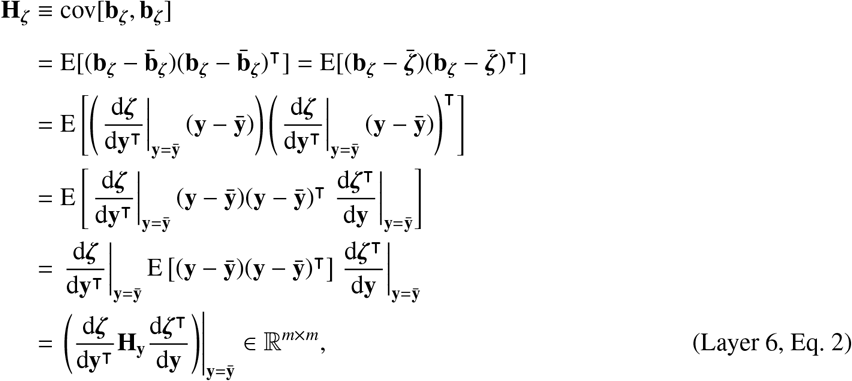

where the fourth line follows from the property of the transpose of a product (i.e., (**AB**)^⊺^ = **B**^⊺^**A**^⊺^) and the last line follows since the mechanistic additive genetic covariance matrix of the genotype **y** is

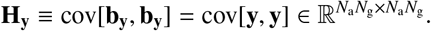

Layer 6, Eq. 2 has the same form of previous expressions for the additive genetic covariance matrix under quantitative genetics assumptions, although using least-square regression coefficients in place of the derivatives if the classic partitioning of phenotypic variance is to hold (see Eq. II of Fisher 1918, Eq. + of Wagner 1984, Eq. 3.5b of Barton and Turelli 1987, and Eq. 4.23b of Lynch and Walsh 1998; see also Eq. 22a of Lande 1980, Eq. 3 of Wagner 1989, and Eq. 9 of Charlesworth 1990). We denote the matrix **H** rather than **G** to note that the two are different, particularly as the former is based on mechanistic breeding value. Note **H**_*ζ*_ is symmetric.

In some cases, Layer 6, Eq. 2 allows one to immediately determine whether a mechanistic additive genetic covariance matrix is singular, which means there are directions in the matrix’s space in which there is no genetic variation. Indeed, a matrix with fewer rows than columns is always singular (Horn and Johnson, 2013, section 0.5 second line), and if the product **AB** is well-defined and **B** is singular, then **AB** is singular (this is easily checked to hold). Hence, from Layer 6, Eq. 2 it follows that **H**_*ζ*_ is necessarily singular if d*ζ*^⊺^/d**y** has fewer rows than columns, that is, if **y** has fewer entries than *ζ*. Since **y** has *N*_a_*N*_g_ entries and *ζ* has *m* entries, then **H**_*ζ*_ is singular if *N*_a_*N*_g_ < *m*. Moreover, Layer 6, Eq. 2 allows one to immediately identify bounds for the “degrees of freedom” of genetic covariation, that is, for the rank of **H**_*ζ*_. Indeed, for a matrix **A** ∈ ℝ^*m*×*n*^, we have that the rank of **A** is at most the smallest value of *m* and *n*, that is, rank(**A**) ≤ min{*m, n*} (Horn and Johnson, 2013, section 0.4.5 (a)). Furthermore, from the Frobenius inequality (Horn and Johnson, 2013, section 0.4.5 (e)), for a well-defined product **AB**, we have that rank(**AB**) ≤ rank(**B**). Therefore, for *ζ* ∈ ℝ^*m*×1^, we have that

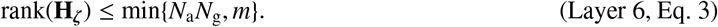

Intuitively, this states that the degrees of freedom of genetic covariation are at most given by the lifetime number of genotypic traits (i.e., *N*_a_*N*_g_). So if there are more entries in *ζ* than there are lifetime genotypic traits, then there are fewer degrees of freedom of genetic covariation than traits. This point is mathematically trivial but biologically crucial because the evolutionary dynamic equations in gradient form that are generally dynamically sufficient involve an **H**_*ζ*_ whose *ζ* necessarily has more entries than **y**. These points on the singularity and rank of **H**_*ζ*_ also hold under quantitative genetics assumptions, where the same structure (Layer 6, Eq. 2) holds, except that **H**_**y**_ does not refer to mutational variation but to standing variation in allele frequency and total effects are measured with regression coefficients. Considering standing variation in **H**_**y**_ and regression coefficients does not affect the points made in this paragraph.

Consider the following slight generalization of the mechanistic additive genetic covariance matrix. We define the mechanistic additive genetic cross-covariance matrix between a vector *ζ* ∈ R^*m*×1^ and a vector *ξ* ∈ R^*n*×1^ as the cross-covariance matrix of their mechanistic breeding value:

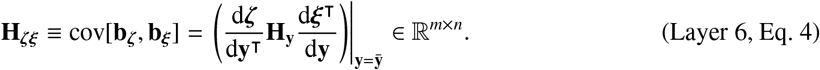

Thus, **H**_*ζζ*_ = **H**_*ζ*_. Note **H**_*ζξ*_ may be rectangular, and if square, asymmetric. Again, from Layer 6, Eq. 4 it follows that **H**_*ζξ*_ is necessarily singular if there are fewer entries in **y** than in *ξ* (i.e., if *N*_a_*N*_g_ < *n*). Also, for *ξ* ∈ ℝ^*n*×1^, have that

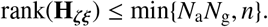

In words, the degrees of freedom of genetic cross-covariation are at most given by the lifetime number of genotypic traits.

The mechanistic additive genetic covariance matrix of the phenotype takes the following form. Evaluating Layer 6, Eq. 2 at *ζ* = **x**, the mechanistic additive genetic covariance matrix of the phenotype 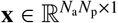 is

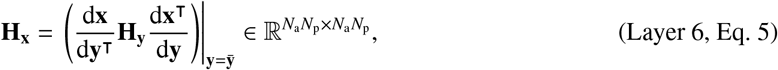

which is singular because the developmental matrix 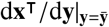 is singular since the developmentally initial phenotype is not affected by the genotype and the developmentally final genotypic traits do not affect the phenotype (SI section S5.2, Eq. S5.2.16). This singularity is immaterial for our purposes because a dynamical system consisting only of evolutionary dynamic equations for the phenotype thus having an associated **H**_**x**_-matrix is underdetermined in general because the system has fewer dynamic equations (i.e., the number of entries in **x**) than dynamic variables (i.e., the number of entries in (**x**; **y**; **ϵ**)). Indeed, the evolutionary dynamics of the phenotype generally depends on the resident genotype, in particular, because the developmental matrix depends on the resident genotype (Layer 4, Eq. S2), for instance, due to non-linearities in the developmental map involving products between genotypic traits, or between genotypic traits and phenotypes, or between genotypic traits and environmental traits, that is, gene-gene interaction, gene-phenotype interaction, and gene-environment interaction, respectively. Thus, evolutionary dynamic equations of the phenotype alone generally have either zero or an infinite number of solutions for any given initial condition and are thus dynamically insufficient. To have a determined system in gradient form that is dynamically sufficient in general, we follow the evolutionary dynamics of both the phenotype and the genotype, that is, of the geno-phenotype, which depends on **H**_**z**_ rather than **H**_**x**_ alone.

The mechanistic additive genetic covariance matrix of the geno-phenotype takes the following form. Evaluating Layer 6, Eq. 2 at *ζ* = **z**, the mechanistic additive genetic covariance matrix of the geno-phenotype 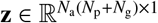 is

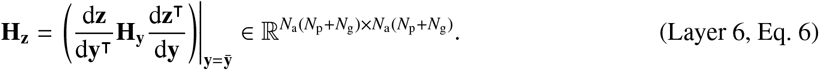

This matrix is necessarily singular because the geno-phenotype **z** includes the genotype **y** so d**z**^⊺^/d**y** has fewer rows than columns (Layer 4, Eq. S7). Intuitively, Layer 6, Eq. 6 has this form because the phenotype is related to the genotype by the developmental constraint (1). From Layer 6, Eq. 3, the rank of **H**_**z**_ has an upper bound given by the number of genotypic traits across life (i.e., *N*_a_*N*_g_), so **H**_**z**_ has at least *N*_a_*N*_p_ eigenvalues that are exactly zero. Thus, **H**_**z**_ is singular if there is at least one trait that is developmentally constructed according to the developmental constraint (1) (i.e., if *N*_p_ > 0). This is a mathematically trivial singularity, but it is biologically key because it is **H**_**z**_ rather than **H**_**x**_ that occurs in a generally dynamically sufficient evolutionary system in gradient form (provided the environment is constant; if the environment is not constant, the relevant matrix is **H**_**m**_ which is also always singular if there is at least one phenotype or one environmental trait).

Another way to see the singularity of **H**_**z**_ is the following. From Layer 6, Eq. 6, we can write the mechanistic additive genetic covariance matrix of the geno-phenotype as

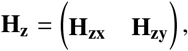

where the mechanistic additive genetic cross-covariance matrix between **z** and **x** is

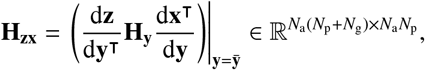

and the mechanistic additive genetic cross-covariance matrix between **z** and **y** is

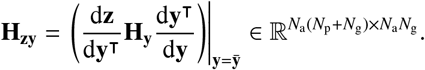

Thus, using Layer 4, Eq. S5, we have that

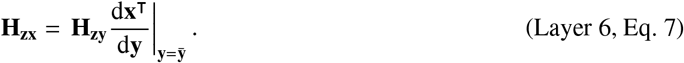

That is, some columns of **H**_**z**_ (i.e., those in **H**_**zx**_) are linear combinations of other columns of **H**_**z**_ (i.e., those in **H**_**zy**_). Hence, **H**_**z**_ is singular.

The mechanistic additive genetic covariance matrix of the geno-phenotype is singular because the geno-phenotype includes the genotype (“gene content”). The singularity arises because the mechanistic breeding value of the pheno-type is a linear combination of the mechanistic breeding value of the genotype by definition of mechanistic breeding value, regardless of whether the phenotype is a linear function of the genotype and regardless of the number of phenotypic or genotypic traits. In quantitative genetics terms, the **G**-matrix is a function of allele frequencies (which corresponds to our 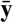), so a generally dynamically sufficient Lande system would require that allele frequencies are part of the dynamic variables considered; consequently, if the geno-phenotypic vector 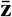 includes allele frequencies 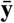, then the associated **G** is necessarily singular since by definition, breeding value under quantitative genetics assumptions is a linear combination of gene content.

The definition of mechanistic breeding value implies that if there is only one phenotype and one genotypic trait, with a single age each, then there is a perfect correlation between their mechanistic breeding values (i.e., their correlation coefficient is 1). This also holds under quantitative genetics assumptions (Via and Lande, 1985), in which case the breeding value *a* of a trait *x* is a linear combination of a single predictor *y*, so the breeding value *a* and predictor *y* are perfectly correlated (i.e., 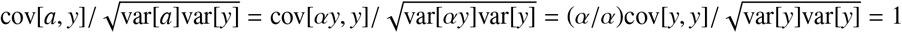). The perfect correlation between a single breeding value and a single predictor arises because, by definition, breeding value excludes the residual *e*. Note this does not mean that the phenotype and genotype are linearly related: it is (mechanistic) breeding values and the genotype that are linearly related by definition of (mechanistic) breeding value (Layer 6, Eq. 1).

A standard approach to remove the singularity of an additive genetic covariance matrix is to remove some traits from the analysis (Lande, 1979). To remove the singularity of **H**_**z**_ we would need to remove at least either all phenotypic traits or all genotypic traits from the analysis. However, removing all phenotypic traits from the analysis prevents analyzing phenotypic evolution as the climbing of a fitness landscape whereas removing all genotypic traits from the analysis renders the analysis dynamically insufficient in general because the evolutionary dynamics of some variables is not described. Thus, in general, to analyze a dynamically sufficient description of phenotypic evolution as the climbing of a fitness landscape, we must keep the singularity of **H**_**z**_.

We now extend the notion of mechanistic breeding value to consider social development. As stated above, we denote a matrix of stabilized effects as s*ζ*^⊺^/s*ξ*, which gives the total effects of *ξ* on *ζ*^⊺^ over the individual’s development and after the effects of the perturbation have stabilized in the population. We define the stabilized mechanistic breeding value of a vector *ζ* as:

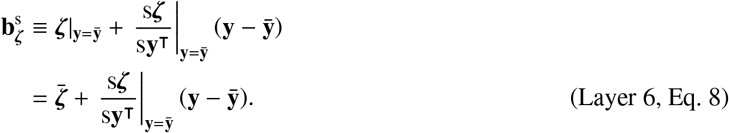

The stabilized-effect matrix 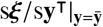 equals the total-effect matrix 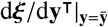 if development is non-social (SI section S3.5). Thus, if development is non-social, the stabilized mechanistic breeding value 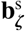 equals the mechanistic breeding value **b**_*ζ*_. Also, note that 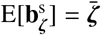.

We use this definition to extend the notion of mechanistic additive genetic covariance matrix to include the effects of social development as follows. We define the *mechanistic additive socio-genetic cross-covariance matrix of ζ* ∈ ℝ^*m*×1^ as

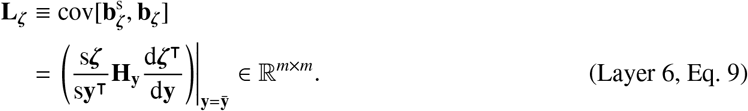

Note **L**_*ζ*_ may be asymmetric and its main diagonal entries may be negative (unlike variances). If development is non-social, **L**_*ζ*_ equals **H**_*ζ*_. As before, **L**_*ζ*_ is singular if *ζ* has fewer entries than **y**. Also, for *ζ* ∈ ℝ^*m*×1^, have that

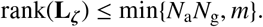

That is, the degrees of freedom of socio-genetic covariation are at most also given by the lifetime number of genotypic traits.

As before, we generalize this notion and define the *mechanistic additive socio-genetic cross-covariance matrix between ζ* ∈ ℝ^*m*×1^ *and ξ* ∈ ℝ^*n*×1^ as

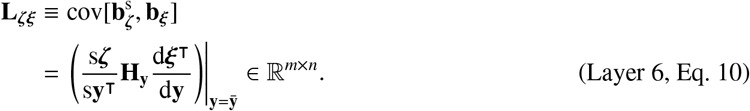

Again, if development is non-social, **L**_*ζξ*_ equals **H**_*ζξ*_. Note **L**_*ζξ*_ may be rectangular and, if square, asymmetric. Also, **L**_*ζ*ξ_ is singular if *ξ* has fewer entries than **y**. For *ξ* ∈ ℝ^*n*×1^, have that

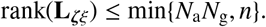

That is, the degrees of freedom of socio-genetic cross-covariation are at most still given by the lifetime number of genotypic traits.

In particular, some **L**_*ζ*ξ_ matrices are singular or not as follows. The mechanistic additive socio-genetic cross-covariance matrix between *ζ* and the geno-phenotype z

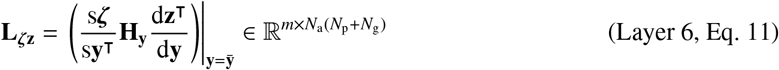

is singular if there is at least one phenotype (i.e., if *N*_p_ > 0). Thus, **L**_*ζ***z**_ has at least *N*_a_*N*_p_ eigenvalues that are exactly zero. Also, the mechanistic additive socio-genetic cross-covariance matrix between *ζ* and the geno-envo-phenotype **m**

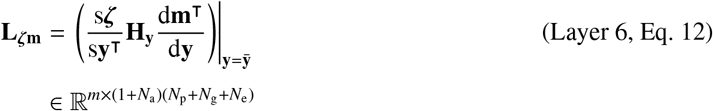

is singular if there is at least one phenotype or one environmental trait (i.e., if *N*_p_ > 0 or *N*_e_ > 0). Thus, **L**_*ζ***m**_ has at least *N*_a_(*N*_p_ + *N*_e_) eigenvalues that are exactly zero. In important contrast, the mechanistic additive socio-genetic cross-covariance matrix between a vector *ζ* ∈ {**y, z, m**} and the genotype **y**

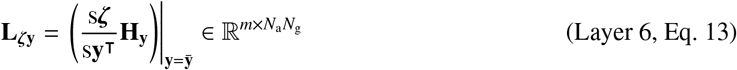

is non-singular if **H**_**y**_ is non-singular because the genotype is developmentally independent (SI sections S5.7 and S5.9). The **L**-matrices share various properties with similar generalizations of the **G**-matrix arising in the indirect genetic effects literature (Kirkpatrick and Lande, 1989; Moore et al., 1997; Townley and Ezard, 2013).

### 5.2. Layer 7: evolutionary dynamics

Now we describe the top layer of the evo-devo process, that of the evolutionary dynamics. This layer contains equations describing the evolutionary dynamics under explicit developmental and environmental constraints. In the SI sections S2.5 and S5.6-S5.9, we show that, in the limit as Δ*τ* → 0, the evolutionary dynamics of the resident phenotype, genotype, geno-phenotype, environment, and geno-envo-phenotype (i.e., for *ζ* ∈ {**x, y, z, ϵ, m**}) are given by

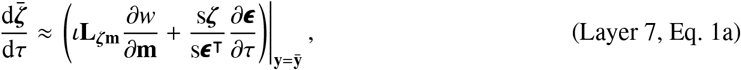

which must satisfy both the developmental constraint

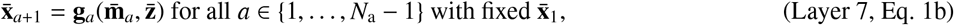

and the environmental constraint

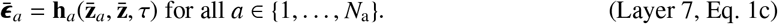

If *ζ* = **z** in Layer 7, Eq. 1a, then the equations in Layers 2-6 guarantee that the developmental constraint is satisfied for all τ > τ_1_ given that it is satisfied at the initial evolutionary time τ_1_. If *ζ* = **m** in Layer 7, Eq. 1a, then the equations in Layers 2-6 guarantee that both the developmental and environmental constraints are satisfied for all τ > τ_1_ given that they are satisfied at the initial evolutionary time τ_1_. Both the developmental and environmental constraints can evolve as the genotype, phenotype, and environment evolve and such constraints can involve any family of curves as long as they are continuously differentiable.

Importantly, although Layer 7, Eq. 1a describes the evolutionary dynamics of *ζ*, such equation is guaranteed to be dynamically sufficient only for certain *ζ*. Layer 7, Eq. 1a is dynamically sufficient if *ζ* is the genotype **y**, the geno-phenotype **z**, or the geno-envo-phenotype **m**, provided that the developmental and environmental constrains are satisfied throughout. In contrast, Layer 7, Eq. 1a is dynamically insufficient if *ζ* is the phenotype **x** or the environment ***ϵ***, because the evolution of the genotype is not followed but it generally affects the system.

Layer 7, Eq. 1a describes the evolutionary dynamics as consisting of selection response and exogenous plastic response. Layer 7, Eq. 1a contains the term

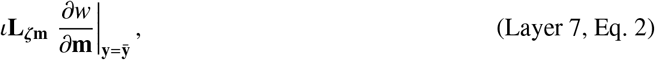

which comprises direct directional selection on the geno-envo-phenotype 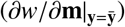, socio-genetic cross-covariation between *ζ* and the geno-envo-phenotype (**L**_*ζ***m**_), and mutational input (ι). The term in Layer 7, Eq. 2 is the *selection response* of 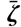 and is a mechanistic generalization of Lande’s (1979) generalization of the univariate breeder’s equation (Lush, 1937; Walsh and Lynch, 2018). Additionally, Layer 7, Eq. 1a contains the term

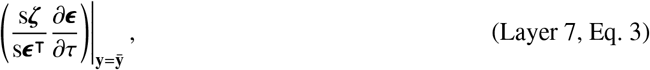

which comprises the vector of environmental change due to exogenous causes 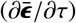 and the matrix of stabilized plasticity 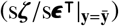. The term in Layer 7, Eq. 3 is the *exogenous plastic response* of 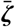 and is a mechanistic generalization of previous expressions (cf. Eq. A3 of Chevin et al. 2010). Note that the *endogenous* plastic response of 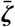 (i.e., the plastic response due to endogenous environmental change arising from niche construction) is part of both the selection response and the exogenous plastic response (Layers 2-6).

Selection response is relatively incompletely described by direct directional selection on the geno-envo-phenotype. Indeed, we saw that the matrix **L**_*ζ***m**_ is always singular if there is at least one phenotype or one environmental trait (Layer 6, Eq. 12). Consequently, the selection response of 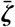 can invariably vanish with persistent direct directional selection on the geno-envo-phenotype.

Selection response is also relatively incompletely described by total immediate selection on the geno-phenotype. We can rewrite the selection response, so the evolutionary dynamics of 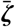for *ζ* ∈ {**x, y, z, ϵ, m**} (Layer 7, Eq. 1a) is equivalently given by

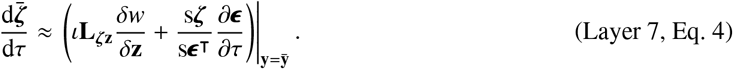

This equation now depends on total immediate selection on the geno-phenotype 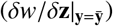, which measures total immediate directional selection on the geno-phenotype (or in a quantitative genetics framework, it is Lande’s (1979) selection gradient of the allele frequency and phenotype if environmental traits are not explicitly included in the analysis). The total immediate selection gradient of the geno-phenotype can be interpreted as pointing in the direction of steepest ascent on the fitness landscape in geno-phenotype space after the landscape is modified by the interaction of direct niche construction and environmental sensitivity of selection (Layer 3, Eq. S1). We saw that the matrix **L**_*ζ***z**_ is always singular if there is at least one phenotype (Layer 6, Eq. 11). Consequently, the selection response of 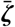 can invariably vanish with persistent directional selection on the geno-phenotype after niche construction has modified the geno-phenotype’s fitness landscape.

In contrast, selection response is relatively completely described by total genotypic selection. We can further rewrite the selection response, so the evolutionary dynamics of 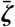 for *ζ* ∈ {**x, y, z, ϵ, m**} (Layer 7, Eq. 1a) is equivalently given by

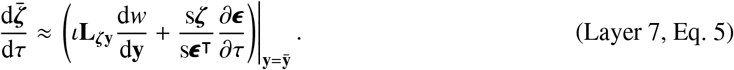

This equation now depends on total genotypic selection 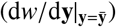, which measures total directional selection on the genotype considering downstream developmental effects (or in a quantitative genetics framework, it is Lande’s (1979) selection gradient of allele frequency if neither the phenotype nor environmental traits are explicitly included in the analysis). In Eq. 9 we saw that the total selection gradient of the genotype can be interpreted as pointing in the direction of steepest ascent on the fitness landscape in genotype space after the landscape is modified by: (1) the interaction of total developmental bias from the genotype and directional selection on the phenotype and (2) the interaction of total niche construction by the genotype and environmental sensitivity of selection. In contrast to the other arrangements of selection response, in SI sections S5.7 and S5.9 we show that **L**_*ζ***y**_ is non-singular for all *ζ* ∈ {**y, z, m**} if **H**_**y**_ is non-singular (i.e., if there is mutational variation in all directions of genotype space); this non-singularity of **L**_*ζ***y**_ arises because genotypic traits are developmentally independent by assumption. Consequently, the selection response of the genotype, geno-phenotype, or geno-envo-phenotype can only vanish when total genotypic selection vanishes if there is mutational variation in all directions of genotype space.

Let us revisit why following the evolutionary dynamics of the phenotype alone is dynamically insufficient in order to compare such an approach with a dynamically sufficient extension. As stated above, Layer 7, Eq. 1a and its equivalents are generally dynamically insufficient if only the evolutionary dynamics of the phenotype are considered (i.e., if *ζ* = **x**). Let us temporarily assume that the following four conditions hold: (I) development is non-social 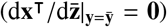, and there is (II) no exogenous plastic response of the phenotype 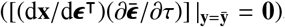, (III) no total immediate selection on the genotype 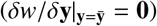, and (IV) no niche-constructed effects of the phenotype on fitness 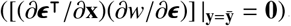. Then, the evolutionary dynamics of the phenotype reduces to

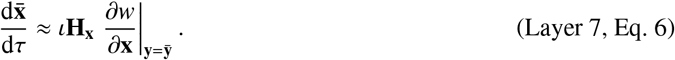

This is a mechanistic version of the Lande equation for the phenotype. The mechanistic additive genetic covariance matrix of the phenotype (Layer 6, Eq. 5) in this equation is singular because the developmentally initial phenotype is not affected by the genotype and the developmentally final genotypic traits do not affect the phenotype (so 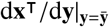 has rows and columns that are zero; SI section S5.2, Eq. S5.2.16). This singularity might disappear by removing from the analysis the developmentally initial phenotype and developmentally final genotypic traits, provided additional conditions hold. Yet, the key point here is that a system describing the evolutionary dynamics of the phenotype alone is dynamically insufficient because such system depends on the resident genotype whose evolution must also be followed. In particular, setting 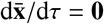does not generally imply an evolutionary equilibrium, or evolutionary stasis, but only an evolutionary nullcline in the phenotype, that is, a transient lack of evolutionary change in the phenotype. To guarantee a dynamically sufficient description of the evolutionary dynamics of the phenotype, we simultaneously consider the evolutionary dynamics of the phenotype and genotype, that is, the geno-phenotype.

A dynamically sufficient system can be obtained by describing the dynamics of the geno-phenotype alone if the environment is constant or has no evolutionary effect. Let us now assume that the following three conditions hold: (i) development is non-social 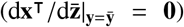, and there is (ii) no exogenous plastic response of the phenotype 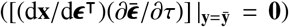, and (iii) no niche-constructed effects of the geno-phenotype on fitness 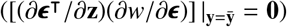. Then, the evolutionary dynamics of the geno-phenotype reduces to

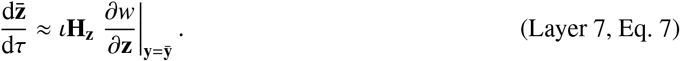

This is an extension of the mechanistic version of the Lande equation to consider the geno-phenotype. The mechanistic additive genetic covariance matrix of the geno-phenotype (Layer 6, Eq. 6) in this equation is singular because the geno-phenotype **z** includes the genotype **y** (so d**z**^⊺^/d**y** has fewer rows than columns; Layer 4, Eq. S7). Hence, the degrees of freedom of genetic covariation in geno-phenotype space are at most given by the number of lifetime genotypic traits, so these degrees of freedom are bounded by genotypic space in a necessarily larger geno-phenotype space. Thus, **H**_**z**_ is singular if there is at least one trait that is developmentally constructed according to the developmental map (Layer 7, Eq. 1b). The evolutionary dynamics of the geno-phenotype is now fully determined by Layer 7, Eq. 7 provided that i-iii hold and that the developmental (Layer 7, Eq. 1b) and environmental (Layer 7, Eq. 1c) constraints are met, which **H**_**z**_ guarantees they are if they are met at the initial evolutionary time τ_1_. In such case, setting 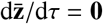 does imply an evolutionary equilibrium to first order of approximation, but this does not imply absence of direct directional selection on the geno-phenotype (i.e., it is possible that 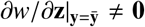) since **H**_**z**_ is always singular. Due to this singularity, if there is any evolutionary equilibrium, there is an infinite number of them. Kirkpatrick and Lofsvold (1992) showed that if **G** is singular and constant, then the evolutionary equilibrium that is achieved depends on the initial conditions. Our results extend the relevance of Kirkpatrick and Lofsvold’s (1992) observation by showing that **H**_**z**_ is always singular and remains so as it evolves. Moreover, since both the developmental (Layer 7, Eq. 1b) and environmental (Layer 7, Eq. 1c) constraints must be satisfied throughout the evolutionary process, the developmental and environmental constraints determine the admissible evolutionary trajectory and the admissible evolutionary equilibria if mutational variation exists in all directions of genotype space. Therefore, developmental and environmental constraints together with direct directional selection jointly define the evolutionary outcome if mutational variation exists in all directions of genotype space.

Since selection response is relatively completely described by total genotypic selection, further insight can be gained by rearranging the extended mechanistic Lande equation for the geno-phenotype (Layer 7, Eq. 7) in terms of total genotypic selection. Using the rearrangement in Layer 7, Eq. 5 and making the assumptions i-iii in the previous paragraph, the extended mechanistic Lande equation in Layer 7, Eq. 7 becomes

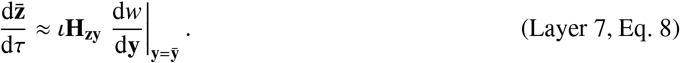

If the mutational covariance matrix **H**_**y**_ is non-singular, then the mechanistic additive genetic cross-covariance matrix between geno-phenotype and genotype **H**_**zy**_ is non-singular so evolutionary equilibrium 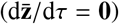 implies absence of total genotypic selection (i.e., 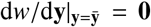) to first order of approximation. Moreover, dropping assumptions i-iii and to first order, lack of total genotypic selection provides a necessary and sufficient condition for evolutionary equilibria in the absence of exogenous environmental change and of absolute mutational constraints (Layer 7, Eq. 5). Consequently, evolutionary equilibria depend on development and niche construction since total genotypic selection depends on the developmental matrix and on total niche construction by the genotype (Eq. 9). However, since 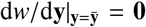 has only as many equations as there are lifetime genotypic traits and since not only the genotype but also the phenotype and environmental traits must be determined, then 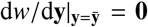 provides fewer equations than variables to solve for. Hence, absence of total genotypic selection still implies an infinite number of evolutionary equilibria. Again, only the subset of evolutionary equilibria that satisfy the developmental (Layer 7, Eq. 1b) and environmental (Layer 7, Eq. 1c) constraints are admissible, and so the number of admissible evolutionary equilibria may be finite. Therefore, admissible evolutionary equilibria have a dual dependence on developmental and environmental constraints: first, by the constraints’ influence on total genotypic selection and so on evolutionary equilibria; and second, by the constraints’ specification of which evolutionary equilibria are admissible.

Because we assume that mutants arise when residents are at carrying capacity, the analogous statements can be made for the evolutionary dynamics of a resident vector in terms of lifetime reproductive success (Eq. 8). Using the relationship between selection gradients in terms of fitness and of expected lifetime reproductive success (Eq. S2.3.12), the evolutionary dynamics of 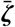 for *ζ* ∈ {**x, y, z, ϵ, m**} (Layer 7, Eq. 1a) are equivalently given by

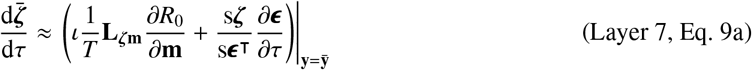

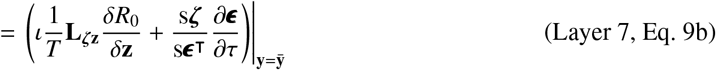

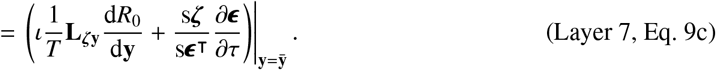

To close, the evolutionary dynamics of the environment can be written in a particular form that is insightful. In SI section S5.8, we show that the evolutionary dynamics of the environment satisfy

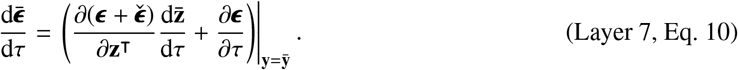

Thus, the evolutionary change of the environment comprises “joint” direct niche construction and exogenous environmental change.

## 6. Example: allocation to growth vs reproduction

We now provide an example that illustrates the method and some of the points above. To do this, we use a life-history model rather than a model of morphological development as the former is simpler yet sufficient to illustrate the points. In particular, this example shows that our results above enable direct calculation of the evo-devo dynamics and the evolution of the constraining matrices **H** and **L** and provide an alternative method to dynamic optimization to identify the evolutionary outcomes under explicit developmental constraints. We first describe the example where development is non-social and then extend the example to make development social.

### 6.1. Non-social development

We consider the classic life-history problem of modeling the evolution of resource allocation to growth vs reproduction (Gadgil and Bossert, 1970; León, 1976; Schaffer, 1983; Stearns, 1992; Roff, 1992; Kozłowski and Teriokhin, 1999). Let there be one phenotype (or state variable), one genotypic trait (or control variable), and no environmental traits. In particular, let *x*_*a*_ be a mutant’s phenotype at age *a* (e.g., body size or resources available) and *y*_*a*_ ∈ [0, 1] be the mutant’s fraction of resource allocated to phenotype growth at that age. Let the mutant survival probability *p*_*a*_ = *p* be constant for all *a* ∈ {1, …, *N*_a_ − 1} with 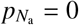, so survivorship is *ℓ*_*a*_ = *p*^*a*−1^ for all *a* ∈ {1, …, *N*_a_} with 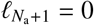. Let the mutant fertility be

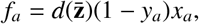

where (1 − *y*_*a*_)*x*_*a*_ is the resource a mutant allocates to reproduction at age *a* and 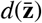 is a positive density-dependent scalar that brings the resident population size to carrying capacity. Let the developmental constraint be

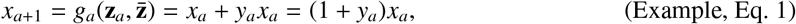

with initial condition 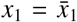, where *y*_*a*_ *x*_*a*_ is the resource a mutant allocates to growth at age *a*. These equations are a simplification of those used in the classic life-history problem of finding the optimal resource allocation to growth vs reproduction in discrete age (Gadgil and Bossert, 1970; León, 1976; Schaffer, 1983; Stearns, 1992; Roff, 1992; Kozłowski and Teriokhin, 1999). In life-history theory, one assumes that at evolutionary equilibrium, a measure of fitness such as lifetime reproductive success is maximized by an optimal control **y**^*^ yielding an optimal pair (**x**^*^, **y**^*^) that is obtained with dynamic programming or optimal control theory (Sydsæter et al., 2008). Instead, here we illustrate how the evolutionary dynamics of 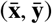 can be analyzed with the equations derived in this paper, including identification of an optimal pair (**x**^*^, **y**^*^).

We begin by obtaining analytical expressions for the phenotype and the density dependent scalar. Although an analytical expression for the phenotype is possible for this simple model, numerical solution is sufficient for more complex models. Iterating the recurrence given by the developmental constraint (Example, Eq. 1) yields the mutant phenotype at age *a*

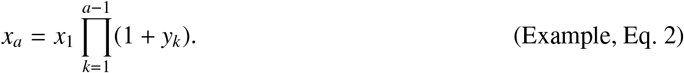

To find the density-dependent scalar, we note that a resident at carrying capacity satisfies the Euler-Lotka equation 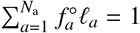 (Eq. S2.4.9), which yields

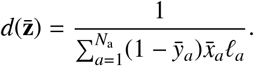

We now calculate the elements of the bottom layers that we need to calculate genetic covariation and the evolutionary dynamics. Because there are no environmental traits, total immediate effects equal direct effects (i.e., layer 3 is immaterial). Also, because development is non-social, stabilized effects equal total effects (except for social feedback, which is simply the identity matrix; i.e., layer 5 is immaterial). So, we only need to compute layers 2 and 4, that is, direct and total effects. Using Eq. (5a), the entries of the direct selection gradients are given by

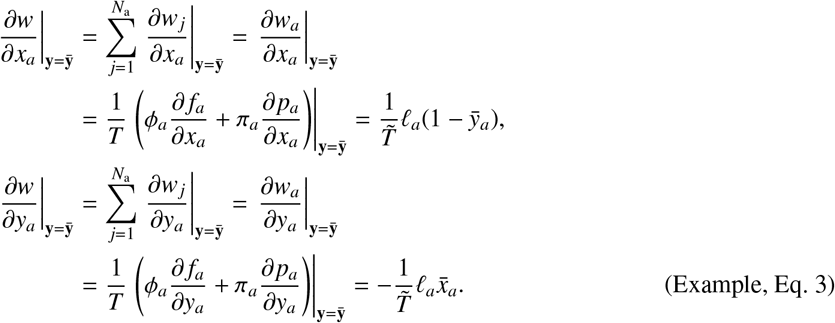

where the generation time without density dependence is

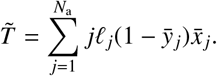

Thus, there is always direct selection for increased phenotype and against allocation to growth (except at the boundaries where 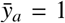 or 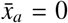). The entries of the matrices of direct effects on the phenotype (*a*: row, *j*: column) are given by

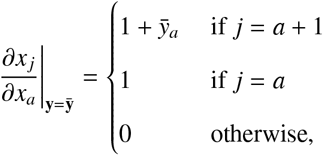

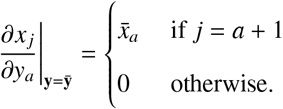

Using Eq. (12) and Eq. S5.2.15, the entries of the matrices of total effects on the phenotype are given by

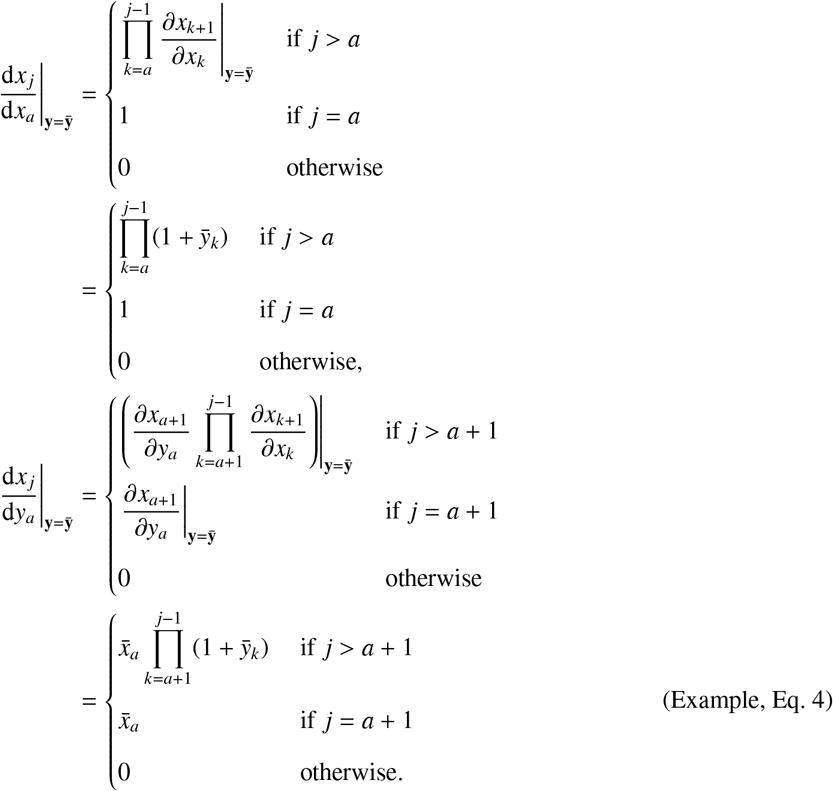

Then, using Layer 4, Eq. S20 and Eq. (9), the entries of the total selection gradients are given by

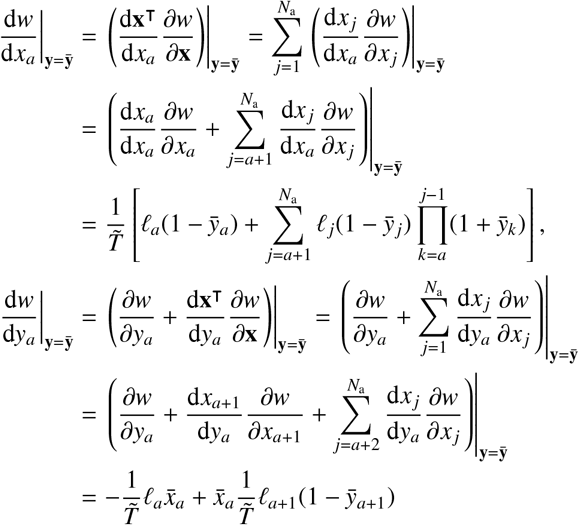

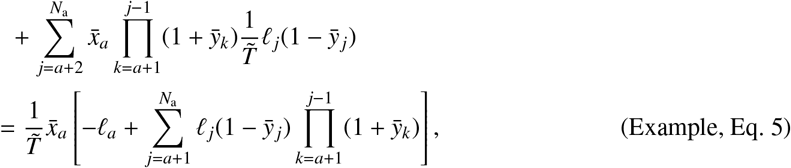

where we use the empty-product notation such that 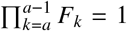 and the empty-sum notation such that 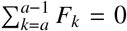 for any *F*_*k*_. There is thus always total selection for increased phenotype (except at the boundaries), although total selection for allocation to growth may be positive or negative.

Now, using Eqs. (1) and (3), the evo-devo dynamics are given by

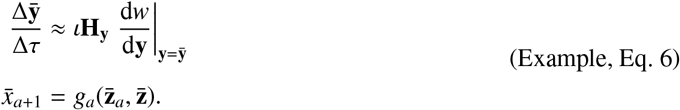

Using Layer 7, Eq. 1a, Layer 7, Eq. 4, and Layer 7, Eq. 5, the evolutionary dynamics of the resident phenotype in the limit as Δ*τ* → 0 are given by

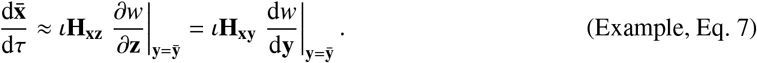

Note these are not equations in Lande’s form. In particular, the mechanistic additive genetic-cross covariance matrices involved are not symmetric and the selection gradients are not those of the evolving trait in the left-hand side; Example, Eq. 7 cannot be arranged in Lande’s form because the genotypic trait directly affects fitness (i.e., 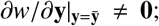 Example, Eq. 3). Importantly, **H**_**xz**_ and **H**_**xy**_ depend on 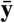 because of gene-phenotype interaction in development (i.e., the developmental map involves a product *y*_*a*_ *x*_*a*_ such that the total effect of the genotype on the phenotype depends on the genotype; Example, Eq. 4); consequently, Example, Eq. 7 is dynamically insufficient because the system does not describe the evolution of 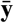. In turn, the evolutionary dynamics of the geno-phenotype are given by

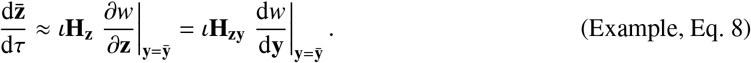

This system contains dynamic equations for all the evolutionarily dynamic variables, namely both the resident phenotype 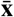 and the resident genotype 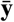, so it is determined and dynamically sufficient. The first equality in Example, Eq. 8 is in Lande’s form, but **H**_**z**_ is always singular. In contrast, the matrix **H**_**zy**_ in the second equality is non-singular if the mutational covariance matrix **H**_**y**_ is non-singular. Thus, the total selection gradient of the genotype provides a relatively complete description of the evolutionary process of the geno-phenotype.

Let the entries of the mutational covariance matrix be given by

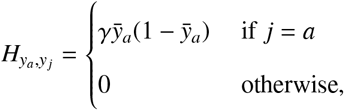

where 0 < γ ≪ 1 so the assumption of marginally small mutational variance, namely 0 < tr(**H**_**y**_) ≪ 1, holds. Thus, **H**_**y**_ is diagonal and becomes singular only at the boundaries where the resident genotype is zero or one. Then, from Example, Eq. 6, the evolutionary equilibria of the genotypic trait at a given age and their stability are given by the sign of its corresponding total selection gradient.

Let us now find the evolutionary equilibria and their stability for the genotypic trait. Using Example, Eq. 5, starting from the last age, the total selection on the genotypic trait at this age is

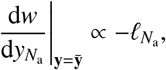

which is always negative so the stable resident genotypic trait at the last age is

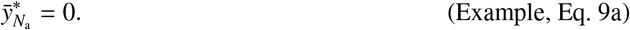

That is, no allocation to growth at the last age. Continuing with the second-to-last age, the total selection on the genotypic trait at this age is

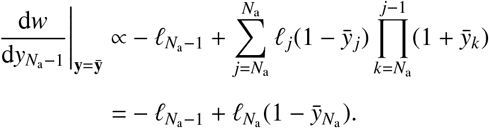

Evaluating at the optimal genotypic trait at the last age (Example, Eq. 9a) and substituting *ℓ*_*a*_ = *p*^*a*−1^ yields

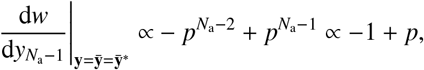

which is negative (assuming *p* < 1) so the stable resident genotypic trait at the second-to-last age is

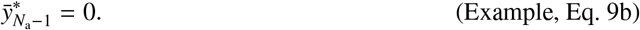

Continuing with the third-to-last age, the total selection on the genotypic trait at this age is

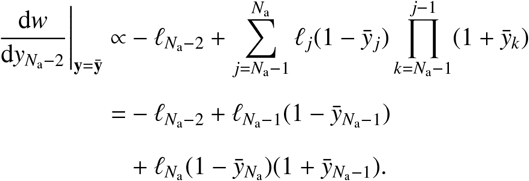

Evaluating at the optimal genotypic trait at the last two ages (Example, Eq. 9a and Example, Eq. 9b) and substituting *ℓ*_*a*_ = *p*^*a*−1^ yields

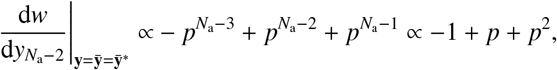

which is positive if

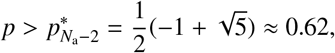

(thus, 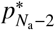 is the reciprocal of the golden ratio). So the stable resident genotypic trait at the third-to-last age is

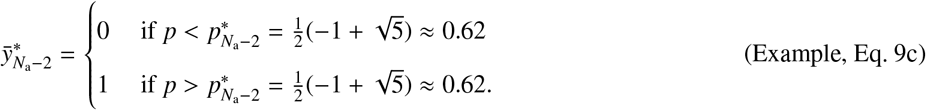

If 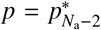, the genotypic trait at such age is selectively neutral, but we ignore this case as without an evolutionary model for *p* it is biologically unlikely that survival is and remains at such precise value. Hence, there is no allocation to growth at this age for low survival and full allocation for high survival. Continuing with the fourth-to-last age, the total selection on the genotypic trait at this age is

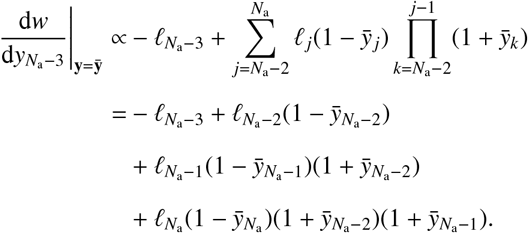

Evaluating at the optimal genotypic trait at the last three ages (Example, Eq. 9a-Example, Eq. 9c) and substituting *ℓ*_*a*_ = *p*^*a*−1^ yields

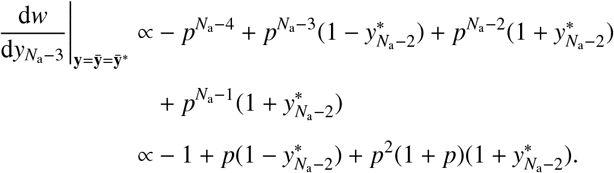

If 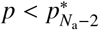, this is

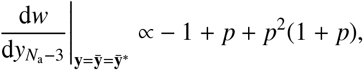

which is positive if

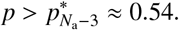

If 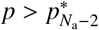, the gradient is

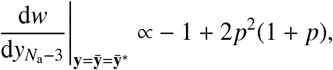

which is positive if

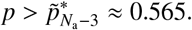

Hence, the stable resident genotypic trait at the fourth-to-last age is

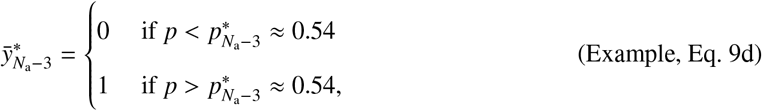

for 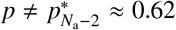. Again, this is no allocation to growth for low survival, although at this earlier age survival can be smaller for allocation to growth to evolve. Numerical solution for the evo-devo dynamics using Example, Eq. 6 is given in Fig. 5. The associated evolution of the **H**_**z**_ matrix, plotting Layer 6, Eq. 6, is given in Fig. 6. The code used to generate these figures is in the SI.

**Figure 5.**
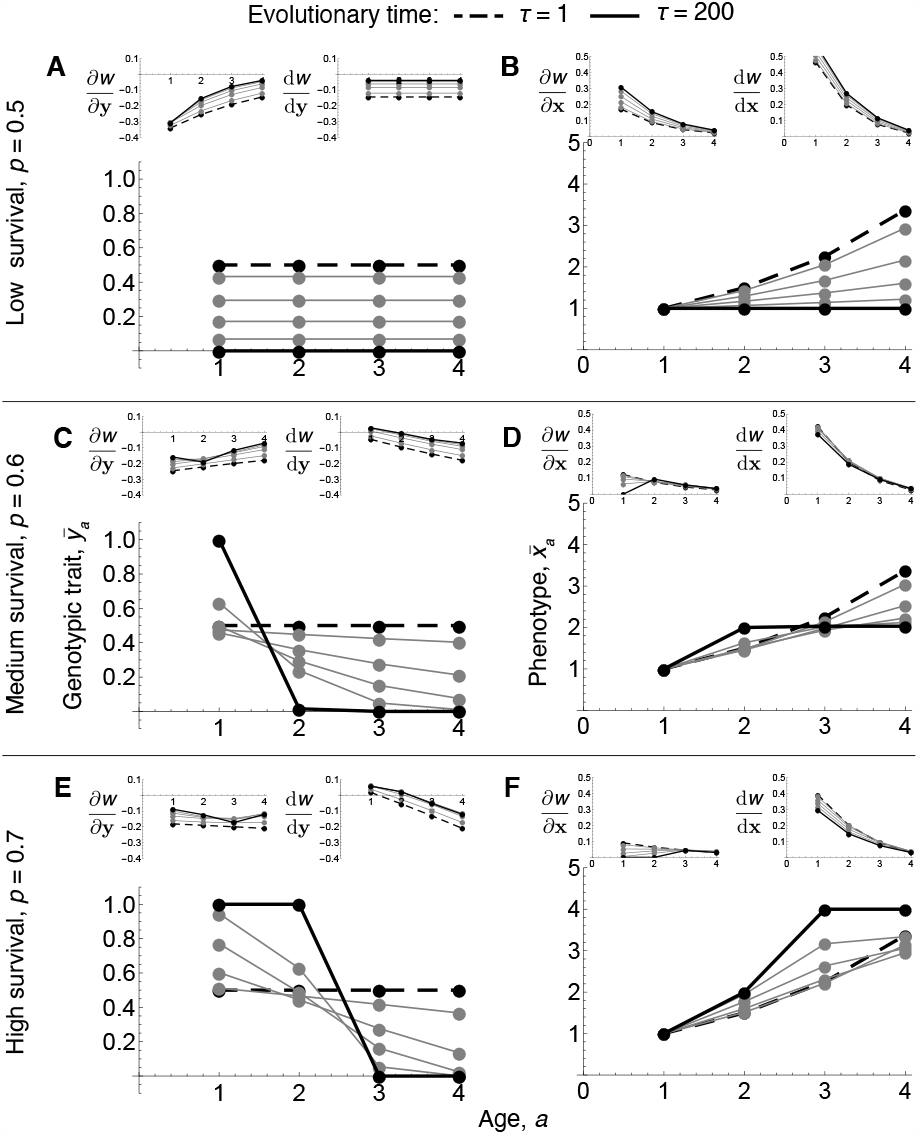
Example. Numerical solution of the evolutionary dynamics of the genotype and associated developmental dynamics of the phenotype. Large plots give the resident genotype (left) or phenotype (right) vs age over evolutionary time for various survival probabilities *p*. For low survival, (A) the genotypic trait evolves from mid allocation to growth over life to no allocation to growth over life. (B) This leads the phenotype to evolve from indeterminate growth to no growth over life. For medium survival, (C) the genotypic trait evolves from mid allocation to growth over life to full allocation to growth at the first age and no allocation to growth at later ages. (D) This leads the phenotype to evolve from indeterminate growth to determinate growth for one age. For high survival, (E) the genotypic trait evolves from mid allocation to growth over life to full allocation to growth at the first two ages and no allocation to growth at later ages. (F) This leads the phenotype to evolve from indeterminate growth to determinate growth for two ages, thus reaching the largest size of all these scenarios. Small plots give the associated direct and total selection gradients; the costate variable of *x* is proportional to the total selection gradient of the phenotype at large τ. In all scenarios, there is always negative direct selection for allocation to growth (small left plots in A,C,E) and non-negative direct selection for the phenotype (small left plots in B,D,F); hence, the genotype does not reach a fitness peak in geno-phenotype space, but for medium and high survival the phenotype at early ages reaches a fitness peak. Total selection for growth (small right plots in A,C,E) is positive only for early ages and sufficiently high survival; it remains non-zero at evolutionary equilibrium because 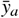 cannot evolve further as mutational variation vanishes as 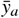approaches 0 or 1. Total selection for the phenotype (small right plots in B,D,F) is always positive, even at evolutionary equilibrium; the phenotype cannot evolve further despite persistent total selection for it because genetic variation vanishes as mutational variation vanishes. The numerical evolutionary outcomes match the analytical expressions for the genotype (Example, Eq. 9) and associated phenotype (Example, Eq. 2). 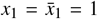. From Eq. S2.3.5a, the carrying capacity is 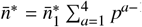. We let 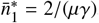, so 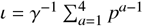.

**Figure 6.**
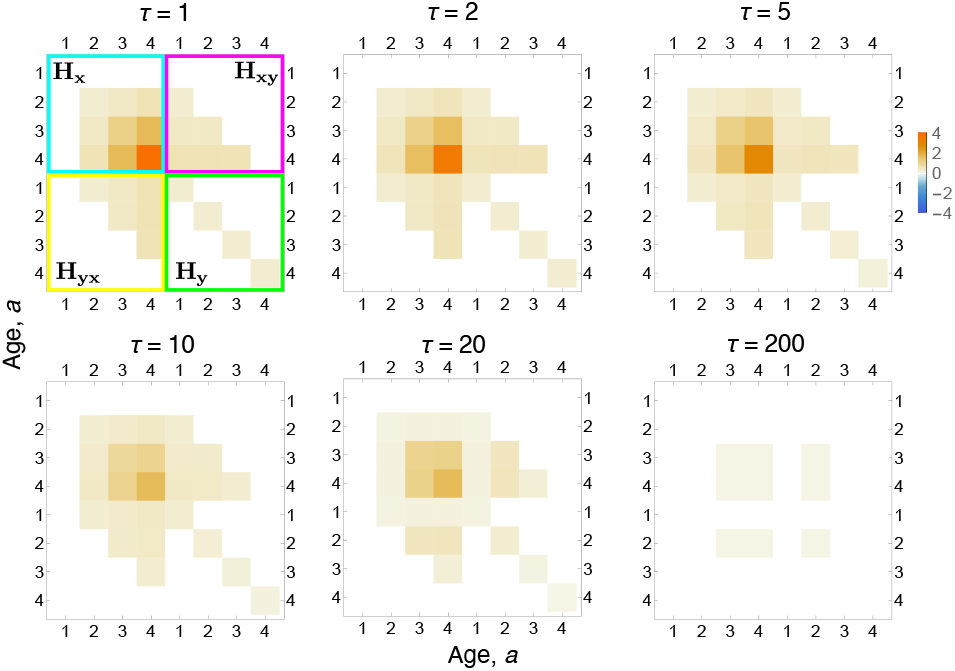
Resulting evolutionary dynamics of the mechanistic additive genetic covariance matrix **H**_**z**_. The upper-left quadrant (blue) is the mechanistic additive genetic covariance matrix **H**_**x**_ of the phenotype, that is, of the state variable. For instance, at the initial evolutionary time, the genetic variance for the phenotype is higher at later ages, and the phenotype at age 3 is highly genetically correlated with the phenotype at age 4. As evolutionary time progresses, genetic covariation vanishes as mutational covariation vanishes (**H**_**y**_ becomes singular) as genotypic traits approach their boundary values. *p* = 0.7. The evolutionary times τ shown correspond to those of Fig. 5.

### 6.2. Social development

Consider a slight modification of the previous example, so that development is social. Let the mutant fertility be

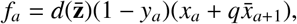

where the available resource is now given by 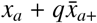 for some constant *q* (positive, negative, or zero). Here the source of social development can be variously interpreted, including that an immediately older resident contributes to (positive *q*) or takes from (negative *q*) the resource of the focal individual, or that the focal individual learns from the older resident (positive or negative *q* depending on whether learning increases or increases the phenotype). Let the developmental constraint be

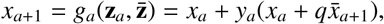

with initial condition 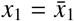.

To see what the stabilization of social development is, imagine the first individual in the population having a resident genotype developing with such developmental constraint. As it is the first individual in the population, such individual has no social partners so the developed phenotype is

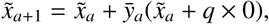

with 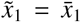. Imagine a next resident individual who develops in the context of such initial individual. This next individual develops the phenotype

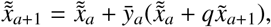

with 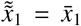, but then 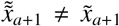. Iterating this process, the resident phenotype 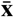 may converge to a socio-devo equilibrium 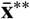, which satisfies

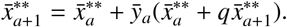

Solving for 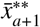 yields a recurrence for the resident phenotype at socio-devo equilibrium

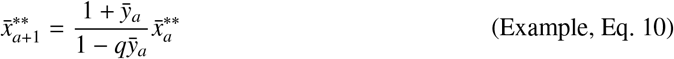

provided that 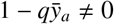.

Iterating Example, Eq. 10 yields the resident phenotype at socio-devo equilibrium

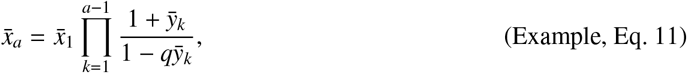

where we drop the ^**^ for simplicity. To determine when this socio-devo equilibrium is socio-devo stable, we find the eigenvalues of 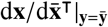 (Eq. 28) as follows. The entries of the matrix of the direct social effects on the phenotype are given by

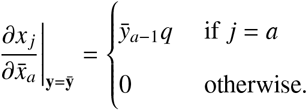

Hence, from Eq. S5.6.8 and Eq. S5.6.9, 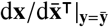 is lower-triangular, so its eigenvalues are the values in its main diagonal, which are given by 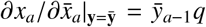. Thus, the eigenvalues of 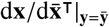 have absolute value strictly less than one if |*q*| < 1, in which case the socio-devo equilibrium in Example, Eq. 11 is socio-devo stable.

Therefore, let 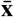 be the SDS resident phenotype given by Example, Eq. 11 with |*q*| < 1. Then, the evo-devo dynamics are given by Example, Eq. 6. Using Layer 7, Eq. 1a, Layer 7, Eq. 4, and Layer 7, Eq. 5, the evolutionary dynamics of the phenotype in the limit as Δτ → 0 are now given by

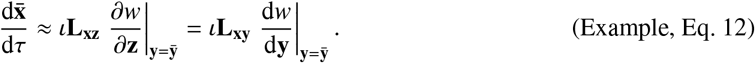

This system is dynamically insufficient as **L**_**xz**_ and **L**_**xy**_ depend on 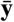 because of gene-phenotype interaction in development. In turn, the evolutionary dynamics of the geno-phenotype are given by

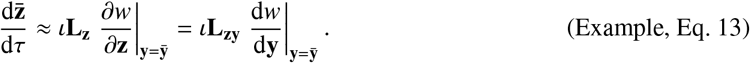

This system is dynamically sufficient as it contains dynamic equations for all evolutionarily dynamic variables, namely both 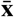 and 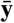. While **L**_**z**_ in the first equality is always singular, the matrix **L**_**zy**_ in the second equality is non-singular if the mutational covariance matrix **H**_**y**_ is non-singular. Thus, the total selection gradient of the genotype still provides a relatively complete description of the evolutionary process of the geno-phenotype.

We can similarly find that the total selection gradient of the genotypic trait at age *a* is

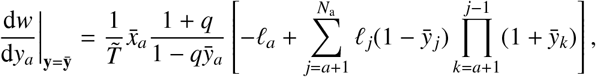

where the generation time without density dependence is now

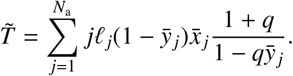

This total selection gradient of the genotypic trait at age *a* has the same sign as that found in the model for non-social development (Example, Eq. 5). Hence, the stable evolutionary equilibria for the genotype are still given by Example, Eq. 9. Yet, the associated phenotype, given by Example, Eq. 11, may be different due to social development (Fig. 7). That is, social development here does not affect the evolutionary equilibria, as it does not affect the zeros of the total selection gradient of the genotype which gives the zeros of the evolutionary dynamics of the geno-phenotype (Example, Eq. 13). Instead, social development affects here the developmental constraint so it affects the admissible evolutionary equilibria of the phenotype. Numerical solution for the evo-devo dynamics using Example, Eq. 6 is given in Fig. 7. For the *q* chosen, the phenotype evolves to much larger values due to social feedback than with non-social development although the genotype evolves to the same values. The associated evolution of the **L**_**z**_ matrix, using Layer 6, Eq. 9, is given in Fig. 8. The code used to generate these figures is in the SI.

**Figure 7.**
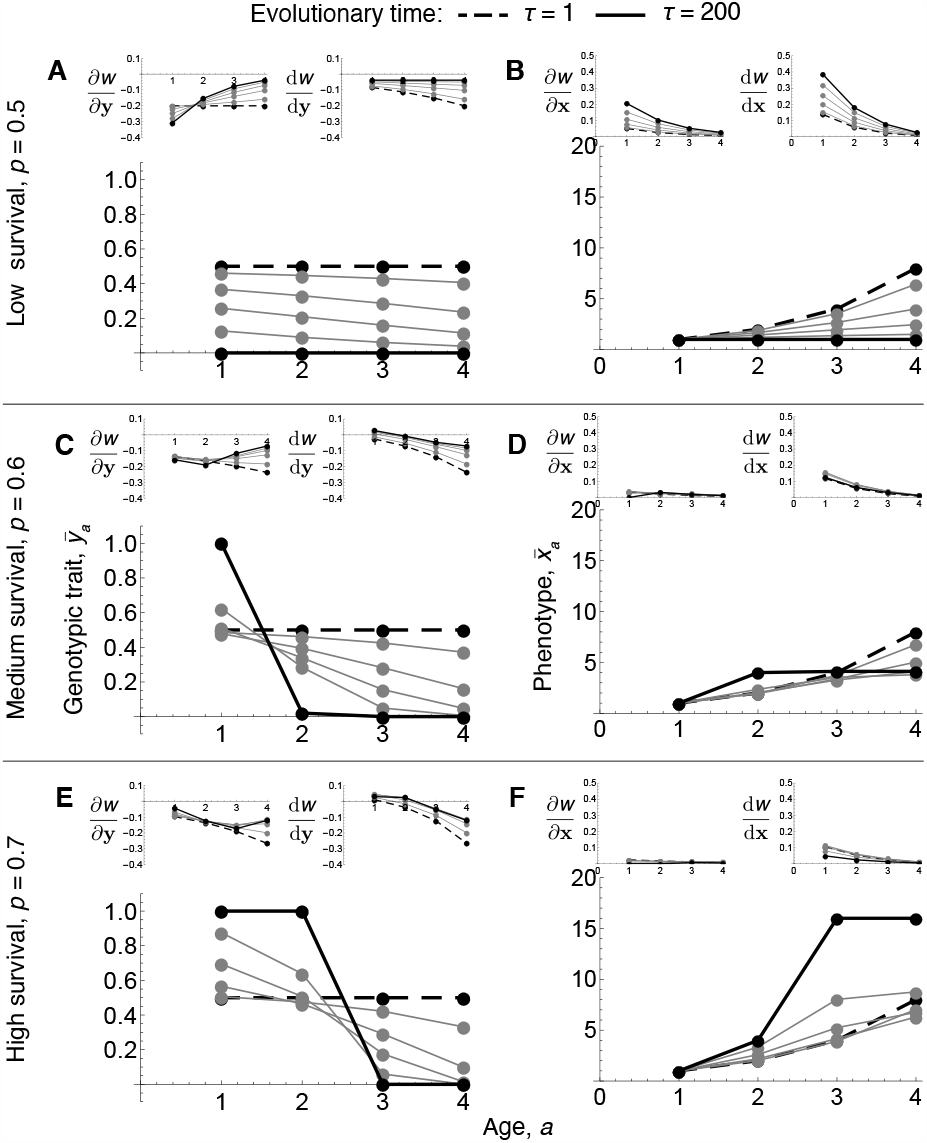
Example with social development. The genotype evolves to the same values as those with non-social development in Fig. 5. However, the phenotype evolves to much larger values due to social development. Total and direct selection for the phenotype (small plots in B,D,F) are smaller than with non-social development, to the point of there being almost no direct selection on the phenotype with high survival indicating that the phenotype is near a fitness peak even for late ages (small left plot in F). Large plots give the resident genotype (left) or phenotype (right) vs age over evolutionary time for various survival probabilities *p*. Small plots give the associated direct and total selection gradients. The numerical evolutionary dynamics of the genotype match the analytical expressions for the genotype (Example, Eq. 9) and associated phenotype (Example, Eq. 11). ι is as in Fig. 5. *q* = 0.5.

**Figure 8.**
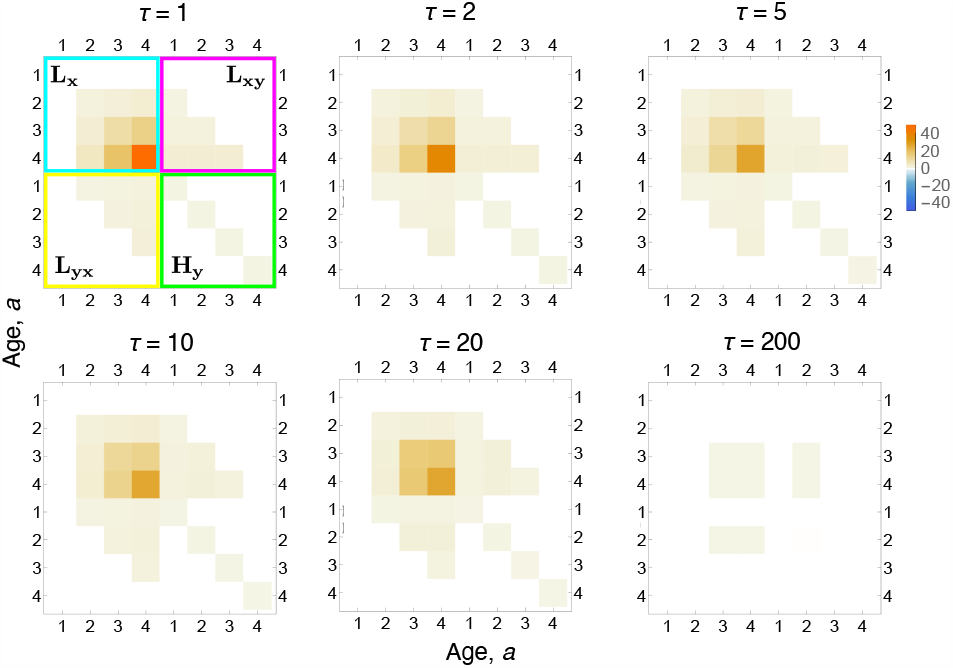
Resulting dynamics of the mechanistic additive socio-genetic cross-covariance matrix **L**_**z**_. The structure and dynamics of **L**_**z**_ here are similar to those of **H**_**z**_ in Fig. 8 but the magnitudes are an order of magnitude larger (compare bar legends). *p* = 0.7, *q* = 0.5. The evolutionary times τ shown correspond to those of Fig. 7.

## 7. Discussion

We have addressed the question of how development affects evolution by formulating a mathematical framework that integrates explicit developmental dynamics into evolutionary dynamics. The framework integrates age progression, explicit developmental constraints according to which the phenotype is constructed across life, and evolutionary dynamics. This framework yields a description of the structure of genetic covariation, including the additive effects of allelic substitution 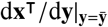, from mechanistic processes. The framework also yields a dynamically sufficient description of the evolution of developed phenotypes in gradient form, such that their long-term evolution can be described as the climbing of a fitness landscape within the assumptions made. This framework provides a tractable method to model the evo-devo dynamics for a broad class of models. We also obtain formulas to compute the sensitivity of the solution of a recurrence (here, the phenotype) to perturbations in the solution or parameters at earlier times (here, ages), which are given by d**x**^⊺^/d*ζ* for *ζ* ∈ {**x, y**}. Overall, the framework provides a theory of long-term constrained evolutionary dynamics, where the developmental and environmental constraints determine the admissible evolutionary path (Layer 7, Eq. 1).

Previous understanding suggested that development affects evolution by inducing genetic covariation and genetic constraints, although the nature of such constraints had remained uncertain. We find that genetic constraints are necessarily absolute in a generally dynamically sufficient description of long-term phenotypic evolution in gradient form. This is because dynamic sufficiency in general requires that not only phenotypic but also genotypic evolution is followed. Because the phenotype is related to the genotype via development, simultaneously describing the evolution of the genotype and phenotype in gradient form entails that the associated constraining matrix (**H**_**z**_ or **L**_**z**_) is necessarily singular with a maximum number of degrees of freedom given by the number of lifetime genotypic traits (*N*_a_*N*_g_). Consequently, genetic covariation is necessarily absent in as many directions of geno-phenotype space as there are lifetime developed traits (*N*_a_*N*_p_). Since the constraining matrix is singular, direct directional selection is insufficient to identify evolutionary equilibria in contrast to common practice. Instead, total genotypic selection, which depends on development, is sufficient to identify evolutionary equilibria if there are no absolute mutational constraints and no exogenous plastic response. Yet, absence of total genotypic selection is insufficient to identify evolutionary outcomes because the singularity of the constraining matrix associated to direct geno-phenotypic selection entails that if there is any evolutionary equilibrium and no exogenous plastic response, then there is an infinite number of evolutionary equilibria that depend on development. Which of these equilibria is attained also depends on development as it determines the admissible evolutionary trajectory and so the admissible equilibria. The adaptive topography in phenotype space is often assumed to involve a non-singular **G**-matrix where evolutionary outcomes occur at fitness landscape peaks (i.e., where 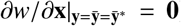). In contrast, we find that the evolutionary dynamics differ from that representation in that evolutionary outcomes occur at best (i.e., without absolute mutational constraints) at peaks in the admissible evolutionary path determined by development (i.e., where 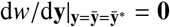), and that such path peaks do not typically occur at landscape peaks (so generally 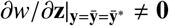).

The singularity of the constraining matrix (**H**_**z**_ or **L**_**z**_) is not due to our adaptive dynamics assumptions. Under quantitative genetics assumptions, the additive genetic covariance matrix of phenotype **x** is 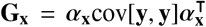as described in the introduction, and here we use the subscripts **x** to highlight that this *α* matrix is for the regression coefficients of the phenotype with respect to gene content. Under quantitative genetics assumptions, the matrix cov[**y, y**] describes the observed covariance in allele frequency due to any source, so it describes standing covariation in allele frequency. Under our adaptive dynamics assumptions, we obtain an **H**_**x**_ matrix that has the same form of **G**_**x**_, but where cov[**y, y**] describes the covariance in genotypic traits only due to mutation at the current evolutionary time step among the possible mutations, so it describes (expected) mutational covariation. Regardless of whether cov[**y, y**] describes standing covariation in allele frequency or mutational covariation, the additive genetic covariance matrix in genophenotype space 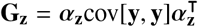is always singular because the developmental matrix of the geno-phenotype 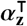 has fewer rows than columns: that is, the degrees of freedom of **G**_**z**_ have an upper bound given by the number of loci (or genetic predictors) while the size of **G**_**z**_ is given by the number of loci and of phenotypes. Thus, whether one considers standing or mutational covariation, the additive genetic covariance matrix of the geno-phenotype is always singular. Eliminating traits from the analysis to render **G**_**z**_ non-singular as traditionally recommended (Lande, 1979) either renders the gradient system underdetermined and so dynamically insufficient in general (if allele frequency 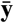 is removed), or prevents a description of phenotypic evolution as the climbing of a fitness landscape (if the mean phenotype 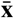 is removed). The singularity of **H** and **L** in geno-phenotype space persists despite evolution of the developmental map, regardless of the number of genotypic traits or phenotypes provided there is any phenotype, and in the presence of endogenous or exogenous environmental change. Thus, we find that a dynamically sufficient description of phenotypic evolution in gradient form generally requires a singular constraining matrix. It remains to be seen whether these conclusions hold under other types of developmental constraints, for instance, inequality constraints or stochastic development.

In our results, dynamic sufficiency for phenotypic evolution in gradient form requires that the constraining matrix is in geno-phenotype space particularly because of non-linear development. The **H**-matrix in phenotype space generally depends on the resident genotype via both the mutational covariance matrix and the developmental matrix. The developmental matrix depends on the resident genotype due to non-linear development, particularly gene-gene interaction, gene-phenotype interaction, and gene-environment interaction (see text below Layer 6, Eq. 5). The analogous dependence of **G** on allele frequency holds under quantitative genetics assumptions for the same reasons (Turelli, 1988; Service and Rose, 1985). If development is linear (i.e., the developmental map for all phenotypes is a linear function in all its variables at all ages), the developmental matrix no longer depends on the resident genotype (or allele frequency under quantitative genetics assumptions). If in addition the mutational covariance matrix is independent of the resident genotype, then the constraining matrix **H** in phenotype space is no longer dependent on the resident genotype. Thus, if one assumes linear development and both mutational covariation and phenotypic selection being independent of the resident genotype (in addition to no social interactions, no exogenous plastic response, no total immediate genotypic selection, and no niche-constructed effects of the phenotype on fitness; Layer 7, Eq. 6), the **H** matrix in phenotype space becomes constant and the mechanistic Lande equation (Layer 7, Eq. 6) becomes dynamically sufficient. However, even simple models of explicit development involve non-linearities (e.g., Example, Eq. 1) and mutational covariation depends on the resident genotype whenever the genotype is constrained to take values within a finite range (e.g., between zero and one). Thus, consideration of even slightly realistic models of development seems unlikely to allow for a long-term dynamically sufficient mechanistic Lande equation (i.e., following only phenotypic evolution).

Extensive research efforts have been devoted to determining the relevance of constraints in adaptive evolution (Arnold, 1992; Hine and Blows, 2006; Hansen and Houle, 2008; Jones et al., 2014; Hine et al., 2014; Engen and Sæther, 2021). Empirical research has found that the smallest eigenvalue of **G** in phenotype space is often close to zero (Kirkpatrick and Lofsvold, 1992; Hine and Blows, 2006; McGuigan and Blows, 2007). Mezey and Houle (2005) found a non-singular **G**-matrix for 20 morphological (so, developed) traits in fruit flies. Our results suggest **G** singularity would still arise in these studies if enough traits are included to provide a dynamically sufficient description of long-term phenotypic evolution on an adaptive topography (i.e., if allele frequency were included in the analysis as part of the multivariate “geno-phenotype”).

Previous theory has offered limited predictions as to when the **G**-matrix would be singular. These include that incorporating more traits in the analysis renders **G** more likely to be singular as the traits are more likely to be genetically correlated, such as in infinite-dimensional traits (Gomulkiewicz and Kirkpatrick, 1992; Kirkpatrick and Lofsvold, 1992). Suggestions to include gene frequency as part of the trait vector in the classic Lande equation (e.g., Barfield et al., 2011) have been made without noticing that doing so entails that the associated **G**-matrix is necessarily singular. Kirkpatrick and Lofsvold (1992, p. 962 onwards) showed that, assuming that **G** in phenotypic space is singular and constant, then the evolutionary trajectory and equilibria depend on the evolutionarily initial conditions of the phenotype. Such dependence on the evolutionarily initial conditions is sometimes called phylogenetic constraints (Hansen, 1997). Our results extend the relevance of Kirkpatrick and Lofsvold’s (1992) analysis by our observation that the constraining matrix is always singular in a dynamically sufficient gradient system for long-term phenotypic evolution, even with few traits and evolution of the constraining matrix. In our framework, the evolutionarily initial conditions of the phenotype are given by the developmental constraint evaluated at the evolutionarily initial genotype and environment. Hence, the evolutionary trajectory, equilibria, and outcomes depend on the developmental constraint, which provides the admissible evolutionary path.

Multiple mathematical models have studied whether **G** is singular. Recently, simulation work studying the effect of pleiotropy on the structure of the **G**-matrix found that the smallest eigenvalue of **G** is very small but positive (Engen and Sæther, 2021, Tables 3 and 5). Our findings indicate that this model and others (e.g., Wagner, 1984; Barton and Turelli, 1987; Wagner, 1989; Wagner and Mezey, 2000; Martin, 2014; Morrissey, 2014, 2015) would recover **G**-singularity by considering the geno-phenotype, so that both allele frequency and phenotype change are part of the gradient system. Other recent simulation work found that a singular **G**-matrix due to few segregating alleles still allows the phenotype to reach its unconstrained optimum if all loci have segregating alleles at some point over the long run, thus allowing for evolutionary change in all directions of phenotype space in the long run (Barton, 2017, Fig. 3). Our results indicate that such a model attains the unconstrained optimum because it assumes that fitness depends on a single phenotype at a single age, that additive effects of allelic substitution are always positive, and that there is no direct genotypic selection and no niche-constructed effects of the genotype on fitness (i.e., there ∂*w*/∂**y** = **0** and (d**ϵ**^⊺^/d**y**)(∂*w*/∂**ϵ**) = **0**, so 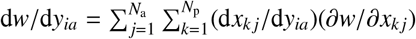, which since fitness depends on a single trait k at a single age j further reduces to (d*x*_kj_/d*y*_*ia*_)(∂*w*/∂*x*_kj_); hence, assuming d*x*_kj_/d*y*_*ia*_ > 0, there we have that d*w*/d*y*_*i*j_ = 0 for any locus *i* implies ∂*w*/∂*x*_kj_ = 0; Layer 4, Eq. S21). Our results show that when at least one of these assumptions does not hold, the unconstrained optimum is not necessarily achieved (as illustrated in Example, Eq. 3 and Fig. 5). In our framework, phenotypic evolution converges at best to constrained fitness optima, which may under certain conditions coincide with unconstrained fitness optima. Convergence to constrained fitness optima under no absolute mutational constraints occurs even with the fewest number of traits allowed in our framework: two, that is, one genotypic trait and one phenotype with one age each (or in a standard quantitative genetics framework, allele frequency at a single locus and one quantitative trait that is a function of such allele frequency). Such constrained adaptation has important implications for biological understanding (see e.g., Kirkpatrick and Lofsvold, 1992; Gomulkiewicz and Kirkpatrick, 1992). It is also consistent with empirical observations of lack of selection response in the wild despite selection and genetic variation (Merilä et al., 2001; Hansen and Houle, 2004; Pujol et al., 2018), and of relative lack of stabilizing selection across many species (Kingsolver et al., 2001; Kingsolver and Diamond, 2011).

Other modelling work studied the evolutionary relevance of developmental factors without concluding that there are necessarily absolute genetic constraints. Wagner (1984, 1989) constructed and analyzed evolutionary models considering developmental maps, and wrote the **G**-matrix in terms of his developmental matrix to assess its impact on the maintenance of genetic variation. Wagner (1984, 1988, 1989) and Wagner and Mezey (2000) did not simultaneously track the evolution of genotypes and phenotypes, so did not conclude that the associated **G**-matrix is necessarily singular or that the developmental matrix affects evolutionary equilibria. Altenberg (1995) used Wagner’s (1984, 1989) developmental matrix to model constrained adaptation (see Altenberg 1995 Fig. 2) without evolutionary dynamics, but found an analytical expression for the constrained optimum phenotype that equals the globally optimum phenotype (i.e., mapping the constrained optimal genotype in his Eq. 10 to phenotype, 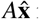 in his notation, yields the globally optimum phenotype). This and other studies (Via and Lande 1985, Houle 1991, his Fig. 2 and Kirkpatrick and Lofsvold 1992, their Fig. 5) have illustrated how constrained evolutionary dynamics would proceed if the **G** matrix is singular, considering it as a possible case rather than a necessary case. Other models have found that epistasis can cause the evolutionary dynamics to take an exponentially long time to reach fitness peaks (Kaznatcheev, 2019). We find that as the constraining matrix in geno-phenotype space has at least as many zero eigenvalues as there are lifetime phenotypes (i.e., *N*_a_*N*_p_), even if there were infinite time, the population does not necessarily reach a fitness peak in geno-phenotype space. The population eventually reaches a fitness peak in genotype space if there are no absolute mutational constraints after the landscape is modified by: (1) the interaction of the total effects of the genotype on phenotype and direct phenotypic selection and (2) the total niche-constructed effects of the genotype on fitness.

We find that total genotypic selection provides more information regarding selection response than direct directional selection or other forms of total selection. We show that evolutionary equilibria occur when total genotypic selection vanishes if there are no absolute mutational constraints and no exogenous plastic response. Direct selection or total selection on the phenotype need not vanish at evolutionary equilibria, even if there are no absolute mutational constraints and no exogenous plastic response. As total genotypic selection depends on development rather than exclusively on direct selection, and as development determines the admissible evolutionary trajectory along which developmental and environmental constraints are satisfied, our findings show that development plays a major evolutionary role by sharing responsibility with selection for defining evolutionary equilibria and for determining the admissible evolutionary path.

Total selection gradients correspond to quantities that have received various names. Such gradients correspond to Caswell’s (1982, 2001) “total derivative of fitness” (denoted by him as dλ), Charlesworth’s (1994) “total differential” (of the population’s growth rate, denoted by him as d*r*), van Tienderen’s (1995) “integrated sensitivity” (of the population’s growth rate, denoted by him as IS), and Morrissey’s (2014, 2015) “extended selection gradient” (denoted by him as *η*). Total selection gradients measure total directional selection, so in our framework they take into account the downstream developmental effects of a trait on fitness. In contrast, Lande’s (1979) selection gradients measure direct directional selection, so in our framework’s terms they do not consider the developmentally immediate total effects of a trait on fitness nor the downstream developmental effects of a trait on fitness. We obtained compact expressions for total selection gradients as linear transformations of direct selection gradients, arising from the chain rule in matrix calculus notation (Layer 4, Eq. S19), analogously to previous expressions in terms of vital rates that have no developmental constraints (Caswell, 2001, Eq. 9.38).

Our mechanistic approach to total selection recovers the regression approach to total selection of Morrissey (2014). Morrissey (2014) defined the extended selection gradient as *η* = Φ*β*, where *β* is Lande’s selection gradient and Φ is the matrix of total effects of all traits on themselves (computed as regression coefficients between variables related by a path diagram rather than as total derivatives, which entails material differences with our approach as explained above). Morrissey (2014) used an equation for the total-effect matrix Φ (his Eq. 2) from path analysis (Greene, 1977, p. 380), which has the form of our matrices describing developmental feedback of the phenotype and the geno-phenotype (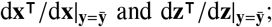 Eq. (11) and Layer 4, Eq. S8). Thus, interpreting Morrissey’s (2014) Φ as our 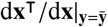(resp. 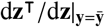) and *β* as our 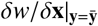(resp. 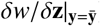) (i.e., Lande’s selection gradient of the phenotype or the geno-phenotype if environmental traits are not explicitly included in the analysis), then Layer 4, Eq. S20 (resp. Layer 4, Eq. S23) shows that the extended selection gradient *η* = Φ*β* corresponds to the total selection gradient of the phenotype 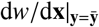(resp. of the geno-phenotype 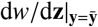). We did not show that 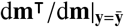 has the form of the equation for Φ provided by Morrissey (2014) (his Eq. 2), but it might indeed hold. If we interpret Φ as our 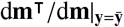 and *β* as our 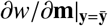(i.e., Lande’s selection gradient of the geno-envo-phenotype thus explicitly including environmental traits in the analysis), then Layer 4, Eq. S24 shows that the extended selection gradient *η* = Φ*β* corresponds to the total selection gradient of the geno-envo-phenotype 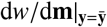.

Not all total selection gradients provide a relatively complete description of the selection response. We show in the SI sections S5.7 (Eq. S5.7.5) and S5.9 (Eq. S5.9.5) that the selection response of the geno-phenotype or the geno-envo-phenotype can respectively be written in terms of the total selection gradients of the geno-phenotype 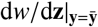 or the geno-envo-phenotype 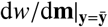, but such total selection gradients are insufficient to predict lack of selection response because they are premultiplied by a singular socio-genetic cross-covariance matrix. Also, the selection response of the phenotype can be written in terms of the total selection gradient of the phenotype 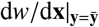, but this expression for the selection response has an additional term involving the total immediate selection gradient of the genotype 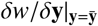, so the total selection gradient of the phenotype is insufficient to predict lack of selection response (even more considering that following the evolutionary dynamics of the phenotype alone is generally dynamically insufficient). In contrast, we have shown that the total selection gradient of the genotype 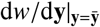 predicts lack of selection response if there are no absolute mutational constraints. Thus, out of all total selection gradients considered, only total genotypic selection provides a relatively complete description of the selection response. Morrissey (2015) considers that the total selection gradient of the genotype (his “inputs”) and of the phenotype (his “traits”) would be equal, but the last line of Layer 4, Eq. S21 shows that the total selection gradients of the phenotype and genotype are different in general, particularly due to direct genotypic selection and the total effects of genotype on phenotype.

Our results allow for the modeling of evo-devo dynamics in a wide array of settings. First, developmental and environmental constraints (Eqs. 1 and 2) can mechanistically describe development, gene-gene interaction, and gene-environment interaction, while allowing for arbitrary non-linearities and evolution of the developmental map. Several previous approaches have modelled gene-gene interaction, such as by considering multiplicative gene effects, but general analytical frameworks mechanistically linking gene-gene interaction, gene-environment interaction, developmental dynamics, and evolutionary dynamics have previously remained elusive (Rice, 1990; Hansen and Wagner, 2001; Rice, 2002; Hermisson et al., 2003; Carter et al., 2005; Rice, 2011). A historically dominant yet debated view is that gene-gene interaction has minor evolutionary effects (Hansen, 2013; Nelson et al., 2013; Paixão and Barton, 2016; Barton, 2017). Our finding that the constraining matrix (**H** or **L**) is necessarily singular in a long-term dynamically sufficient phenotypic adaptive topography entails that evolutionary equilibria depend on development and consequently on gene-gene and gene-environment interactions. Hence, gene-gene and gene-environment interaction can generally have strong and permanent evolutionary effects in the sense of defining together with selection what the evolutionary equilibria are (e.g., via developmental feedbacks described by 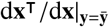 even by altering the constraining matrix alone. This contrasts with a non-singular constraining matrix whereby evolutionary equilibria are pre-determined by selection.

Second, our results allow for the study of long-term evolution of the constraining matrix as an emergent property of the evolution of the genotype, phenotype, and environment (i.e., the geno-envo-phenotype). In contrast, it has been traditional to study short-term evolution of **G** by treating it as another dynamic variable under constant allele frequency (Bulmer, 1971; Lande, 1979; Bulmer, 1980; Lande, 1980; Lande and Arnold, 1983; Barton and Turelli, 1987; Turelli, 1988; Gavrilets and Hastings, 1994; Carter et al., 2005; Débarre et al., 2014). Third, our results allow for the study of the effects of developmental bias, biased genetic variation, and modularity (Wagner, 1996; Pavlicev and Hansen, 2011; Pavlicev et al., 2011; Wagner and Zhang, 2011; Pavlicev and Wagner, 2012; Watson et al., 2013). While we have assumed that mutation is unbiased for the genotype, our equations allow for the developmental map to bias the phenotype distribution and the genetic variation of the phenotype. This may lead to modular effects of mutations, whereby altering a genotypic trait at a given age tends to affect some phenotypes but not others.

Fourth, our equations facilitate the study of life-history models with dynamic constraints. Life-history models with dynamic constraints have typically assumed evolutionary equilibrium, so they are analyzed using dynamic optimization techniques such as dynamic programming and optimal control (e.g., León, 1976; Iwasa and Roughgarden, 1984; Houston and McNamara, 1999; González-Forero et al., 2017; Avila et al., 2021). In recent years, mathematically modeling the evolutionary dynamics of life-history models with dynamic constraints, that is, of what we call the evo-devo dynamics, has been made possible with the canonical equation of adaptive dynamics for function-valued traits (Dieckmann et al., 2006; Parvinen et al., 2013; Metz et al., 2016). However, such an approach poses substantial mathematical challenges by requiring derivation of functional derivatives and solution of associated differential equations for costate variables (Parvinen et al., 2013; Metz et al., 2016; Avila et al., 2021). By using discrete age, we have obtained closed-form equations that facilitate modeling the evo-devo dynamics. By doing so, our framework yields an alternative method to dynamic optimization to analyze a broad class of life-history models with dynamic constraints (Example and González-Forero 2023).

Fifth, our framework allows for the modeling of the evo-devo dynamics of pattern formation by allowing the implementation of reaction-diffusion equations in *discrete space* in the developmental map, once equations are suitably written (e.g., Eq. 6.1 of Turing, 1952; Tomlin and Axelrod, 2007; Deutsch and Dormann, 2017; SI section S1.2). Thus, the framework may allow one to implement and analyze the evo-devo dynamics of existing detailed models of the development of morphology (e.g., Salazar-Ciudad and Jernvall, 2010; Sheth et al., 2012), to the extent that developmental maps can be written in the form of Eq. (1). Sixth, our framework allows for the mechanistic modeling of adaptive plasticity, for instance, by implementing reinforcement learning or supervised learning in the developmental map (Sutton and Barto, 2018; Paenke et al., 2007).

Our approach might also enable modelling developmental innovation, whereby fully formed traits emerge de novo in an individual, such as an extra digit (Goldschmidt, 1940; Gould, 1977; Orr and Coyne, 1992; Orr, 2005; Müller, 2010). The developmental constraint (1) admits that a slight perturbation in the geno-envo-phenotype at an early age yields a large change in the phenotype at a later age, possibly changing it from zero to an appreciable value. This may be used to model developmental innovation, possibly via exploratory processes described by Gerhart and Kirschner (2007) and Kirschner and Gerhart (2010) provided that a mathematical model of such processes satisfies Eq. 1. However, slight perturbations yielding large phenotypic effects might violate our assumption that invasion implies fixation. It has previously been established that invasion implies fixation if mutant *genotypes* **y** do not deviate substantially from resident genotypes 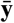 (Geritz et al., 2002; Geritz, 2005; Dieckmann et al., 2006; Priklopil and Lehmann, 2020; Avila et al., 2021), which we assume. Application of our framework to model developmental innovation will require explicitly determining whether large deviations in mutant phenotypes in our sense of the word still entail that invasion implies fixation because of small deviations in mutant genotypes.

By allowing development to be social, our framework allows for a mechanistic description of extra-genetic inheritance and indirect genetic effects. Extra-genetic inheritance can be described since the phenotype at a given age can be an identical or modified copy of the geno-phenotype of social partners. Thus, social development allows for the modeling of social learning (Sutton and Barto, 2018; Paenke et al., 2007) and epigenetic inheritance (Jablonka et al., 1992; Slatkin, 2009; Day and Bonduriansky, 2011). However, in our framework extra-genetic inheritance is insufficient to yield phenotypic evolution that is independent of both genetic evolution and exogenous plastic change (e.g., in the framework, there cannot be cultural evolution without genetic evolution or exogenous environmental change). This is seen by setting mutational covariation and exogenous environmental change to zero (i.e., **H**_**y**_ = **0** and 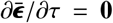), which eliminates evolutionary change (i.e., 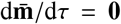). The reason seems to be that although there is extra-genetic *inheritance* in our framework, there is no extra-genetic *variation* because both development is deterministic and we use adaptive dynamics assumptions: without mutation, every SDS resident develops the same phenotype as every other resident. Extensions to consider stochastic development might enable extra-genetic variation and possibly phenotypic evolution that is independent of genetic and exogenously plastic evolution. Also, we have only considered social interactions among non-relatives, so our framework at present only allows for social learning or epigenetic inheritance from non-relatives.

Our framework can mechanistically describe indirect genetic effects via social development because the developed phenotype can be mechanistically influenced by the genotype or phenotype of social partners. Indirect genetic effects mean that a phenotype may be partly or completely caused by genes located in another individual (Moore et al., 1997). Indirect genetic effect approaches model the phenotype considering a linear regression of individual’s phenotype on social partner’s phenotype (Kirkpatrick and Lande, 1989; Moore et al., 1997; Townley and Ezard, 2013), whereas our approach constructs individual’s phenotype from development depending on social partners’ genotype and phenotypes. We found that social development generates social feedback (described by 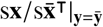Eq. 27), which closely though not entirely corresponds to social feedback found in the indirect genetic effects literature (Moore et al., 1997, Eq. 19b and subsequent text). The social feedback we obtain depends on total social developmental bias from the phenotype (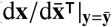Eq. 28); analogously, social feedback in the indirect genetic effects literature depends on the matrix of interaction coefficients (Ψ) which contains the regression coefficients of phenotype on social partner’s phenotype. Social development leads to a generalization of mechanistic additive genetic covariance matrices **H** = cov[**b, b**] into mechanistic additive socio-genetic cross-covariance matrices **L** = cov[**b**^s^, **b**]; similarly, indirect genetic effects involve a generalization of the **G**-matrix, which includes **C**_**ax**_ = cov[**a, x**], namely the cross-covariance matrix between multivariate breeding value and phenotype (Kirkpatrick and Lande, 1989; Moore et al., 1997; Townley and Ezard, 2013).

However, there are differences between our results and those in the indirect genetic effects literature. First, social feedback (in the sense of inverse matrices involving Ψ) appears twice in the evolutionary dynamics under indirect genetic effects (see Eqs. 20 and 21 of Moore et al. 1997) while it only appears once in our evolutionary dynamics equations through 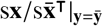,(Layer 6, Eq. 10). This difference may stem from the assumption in the indirect genetic effects literature that social interactions are reciprocal, while we assume that they are asymmetric in the sense that, since mutants are rare, mutant’s development depends on residents but resident’s development does not depend on mutants (we thank J. W. McGlothlin for pointing this out). Second, our **L** matrices make the evolutionary dynamics equations depend on total social developmental bias from the genotype (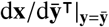, Eq. 26) in a non-feedback manner (specifically, not in an inverse matrix) but this type of dependence does not occur in the evolutionary dynamics under indirect genetic effects (Eqs. 20 and 21 of Moore et al. 1997). This difference might stem from the absence of explicit tracking of allele frequency in the indirect genetic effects literature in keeping with the tradition of quantitative genetics, whereas we explicitly track genotypic traits. Third, “social selection” (i.e., 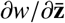) plays no role in our results consistently with our assumption of a well-mixed population, but social selection plays an important role in the indirect genetic effects literature even if relatedness is zero (McGlothlin et al., 2010, e.g., setting *r* = 0 in their Eq. 10 still leaves an effect of social selection on selection response due to “phenotypic” kin selection).

Our framework offers formalizations to the notions of developmental constraints and developmental bias. The two notions have been often interpreted as equivalents (e.g., Brakefield, 2006), or with a distinction such that constraints entail a negative, prohibiting effect while bias entails a positive, directive effect of development on the generation of phenotypic variation (Uller et al., 2018; Salazar-Ciudad, 2021). We defined developmental constraint as the condition that the phenotype at a given age is a function of the individual’s condition at their immediately previous age, which both prohibits certain values of the phenotype and has a “directive” effect on the generation of phenotypic variation. We offered quantification of developmental bias in terms of the slope of the phenotype with respect to itself at subsequent ages. No bias would lead to zero slopes thus to identity matrices (e.g., 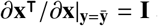 and 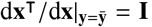) and deviations from the identity matrix would constitute bias.

Our results clarify the role of several developmental factors previously suggested to be evolutionarily important. We have arranged the evo-devo process in a layered structure, where a given layer is formed by components of layers below (Fig. 4). This layered structure helps see that several developmental factors previously suggested to have important evolutionary effects (Laland et al., 2014) but with little clear connection (Welch, 2017) can be viewed as basic elements of the evolutionary process. Direct-effect matrices (Layer 2) are basic in that they form all the components of the evolutionary dynamics (Layer 7) except mutational covariation and exogenous environmental change. Direct-effect matrices quantify direct (i) directional selection, (ii) developmental bias, (iii) niche construction, (iv) social developmental bias (e.g., extra-genetic inheritance and indirect genetic effects; Moore et al. 1997), (v) social niche construction, (vi) environmental sensitivity of selection (Chevin et al., 2010), and (vii) phenotypic plasticity. These factors variously affect selection and development, thus affecting evolutionary equilibria and the admissible evolutionary trajectory.

Our approach uses discrete rather than continuous age, which substantially simplifies the mathematics. This treatment allows for the derivation of closed-form expressions for what can otherwise be a difficult mathematical challenge if age is continuous (Kirkpatrick and Heckman, 1989; Dieckmann et al., 2006; Parvinen et al., 2013; Metz et al., 2016; Avila et al., 2021). For instance, costate variables are key in dynamic optimization as used in life-history models (Gadgil and Bossert, 1970; León, 1976; Schaffer, 1983; Stearns, 1992; Roff, 1992; Kozłowski and Teriokhin, 1999; Sydsæter et al., 2008), but general closed-form formulas for costate variables were previously unavailable and their calculation often limits the analysis of such models. In SI section S4, we show that our results recover the key elements of Pontryagin’s maximum principle, which is a central tool of optimal control theory to analytically solve dynamic optimization problems (Sydsæter et al., 2008). Under the assumption that there are no environmental traits (hence, no exogenous plastic response), in SI section S4, we show that an admissible locally stable evolutionary equilibrium solves a local, dynamic optimization problem of finding a genotype that both “totally” maximises a mutant’s lifetime reproductive success *R*_0_ and “directly” maximises the Hamiltonian of Pontryagin’s maximum principle. We show that this Hamiltonian depends on costate variables that are proportional to the total selection gradient of the phenotype at evolutionary equilibrium (Eq. S4.3), and that the costate variables satisfy the costate equations of Pontryagin’s maximum principle. Thus, our approach offers an alternative method to optimal control theory to find admissible evolutionary equilibria for the broad class of models considered here. By exploiting the discretization of age, we have obtained various formulas that can be computed directly for the total selection gradient of the phenotype (Layer 4, Eq. S20), so for costate variables, and of their relationship to total genotypic selection (fifth line of Layer 4, Eq. S21), thus facilitating analytic and numerical treatment of life-history models with dynamic constraints. Although discretization of age may induce numerical imprecision relative to continuous age (Kirkpatrick and Heckman, 1989), numerical and empirical treatment of continuous age typically involves discretization at one point or another, with continuous curves often achieved by interpolation (e.g., Kirkpatrick et al., 1990; Rao, 2009). Numerical precision with discrete age may be increased by reducing the age bin size (e.g., to represent months or days rather than years; Caswell, 2001), at a computational cost.

By simplifying the mathematics, our approach yields insight that may be otherwise challenging to gain. Life-history models with dynamic constraints generally find that costate variables are non-zero under optimal controls (Gadgil and Bossert, 1970; Taylor et al., 1974; León, 1976; Schaffer, 1983; Houston et al., 1988; Houston and McNamara, 1999; Sydsæter et al., 2008). This means that there is persistent total selection on the phenotype at evolutionary equilibrium. Our findings show that this is to be expected for various reasons including if there are absolute mutational constraints (i.e., active path constraints so controls remain between zero and one, as in the Example), direct genotypic selection, or more state variables than control variables (in which case δ**x**^⊺^/δ**y** is singular as it has more rows than columns, even after removing initial states and final controls from the analysis; Eq. S5.2.10) (fifth line of Layer 4, Eq. S21). Thus, zero total genotypic selection at equilibrium may involve persistent total phenotypic selection. Moreover, life-history models with explicit developmental constraints have found that their predictions can be substantially different from those found without explicit developmental constraints. In particular, without developmental constraints, the outcome of parent-offspring conflict over sex allocation has been found to be an intermediate between the outcomes preferred by mother and offspring (Reuter and Keller, 2001), whereas with developmental constraints, the outcome has been found to be that preferred by the mother (Avila et al., 2019). Our results show that changing the particular form of the developmental map may induce substantial changes in predictions by influencing total genotypic selection and the admissible evolutionary equilibria. In other words, the developmental map used alters the evolutionary outcome because it modulates absolute socio-genetic constraints (i.e., the **H** or **L** matrices in geno-phenotype space).

We have obtained a term that we refer to as exogenous plastic response, which is the plastic response to exogenous environmental change over an evolutionary time step (Layer 7, Eq. 3). An analogous term occurs in previous equations (Eq. A3 of Chevin et al. 2010). Additionally, our framework considers *endogenous* plastic response due to niche construction (i.e., endogenous environmental change), which affects both the selection response and the exogenous plastic response. Exogenous plastic response affects the evolutionary dynamics even though it is not ultimately caused by change in the resident genotype (or in gene frequency), but by exogenous environmental change. In particular, exogenous plastic response allows for a straightforward form of “plasticity-first” evolution (Waddington, 1942, 1961; West-Eberhard, 2003) as follows. At an evolutionary equilibrium where exogenous plastic response is absent, the introduction of exogenous plastic response generally changes socio-genetic covariation or directional selection at a subsequent evolutionary time, thereby inducing selection response. This constitutes a simple form of plasticity-first evolution, whereby plastic change precedes genetic change, although the plastic change may not be adaptive and the induced genetic change may have a different direction to that of the plastic change.

Empirical estimation of the developmental map may be facilitated by it defining a dynamic equation. Whereas the developmental map defines a dynamic equation to construct the phenotype, the genotype-phenotype map corresponds to the solution of such dynamic equation. It is often impractical or impossible to write the solution of a dynamic equation, even if the dynamic equation can be written in practice. Accordingly, it may often prove impractical to empirically estimate the genotype-phenotype map, whereas it may be more tractable to empirically infer developmental maps. Inference of developmental maps from empirical data can be pursued via the growing number of methods to infer dynamic equations from data (Schmidt and Lipson, 2009; Brunton et al., 2016; Ghadami and Epureanu, 2022; Course and Nair, 2023).

To conclude, we have formulated a framework that synthesizes developmental and evolutionary dynamics yielding a theory of long-term phenotypic evolution on an adaptive topography, which also mechanistically describes the long-term evolution of genetic covariation. This framework shows that development has major evolutionary effects by finding that selection and development jointly define the evolutionary outcomes if mutation is not absolutely constrained and exogenous plastic response is absent, rather than the outcomes being defined only by selection. Our results provide a tool to chart major territory on how development affects evolution.

## Supporting information

Computer Code

## 8. Acknowledgements

I thank Andy Gardner for extensive support throughout this project, by discussing, reading, and commenting on many early drafts, and generously offering interpretation, advice, and funding. I thank K.N. Laland, R. Lande, L.C. Mikula, A.J. Moore, and M.B. Morrissey for comments on previous versions of the manuscript, and A. Krause, J.W. McGlothlin, J.A.J. Metz, L. Milocco, I. Salazar-Ciudad, and D.M. Shuker for discussion. I thank T.F. Hansen, M. Pavlicev, S.J. Schreiber, and three anonymous reviewers for comments that helped to greatly improve the manuscript. I thank M.B. Morrissey for discussion and explanation of his work. This work was funded by an ERC Consolidator Grant to A. Gardner (grant no. 771387), by the School of Biology of the University of St Andrews, and by a John Templeton Foundation grant to K.N. Laland and T. Uller (grant ID 60501). A preprint of the manuscript is available at https://www.biorxiv.org/content/10.1101/2021.05.17.444499.

## Appendix A Matrix calculus notation

Following Caswell (2019), for vectors **a** ∈ ℝ^*n*×1^ and **b** ∈ ℝ^*m*×1^, we denote

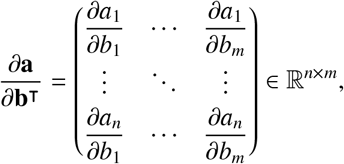

so (∂**a**/∂**b**^⊺^)^⊺^ = ∂**a**⊺/∂**b**. The same notation applies with total derivatives.

