## Supplementary Information for "A mathematical framework for evo-devo dynamics"

### Supplementary information for:
### A mathematical framework for evo-devo dynamics

Mauricio González-Forero[1]

[1]School of Biology, University of St Andrews, Dyers Brae, St Andrews, KY16 9TH, Fife, UK

**S4 Connection to dynamic optimization** **19**

**S5 Derivation of results** **21**
S5.1 Total selection gradient of the phenotype . . 21
S5.1.1 Total selection gradient of the phenotype in terms of direct fitness effects 21

### Outline

This Supplementary Information contains the following elements. Section S1 provides further discussion of how our approach relates to closely related previous work and of how the developmental map admits reaction-diffusion models. Sections S2 presents the model in detail. In particular, section S2 includes a derivation under our assumptions, including deterministic population dynamics, of the evolutionary dynamics of the genotype, namely, the canonical equation for adaptive dynamics, which depends on the total selection gradient of the genotype. Section S3 describes the results for the bottom layers of the evo-devo process, namely, layers 1-5 (the results for the top layers 6-7 are presented in the main text). Section S4 shows how our approach relates to dynamic optimization, specifically, to optimal control. Finally, section S5 presents the derivation of all the results. More specifically, in section S5, we derive equations for the total selection gradient of the phenotype, genotype, environment, geno-phenotype, and geno-envo-phenotype, and equations for the evolutionary dynamics of the phenotype, environment, geno-phenotype, and geno-envo-phenotype.

### S1 Further discussion

#### S1.1 Relationship to closely related work

Our approach to describe the evo-devo dynamics by an evolutionary dynamic equation of genetic traits (Eq. 3) and a developmentally dynamic equation of developed traits (Eq. 1) further relates to previous work as follows. Because of the multivariate age structure of $\mathbf{y}$, Eq. (3) can be seen as a canonical equation for multivariate discrete function-valued traits. Dieckmann *et al.* (2006) derive the canonical equation for univariate continuous function-valued traits. Both Eq. (3) and the canonical equation of Dieckmann *et al.* (2006) are dynamic equations in gradient form for control variables $\bar{\mathbf{y}}$, but not for state variables $\bar{\mathbf{x}}$, thus leaving unanswered whether and to what extent the evolution of developed traits can be described as the climbing of a fitness landscape. Dieckmann *et al.* (2006, p. 378) note that the mutational covariance matrix in their canonical equation is singular if and only if the function-valued trait has equality constraints. As their canonical equation describes the evolutionary dynamics in gradient form for control variables, Dieckmann *et al.*'s (2006) point is that the mutational covariance matrix $\mathrm{cov}[\mathbf{y},\mathbf{y}]$ is singular if and only if $\mathbf{y}$ has equality constraints. Developmental constraints (1) are equality constraints on state variables $\mathbf{x}$, so Dieckmann *et al.*'s (2006) point suggests that an analogous singularity would appear in the covariance matrices involved in a canonical equation describing the evolutionary dynamics of state variables in gradient form, although such canonical equation has not been derived to our knowledge (and we show it to have a different form to that of Eq. 3). Parvinen *et al.* (2013) and Metz *et al.* (2016) derive the total selection gradient $\mathrm{d}\lambda/\mathrm{d}\mathbf{y}|_{\mathbf{y}=\bar{\mathbf{y}}}$ for specific models for a univariate function-valued control with continuous domain where $\lambda$ depends on a multivariate state variable with continuous domain subject to developmental constraints analogous to those in Eq. (1) in continuous age (Parvinen et al's Eq. 2a). Avila *et al.* (2021) derive $\mathrm{d}\lambda/\mathrm{d}\mathbf{y}|_{\mathbf{y}=\bar{\mathbf{y}}}$ for a broad class of models for a univariate control with continuous domain where $\lambda$ depends on a univariate state variable with continuous domain in a group-structured population (their Eqs. 7,23,24); the resulting equation depends on an unknown univariate costate variable, which at evolutionary equilibrium can be calculated by solving an associated differential equation (their Eq. 32). Yet, calculating the selection gradient $\mathrm{d}\lambda/\mathrm{d}\mathbf{y}|_{\mathbf{y}=\bar{\mathbf{y}}}$ when invasion fitness depends on state variables in continuous age remains challenging as functional derivatives, integrals, and associated differential equations must be computed. We circumvent these difficulties by using discrete age, so calculations reduce to basic derivatives and matrix algebra. This enables us to derive simple expressions that can be calculated directly for both the total selection gradient of controls and the evolutionary dynamic equations in gradient form for state variables subject to dynamic constraints.

#### S1.2 Reaction-diffusion in the developmental constraint

Here we illustrate how the developmental constraint (1) admits spatially explicit reaction-diffusion models.

Let $t \in \mathbb{R}$ be time, $x \in \mathbb{R}$ be unidimensional space, $u(t,x) \in \mathbb{R}$ be the concentration of a chemical at time $t$ and location $x$, $D$ be a diffusion constant, and $f(u)$ be a reaction function. Consider then a Turing reaction-diffusion partial differential equation in unidimensional space of the form

$$\frac{\partial u(t,x)}{\partial t} = f(u) + D\frac{\partial^2 u}{\partial x^2}. \qquad \text{(Eq. S1.2.1)}$$

We may write this equation in the form of the developmental constraint (1) as follows.

First, we discretize time and space. To do this, let $v(t,x) = u(t,x+1) - u(t,x)$. Then, Eq. S1.2.1 in discrete time and space is

$$\begin{aligned} &u(t+1,x) - u(t,x) \\ &= f(u(t,x)) + D[v(t,x+1) - v(t,x)] \\ &= f(u(t,x)) + D\{[u(t,x+2) - u(t,x+1)] \\ &\qquad\qquad - [u(t,x+1) - u(t,x)]\} \\ &= f(u(t,x)) + D[u(t,x+2) - 2u(t,x+1) + u(t,x)]. \end{aligned} \qquad \text{(Eq. S1.2.2)}$$

This corresponds to Turing's (1952) Eq. 6.1, which uses discrete space (cell) but continuous time.

Second, we write this in our notation. Let $x_{ia}$ be the concentration of the chemical at age $a \in \{1,\ldots,N_\mathrm{a}\}$ at spatial location $i \in \{1,\ldots,N_\mathrm{p}\}$. Then, rearranging Eq. S1.2.2 and using our notation, the developmental constraint for the concentration of the chemical is

$$\begin{aligned} x_{i,a+1} &= g_{ia}(\mathbf{x}_a) \\ &= x_{ia} + f(x_{ia}) + D[x_{i+2,a} - 2x_{i+1,a} + x_{ia}], \end{aligned} \qquad \text{(Eq. S1.2.3a)}$$

for $i \in \{1,\dots,N_\mathrm{p}-2\}$ where $\mathbf{x}_a = (x_{1a},\dots,x_{N_\mathrm{p}a})^\intercal$. Evaluating this equation at the spatial locations $N_\mathrm{p}$ and $N_\mathrm{p}-1$ we find the developmental constraint at these locations:

$$\begin{aligned} x_{N_\mathrm{p},a+1} &= g_{N_\mathrm{p}a}(\mathbf{x}_a) \\ &= x_{N_\mathrm{p}a} + f(x_{N_\mathrm{p}a}) + Dx_{N_\mathrm{p}a}, \qquad \text{(Eq. S1.2.3b)} \\ x_{N_\mathrm{p}-1,a+1} &= g_{N_\mathrm{p}-1,a}(\mathbf{x}_a) \\ &= x_{N_\mathrm{p}-1,a} + f(x_{N_\mathrm{p}-1,a}) + D[-2x_{N_\mathrm{p}a} + x_{N_\mathrm{p}-1,a}], \qquad \text{(Eq. S1.2.3c)} \end{aligned}$$

where we let $x_{N_\mathrm{p}+2,a} = 0$ and $x_{N_\mathrm{p}+1,a} = 0$. Eq. S1.2.3 describe a reaction-diffusion mechanism in unidimensional space in the form of the developmental constraint (1).

A reaction-diffusion system in multidimensional space may be similarly written in the form of the developmental constraint (1) by mapping the discrete spatial vector into a discrete scalar $i$.

### S2 Model

In this section, we describe our methods in detail. It consists of the following subsections. First, we formally describe the three phases in which we divide an evolutionary time step. We address the complication introduced by social development by adding a phase to the standard separation of time scales in adaptive dynamics. Thus, we divide an evolutionary time step in three phases rather than the usual two of resident population dynamics and mutant invasion dynamics; analogues of such additional phase have been used in modeling the evolution of social learning (Aoki *et al.*, 2012; Kobayashi *et al.*, 2015). Second, we obtain a first-order approximation of invasion fitness and use it to derive the canonical equation describing the evolutionary dynamics of the genotype. Third, we derive the selection gradient in age structured populations, which we use to calculate the total selection gradient of the genotype. Fourth, we derive the expressions for the forces of selection involved in the selection gradient for age structured populations, and expressions for the selection gradient in terms of lifetime reproductive success. Fifth, we derive the canonical equation of adaptive dynamics under our assumptions, in particular, deterministic population dynamics.

#### S2.1 Phases of the evolutionary cycle

We now formally describe the three phases in which we partition an evolutionary time step (Fig. 3). We start with the socio-devo stabilization dynamics phase, which yields the notions of socio-devo equilibrium and socio-devo stability.

##### S2.1.1 Socio-devo stabilization dynamics phase

Socio-devo stabilization dynamics occur as follows. For a resident geno-envo-phenotype $\bar{\mathbf{m}} = (\bar{\mathbf{x}};\bar{\mathbf{y}};\boldsymbol{\epsilon})$, a new resident phenotype is obtained using the developmental (Eq. 1) and environmental (Eq. 2) constraints; the resulting phenotype, its genotype, and the resulting environment are set as the new resident; and this is iterated. To write this formally, let $\theta$ denote time for the socio-devo stabilization dynamics (we do not explicitly address how socio-devo time $\theta$ relates to ecological time $t$). During the socio-devo stabilization phase, denote the resident phenotype at socio-devo time $\theta$ as $\bar{\mathbf{x}}(\theta)$. Then, using the developmental (Eq. 1) and environmental (Eq. 2) constraints, the resident phenotype at socio-devo time $\theta+1$ is given by

$$\begin{aligned} &\bar{\mathbf{x}}_{a+1}(\theta+1) \\ &\quad = \mathbf{g}_a(\bar{\mathbf{x}}_a(\theta+1),\bar{\mathbf{y}}_a,\mathbf{h}_a(\bar{\mathbf{x}}_a(\theta+1),\bar{\mathbf{y}}_a,\bar{\mathbf{x}}(\theta),\bar{\mathbf{y}},\tau),\bar{\mathbf{x}}(\theta),\bar{\mathbf{y}}), \qquad \text{(Eq. S2.1.1)} \end{aligned}$$

for all $a \in \{1,\dots,N_\mathrm{a}-1\}$ and with given initial conditions $\bar{\mathbf{x}}(1)$ and $\bar{\mathbf{x}}_1(\theta+1) = \bar{\mathbf{x}}_1$. If $\lim_{\theta\to\infty}\bar{\mathbf{x}}(\theta)$ converges, then this limit, the resident genotype, and the resulting environment yield a socio-devo stable geno-envo-phenotype as defined below.

We say a geno-envo-phenotype $\bar{\mathbf{m}} = (\bar{\mathbf{x}};\bar{\mathbf{y}};\bar{\boldsymbol{\epsilon}})$ is a socio-devo equilibrium if and only if $\bar{\mathbf{x}}$ is produced by development when the individual has such genotype $\bar{\mathbf{y}}$ and everyone else in the population has that same genotype, phenotype, and environment; specifically, a socio-devo equilibrium $\bar{\mathbf{m}} = (\bar{\mathbf{x}};\bar{\mathbf{y}};\bar{\boldsymbol{\epsilon}})$ satisfies

$$\begin{aligned} \bar{\mathbf{x}}_{a+1} &= \mathbf{g}_a(\bar{\mathbf{x}}_a,\bar{\mathbf{y}}_a,\bar{\boldsymbol{\epsilon}}_a,\bar{\mathbf{x}},\bar{\mathbf{y}}) \quad \forall a \in \{1,\dots,N_\mathrm{a}-1\}, \bar{\mathbf{x}}_1 \text{ fixed}, \\ \bar{\boldsymbol{\epsilon}}_a &= \mathbf{h}_a(\bar{\mathbf{x}}_a,\bar{\mathbf{y}}_a,\bar{\mathbf{x}},\bar{\mathbf{y}},\tau) \quad \forall a \in \{1,\dots,N_\mathrm{a}\}. \qquad \text{(Eq. S2.1.2)} \end{aligned}$$

We assume that there is at least one socio-devo equilibrium for a given developmental map at every evolutionary time $\tau$.

If the resident geno-envo-phenotype is a socio-devo equilibrium, from Eq. (1), Eq. (2), and Eq. S2.1.2, it follows that evaluation of the mutant genotype at the resident genotype yields resident variables. That is, if $\bar{\mathbf{m}} = (\bar{\mathbf{x}};\bar{\mathbf{y}};\bar{\boldsymbol{\epsilon}})$ is a socio-devo equilibrium, then

$$\begin{aligned} \mathbf{x}|_{\mathbf{y}=\bar{\mathbf{y}}} &= \bar{\mathbf{x}} \\ \boldsymbol{\epsilon}|_{\mathbf{y}=\bar{\mathbf{y}}} &= \bar{\boldsymbol{\epsilon}} \\ \mathbf{z}|_{\mathbf{y}=\bar{\mathbf{y}}} &= \bar{\mathbf{z}} \\ \mathbf{m}|_{\mathbf{y}=\bar{\mathbf{y}}} &= \bar{\mathbf{m}}. \end{aligned}$$

More specifically, if the resident geno-envo-phenotype is a socio-devo equilibrium, the resident phenotype at age $a+1$ is given by Eq. (1) evaluating the developmental map at the resident geno-envo-phenotype (i.e., $\bar{\mathbf{x}}_{a+1} = \mathbf{g}_a(\bar{\mathbf{m}}_a,\bar{\mathbf{z}})$), and the resident environment at age $a$ is given by Eq. (2) evaluating the environmental map at the resident geno-phenotype (i.e., $\bar{\boldsymbol{\epsilon}}_a = \mathbf{h}_a(\bar{\mathbf{z}}_a,\bar{\mathbf{z}},\tau)$).

We say that a geno-envo-phenotype $\bar{\mathbf{m}} = (\bar{\mathbf{x}};\bar{\mathbf{y}};\bar{\boldsymbol{\epsilon}})$ is socio-devo stable (SDS) if and only if $\bar{\mathbf{m}}$ is a locally stable socio-devo equilibrium. A socio-devo equilibrium $\bar{\mathbf{m}} = (\bar{\mathbf{x}};\bar{\mathbf{y}};\bar{\boldsymbol{\epsilon}})$ is locally stable if and only if a marginally small deviation in the initial phenotype $\bar{\mathbf{x}}(1)$ from the socio-devo equilibrium keeping the same genotype leads the socio-devo stabilization dynamics to the same equilibrium. Thus, a socio-devo equilibrium $\bar{\mathbf{m}}$ is locally stable if all the eigenvalues of the matrix

$$\left.\frac{\mathrm{d}\mathbf{x}}{\mathrm{d}\bar{\mathbf{x}}^{\intercal}}\right|_{\mathbf{y}=\bar{\mathbf{y}}} \qquad \text{(Eq. S2.1.3)}$$

have absolute value (or modulus) strictly less than one. The requirement that this matrix has such eigenvalues arises naturally in the derivation of the evolutionary dynamics of the resident phenotype $\bar{\mathbf{x}}$ (section S5.6). We assume that there is a unique SDS geno-envo-phenotype for a given developmental map at every evolutionary time $\tau$.

To see why the matrix in Eq. S2.1.3 is sufficient to determine socio-devo stability, consider the following. Let $\bar{\mathbf{x}}(\theta+1) = \tilde{\mathbf{g}}(\bar{\mathbf{x}}(\theta))$ denote the solution of iterating Eq. S2.1.1 over $a$, where we highlight only the argument corresponding to the phenotype of social partners. An equilibrium $\bar{\mathbf{x}}^{**}$ of the socio-devo stabilization dynamics satisfies $\bar{\mathbf{x}}^{**} = \tilde{\mathbf{g}}(\bar{\mathbf{x}}^{**})$. Taylor-expanding $\bar{\mathbf{x}}(\theta+1)$ to first-order around $\bar{\mathbf{x}}^{**}$, we have

$$\begin{aligned}\bar{\mathbf{x}}(\theta+1) &= \tilde{\mathbf{g}}(\bar{\mathbf{x}}^{**}) \\ &\quad + \left.\frac{\mathrm{d}\tilde{\mathbf{g}}}{\mathrm{d}\bar{\mathbf{x}}^{\intercal}}\right|_{\bar{\mathbf{x}}=\bar{\mathbf{x}}^{**}} \left(\bar{\mathbf{x}}(\theta) - \bar{\mathbf{x}}^{**}\right) + O(||\bar{\mathbf{x}}(\theta) - \bar{\mathbf{x}}^{**}||^2),\end{aligned}$$

where the operator $\mathrm{d}/\mathrm{d}\bar{\mathbf{x}}^{\intercal}$ takes the total derivative to take into account developmental and environmental constraints. Noting that $\mathrm{d}\tilde{\mathbf{g}}/\mathrm{d}\bar{\mathbf{x}}^{\intercal}|_{\bar{\mathbf{x}}=\bar{\mathbf{x}}^{**}} = \mathrm{d}\mathbf{x}/\mathrm{d}\bar{\mathbf{x}}^{\intercal}|_{\mathbf{y}=\bar{\mathbf{y}}}$ since the resident is a socio-devo equilibrium, we have that a perturbation from a socio-devo equilibrium is approximately

$$\bar{\mathbf{x}}(\theta+1) - \tilde{\mathbf{g}}(\bar{\mathbf{x}}^{**}) \approx \left.\frac{\mathrm{d}\mathbf{x}}{\mathrm{d}\bar{\mathbf{x}}^{\intercal}}\right|_{\mathbf{y}=\bar{\mathbf{y}}} \left(\bar{\mathbf{x}}(\theta) - \bar{\mathbf{x}}^{**}\right),$$

which asymptotically converges to $\mathbf{0}$ (i.e., $\bar{\mathbf{x}}^{**}$ is locally stable) if all the eigenvalues of the matrix in Eq. S2.1.3 have absolute value (or modulus) strictly less than one.

##### S2.1.2 Resident population dynamics phase

Once the SDS resident is reached (or, more strictly, sufficiently approached) in the socio-devo stabilization phase, we continue to the resident population dynamics phase (Fig. 3). Let the resident geno-envo-phenotype $\bar{\mathbf{m}}$ be SDS. Let $\bar{n}_a(t)$ denote the population density of SDS residents of age $a \in \{1,\ldots,N_\mathrm{a}\}$ at ecological time $t$. The vector of resident density at $t$ is $\bar{\mathbf{n}}(t) = (\bar{n}_1(t),\ldots,\bar{n}_{N_\mathrm{a}}(t))^{\intercal}$. The life cycle is age-structured (Fig. S1). At age $a$, an SDS resident produces a number $A_{1a}(\bar{\mathbf{m}}_a,\bar{\mathbf{z}},\bar{\mathbf{n}}(t))$ of offspring and survives to age $a+1$ with probability $A_{a+1,a}(\bar{\mathbf{m}}_a,\bar{\mathbf{z}},\bar{\mathbf{n}}(t))$ (where we set $A_{N_\mathrm{a}+1,N_\mathrm{a}}(\bar{\mathbf{m}}_a,\bar{\mathbf{z}},\bar{\mathbf{n}}(t)) = 0$). The first argument of these two functions is the geno-envo-phenotype of the individual at that age, the second argument is the geno-phenotype of the individual's social partners who can be of any age, and the third argument is density dependence; thus, an individual's fertility and survival directly depend on the individual's local environment (via the first argument $\bar{\mathbf{m}}_a$) but not directly on the local environment of social partners (the second argument $\bar{\mathbf{z}}$ does not include the environment). These expressions for survival and fertility use the assumption that exogenous environmental change is slow so $\bar{\mathbf{m}}$ is constant with respect

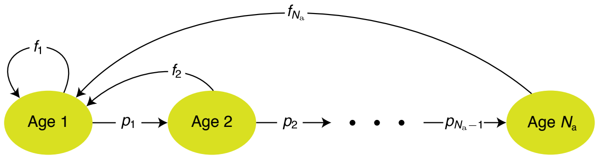

Figure S1: Age-structured life cycle. The vital rates shown are those of rare mutants: a mutant of age $a$ produces $f_a$ offspring and survives to age $a+1$ with probability $p_a$. See text for the vital rates of the resident.

to the ecological time $t$. The SDS resident population thus has deterministic dynamics given by

$$\bar{\mathbf{n}}(t+1) = \mathbf{A}(\bar{\mathbf{m}},\bar{\mathbf{z}},\bar{\mathbf{n}}(t))\bar{\mathbf{n}}(t), \qquad \text{(Eq. S2.1.4)}$$

where $\mathbf{A}(\bar{\mathbf{m}},\bar{\mathbf{z}},\bar{\mathbf{n}}(t))$ is a density-dependent Leslie matrix whose entries $A_{ij}(\bar{\mathbf{m}}_j,\bar{\mathbf{z}},\bar{\mathbf{n}}(t))$ give the age-specific survival probabilities and fertilities of SDS resident individuals; the first argument of $\mathbf{A}(\bar{\mathbf{m}},\bar{\mathbf{z}},\bar{\mathbf{n}}(t))$ is the geno-envo-phenotype vector formed by the first argument of $A_{ij}(\bar{\mathbf{m}}_j,\bar{\mathbf{z}},\bar{\mathbf{n}}(t))$ for all $i,j \in \{1,\ldots,N_\mathrm{a}\}$. We assume that density dependence is such that the population dynamics of the SDS resident (Eq. S2.1.4) have a unique stable non-trivial equilibrium $\bar{\mathbf{n}}^*(\bar{\mathbf{m}})$ (a vector of non-negative entries some of which are positive), which solves

$$\bar{\mathbf{n}}^*(\bar{\mathbf{m}}) = \mathbf{A}(\bar{\mathbf{m}},\bar{\mathbf{z}},\bar{\mathbf{n}}^*(\bar{\mathbf{m}}))\bar{\mathbf{n}}^*(\bar{\mathbf{m}}). \qquad \text{(Eq. S2.1.5)}$$

The carrying capacity is given by $\bar{n}^*(\bar{\mathbf{m}}) = \sum_{a=1}^{N_\mathrm{a}} \bar{n}_a^*(\bar{\mathbf{m}})$, which depends on the SDS resident geno-envo-phenotype. We assume that $\mathbf{A}(\bar{\mathbf{m}},\bar{\mathbf{z}},\bar{\mathbf{n}}^*(\bar{\mathbf{m}}))$ is primitive (i.e., raising $\mathbf{A}(\bar{\mathbf{m}},\bar{\mathbf{z}},\bar{\mathbf{n}}^*(\bar{\mathbf{m}}))$ to a sufficiently high power yields a matrix whose entries are all positive). For instance, $\mathbf{A}(\bar{\mathbf{m}},\bar{\mathbf{z}},\bar{\mathbf{n}}^*(\bar{\mathbf{m}}))$ is primitive if residents can survive to the last age with non-zero probability (i.e., $A_{a+1,a}(\bar{\mathbf{m}}_a,\bar{\mathbf{z}},\bar{\mathbf{n}}(t)) > 0$ for all $a \in \{1,\ldots,N_\mathrm{a}-1\}$), residents in the last age reproduce (i.e., $A_{1N_\mathrm{a}}(\bar{\mathbf{m}}_{N_\mathrm{a}},\bar{\mathbf{z}},\bar{\mathbf{n}}(t)) > 0$), and residents of at least two consecutive age classes have non-zero fertility (i.e., $A_{1a}(\bar{\mathbf{m}}_a,\bar{\mathbf{z}},\bar{\mathbf{n}}(t)) > 0$ and $A_{1,a+1}(\bar{\mathbf{m}}_{a+1},\bar{\mathbf{z}},\bar{\mathbf{n}}(t)) > 0$ for some $a \in \{1,\ldots,N_\mathrm{a}-1\}$) (Sternberg, 2010, section 9.4.1). Hence, from the Perron-Frobenius theorem (Sternberg, 2010, theorem 9.1.1), it follows that $\mathbf{A}(\bar{\mathbf{m}},\bar{\mathbf{z}},\bar{\mathbf{n}}^*(\bar{\mathbf{m}}))$ has an eigenvalue $\bar{\lambda} = 1$ that is strictly greater than the absolute value of any other eigenvalue of the matrix. This $\bar{\lambda}$ describes the asymptotic growth rate of the resident population, as the resident population dynamics equilibrium $\bar{\mathbf{n}}^*(\bar{\mathbf{m}})$ is achieved.

##### S2.1.3 Resident-mutant population dynamics phase

Once the resident population has reached (or, more strictly, sufficiently approached) the equilibrium $\bar{\mathbf{n}}^*(\bar{\mathbf{m}})$, we move on to the resident-mutant population dynamics phase (Fig. 3). After some time inversely proportional to $\mu(\bar{\mathbf{m}})n^*(\bar{\mathbf{m}})$, where $\mu(\bar{\mathbf{m}})$ is the mutation rate, a rare mutant genotype $\mathbf{y}$ arises, where $\mathbf{y}$ is a realization of a multivariate random variable. A mutant has geno-envo-phenotype $\mathbf{m} = (\mathbf{x};\mathbf{y};\boldsymbol{\epsilon})$ where the mutant phenotype $\mathbf{x}$ is given by the developmental constraint (Eq. 1) and the mutant environment $\boldsymbol{\epsilon}$ is given by the environmental constraint (Eq. 2).

Let $n_a(t)$ denote the density of mutant individuals of age $a \in \{1, \ldots, N_\mathrm{a}\}$ at ecological time $t$. The vector of mutant density at $t$ is $\mathbf{n}(t) = (n_1(t), \ldots, n_{N_\mathrm{a}}(t))^\intercal$. Given clonal reproduction, the population dynamics of the resident and rare mutant subpopulations are then given by the expanded system

$$\begin{pmatrix} \bar{\mathbf{n}}(t+1) \\ \mathbf{n}(t+1) \end{pmatrix} = \begin{pmatrix} \mathbf{A}(\bar{\mathbf{m}}, \bar{\mathbf{z}}, \bar{\mathbf{n}}(t)) & \mathbf{0} \\ \mathbf{0} & \mathbf{A}(\mathbf{m}, \bar{\mathbf{z}}, \bar{\mathbf{n}}(t)) \end{pmatrix} \begin{pmatrix} \bar{\mathbf{n}}(t) \\ \mathbf{n}(t) \end{pmatrix},$$

where the mutant projection matrix $\mathbf{A}(\mathbf{m}, \bar{\mathbf{z}}, \bar{\mathbf{n}}(t))$ is given by evaluating the first argument of $\mathbf{A}(\bar{\mathbf{m}}, \bar{\mathbf{z}}, \bar{\mathbf{n}}(t))$ at the mutant geno-envo-phenotype. Hence, $\mathbf{A}(\mathbf{m}, \bar{\mathbf{z}}, \bar{\mathbf{n}}(t))$ is a density-dependent Leslie matrix whose $ij$-th entry is $A_{ij}(\mathbf{m}_j, \bar{\mathbf{z}}, \bar{\mathbf{n}}(t))$ that gives either the age-specific survival probability (for $i > 1$) or the age-specific fertility (for $i = 1$) of mutant individuals in the context of the resident. The rare mutant subpopulation thus has population dynamics given by $\mathbf{n}(t+1) = \mathbf{A}(\mathbf{m}, \bar{\mathbf{z}}, \bar{\mathbf{n}}(t))\mathbf{n}(t)$.

The mutant population dynamics around the resident equilibrium $\bar{\mathbf{n}}^*(\bar{\mathbf{m}})$ are to first order of approximation given by

$$\mathbf{n}(t+1) \approx \mathbf{J}\mathbf{n}(t), \qquad \text{(Eq. S2.1.6)}$$

where the local stability matrix of the mutant (Appendix A) is

$$\mathbf{J} = \left. \frac{\partial \mathbf{A}(\mathbf{m}, \bar{\mathbf{z}}, \bar{\mathbf{n}})\mathbf{n}}{\partial \mathbf{n}^\intercal} \right|_{\bar{\mathbf{n}} = \bar{\mathbf{n}}^*} = \left( \frac{\partial}{\partial n_j} \sum_{k=1}^{N_\mathrm{a}} A_{ik}(\mathbf{m}_k, \bar{\mathbf{z}}, \bar{\mathbf{n}}) n_k \bigg|_{\bar{\mathbf{n}} = \bar{\mathbf{n}}^*} \right) = \left( A_{ij}(\mathbf{m}_j, \bar{\mathbf{z}}, \bar{\mathbf{n}}^*(\bar{\mathbf{m}})) \right).$$

Explicitly,

$$\mathbf{J} = \begin{pmatrix} f_1 & f_2 & \cdots & f_{N_\mathrm{a}-1} & f_{N_\mathrm{a}} \\ p_1 & 0 & \cdots & 0 & 0 \\ 0 & p_2 & \cdots & 0 & 0 \\ \vdots & \vdots & \ddots & \vdots & \vdots \\ 0 & 0 & \cdots & p_{N_\mathrm{a}-1} & 0 \end{pmatrix}, \qquad \text{(Eq. S2.1.7)}$$

where we denote the mutant's fertility at age $a$ at the resident population dynamics equilibrium as

$$f_a = f_a(\mathbf{m}_a, \bar{\mathbf{m}}) = A_{1a}(\mathbf{m}_a, \bar{\mathbf{z}}, \bar{\mathbf{n}}^*(\bar{\mathbf{m}})) \qquad \text{(Eq. S2.1.8a)}$$

and the mutant's survival probability from age $a$ to $a+1$ as

$$p_a = p_a(\mathbf{m}_a, \bar{\mathbf{m}}) = A_{a+1,a}(\mathbf{m}_a, \bar{\mathbf{z}}, \bar{\mathbf{n}}^*(\bar{\mathbf{m}})). \qquad \text{(Eq. S2.1.8b)}$$

We denote the fertility of a neutral mutant of age $a$ as $f_a^\circ = f_a(\bar{\mathbf{m}}_a, \bar{\mathbf{m}})$ and the survival probability from age $a$ to $a+1$ of a neutral mutant as $p_a^\circ = p_a(\bar{\mathbf{m}}_a, \bar{\mathbf{m}})$.

#### S2.2 Evolutionary dynamics of the genotype

We now obtain a first-order approximation of invasion fitness and use it to write an equation describing the evolutionary dynamics of the genotype. Invasion fitness is the asymptotic growth rate of the mutant population and it enables the determination of whether the mutant invades the resident population (i.e., whether the mutation increases in frequency) (Otto and Day, 2007). From Eq. S2.1.6, the asymptotic population dynamics of the mutant subpopulation around the resident equilibrium are given to first order of approximation by the eigenvalues and eigenvectors of $\mathbf{J}$. As for residents, we assume that $\mathbf{J}$ is primitive. For instance, $\mathbf{J}$ is primitive if rare mutants can survive to the last age with non-zero probability (i.e., $p_a > 0$ for all $a \in \{1, \ldots, N_\mathrm{a} - 1\}$), rare mutants in the last age reproduce ($f_{N_\mathrm{a}} > 0$), and rare mutants of at least two consecutive age classes have non-zero fertility (i.e., $f_a > 0$ and $f_{a+1} > 0$ for some $a \in \{1, \ldots, N_\mathrm{a} - 1\}$) (Sternberg, 2010, section 9.4.1). Then, from the Perron-Frobenius theorem (Sternberg, 2010, theorem 9.1.1), $\mathbf{J}$ has a real positive eigenvalue $\lambda = \lambda(\mathbf{m}, \bar{\mathbf{m}})$ whose magnitude is strictly larger than that of the other eigenvalues of $\mathbf{J}$. Such leading eigenvalue $\lambda$ is the asymptotic growth rate of the mutant population around the resident equilibrium, and thus gives the mutant's invasion fitness. Since the population dynamics of rare mutants are locally given by Eq. S2.1.6 where $\mathbf{J}$ projects the mutant population to the next ecological time step, the mutant population invades when invasion fitness satisfies $\lambda > 1$.

We consider the evolutionary change in the genotype from the evolutionary time $\tau$, specifically the point at which the socio-devo stable resident is at carrying capacity as marked in Fig. 3, to the evolutionary time $\tau + \Delta\tau$ at which a new socio-devo stable resident is at carrying capacity. The vector $\mathbf{y}$ is a realization of a multivariate random variable $\mathbf{y}$ with probability density $M(\mathbf{y}, \bar{\mathbf{y}})$ called *the mutational distribution* (Dieckmann and Law, 1996), with support in $\mathbb{R}^{N_\mathrm{a} N_\mathrm{g}}$ (abusing notation, we denote a random variable and its realization with the same symbol, as has been common practice—e.g., Lande 1979 and Lynch and Walsh 1998, p. 192). We assume that the mutational distribution is such that (i) the expected mutant genotype is the resident, $\mathrm{E}[\mathbf{y}] = \bar{\mathbf{y}}$; (ii) mutational variance is marginally small (i.e., selection is $\delta$-weak) such that $0 < \mathrm{E}[||\mathbf{y} - \bar{\mathbf{y}}||^2] = \mathrm{tr}(\mathrm{cov}[\mathbf{y}, \mathbf{y}]) = \sum_{i=1}^{N_\mathrm{g}} \sum_{a=1}^{N_\mathrm{a}} \mathrm{E}[(y_{ia} - \bar{y}_{ia})^2] \ll 1$; and (iii) mutation is unbiased, that is, the mutational distribution is even $M(\mathbf{y} - \bar{\mathbf{y}}) = M(\bar{\mathbf{y}} - \mathbf{y})$. Given small mutational variance, Taylor-expanding $\lambda$ with respect to $\mathbf{y}$ around $\bar{\mathbf{y}}$, invasion fitness is to first order of approximation given by

$$\lambda = 1 + (\mathbf{y} - \bar{\mathbf{y}})^\intercal \left. \frac{\mathrm{d}\lambda}{\mathrm{d}\mathbf{y}} \right|_{\mathbf{y} = \bar{\mathbf{y}}} + O(||\mathbf{y} - \bar{\mathbf{y}}||^2), \qquad \text{(Eq. S2.2.1)}$$

where we use the fact that $\lambda|_{\mathbf{y}=\bar{\mathbf{y}}} = 1$ due to density dependence. A given entry of the operator $\mathrm{d}/\mathrm{d}\mathbf{y}|_{\mathbf{y}=\bar{\mathbf{y}}}$, say $\mathrm{d}/\mathrm{d}y_{ia}|_{\mathbf{y}=\bar{\mathbf{y}}}$, takes the total derivative with respect to $y_{ia}$ while keeping all the other genotypic trait values $y_{kj}$ constant. Hence, we refer to $\mathrm{d}\lambda/\mathrm{d}\mathbf{y}|_{\mathbf{y}=\bar{\mathbf{y}}}$ as the *total selection gradient of the genotype* $\mathbf{y}$, which takes the total derivative considering both developmental constraints (Eq. 1) and environmental constraints (Eq. 2). Thus, the total selection gradient of the genotype can be interpreted as measuring *total genotypic selection*. Since the mutant population invades when $\lambda > 1$ and mutational variances are marginally small (i.e., selection is $\delta$-weak), the mutant population invades if and only if

$$(\mathbf{y} - \bar{\mathbf{y}})^\intercal \left. \frac{\mathrm{d}\lambda}{\mathrm{d}\mathbf{y}} \right|_{\mathbf{y} = \bar{\mathbf{y}}} > 0,$$

to first-order of approximation. The left-hand side of this inequality is the dot product of total genotypic selection and the realized mutational effect on the genotype $(\mathbf{y}-\bar{\mathbf{y}})$. The dot product is positive if and only if the absolute value of the smallest angle between two non-zero vectors is smaller than 90 degrees. Hence, to first-order of approximation, the mutant population invades if and only if the mutational effect on the genotype has a vector component in the direction of total genotypic selection.

Using Eq. S2.2.1 and closely following Dieckmann and Law (1996), in section S2.5 we provide a short derivation showing that the evolutionary dynamics of the genotype are given by the canonical equation of adaptive dynamics:

$$\frac{\Delta\bar{\mathbf{y}}}{\Delta\tau} \approx \iota \mathbf{H}_{\mathbf{y}} \left.\frac{\mathrm{d}\lambda}{\mathrm{d}\mathbf{y}}\right|_{\mathbf{y}=\bar{\mathbf{y}}}, \qquad \text{(Eq. S2.2.2a)}$$

where $\iota = \frac{1}{2}\mu(\bar{\mathbf{m}})\bar{n}^*(\bar{\mathbf{m}})$ is a scalar that is proportional to the mutation rate and carrying capacity, whereas

$$\mathbf{H}_{\mathbf{y}} = \mathrm{cov}[\mathbf{y},\mathbf{y}] \qquad \text{(Eq. S2.2.2b)}$$

is equivalently the mutational covariance matrix (of the genotype) and the mechanistic additive genetic covariance matrix of the genotype under our adaptive dynamics assumptions (cf. Eq. 6.1 of Dieckmann and Law 1996 and Eq. 23 of Durinx *et al.* 2008, both of which refer to the evolution of what we call genotypic traits). If the total selection gradient of the genotype is zero, then there is no evolutionary change in the (expected) resident genotype to first order of approximation. However, if the total selection gradient of the genotype is zero, mutants may still invade symmetrically around the resident genotype to second-order of approximation (i.e., $\lambda > 1$ may still hold; Eq. S2.2.1), as in evolutionary branching (Geritz *et al.*, 1998; Leimar, 2009; Débarre *et al.*, 2014). Here we will be concerned with describing the evolutionary dynamics to first-order of approximation, so we will treat the approximation in Eq. S2.2.2a as an equality although we keep the approximation symbol to distinguish what is and what is not an approximation.

Owing to the block structure of $\mathbf{y}$, $\mathbf{H}_{\mathbf{y}}$ is a block matrix whose $aj$-th block entry is the matrix $\mathbf{H}_{\mathbf{y}_a,\mathbf{y}_j} = \mathrm{cov}[\mathbf{y}_a,\mathbf{y}_j]$, which is the mutational or mechanistic additive genetic cross-covariance matrix between the genotypic trait values $\mathbf{y}_a$ at age $a$ and the genotypic trait values $\mathbf{y}_j$ at age $j$. In turn, the $ik$-th entry of $\mathbf{H}_{\mathbf{y}_a,\mathbf{y}_j}$ is $H_{y_{ia},y_{kj}} = \mathrm{cov}[y_{ia},y_{kj}]$ which is the mutational or mechanistic additive genetic covariance between genotypic trait value $y_{ia}$ and genotypic trait value $y_{kj}$. Since $\mathbf{y} \in \mathbb{R}^{N_\mathrm{a}N_\mathrm{g}\times 1}$, then $\mathbf{H}_{\mathbf{y}} \in \mathbb{R}^{N_\mathrm{a}N_\mathrm{g}\times N_\mathrm{a}N_\mathrm{g}}$.

We will define the mechanistic additive genetic covariance matrix $\mathbf{H}_{\boldsymbol{\zeta}}$ of a trait vector $\boldsymbol{\zeta}$ once we define mechanistic breeding value under our adaptive dynamics assumptions. Using a modification of the terminology of Houle (2001) and Klingenberg (2005, 2010), we say that there are no genetic constraints for a vector $\boldsymbol{\zeta}$ if and only if all the eigenvalues of its mechanistic additive genetic covariance matrix $\mathbf{H}_{\boldsymbol{\zeta}}$ are equal and positive; that there are only relative genetic constraints if and only if $\mathbf{H}_{\boldsymbol{\zeta}}$ has different eigenvalues but all are positive; and that there are absolute genetic constraints if and only if $\mathbf{H}_{\boldsymbol{\zeta}}$ has at least one zero eigenvalue (i.e., $\mathbf{H}_{\boldsymbol{\zeta}}$ is singular). If $\boldsymbol{\zeta} = \mathbf{y}$, we speak of mutational rather than genetic constraints. For example, we say there are absolute mutational constraints if and only if $\mathbf{H}_{\mathbf{y}}$ is singular, in which case there is no mutational variation in some directions of genotype space. Hence, from Eq. S2.2.2a, if there are absolute mutational constraints (i.e., $\mathbf{H}_{\mathbf{y}}$ is singular), the evolutionary dynamics of the genotype can stop (i.e., $\Delta\bar{\mathbf{y}}/\Delta\tau \approx \mathbf{0}$) with a non-zero total selection gradient of the genotype (i.e., $\mathrm{d}\lambda/\mathrm{d}\mathbf{y}|_{\mathbf{y}=\bar{\mathbf{y}}} \neq \mathbf{0}$) (because a homogeneous system $\mathbf{Ax} = \mathbf{0}$ has non-zero solutions $\mathbf{x}$ with $\mathbf{A}$ singular if there is any solution to the system). Similarly, we will define the mechanistic additive socio-genetic cross-covariance matrix $\mathbf{L}_{\boldsymbol{\zeta}}$ of a trait vector $\boldsymbol{\zeta}$, and we make the analogous statements as "socio-genetic" constraints of $\boldsymbol{\zeta}$ in terms of $\mathbf{L}_{\boldsymbol{\zeta}}$.

As the resident genotype evolves, the resident phenotype evolves. Specifically, at a given evolutionary time $\tau$, from Eq. (1) the resident phenotype is given by the recurrence equation

$$\bar{\mathbf{x}}_{a+1} = \mathbf{g}_a(\bar{\mathbf{m}}_a,\bar{\mathbf{z}}) \qquad \text{(Eq. S2.2.2c)}$$

for all $a \in \{1,\ldots,N_\mathrm{a}-1\}$ with $\bar{\mathbf{x}}_1$ constant, and where the resident environment is given by

$$\bar{\boldsymbol{\epsilon}}_a = \mathbf{h}_a(\bar{\mathbf{z}}_a,\bar{\mathbf{z}},\tau) \qquad \text{(Eq. S2.2.2d)}$$

for all $a \in \{1,\ldots,N_\mathrm{a}\}$. Intuitively, the evolutionary dynamics of the phenotype thus occur as an outgrowth of the evolutionary dynamics of the genotype and are modulated by the environmental evolutionary dynamics.

Eq. S2.2.2a describes the evolutionary dynamics of the genotype and Eq. S2.2.2c describes the developmental dynamics of the phenotype, so together Eq. S2.2.2 describe the evo-devo dynamics. To characterize the evo-devo process, we obtain general expressions for the total selection gradient of the genotype and for the evolutionary dynamics of the phenotype, environment, geno-phenotype, and geno-envo-phenotype. To do this, we first re-derive the classical form of the selection gradient in age-structured populations, upon which we build our derivations.

#### S2.3 Selection gradient in age-structured populations

To calculate the evo-devo dynamics given by Eq. S2.2.2, we need to calculate the total selection gradient of the genotype $\mathrm{d}\lambda/\mathrm{d}\mathbf{y}|_{\mathbf{y}=\bar{\mathbf{y}}}$. Since the life cycle is age structured (Eq. S2.1.7 and Fig. S1), the total selection gradient of the genotype has the form of the selection gradient in age structured populations, which is well-known but we re-derive it here for ease of reference.

We first use an eigenvalue perturbation theorem to write the selection gradient, which suggests a definition of relative fitness as a quantity that has the same first-order derivatives of invasion fitness. Let $\bar{\zeta}$ and $\zeta$ respectively denote resident and mutant trait values (i.e., $\bar{\zeta}$ is an entry of $\bar{\mathbf{m}}$ and $\zeta$ is an entry of $\mathbf{m}$). From a theorem on eigenvalue perturbation (Eq. 9 of Caswell 1978 or Eq. 9.10 of Caswell 2001), the selection gradient of $\zeta$ is

$$\left.\frac{\partial\lambda}{\partial\zeta}\right|_{\mathbf{y}=\bar{\mathbf{y}}} = \frac{1}{\mathbf{v}^{\circ\intercal}\mathbf{u}^\circ}\mathbf{v}^{\circ\intercal}\left(\left.\frac{\partial\mathbf{J}}{\partial\zeta}\right|_{\mathbf{y}=\bar{\mathbf{y}}}\right)\mathbf{u}^\circ$$

$$= \frac{1}{\mathbf{v}^{\circ\intercal}\mathbf{u}^{\circ}} \sum_{i=1}^{N_a}\sum_{j=1}^{N_a} v_i^{\circ} \left( \frac{\partial J_{ij}}{\partial \zeta}\bigg|_{\mathbf{y}=\bar{\mathbf{y}}} \right) u_j^{\circ}, \quad \text{(Eq. S2.3.1)}$$

where $\mathbf{v}$ and $\mathbf{u}$ are respectively dominant left and right eigenvectors of $\mathbf{J}$ (Eq. S2.1.7). The vector $\mathbf{v}$ lists the mutant reproductive values and the vector $\mathbf{u}$ lists the mutant stable age distribution. In turn, $\mathbf{v}^{\circ} = \mathbf{v}|_{\mathbf{y}=\bar{\mathbf{y}}}$ lists the neutral (mutant) reproductive values and $\mathbf{u}^{\circ} = \mathbf{u}|_{\mathbf{y}=\bar{\mathbf{y}}}$ lists the neutral (mutant) stable age distribution. Substituting $J_{ij}$ for the entries in Eq. S2.1.7 yields

$$\frac{\partial \lambda}{\partial \zeta}\bigg|_{\mathbf{y}=\bar{\mathbf{y}}} = \frac{1}{\mathbf{v}^{\circ\intercal}\mathbf{u}^{\circ}} \sum_{j=1}^{N_a} u_j^{\circ} \left( v_1^{\circ} \frac{\partial f_j}{\partial \zeta}\bigg|_{\mathbf{y}=\bar{\mathbf{y}}} + v_{j+1}^{\circ} \frac{\partial p_j}{\partial \zeta}\bigg|_{\mathbf{y}=\bar{\mathbf{y}}} \right), \quad \text{(Eq. S2.3.2)}$$

where we let $v_{N_a+1} = 0$. Eq. S2.3.1 motivates the definition of the relative fitness of a mutant individual per unit of generation time as

$$w = \frac{1}{\mathbf{v}^{\circ\intercal}\mathbf{u}^{\circ}} \mathbf{v}^{\circ\intercal}\mathbf{J}\mathbf{u}^{\circ} = \frac{1}{\mathbf{v}^{\circ\intercal}\mathbf{u}^{\circ}} \sum_{i=1}^{N_a}\sum_{j=1}^{N_a} v_i^{\circ} J_{ij} u_j^{\circ} \quad \text{(Eq. S2.3.3)}$$

(cf. Lande, 1982, his Eq. 12c) and of the relative fitness of a mutant individual of age $j$ per unit of generation time as

$$w_j = \frac{1}{\mathbf{v}^{\circ\intercal}\mathbf{u}^{\circ}} \sum_{i=1}^{N_a} v_i^{\circ} J_{ij} u_j^{\circ} = \frac{1}{\mathbf{v}^{\circ\intercal}\mathbf{u}^{\circ}} u_j^{\circ} \left( v_1^{\circ} f_j + v_{j+1}^{\circ} p_j \right). \quad \text{(Eq. S2.3.4)}$$

Note, however, that relative fitness $w$ in Eq. S2.3.3 is defined here by its property of having the same first derivative as invasion fitness, so it is a first-order approximation of invasion fitness and is not adequate to assess second-order selection, namely, stabilizing or disruptive selection, in particular, evolutionary branching.

We now obtain that relative fitness depends on the forces of selection, which decrease with age. Age-specific relative fitness (Eq. S2.3.4) depends on the neutral stable age distribution $u_j^{\circ}$ and the neutral reproductive value $v_{j+1}^{\circ}$, which are well-known quantities but we re-derive them in section S2.4.1 for ease of reference. We obtain that the neutral stable age distribution and neutral reproductive value are

$$u_j^{\circ} = \ell_j^{\circ} u_1^{\circ} \quad \text{(Eq. S2.3.5a)}$$

$$v_j^{\circ} = \frac{1}{\ell_j^{\circ}} v_1^{\circ} \sum_{k=j}^{N_a} \ell_k^{\circ} f_k^{\circ}, \quad \text{(Eq. S2.3.5b)}$$

for $j \in \{1, \ldots, N_a\}$ and where $u_1^{\circ}$ and $v_1^{\circ}$ can take any positive value. Hence, the weights on fertility and survival in Eq. S2.3.4 are

$$\frac{u_j^{\circ} v_1^{\circ}}{\mathbf{v}^{\circ\intercal}\mathbf{u}^{\circ}} = \frac{1}{T} \ell_j^{\circ} \quad \text{(Eq. S2.3.6a)}$$

$$\frac{u_j^{\circ} v_{j+1}^{\circ}}{\mathbf{v}^{\circ\intercal}\mathbf{u}^{\circ}} = \frac{1}{T} \frac{1}{p_j^{\circ}} \sum_{k=j+1}^{N_a} \ell_k^{\circ} f_k^{\circ}, \quad \text{(Eq. S2.3.6b)}$$

where generation time is given by Eq. (6). Eq. S2.3.5 and Eq. S2.3.6 recover classic equations (Hamilton 1966 and Caswell 1978, his Eqs. 11 and 12).

We can then obtain a biologically informative expression for the selection gradient in terms relative fitness. Using Eq. S2.3.4, Eq. S2.3.6, and Eq. (7), a mutant's relative fitness at age $j$ is given by Eq. (5b) or with explicit arguments using Eq. S2.1.8,

$$w_j(\mathbf{m}_j, \bar{\mathbf{m}}) = \frac{1}{T(\bar{\mathbf{m}})} \left[ \phi_j(\bar{\mathbf{m}}) f_j(\mathbf{m}_j, \bar{\mathbf{m}}) + \pi_j(\bar{\mathbf{m}}) p_j(\mathbf{m}_j, \bar{\mathbf{m}}) \right]. \quad \text{(Eq. S2.3.7)}$$

Using Eq. S2.3.3, Eq. S2.3.4, and Eq. 5b, a mutant's relative fitness is given by Eq. (5a), or with explicit arguments,

$$w(\mathbf{m}, \bar{\mathbf{m}}) = \sum_{j=1}^{N_a} w_j(\mathbf{m}_j, \bar{\mathbf{m}}). \quad \text{(Eq. S2.3.8)}$$

Hence, from Eq. S2.3.1 and Eq. S2.3.3, the selection gradient entry for trait $\zeta$ is

$$\frac{\partial \lambda}{\partial \zeta}\bigg|_{\mathbf{y}=\bar{\mathbf{y}}} = \frac{\partial w}{\partial \zeta}\bigg|_{\mathbf{y}=\bar{\mathbf{y}}} = \sum_{j=1}^{N_a} \frac{\partial w_j}{\partial \zeta}\bigg|_{\mathbf{y}=\bar{\mathbf{y}}}. \quad \text{(Eq. S2.3.9a)}$$

The same procedure applies for total rather than partial derivatives, so the total selection gradient of $\zeta$ is

$$\frac{\mathrm{d}\lambda}{\mathrm{d}\zeta}\bigg|_{\mathbf{y}=\bar{\mathbf{y}}} = \frac{\mathrm{d}w}{\mathrm{d}\zeta}\bigg|_{\mathbf{y}=\bar{\mathbf{y}}} = \sum_{j=1}^{N_a} \frac{\mathrm{d}w_j}{\mathrm{d}\zeta}\bigg|_{\mathbf{y}=\bar{\mathbf{y}}}. \quad \text{(Eq. S2.3.9b)}$$

It follows that invasion fitness equals a mutant's relative fitness to first-order of approximation around the resident genotype. Indeed, substituting $\zeta$ for $y_{ia}$ for all $i \in \{1, \ldots, N_g\}$ and all $a \in \{1, \ldots, N_a\}$ in Eq. S2.3.9 and using this in Eq. S2.2.1 yields

$$\lambda = 1 + (\mathbf{y} - \bar{\mathbf{y}})^{\intercal} \frac{\mathrm{d}w}{\mathrm{d}\mathbf{y}}\bigg|_{\mathbf{y}=\bar{\mathbf{y}}} + O(||\mathbf{y} - \bar{\mathbf{y}}||^2). \quad \text{(Eq. S2.3.10)}$$

Similarly, Taylor expanding fitness $w$ around the resident genotype with respect to the mutant genotype yields

$$w = 1 + (\mathbf{y} - \bar{\mathbf{y}})^{\intercal} \frac{\mathrm{d}w}{\mathrm{d}\mathbf{y}}\bigg|_{\mathbf{y}=\bar{\mathbf{y}}} + O(||\mathbf{y} - \bar{\mathbf{y}}||^2),$$

so

$$(\mathbf{y} - \bar{\mathbf{y}})^{\intercal} \frac{\mathrm{d}w}{\mathrm{d}\mathbf{y}}\bigg|_{\mathbf{y}=\bar{\mathbf{y}}} = w - 1 + O(||\mathbf{y} - \bar{\mathbf{y}}||^2).$$

Substituting this in Eq. S2.3.10 yields

$$\lambda = w + O(||\mathbf{y} - \bar{\mathbf{y}}||^2). \quad \text{(Eq. S2.3.11)}$$

It is often useful to write selection gradients in terms of lifetime reproductive success if possible. In section S2.4.2, we re-derive that the selection gradients can be expressed in terms of expected lifetime reproductive success, as previously known (Bulmer, 1994; Caswell, 2009), because of our assumption that mutants arise when residents are at carrying capacity (Mylius and Diekmann, 1995). In section S2.4.2, we show that the selection gradient can be written as

$$\frac{\partial \lambda}{\partial \zeta}\bigg|_{\mathbf{y}=\bar{\mathbf{y}}} = \frac{1}{T} \frac{\partial R_0}{\partial \zeta}\bigg|_{\mathbf{y}=\bar{\mathbf{y}}}, \quad \text{(Eq. S2.3.12a)}$$

and that the total selection gradient can be written as

$$\frac{\mathrm{d}\lambda}{\mathrm{d}\zeta}\bigg|_{\mathbf{y}=\bar{\mathbf{y}}} = \frac{1}{T} \frac{\mathrm{d}R_0}{\mathrm{d}\zeta}\bigg|_{\mathbf{y}=\bar{\mathbf{y}}}, \quad \text{(Eq. S2.3.12b)}$$

which recover previous equations (Bulmer 1994, Eq. 25 of Ch. 5; and Caswell 2009, Eqs. 58-61). Proceeding as above, we can obtain a relationship between invasion fitness and lifetime reproductive success. Taylor expanding $R_0$ around the resident genotype with respect to the mutant genotype yields

$$R_0 = 1 + (\mathbf{y} - \bar{\mathbf{y}})^\intercal \left.\frac{\mathrm{d}R_0}{\mathrm{d}\mathbf{y}}\right|_{\mathbf{y}=\bar{\mathbf{y}}} + O(||\mathbf{y} - \bar{\mathbf{y}}||^2),$$

so

$$(\mathbf{y} - \bar{\mathbf{y}})^\intercal \left.\frac{\mathrm{d}R_0}{\mathrm{d}\mathbf{y}}\right|_{\mathbf{y}=\bar{\mathbf{y}}} = R_0 - 1 + O(||\mathbf{y} - \bar{\mathbf{y}}||^2).$$

Using Eq. S2.3.12b, this is

$$(\mathbf{y} - \bar{\mathbf{y}})^\intercal \left.\frac{\mathrm{d}\lambda}{\mathrm{d}\mathbf{y}}\right|_{\mathbf{y}=\bar{\mathbf{y}}} = \frac{1}{T}(R_0 - 1) + O(||\mathbf{y} - \bar{\mathbf{y}}||^2).$$

Substituting this in Eq. S2.2.1 yields

$$\lambda = 1 + \frac{1}{T}(R_0 - 1) + O(||\mathbf{y} - \bar{\mathbf{y}}||^2). \qquad \text{(Eq. S2.3.13)}$$

#### S2.4 Derivation of elements of the selection gradient in age-structured populations

In this section we first derive the stable age distribution, reproductive values, and generation time in age structured populations, which yield Eq. S2.3.5 and Eq. S2.3.6. Then, we derive the selection gradient in age structured populations in terms of lifetime reproductive success. All these quantities are well-known but we re-derive them here for ease of reference.

##### S2.4.1 Stable age distribution, reproductive values, and generation time

The mutant stable age distribution and mutant reproductive value are given by dominant left and right eigenvectors $\mathbf{v}$ and $\mathbf{u}$ of the mutant's local stability matrix $\mathbf{J}$ in Eq. S2.1.7. That is, $\mathbf{v}$ and $\mathbf{u}$ are defined respectively by $\lambda\mathbf{u} = \mathbf{J}\mathbf{u}$ and $\lambda\mathbf{v}^\intercal = \mathbf{v}^\intercal\mathbf{J}$. Expanding these equations yields

$$\lambda u_1 = \sum_{j=1}^{N_\mathrm{a}} f_j u_j \qquad \text{(Eq. S2.4.1a)}$$

$$\lambda u_j = p_{j-1} u_{j-1} \quad \text{for } j \in \{2,\dots,N_\mathrm{a}\} \qquad \text{(Eq. S2.4.1b)}$$

$$\lambda v_j = v_1 f_j + v_{j+1} p_j \quad \text{for } j \in \{1,\dots,N_\mathrm{a}\}, \qquad \text{(Eq. S2.4.1c)}$$

since $v_{N_\mathrm{a}+1} = 0$. Eq. S2.4.1b and Eq. S2.4.1c give the recurrence equations

$$u_j = \lambda^{-1} p_{j-1} u_{j-1}$$

$$v_j = \frac{1}{p_{j-1}}\lambda v_{j-1} - \frac{1}{p_{j-1}} v_1 f_{j-1},$$

for $j \in \{2,\dots,N_\mathrm{a}\}$, which iterating yield

$$u_j = \lambda^{-j+1}\ell_j u_1 \qquad \text{(Eq. S2.4.3a)}$$

$$v_j = \frac{1}{\ell_j}\lambda^{j-1} v_1 - v_1 \sum_{k=1}^{j-1} \frac{\lambda^{j-1-k}}{\ell_j/\ell_k} f_k$$

$$= \frac{1}{\ell_j}\lambda^{j-1} v_1 \left(1 - \sum_{k=1}^{j-1} \lambda^{-k}\ell_k f_k\right), \qquad \text{(Eq. S2.4.3b)}$$

where $\ell_j = \prod_{k=1}^{j-1} p_k$ is mutant survivorship from age 1 to age $j \in \{2,\dots,N_\mathrm{a}\}$. Eq. S2.4.3b can be rewritten in the standard form of Fisher's (1927) reproductive value in discrete time using the Euler-Lotka equation as follows. Defining $\ell_1 = 1$ and since $\lambda^0 = 1$, substituting Eq. S2.4.3a in Eq. S2.4.1a and dividing both sides of the equation by $\lambda u_1$ yields

$$1 = \sum_{j=1}^{N_\mathrm{a}} \lambda^{-j}\ell_j f_j, \qquad \text{(Eq. S2.4.4)}$$

which is the Euler-Lotka equation in discrete time (Charlesworth 1994, Eq. 1.42 and Caswell 2001, Eq. 4.42). Partitioning the sum in Eq. S2.4.4 yields

$$1 - \sum_{j=1}^{m-1} \lambda^{-j}\ell_j f_j = \sum_{j=m}^{N_\mathrm{a}} \lambda^{-j}\ell_j f_j, \qquad \text{(Eq. S2.4.5)}$$

which substituted in Eq. S2.4.3b yields

$$v_j = \frac{1}{\ell_j}\lambda^{j-1} v_1 \sum_{k=j}^{N_\mathrm{a}} \lambda^{-k}\ell_k f_k. \qquad \text{(Eq. S2.4.6)}$$

This equation is the standard form of Fisher's (1927) reproductive value in discrete time (Eq. 4.89 of Caswell 2001). Hence, from Eq. S2.4.3a and Eq. S2.4.6, we obtain the mutant stable age distribution and mutant reproductive value:

$$u_j = \lambda^{-j+1}\ell_j u_1$$

$$v_j = \frac{1}{\ell_j}\lambda^{j-1} v_1 \sum_{k=j}^{N_\mathrm{a}} \lambda^{-k}\ell_k f_k,$$

for $j \in \{2,\dots,N_\mathrm{a}\}$, where $u_1$ and $v_1$ can take any positive value. Evaluating at neutrality ($\mathbf{y} = \bar{\mathbf{y}}$), we have that $\lambda^\circ = \lambda|_{\mathbf{y}=\bar{\mathbf{y}}} = 1$, which yields Eq. S2.3.5.

We now obtain a well-known expression of generation time to be used in our expressions for the forces of selection. Bienvenu and Legendre (2015) find that generation time can be measured by

$$T = \frac{\mathbf{v}^{\circ\intercal}\mathbf{u}^\circ}{\mathbf{v}^{\circ\intercal}\mathbf{F}^\circ\mathbf{u}^\circ},$$

where we evaluate at resident trait values given our adaptive dynamics assumptions, and where $\mathbf{F}$ is given by Eq. S2.1.7 setting all $p_j$ to zero. Using Eq. S2.4.1a, it is easily checked that $\mathbf{v}^{\circ\intercal}\mathbf{F}^\circ\mathbf{u}^\circ = v_1^\circ u_1^\circ$. In turn, we have that the numerator is

$$\mathbf{v}^{\circ\intercal}\mathbf{u}^\circ = \sum_{j=1}^{N_\mathrm{a}} v_j^\circ u_j^\circ.$$

Thus, using Eq. S2.3.5 yields

$$T = \frac{\mathbf{v}^{\circ\intercal}\mathbf{u}^\circ}{v_1^\circ u_1^\circ} = \frac{v_1^\circ u_1^\circ + \sum_{j=2}^{N_\mathrm{a}} v_j^\circ u_j^\circ}{v_1^\circ u_1^\circ}$$

$$= \frac{v_1^\circ u_1^\circ + v_1^\circ u_1^\circ \sum_{j=2}^{N_\mathrm{a}}\sum_{k=j}^{N_\mathrm{a}} \ell_k^\circ f_k^\circ}{v_1^\circ u_1^\circ}$$

$$= 1 + \sum_{j=2}^{N_a} \sum_{k=j}^{N_a} \ell_k^\circ f_k^\circ. \qquad \text{(Eq. S2.4.8)}$$

We further manipulate this expression to recover a standard expression of generation time (Charlesworth 1994, Eq. 1.47c; Bulmer 1994, Eq. 25, Ch. 25; Bienvenu and Legendre 2015, Eq. 5). Evaluating the Euler-Lotka equation (Eq. S2.4.4) at the resident genotype (so $\lambda|_{\mathbf{y}=\bar{\mathbf{y}}} = 1$), we obtain that a neutral mutant's expected lifetime reproductive success is

$$R_0^\circ = \sum_{j=1}^{N_a} \ell_j^\circ f_j^\circ = 1. \qquad \text{(Eq. S2.4.9)}$$

Therefore, Eq. S2.4.8 is

$$\begin{aligned}
T &= 1 + \sum_{j=2}^{N_a} \sum_{k=j}^{N_a} \ell_k^\circ f_k^\circ = R_0^\circ + \sum_{j=2}^{N_a} \sum_{k=j}^{N_a} \ell_k^\circ f_k^\circ \\
&= \sum_{j=1}^{N_a} \ell_j^\circ f_j^\circ + \sum_{j=2}^{N_a} \sum_{k=j}^{N_a} \ell_k^\circ f_k^\circ \\
&= \ell_1^\circ f_1^\circ + \sum_{j=2}^{N_a} \ell_j^\circ f_j^\circ + \sum_{j=2}^{N_a} \sum_{k=j}^{N_a} \ell_k^\circ f_k^\circ \\
&= \ell_1^\circ f_1^\circ + \sum_{j=2}^{N_a} \left( \ell_j^\circ f_j^\circ + \sum_{k=j}^{N_a} \ell_k^\circ f_k^\circ \right).
\end{aligned}$$

Expanding the rightmost sum yields

$$T = \ell_1^\circ f_1^\circ + \sum_{j=2}^{N_a} \left( \ell_j^\circ f_j^\circ + \ell_j^\circ f_j^\circ + \ell_{j+1}^\circ f_{j+1}^\circ + \cdots + \ell_{N_a}^\circ f_{N_a}^\circ \right)$$

Expanding the remaining sum yields

$$\begin{aligned}
T = \ell_1^\circ f_1^\circ &+ \left( \ell_2^\circ f_2^\circ + \ell_2^\circ f_2^\circ + \ell_3^\circ f_3^\circ + \cdots + \ell_{N_a}^\circ f_{N_a}^\circ \right) \\
&+ \left( \ell_3^\circ f_3^\circ + \ell_3^\circ f_3^\circ + \ell_4^\circ f_4^\circ + \cdots + \ell_{N_a}^\circ f_{N_a}^\circ \right) \\
&+ \cdots \\
&+ \left( \ell_{N_a-1}^\circ f_{N_a-1}^\circ + \ell_{N_a-1}^\circ f_{N_a-1}^\circ + \ell_{N_a}^\circ f_{N_a}^\circ \right) \\
&+ \left( \ell_{N_a}^\circ f_{N_a}^\circ + \ell_{N_a}^\circ f_{N_a}^\circ \right).
\end{aligned}$$

Collecting common terms yields

$$\begin{aligned}
T &= \ell_1^\circ f_1^\circ + 2\ell_2^\circ f_2^\circ + 3\ell_3^\circ f_3^\circ + 4\ell_4^\circ f_4^\circ \\
&\quad + \cdots + N_a \ell_{N_a}^\circ f_{N_a}^\circ \\
&= \sum_{j=1}^{N_a} j \ell_j^\circ f_j^\circ, \qquad \text{(Eq. S2.4.10)}
\end{aligned}$$

which is Eq. (6). This expression recovers a standard measure of generation time (Charlesworth 1994, Eq. 1.47c; Bulmer 1994, Eq. 25, Ch. 25; Bienvenu and Legendre 2015, Eq. 5).

##### S2.4.2 Selection gradient in terms of $R_0$

Following Hamilton (1966) (see also Eqs. 58-61 in Caswell 2009), we differentiate the Euler-Lotka equation (Eq. S2.4.4) implicitly with respect to a mutant trait value $\zeta$, which yields

$$0 = \sum_{j=1}^{N_a} \left( \lambda^{-j} \frac{\partial \ell_j f_j}{\partial \zeta} - j \ell_j f_j \lambda^{-j-1} \frac{\partial \lambda}{\partial \zeta} \right) \Bigg|_{\mathbf{y}=\bar{\mathbf{y}}}.$$

Noting that $\lambda|_{\mathbf{y}=\bar{\mathbf{y}}} = 1$ and solving for the selection gradient, we obtain

$$\begin{aligned}
\frac{\partial \lambda}{\partial \zeta}\bigg|_{\mathbf{y}=\bar{\mathbf{y}}} &= \frac{1}{\sum_{j=1}^{N_a} j \ell_j^\circ f_j^\circ} \sum_{j=1}^{N_a} \frac{\partial \ell_j f_j}{\partial \zeta}\bigg|_{\mathbf{y}=\bar{\mathbf{y}}} \\
&= \frac{1}{T} \frac{\partial R_0}{\partial \zeta}\bigg|_{\mathbf{y}=\bar{\mathbf{y}}}, \qquad \text{(Eq. S2.4.11)}
\end{aligned}$$

where we use Eq. (8) and Eq. S2.4.10. This is Eq. S2.3.12a. The same procedure using total derivatives yields Eq. S2.3.12b.

#### S2.5 Derivation of the canonical equation under our assumptions

In this section we derive the equation describing the evolutionary dynamics of the genotype. This derivation closely follows that of Dieckmann and Law (1996) except in a few places, particularly in that we consider deterministic population dynamics so the only source of stochasticity in our framework is due to mutation. Denote by $\bar{\mathbf{y}}'(\tau + \Delta\tau)$ a multivariate random variable describing the possible residents at time $\tau + \Delta\tau$ following fixation of mutants arising at time $\tau$. Let this random variable have probability density function $P(\bar{\mathbf{y}}', \tau + \Delta\tau)$ at time $\tau + \Delta\tau$, with support in $\mathbb{R}^{N_a N_g}$. Hence, the expected resident genotype at time $\tau + \Delta\tau$ is

$$\bar{\mathbf{y}}(\tau + \Delta\tau) \equiv \mathrm{E}[\bar{\mathbf{y}}'(\tau + \Delta\tau)] = \int_{\mathbb{R}^{N_a N_g}} \bar{\mathbf{y}}' P(\bar{\mathbf{y}}', \tau + \Delta\tau) \mathrm{d}\bar{\mathbf{y}}'.$$

The evolutionary change in the resident genotype thus satisfies

$$\begin{aligned}
\frac{\Delta \bar{\mathbf{y}}}{\Delta \tau} &= \frac{\mathrm{E}[\bar{\mathbf{y}}'(\tau + \Delta\tau)] - \mathrm{E}[\bar{\mathbf{y}}'(\tau)]}{\Delta\tau} \\
&= \frac{1}{\Delta\tau} \left( \int_{\mathbb{R}^{N_a N_g}} \bar{\mathbf{y}}' P(\bar{\mathbf{y}}', \tau + \Delta\tau) \mathrm{d}\bar{\mathbf{y}}' - \int_{\mathbb{R}^{N_a N_g}} \bar{\mathbf{y}}' P(\bar{\mathbf{y}}', \tau) \mathrm{d}\bar{\mathbf{y}}' \right).
\end{aligned}$$

Factorizing yields

$$\begin{aligned}
\frac{\Delta \bar{\mathbf{y}}}{\Delta \tau} &= \int_{\mathbb{R}^{N_a N_g}} \bar{\mathbf{y}}' \frac{P(\bar{\mathbf{y}}', \tau + \Delta\tau) - P(\bar{\mathbf{y}}', \tau)}{\Delta\tau} \mathrm{d}\bar{\mathbf{y}}' \\
&= \int_{\mathbb{R}^{N_a N_g}} \bar{\mathbf{y}}' \frac{\Delta P(\bar{\mathbf{y}}', \tau)}{\Delta\tau} \mathrm{d}\bar{\mathbf{y}}'.
\end{aligned}$$

Now, the evolutionary change in the distribution of the resident genotype satisfies the master equation

$$\frac{\Delta P(\bar{\mathbf{y}}', \tau)}{\Delta\tau} = \int_{\mathbb{R}^{N_a N_g}} \left[ \omega(\mathbf{y}, \bar{\mathbf{y}}') P(\mathbf{y}, \tau) - \omega(\bar{\mathbf{y}}', \mathbf{y}) P(\bar{\mathbf{y}}', \tau) \right] \mathrm{d}\mathbf{y},$$

where $\omega(\mathbf{y}, \bar{\mathbf{y}}')$ is the rate at which a resident $\mathbf{y}$ is replaced by $\bar{\mathbf{y}}'$. Then, the evolutionary change in the genotype is

$$\frac{\Delta \bar{\mathbf{y}}}{\Delta \tau} = \int_{\mathbb{R}^{N_a N_g}} \bar{\mathbf{y}}' \Big( \int_{\mathbb{R}^{N_a N_g}} [\omega(\mathbf{y}, \bar{\mathbf{y}}') P(\mathbf{y}, \tau) - \omega(\bar{\mathbf{y}}', \mathbf{y}) P(\bar{\mathbf{y}}', \tau)] \mathrm{d}\mathbf{y} \Big) \mathrm{d}\bar{\mathbf{y}}'.$$

Since the integral is a linear operator, we have

$$\frac{\Delta\bar{\mathbf{y}}}{\Delta\tau} = \int\limits_{\mathbb{R}^{N_a N_g}} \int\limits_{\mathbb{R}^{N_a N_g}} \bar{\mathbf{y}}' \omega(\mathbf{y},\bar{\mathbf{y}}') P(\mathbf{y},\tau) \mathrm{d}\mathbf{y}\mathrm{d}\bar{\mathbf{y}}' - \int\limits_{\mathbb{R}^{N_a N_g}} \int\limits_{\mathbb{R}^{N_a N_g}} \bar{\mathbf{y}}' \omega(\bar{\mathbf{y}}',\mathbf{y}) P(\bar{\mathbf{y}}',\tau) \mathrm{d}\mathbf{y}\mathrm{d}\bar{\mathbf{y}}'.$$

Exchanging $\mathbf{y}$ for $\bar{\mathbf{y}}'$ in the first term since they are dummy variables yields

$$\frac{\Delta\bar{\mathbf{y}}}{\Delta\tau} = \int\limits_{\mathbb{R}^{N_a N_g}} \int\limits_{\mathbb{R}^{N_a N_g}} \mathbf{y} \omega(\bar{\mathbf{y}}',\mathbf{y}) P(\bar{\mathbf{y}}',\tau) \mathrm{d}\mathbf{y}\mathrm{d}\bar{\mathbf{y}}' - \int\limits_{\mathbb{R}^{N_a N_g}} \int\limits_{\mathbb{R}^{N_a N_g}} \bar{\mathbf{y}}' \omega(\bar{\mathbf{y}}',\mathbf{y}) P(\bar{\mathbf{y}}',\tau) \mathrm{d}\mathbf{y}\mathrm{d}\bar{\mathbf{y}}'.$$

Factorizing yields

$$\frac{\Delta\bar{\mathbf{y}}}{\Delta\tau} = \int\limits_{\mathbb{R}^{N_a N_g}} \int\limits_{\mathbb{R}^{N_a N_g}} (\mathbf{y}-\bar{\mathbf{y}}') \omega(\bar{\mathbf{y}}',\mathbf{y}) P(\bar{\mathbf{y}}',\tau) \mathrm{d}\mathbf{y}\mathrm{d}\bar{\mathbf{y}}'. \quad \text{(Eq. S2.5.1)}$$

Assuming that invasion implies fixation, we let the rate at which resident $\bar{\mathbf{y}}'$ is replaced by $\mathbf{y}$ be

$$\omega(\bar{\mathbf{y}}',\mathbf{y}) = \delta(\bar{\mathbf{y}}'-\bar{\mathbf{y}})\mu(\bar{\mathbf{m}}')\bar{n}^*(\bar{\mathbf{m}}')\frac{M(\mathbf{y}-\bar{\mathbf{y}}')}{P(\bar{\mathbf{y}}',\tau)}[\lambda(\mathbf{m},\bar{\mathbf{m}}')-1] \quad \text{(Eq. S2.5.2)}$$

if $\lambda(\mathbf{m},\bar{\mathbf{m}}') > 1$ or $\omega(\bar{\mathbf{y}}',\mathbf{y}) = 0$ otherwise. Here $\delta(\cdot)$ is the Dirac delta function, $\mathbf{m}$ is the mutant geno-envo-phenotype arising from $\mathbf{y}$, and $\bar{\mathbf{m}}'$ is the SDS resident geno-envo-phenotype arising from $\bar{\mathbf{y}}'$. This expression for $\omega(\bar{\mathbf{y}}',\mathbf{y})$ can be understood as comprising the probability density $\delta(\bar{\mathbf{y}}'-\bar{\mathbf{y}})$ that the resident $\bar{\mathbf{y}}'$ is $\bar{\mathbf{y}}$, times the rate of appearance of new mutants given by the carrying capacity $\bar{n}^*(\bar{\mathbf{m}}')$ and the mutation rate $\mu(\bar{\mathbf{m}}') \in [0,1]$, times the conditional probability density $M(\mathbf{y}-\bar{\mathbf{y}}')/P(\bar{\mathbf{y}}',\tau)$ that a mutant is $\mathbf{y}$ given that the resident is $\bar{\mathbf{y}}'$ at time $\tau$, times the rate of substitution $\lambda(\mathbf{m},\bar{\mathbf{m}}')-1$ for a mutant $\mathbf{y}$ in the context of resident $\bar{\mathbf{y}}'$. Substituting Eq. S2.5.2 into Eq. S2.5.1 using Eq. S2.2.1 yields

$$\frac{\Delta\bar{\mathbf{y}}}{\Delta\tau} \approx \int\limits_{Q} \left\{ (\mathbf{y}-\bar{\mathbf{y}}')\delta(\bar{\mathbf{y}}'-\bar{\mathbf{y}})\mu(\bar{\mathbf{m}}')n^*(\bar{\mathbf{m}}')\frac{M(\mathbf{y}-\bar{\mathbf{y}}')}{P(\bar{\mathbf{y}}',\tau)} (\mathbf{y}-\bar{\mathbf{y}}')^\intercal \left.\frac{\mathrm{d}\lambda}{\mathrm{d}\mathbf{y}}\right|_{\mathbf{y}=\bar{\mathbf{y}}'} P(\bar{\mathbf{y}}',\tau)\right\} \mathrm{d}\mathbf{y}\mathrm{d}\bar{\mathbf{y}}',$$

where the integration ranges over the mutant and resident genotypes that render invasion fitness greater than one, that is,

$$Q = \left\{(\mathbf{y},\bar{\mathbf{y}}') | \lambda(\mathbf{m},\bar{\mathbf{m}}') > 1\right\}.$$

Cancelling $P(\bar{\mathbf{y}}',\tau)$ produces

$$\frac{\Delta\bar{\mathbf{y}}}{\Delta\tau} \approx \int\limits_{Q} (\mathbf{y}-\bar{\mathbf{y}}')\delta(\bar{\mathbf{y}}'-\bar{\mathbf{y}})\mu(\bar{\mathbf{m}}')n^*(\bar{\mathbf{m}}')M(\mathbf{y}-\bar{\mathbf{y}}') (\mathbf{y}-\bar{\mathbf{y}}')^\intercal \left.\frac{\mathrm{d}\lambda}{\mathrm{d}\mathbf{y}}\right|_{\mathbf{y}=\bar{\mathbf{y}}'} \mathrm{d}\mathbf{y}\mathrm{d}\bar{\mathbf{y}}'.$$

Using the sifting property of the Dirac delta function [i.e., $\int_{\mathbb{R}^n} F(\mathbf{y})\delta(\mathbf{y}-\bar{\mathbf{y}})\mathrm{d}\mathbf{y} = F(\bar{\mathbf{y}})$ for any function $F(\mathbf{y})$ with $\mathbf{y} \in \mathbb{R}^n$] yields

$$\frac{\Delta\bar{\mathbf{y}}}{\Delta\tau} \approx \int\limits_{R} (\mathbf{y}-\bar{\mathbf{y}})\mu(\bar{\mathbf{m}})n^*(\bar{\mathbf{m}})[M(\mathbf{y}-\bar{\mathbf{y}})](\mathbf{y}-\bar{\mathbf{y}})^\intercal \left.\frac{\mathrm{d}\lambda}{\mathrm{d}\mathbf{y}}\right|_{\mathbf{y}=\bar{\mathbf{y}}} \mathrm{d}\mathbf{y},$$

where the integration ranges over the mutant genotypes that render invasion fitness greater than one, that is,

$$R = \left\{\mathbf{y} | \lambda(\mathbf{m},\bar{\mathbf{m}}) > 1\right\}.$$

Since the integral is a linear operator and because the evaluation at $\mathbf{y} = \bar{\mathbf{y}}$ makes the gradient constant with respect to $\mathbf{y}$, then

$$\frac{\Delta\bar{\mathbf{y}}}{\Delta\tau} \approx \mu(\bar{\mathbf{m}})n^*(\bar{\mathbf{m}}) \int\limits_{R} (\mathbf{y}-\bar{\mathbf{y}})(\mathbf{y}-\bar{\mathbf{y}})^\intercal M(\mathbf{y}-\bar{\mathbf{y}})\mathrm{d}\mathbf{y} \left.\frac{\mathrm{d}\lambda}{\mathrm{d}\mathbf{y}}\right|_{\mathbf{y}=\bar{\mathbf{y}}}.$$

From the assumption of unbiased mutation (i.e., the function $M(\mathbf{y}-\bar{\mathbf{y}})$ is even, so $M(\mathbf{y}-\bar{\mathbf{y}}) = M(\bar{\mathbf{y}}-\mathbf{y})$), the above integral is half the value of the corresponding integral over the genotype space, so

$$\frac{\Delta\bar{\mathbf{y}}}{\Delta\tau} \approx \frac{1}{2}\mu(\bar{\mathbf{m}})n^*(\bar{\mathbf{m}}) \int\limits_{\mathbb{R}^{N_a N_g}} (\mathbf{y}-\bar{\mathbf{y}})(\mathbf{y}-\bar{\mathbf{y}})^\intercal M(\mathbf{y}-\bar{\mathbf{y}})\mathrm{d}\mathbf{y} \left.\frac{\mathrm{d}\lambda}{\mathrm{d}\mathbf{y}}\right|_{\mathbf{y}=\bar{\mathbf{y}}}.$$

Hence, by definition of covariance matrix, we have

$$\frac{\Delta\bar{\mathbf{y}}}{\Delta\tau} \approx \iota(\bar{\mathbf{m}})\mathrm{cov}[\mathbf{y},\mathbf{y}] \left.\frac{\mathrm{d}\lambda}{\mathrm{d}\mathbf{y}}\right|_{\mathbf{y}=\bar{\mathbf{y}}}, \quad \text{(Eq. S2.5.3)}$$

where

$$\iota(\bar{\mathbf{m}}) = \frac{1}{2}\mu(\bar{\mathbf{m}})\bar{n}^*(\bar{\mathbf{m}}).$$

The matrix $\mathrm{cov}[\mathbf{y},\mathbf{y}]$ is *the mutational covariance matrix* (of the genotype). Eq. S2.5.3 recovers the canonical equation of adaptive dynamics (cf. Eq. 6.1 of Dieckmann and Law 1996 and Eq. 23 of Durinx *et al.* 2008).

To see that $\mathrm{cov}[\mathbf{y},\mathbf{y}]$ is also the mechanistic additive genetic covariance of the genotype, denoted by $\mathbf{H}_\mathbf{y}$, note the following. In Layer 6, Eq. 2, we define the *mechanistic additive genetic covariance matrix* $\mathbf{H}_{\boldsymbol{\zeta}}$ of a vector $\boldsymbol{\zeta} \in \mathbb{R}^{m\times 1}$ under our adaptive dynamics assumptions, and show that

$$\mathbf{H}_{\boldsymbol{\zeta}} = \left.\left(\frac{\mathrm{d}\boldsymbol{\zeta}}{\mathrm{d}\mathbf{y}^\intercal}\mathrm{cov}[\mathbf{y},\mathbf{y}]\frac{\mathrm{d}\boldsymbol{\zeta}^\intercal}{\mathrm{d}\mathbf{y}}\right)\right|_{\mathbf{y}=\bar{\mathbf{y}}}.$$

As we will later show, since the genotype is developmentally independent (i.e., it does not have developmental constraints, or is open-loop) so $\mathrm{d}\mathbf{y}^\intercal/\mathrm{d}\mathbf{y}|_{\mathbf{y}=\bar{\mathbf{y}}} = \mathbf{I}$ (Eq. S5.2.13). Hence, the mechanistic additive genetic covariance matrix of the genotype $\mathbf{H}_\mathbf{y}$ equals the mutational covariance matrix $\mathrm{cov}[\mathbf{y},\mathbf{y}]$. This yields Eq. (3) and Eq. S2.2.2a.

### S3 Further results: bottom layers of the evo-devo process

We use the above model to obtain our results. The main results are summarized in the main text section 4. We arrange a comprehensive presentation of the results in a layered structure that we call the evo-devo process (outlined in Fig. 4). In the main text section 5 we present the top layers 6 and 7, corresponding to genetic covariation and evolutionary dynamics. In this section we present the results corresponding to the bottom layers 1-5, corresponding to the elementary components of the evo-devo process, matrices of direct effects, matrices of total immediate effects, matrices of total effects, and matrices of stabilized effects.

#### S3.1 Layer 1: elementary components

All the equations of the evo-devo process can be calculated from ten elementary components. These include five "core" elementary components: the fertility $f_a(\mathbf{m}_a, \bar{\mathbf{m}})$, survival probability $p_a(\mathbf{m}_a, \bar{\mathbf{m}})$, developmental map $\mathbf{g}_a(\mathbf{m}_a, \bar{\mathbf{z}})$, and environmental map $\mathbf{h}_a(\mathbf{z}_a, \bar{\mathbf{z}}, \tau)$ for all ages $a$, as well as the mutational covariance matrix $\mathbf{H}_\mathbf{y}$ (Fig. 4, Layer 1). The remaining five elementary components of the evo-devo process are the mutation rate $\mu$ and the initial conditions for the various dynamical processes, namely, the evolutionarily initial resident genotype $\bar{\mathbf{y}}(\tau = 1)$, the developmentally initial resident phenotype $\bar{\mathbf{x}}_1$, the population density $\bar{n}_1^*$ at carrying capacity of initial-age residents, and the socio-devo initial resident phenotype $\bar{\mathbf{x}}(\theta = 1)$. Once the five core elementary components are available, either from purely theoretical models or using empirical data, the equations of all the remaining layers of the evo-devo process can be derived. The remaining elementary components are then needed to compute the solution of the evo-devo dynamics. The five core elementary components except for $\mathbf{H}_\mathbf{y}$ correspond to the elementary components of physiologically structured models of population dynamics (de Roos, 1997).

#### S3.2 Layer 2: direct effects

In this section, we write the equations for layer 2, that of the direct-effect matrices which constitute nearly elementary components of the evo-devo process. Direct-effect matrices measure the direct effect that a variable has on another variable. Direct-effect matrices capture various effects of age structure, including the declining forces of selection as age advances.

Direct-effect matrices include direct selection gradients, which have the following structure due to age-structure. The *direct selection gradient of the phenotype, genotype, or environment* is

$$\left.\frac{\partial w}{\partial \boldsymbol{\zeta}}\right|_{\mathbf{y}=\bar{\mathbf{y}}} = \left.\left(\frac{\partial w_1}{\partial \boldsymbol{\zeta}_1}; \cdots; \frac{\partial w_{N_\mathrm{a}}}{\partial \boldsymbol{\zeta}_{N_\mathrm{a}}}\right)\right|_{\mathbf{y}=\bar{\mathbf{y}}}, \qquad \text{(Layer 2, Eq. S1)}$$

for $\boldsymbol{\zeta} \in \{\mathbf{x}, \mathbf{y}, \boldsymbol{\epsilon}\}$, with dimensions for $\partial w/\partial \mathbf{x}|_{\mathbf{y}=\bar{\mathbf{y}}} \in \mathbb{R}^{N_\mathrm{a} N_\mathrm{p} \times 1}$, $\partial w/\partial \mathbf{y}|_{\mathbf{y}=\bar{\mathbf{y}}} \in \mathbb{R}^{N_\mathrm{a} N_\mathrm{g} \times 1}$, and $\partial w/\partial \boldsymbol{\epsilon}|_{\mathbf{y}=\bar{\mathbf{y}}} \in \mathbb{R}^{N_\mathrm{a} N_\mathrm{e} \times 1}$. These gradients measure direct directional selection on the phenotype, genotype, or environment, respectively. Analogously, Lande's (1979) selection gradient measures direct directional selection under quantitative genetics assumptions. Also, the direct selection gradient of the environment measures the environmental sensitivity of selection (Chevin *et al.*, 2010). The block entries of Layer 2, Eq. S1 can be computed by differentiating Eq. (5b). Note that Layer 2, Eq. S1 takes the derivative of fitness at each age, so from Eq. (5b) each block entry in Layer 2, Eq. S1 is weighted by the forces of selection at each age. Thus, the selection gradients in Layer 2, Eq. S1 capture the declining forces of selection in that increasingly rightward block entries have smaller magnitude if survival and fertility effects are of the same magnitude as age increases.

We use the above definitions to form the following aggregate direct selection gradients. The *direct selection gradient of the geno-phenotype* is

$$\left.\frac{\partial w}{\partial \mathbf{z}}\right|_{\mathbf{y}=\bar{\mathbf{y}}} \equiv \left.\left(\frac{\partial w}{\partial \mathbf{x}}; \frac{\partial w}{\partial \mathbf{y}}\right)\right|_{\mathbf{y}=\bar{\mathbf{y}}} \in \mathbb{R}^{N_\mathrm{a}(N_\mathrm{p}+N_\mathrm{g}) \times 1},$$

and the *direct selection gradient of the geno-envo-phenotype* is

$$\left.\frac{\partial w}{\partial \mathbf{m}}\right|_{\mathbf{y}=\bar{\mathbf{y}}} \equiv \left.\left(\frac{\partial w}{\partial \mathbf{x}}; \frac{\partial w}{\partial \mathbf{y}}; \frac{\partial w}{\partial \boldsymbol{\epsilon}}\right)\right|_{\mathbf{y}=\bar{\mathbf{y}}} \in \mathbb{R}^{N_\mathrm{a}(N_\mathrm{p}+N_\mathrm{g}+N_\mathrm{e}) \times 1}.$$

Direct-effect matrices also include matrices that measure direct developmental bias. These matrices have specific, sparse structure due to *the arrow of developmental time*: changing a trait at a given age cannot have effects on the developmental past of the individual and only directly affects the developmental present or immediate future. Using matrix calculus notation (Appendix A), the block matrix of *direct effects of a mutant's phenotype on her phenotype* is

$$\left.\frac{\partial \mathbf{x}^\intercal}{\partial \mathbf{x}}\right|_{\mathbf{y}=\bar{\mathbf{y}}} \equiv \left.\begin{pmatrix} \frac{\partial \mathbf{x}_1^\intercal}{\partial \mathbf{x}_1} & \cdots & \frac{\partial \mathbf{x}_{N_\mathrm{a}}^\intercal}{\partial \mathbf{x}_1} \\ \vdots & \ddots & \vdots \\ \frac{\partial \mathbf{x}_1^\intercal}{\partial \mathbf{x}_{N_\mathrm{a}}} & \cdots & \frac{\partial \mathbf{x}_{N_\mathrm{a}}^\intercal}{\partial \mathbf{x}_{N_\mathrm{a}}} \end{pmatrix}\right|_{\mathbf{y}=\bar{\mathbf{y}}}$$

$$= \left.\begin{pmatrix} \mathbf{I} & \frac{\partial \mathbf{x}_2^\intercal}{\partial \mathbf{x}_1} & \cdots & \mathbf{0} & \mathbf{0} \\ \mathbf{0} & \mathbf{I} & \cdots & \mathbf{0} & \mathbf{0} \\ \vdots & \vdots & \ddots & \vdots & \vdots \\ \mathbf{0} & \mathbf{0} & \cdots & \mathbf{I} & \frac{\partial \mathbf{x}_{N_\mathrm{a}}^\intercal}{\partial \mathbf{x}_{N_\mathrm{a}-1}} \\ \mathbf{0} & \mathbf{0} & \cdots & \mathbf{0} & \mathbf{I} \end{pmatrix}\right|_{\mathbf{y}=\bar{\mathbf{y}}} \in \mathbb{R}^{N_\mathrm{a} N_\mathrm{p} \times N_\mathrm{a} N_\mathrm{p}}, \qquad \text{(Layer 2, Eq. S2a)}$$

which can be understood as measuring direct developmental bias from the phenotype. The equality (Layer 2, Eq. S2a) follows because the direct effects of a mutant's phenotype on her phenotype are only non-zero at the next age (from the developmental constraint in Eq. 1) or when the phenotypes are differentiated with respect to themselves. The block entries of Layer 2, Eq. S2a can be computed by differentiating the developmental constraint (Eq. 1). Analogously, the block matrix of *direct* *effects of a mutant's genotype on her phenotype* is

$$\left.\frac{\partial \mathbf{x}^{\intercal}}{\partial \mathbf{y}}\right|_{\mathbf{y}=\bar{\mathbf{y}}} = \left.\begin{pmatrix} \mathbf{0} & \frac{\partial \mathbf{x}_2^{\intercal}}{\partial \mathbf{y}_1} & \cdots & \mathbf{0} & \mathbf{0} \\ \mathbf{0} & \mathbf{0} & \cdots & \mathbf{0} & \mathbf{0} \\ \vdots & \vdots & \ddots & \vdots & \vdots \\ \mathbf{0} & \mathbf{0} & \cdots & \mathbf{0} & \frac{\partial \mathbf{x}_{N_\mathrm{a}}^{\intercal}}{\partial \mathbf{y}_{N_\mathrm{a}-1}} \\ \mathbf{0} & \mathbf{0} & \cdots & \mathbf{0} & \mathbf{0} \end{pmatrix}\right|_{\mathbf{y}=\bar{\mathbf{y}}} \in \mathbb{R}^{N_\mathrm{a}N_\mathrm{g}\times N_\mathrm{a}N_\mathrm{p}}, \quad \text{(Layer 2, Eq. S2b)}$$

which can be understood as measuring direct developmental bias from the genotype. Note that the main block diagonal is zero.

Direct-effect matrices also include matrices measuring direct plasticity and direct niche construction. Indeed, the block matrix of *direct effects of a mutant's environment on her phenotype* is

$$\left.\frac{\partial \mathbf{x}^{\intercal}}{\partial \boldsymbol{\epsilon}}\right|_{\mathbf{y}=\bar{\mathbf{y}}} = \left.\begin{pmatrix} \mathbf{0} & \frac{\partial \mathbf{x}_2^{\intercal}}{\partial \boldsymbol{\epsilon}_1} & \cdots & \mathbf{0} & \mathbf{0} \\ \mathbf{0} & \mathbf{0} & \cdots & \mathbf{0} & \mathbf{0} \\ \vdots & \vdots & \ddots & \vdots & \vdots \\ \mathbf{0} & \mathbf{0} & \cdots & \mathbf{0} & \frac{\partial \mathbf{x}_{N_\mathrm{a}}^{\intercal}}{\partial \boldsymbol{\epsilon}_{N_\mathrm{a}-1}} \\ \mathbf{0} & \mathbf{0} & \cdots & \mathbf{0} & \mathbf{0} \end{pmatrix}\right|_{\mathbf{y}=\bar{\mathbf{y}}} \in \mathbb{R}^{N_\mathrm{a}N_\mathrm{e}\times N_\mathrm{a}N_\mathrm{p}}, \quad \text{(Layer 2, Eq. S2c)}$$

which can be understood as measuring the direct plasticity of the phenotype (Noble *et al.*, 2019). In turn, the block matrix of *direct effects of a mutant's phenotype or genotype on her environment* is

$$\left.\frac{\partial \boldsymbol{\epsilon}^{\intercal}}{\partial \boldsymbol{\zeta}}\right|_{\mathbf{y}=\bar{\mathbf{y}}} = \left.\begin{pmatrix} \frac{\partial \boldsymbol{\epsilon}_1^{\intercal}}{\partial \boldsymbol{\zeta}_1} & \mathbf{0} & \cdots & \mathbf{0} & \mathbf{0} \\ \mathbf{0} & \frac{\partial \boldsymbol{\epsilon}_2^{\intercal}}{\partial \boldsymbol{\zeta}_2} & \cdots & \mathbf{0} & \mathbf{0} \\ \vdots & \vdots & \ddots & \vdots & \vdots \\ \mathbf{0} & \mathbf{0} & \cdots & \frac{\partial \boldsymbol{\epsilon}_{N_\mathrm{a}-1}^{\intercal}}{\partial \boldsymbol{\zeta}_{N_\mathrm{a}-1}} & \mathbf{0} \\ \mathbf{0} & \mathbf{0} & \cdots & \mathbf{0} & \frac{\partial \boldsymbol{\epsilon}_{N_\mathrm{a}}^{\intercal}}{\partial \boldsymbol{\zeta}_{N_\mathrm{a}}} \end{pmatrix}\right|_{\mathbf{y}=\bar{\mathbf{y}}} \quad \text{(Layer 2, Eq. S2d)}$$

for $\boldsymbol{\zeta} \in \{\mathbf{x},\mathbf{y}\}$, which can be understood as measuring direct niche construction by the phenotype or genotype. The equality (Layer 2, Eq. S2d) follows from the environmental constraint in Eq. (2) since the environment faced by a mutant at a given age is directly affected by the mutant phenotype or genotype at the same age only (i.e., $\partial \boldsymbol{\epsilon}_j^{\intercal}/\partial \boldsymbol{\zeta}_a = \mathbf{0}$ for $a \neq j$).

Direct-effect matrices also include a matrix describing direct mutual environmental dependence. This is measured by the block matrix of *direct effects of a mutant's environment on itself*

$$\left.\frac{\partial \boldsymbol{\epsilon}^{\intercal}}{\partial \boldsymbol{\epsilon}}\right|_{\mathbf{y}=\bar{\mathbf{y}}} = \left.\begin{pmatrix} \frac{\partial \boldsymbol{\epsilon}_1^{\intercal}}{\partial \boldsymbol{\epsilon}_1} & \mathbf{0} & \cdots & \mathbf{0} & \mathbf{0} \\ \mathbf{0} & \frac{\partial \boldsymbol{\epsilon}_2^{\intercal}}{\partial \boldsymbol{\epsilon}_2} & \cdots & \mathbf{0} & \mathbf{0} \\ \vdots & \vdots & \ddots & \vdots & \vdots \\ \mathbf{0} & \mathbf{0} & \cdots & \frac{\partial \boldsymbol{\epsilon}_{N_\mathrm{a}-1}^{\intercal}}{\partial \boldsymbol{\epsilon}_{N_\mathrm{a}-1}} & \mathbf{0} \\ \mathbf{0} & \mathbf{0} & \cdots & \mathbf{0} & \frac{\partial \boldsymbol{\epsilon}_{N_\mathrm{a}}^{\intercal}}{\partial \boldsymbol{\epsilon}_{N_\mathrm{a}}} \end{pmatrix}\right|_{\mathbf{y}=\bar{\mathbf{y}}} = \mathbf{I} \in \mathbb{R}^{N_\mathrm{a}N_\mathrm{e}\times N_\mathrm{a}N_\mathrm{e}}. \quad \text{(Layer 2, Eq. S3)}$$

The first equality follows from the environmental constraint (Eq. 2) and the second equality follows from our assumption that environmental traits are mutually independent, so $\partial \boldsymbol{\epsilon}_a^{\intercal}/\partial \boldsymbol{\epsilon}_a|_{\mathbf{y}=\bar{\mathbf{y}}} = \mathbf{I}$ for all $a \in \{1,\ldots,N_\mathrm{a}\}$. It is conceptually useful to write $\partial \boldsymbol{\epsilon}^{\intercal}/\partial \boldsymbol{\epsilon}|_{\mathbf{y}=\bar{\mathbf{y}}}$ rather than only $\mathbf{I}$, and we do so throughout.

Additionally, direct-effect matrices include matrices describing direct social developmental bias, which includes the direct effects of extra-genetic inheritance and indirect genetic effects. The block matrix of *direct effects of social partners' phenotype or genotype on a mutant's phenotype* is

$$\left.\frac{\partial \mathbf{x}^{\intercal}}{\partial \bar{\boldsymbol{\zeta}}}\right|_{\mathbf{y}=\bar{\mathbf{y}}} = \left.\begin{pmatrix} \mathbf{0} & \frac{\partial \mathbf{x}_2^{\intercal}}{\partial \bar{\boldsymbol{\zeta}}_1} & \cdots & \frac{\partial \mathbf{x}_{N_\mathrm{a}}^{\intercal}}{\partial \bar{\boldsymbol{\zeta}}_1} \\ \mathbf{0} & \frac{\partial \mathbf{x}_2^{\intercal}}{\partial \bar{\boldsymbol{\zeta}}_2} & \cdots & \frac{\partial \mathbf{x}_{N_\mathrm{a}}^{\intercal}}{\partial \bar{\boldsymbol{\zeta}}_2} \\ \vdots & \vdots & \ddots & \vdots \\ \mathbf{0} & \frac{\partial \mathbf{x}_2^{\intercal}}{\partial \bar{\boldsymbol{\zeta}}_{N_\mathrm{a}}} & \cdots & \frac{\partial \mathbf{x}_{N_\mathrm{a}}^{\intercal}}{\partial \bar{\boldsymbol{\zeta}}_{N_\mathrm{a}}} \end{pmatrix}\right|_{\mathbf{y}=\bar{\mathbf{y}}}, \quad \text{(Layer 2, Eq. S4)}$$

for $\bar{\boldsymbol{\zeta}} \in \{\bar{\mathbf{x}},\bar{\mathbf{y}}\}$, where the equality follows because the phenotype $\mathbf{x}_1$ at the initial age is constant by assumption. The matrix in Layer 2, Eq. S4 can be understood as measuring direct social developmental bias from either the phenotype or genotype, and mechanistically measures the direct effects of extra-genetic inheritance and indirect genetic effects. This matrix can be less sparse than direct-effect matrices above because the mutant's phenotype can be affected by the phenotype or genotype of social partners of *any* age.

Direct-effect matrices also include matrices describing direct social niche construction. The block matrix of *direct effects of social partners' phenotype or genotype on a mutant's environment* is

$$\left.\frac{\partial \boldsymbol{\epsilon}^{\intercal}}{\partial \bar{\boldsymbol{\zeta}}}\right|_{\mathbf{y}=\bar{\mathbf{y}}} \equiv \left.\begin{pmatrix} \frac{\partial \boldsymbol{\epsilon}_1^{\intercal}}{\partial \bar{\boldsymbol{\zeta}}_1} & \cdots & \frac{\partial \boldsymbol{\epsilon}_{N_\mathrm{a}}^{\intercal}}{\partial \bar{\boldsymbol{\zeta}}_1} \\ \vdots & \ddots & \vdots \\ \frac{\partial \boldsymbol{\epsilon}_1^{\intercal}}{\partial \bar{\boldsymbol{\zeta}}_{N_\mathrm{a}}} & \cdots & \frac{\partial \boldsymbol{\epsilon}_{N_\mathrm{a}}^{\intercal}}{\partial \bar{\boldsymbol{\zeta}}_{N_\mathrm{a}}} \end{pmatrix}\right|_{\mathbf{y}=\bar{\mathbf{y}}}, \quad \text{(Layer 2, Eq. S5)}$$

for $\bar{\boldsymbol{\zeta}} \in \{\bar{\mathbf{x}},\bar{\mathbf{y}}\}$, which can be understood as measuring direct social niche construction by either the phenotype or genotype. This matrix does not contain any zero entries in general because the mutant's environment at any age can be affected by the phenotype or genotype of social partners of any age.

We use the above definitions to form direct-effect matrices involving the geno-phenotype. The block matrix of *direct effects of a mutant's geno-phenotype on her geno-phenotype* is

$$\left.\frac{\partial \mathbf{z}^\intercal}{\partial \mathbf{z}}\right|_{\mathbf{y}=\bar{\mathbf{y}}} \equiv \left.\begin{pmatrix} \frac{\partial \mathbf{x}^\intercal}{\partial \mathbf{x}} & \frac{\partial \mathbf{y}^\intercal}{\partial \mathbf{x}} \\ \frac{\partial \mathbf{x}^\intercal}{\partial \mathbf{y}} & \frac{\partial \mathbf{y}^\intercal}{\partial \mathbf{y}} \end{pmatrix}\right|_{\mathbf{y}=\bar{\mathbf{y}}} = \left.\begin{pmatrix} \frac{\partial \mathbf{x}^\intercal}{\partial \mathbf{x}} & \mathbf{0} \\ \frac{\partial \mathbf{x}^\intercal}{\partial \mathbf{y}} & \mathbf{I} \end{pmatrix}\right|_{\mathbf{y}=\bar{\mathbf{y}}} \quad \text{(Layer 2, Eq. S6)}$$

$$\in \mathbb{R}^{N_\mathrm{a}(N_\mathrm{p}+N_\mathrm{g})\times N_\mathrm{a}(N_\mathrm{p}+N_\mathrm{g})},$$

which measures direct developmental bias of the geno-phenotype, and where the equality follows because genotypic traits are developmentally independent by assumption. The block matrix of *direct effects of a mutant's geno-phenotype on her environment* is

$$\left.\frac{\partial \boldsymbol{\epsilon}^\intercal}{\partial \mathbf{z}}\right|_{\mathbf{y}=\bar{\mathbf{y}}} \equiv \left.\left(\frac{\partial \boldsymbol{\epsilon}^\intercal}{\partial \mathbf{x}}; \frac{\partial \boldsymbol{\epsilon}^\intercal}{\partial \mathbf{y}}\right)\right|_{\mathbf{y}=\bar{\mathbf{y}}} \in \mathbb{R}^{N_\mathrm{a}(N_\mathrm{p}+N_\mathrm{g})\times N_\mathrm{a}N_\mathrm{e}}, \quad \text{(Layer 2, Eq. S7)}$$

which measures direct niche construction by the geno-phenotype. The block matrix of *direct effects of social partners' geno-phenotypes on a mutant's environment* is

$$\left.\frac{\partial \boldsymbol{\epsilon}^\intercal}{\partial \bar{\mathbf{z}}}\right|_{\mathbf{y}=\bar{\mathbf{y}}} \equiv \left.\left(\frac{\partial \boldsymbol{\epsilon}^\intercal}{\partial \bar{\mathbf{x}}}; \frac{\partial \boldsymbol{\epsilon}^\intercal}{\partial \bar{\mathbf{y}}}\right)\right|_{\mathbf{y}=\bar{\mathbf{y}}} \in \mathbb{R}^{N_\mathrm{a}(N_\mathrm{p}+N_\mathrm{g})\times N_\mathrm{a}N_\mathrm{e}}, \quad \text{(Layer 2, Eq. S8)}$$

which measures direct social niche construction by partners' geno-phenotypes. The block matrix of *direct effects of a mutant's environment on her geno-phenotype* is

$$\left.\frac{\partial \mathbf{z}^\intercal}{\partial \boldsymbol{\epsilon}}\right|_{\mathbf{y}=\bar{\mathbf{y}}} \equiv \left.\begin{pmatrix} \frac{\partial \mathbf{x}^\intercal}{\partial \boldsymbol{\epsilon}} & \frac{\partial \mathbf{y}^\intercal}{\partial \boldsymbol{\epsilon}} \end{pmatrix}\right|_{\mathbf{y}=\bar{\mathbf{y}}} = \left.\begin{pmatrix} \frac{\partial \mathbf{x}^\intercal}{\partial \boldsymbol{\epsilon}} & \mathbf{0} \end{pmatrix}\right|_{\mathbf{y}=\bar{\mathbf{y}}} \in \mathbb{R}^{N_\mathrm{a}N_\mathrm{e}\times N_\mathrm{a}(N_\mathrm{p}+N_\mathrm{g})}, \quad \text{(Layer 2, Eq. S9)}$$

which measures the direct plasticity of the geno-phenotype, and where the equality follows because genotypic traits are developmentally independent.

We will see that the evolutionary dynamics of the environment depends on a matrix measuring "joint" direct niche construction. This matrix is the transpose of the matrix of *direct social effects of a focal individual's geno-phenotype on hers and her partners' environment*

$$\left.\frac{\partial(\boldsymbol{\epsilon}+\check{\boldsymbol{\epsilon}})}{\partial \mathbf{z}^\intercal}\right|_{\mathbf{y}=\bar{\mathbf{y}}} = \left.\left(\frac{\partial \boldsymbol{\epsilon}}{\partial \mathbf{z}^\intercal} + \frac{\partial \boldsymbol{\epsilon}}{\partial \bar{\mathbf{z}}^\intercal}\right)\right|_{\mathbf{y}=\bar{\mathbf{y}}} \in \mathbb{R}^{N_\mathrm{a}N_\mathrm{e}\times N_\mathrm{a}(N_\mathrm{p}+N_\mathrm{g})}, \quad \text{(Layer 2, Eq. S10)}$$

where we denote by $\check{\boldsymbol{\epsilon}}$ the environment a resident experiences when she develops in the context of mutants (a donor perspective for the mutant). Thus, this matrix can be interpreted as joint direct niche construction by the geno-phenotype. Note that the second term on the right-hand side of Layer 2, Eq. S10 is the direct effects of social partners' geno-phenotypes on a focal mutant (a recipient perspective for the mutant). Hence, joint direct niche construction by the geno-phenotype as described by Layer 2, Eq. S10 can be equivalently interpreted either from a donor or a recipient perspective.

#### S3.3 Layer 3: total immediate effects

We now proceed to write the equations of the next layer of the evo-devo process, that of total immediate effects. Total-immediate-effect matrices measure the total effects that a variable has on another variable only at a given age, thus without considering the downstream effects over development. With the developmental and environmental constraints assumed, if there are no environmental traits, total immediate effect matrices ($\delta\boldsymbol{\zeta}^\intercal/\delta\boldsymbol{\xi}$) reduce to direct effect matrices ($\partial\boldsymbol{\zeta}^\intercal/\partial\boldsymbol{\xi}$).

Total-immediate-effect matrices include total immediate selection gradients, which capture some of the effects of niche construction. The *total immediate selection gradient of the phenotype, genotype, or geno-phenotype* is

$$\left.\frac{\delta w}{\delta \boldsymbol{\zeta}}\right|_{\mathbf{y}=\bar{\mathbf{y}}} = \left.\left(\frac{\partial w}{\partial \boldsymbol{\zeta}} + \frac{\partial \boldsymbol{\epsilon}^\intercal}{\partial \boldsymbol{\zeta}}\frac{\partial w}{\partial \boldsymbol{\epsilon}}\right)\right|_{\mathbf{y}=\bar{\mathbf{y}}}, \quad \text{(Layer 3, Eq. S1)}$$

for $\boldsymbol{\zeta} \in \{\mathbf{x},\mathbf{y},\mathbf{z}\}$. Here, the total immediate selection gradient of $\boldsymbol{\zeta}$ depends on direct directional selection on $\boldsymbol{\zeta}$, direct niche construction by $\boldsymbol{\zeta}$, and direct environmental sensitivity of selection. Thus, total immediate selection gradients measure total immediate directional selection, which is directional selection in the fitness landscape modified by the interaction of niche construction and environmental sensitivity of selection. In a standard quantitative genetics framework, the total immediate selection gradients correspond to Lande's (1979) selection gradient if the environmental traits are not explicitly included in the analysis.

Total immediate selection on the environment equals direct selection on the environment because we assume environmental traits are mutually independent. The *total immediate selection gradient of the environment* is

$$\left.\frac{\delta w}{\delta \boldsymbol{\epsilon}}\right|_{\mathbf{y}=\bar{\mathbf{y}}} = \left.\left(\frac{\partial \boldsymbol{\epsilon}^\intercal}{\partial \boldsymbol{\epsilon}}\frac{\partial w}{\partial \boldsymbol{\epsilon}}\right)\right|_{\mathbf{y}=\bar{\mathbf{y}}} \in \mathbb{R}^{N_\mathrm{a}N_\mathrm{e}\times 1}. \quad \text{(Layer 3, Eq. S2)}$$

Given our assumption that environmental traits are mutually independent, the matrix of direct effects of the environment on itself is the identity matrix. Thus, the total immediate selection gradient of the environment equals the selection gradient of the environment.

Total-immediate-effect matrices also include matrices describing total immediate developmental bias, which capture additional effects of niche construction. The block matrix of *total immediate effects of the phenotype, genotype, social partner's phenotype, or social partner's genotype on a mutant's phenotype* is

$$\left.\frac{\delta \mathbf{x}^\intercal}{\delta \boldsymbol{\zeta}}\right|_{\mathbf{y}=\bar{\mathbf{y}}} = \left.\left(\frac{\partial \mathbf{x}^\intercal}{\partial \boldsymbol{\zeta}} + \frac{\partial \boldsymbol{\epsilon}^\intercal}{\partial \boldsymbol{\zeta}}\frac{\partial \mathbf{x}^\intercal}{\partial \boldsymbol{\epsilon}}\right)\right|_{\mathbf{y}=\bar{\mathbf{y}}}, \quad \text{(Layer 3, Eq. S3)}$$

for $\boldsymbol{\zeta} \in \{\mathbf{x},\mathbf{y},\bar{\mathbf{x}},\bar{\mathbf{y}}\}$. Here, the total immediate effects of $\boldsymbol{\zeta}$ on the phenotype depend on the direct developmental bias from $\boldsymbol{\zeta}$, direct niche construction by $\boldsymbol{\zeta}$, and the direct plasticity of the phenotype. Consequently, total immediate effects on the phenotype can be interpreted as measuring total immediate developmental bias, which measures developmental bias in the developmental process modified by the interaction of niche construction and plasticity.

Moreover, total immediate-effect matrices include matrices describing total immediate plasticity of the phenotype, which equals plasticity of the phenotype because environmental traits are mutually independent by assumption. The block matrix of *total immediate effects of a mutant's environment on her phenotype* is

$$\left.\frac{\delta \mathbf{x}^\intercal}{\delta \boldsymbol{\epsilon}}\right|_{\mathbf{y}=\bar{\mathbf{y}}} = \left.\frac{\partial \boldsymbol{\epsilon}^\intercal}{\partial \boldsymbol{\epsilon}}\frac{\partial \mathbf{x}^\intercal}{\partial \boldsymbol{\epsilon}}\right|_{\mathbf{y}=\bar{\mathbf{y}}} \in \mathbb{R}^{N_\mathrm{a}N_\mathrm{e}\times N_\mathrm{a}N_\mathrm{p}}. \quad \text{(Layer 3, Eq. S4)}$$

Given our assumption that environmental traits are mutually independent, the matrix of direct effects of the environment on itself is the identity matrix. Thus, the total immediate plasticity of the phenotype equals the direct plasticity of the phenotype.

We use the above definitions to form a matrix quantifying the total immediate developmental bias of the geno-phenotype. This is the block matrix of *total immediate effects of a mutant's geno-phenotype on her geno-phenotype*

$$\left.\frac{\delta \mathbf{z}^\intercal}{\delta \mathbf{z}}\right|_{\mathbf{y}=\bar{\mathbf{y}}} = \left.\left(\frac{\partial \mathbf{z}^\intercal}{\partial \mathbf{z}} + \frac{\partial \boldsymbol{\epsilon}^\intercal}{\partial \mathbf{z}}\frac{\partial \mathbf{z}^\intercal}{\partial \boldsymbol{\epsilon}}\right)\right|_{\mathbf{y}=\bar{\mathbf{y}}} \in \mathbb{R}^{N_\mathrm{a}(N_\mathrm{p}+N_\mathrm{g})\times N_\mathrm{a}(N_\mathrm{p}+N_\mathrm{g})}.$$

(Layer 3, Eq. S5)

Consequently, the total immediate developmental bias of the geno-phenotype depends on the direct developmental bias of the geno-phenotype, direct niche construction by the geno-phenotype, and direct plasticity of the geno-phenotype.

#### S3.4 Layer 4: total effects

We now move to write the equations for the next layer of the evo-devo process, that of total-effect matrices. Total-effect matrices measure the total effects of a variable on another one over the individual's life, thus considering the downstream effects over development, but before the effects of social development have stabilized in the population. More generally, total-effect matrices include matrices that give the sensitivity to perturbations of the solution of a recurrence of the form in Eq. (1).

The total effects of the phenotype on itself describe the *developmental feedback* of the phenotype. This is given by the block matrix of *total effects of a mutant's phenotype on her phenotype*

$$\begin{aligned}\left.\frac{\mathrm{d}\mathbf{x}^\intercal}{\mathrm{d}\mathbf{x}}\right|_{\mathbf{y}=\bar{\mathbf{y}}} &= \left.\left(2\mathbf{I} - \frac{\delta \mathbf{x}^\intercal}{\delta \mathbf{x}}\right)^{-1}\right|_{\mathbf{y}=\bar{\mathbf{y}}} \\ &= \left.\sum_{a=1}^{N_\mathrm{a}}\left(\frac{\delta \mathbf{x}^\intercal}{\delta \mathbf{x}} - \mathbf{I}\right)^{a-1}\right|_{\mathbf{y}=\bar{\mathbf{y}}} \in \mathbb{R}^{N_\mathrm{a}N_\mathrm{p}\times N_\mathrm{a}N_\mathrm{p}},\end{aligned}$$

(Layer 4, Eq. S1)

which is always invertible (section S5.1, Eq. S5.1.15) and where the last equality follows by the geometric series of matrices. This matrix can be interpreted as a lifetime collection of total immediate effects of the phenotype on itself. Also, the developmental feedback of the phenotype can be seen as describing the total developmental bias of the phenotype. More generally, Layer 4, Eq. S1 gives the sensitivity of the solution $\mathbf{x}$ of the recurrence (1) to perturbations in the solution at other times (ages): in particular, $\mathrm{d}x_{kj}/\mathrm{d}x_{ia}$ gives the sensitivity of the solution $x_{kj}$ of the $k$-th variable at time $j$ to perturbations in the solution $x_{ia}$ of the $i$-th variable at time $a$. Developmental feedback may cause major phenotypic effects at subsequent ages as its block entries involve matrix products (Eq. 12).

The total effects of the genotype on the phenotype are a mechanistic analogue of Fisher's additive effects of allelic substitution and of Wagner's developmental matrix. The block matrix of *total effects of a mutant's genotype on her phenotype* is given by

$$\left.\frac{\mathrm{d}\mathbf{x}^\intercal}{\mathrm{d}\mathbf{y}}\right|_{\mathbf{y}=\bar{\mathbf{y}}} = \left.\left(\frac{\delta \mathbf{x}^\intercal}{\delta \mathbf{y}}\frac{\mathrm{d}\mathbf{x}^\intercal}{\mathrm{d}\mathbf{x}}\right)\right|_{\mathbf{y}=\bar{\mathbf{y}}} \in \mathbb{R}^{N_\mathrm{a}N_\mathrm{g}\times N_\mathrm{a}N_\mathrm{p}},$$

(Layer 4, Eq. S2)

which is singular because the developmentally initial phenotype is not affected by the genotype (by our assumption that the initial phenotype is constant) and the developmentally final genotypic traits do not affect the phenotype (by our assumption that individuals do not survive after the final age; so $\mathrm{d}\mathbf{x}^\intercal/\mathrm{d}\mathbf{y}|_{\mathbf{y}=\bar{\mathbf{y}}}$ has rows and columns that are zero; section S5.2, Eq. S5.2.16). From Layer 4, Eq. S2, this matrix can be interpreted as involving a developmentally immediate pulse caused by a change in genotypic traits followed by the triggered developmental feedback of the phenotype. By giving the total effects of a perturbation in the genotype on the phenotype, the entries of this matrix are a mechanistic analogue of Fisher's additive effect of allelic substitution, which he defined as regression coefficients (his $\alpha$; see Eq. I of Fisher 1918 and p. 72 of Lynch and Walsh 1998). Also, this matrix is a mechanistic analogue of Wagner's (1984, 1989) developmental matrix (his $\mathbf{B}$) (see also Martin 2014), Rice's (2002) rank-1 $\mathbf{D}$ tensor, and Morrissey's (2015) total effect matrix (his $\boldsymbol{\Phi}$, but not Morrissey's (2014) $\boldsymbol{\Phi}$, which is a regression-based form of $\mathrm{d}\mathbf{x}^\intercal/\mathrm{d}\mathbf{x}$) (interpreting these authors' partial derivatives as total derivatives, although using derivatives rather than regression coefficients violates the standard partition of phenotypic variance into genetic and "environmental" variances, as explained in the main text section 5.1). More generally, interpreting $\mathbf{y}$ as parameters affecting the recurrence (1) over $\mathbf{x}$, Layer 4, Eq. S2 gives the sensitivity of the solution $\mathbf{x}$ to perturbation in the parameters at other times (ages): in particular, $\mathrm{d}x_{kj}/\mathrm{d}y_{ia}$ gives the sensitivity of the solution $x_{kj}$ of the $k$-th variable at time $j$ to perturbations in the $i$-th parameter $y_{ia}$ at time $a$. The definition of total effects of the genotype on the phenotype in terms of derivatives (Layer 4, Eq. S2) differs from Fisher's in terms of regression coefficients both in that it reveals its structure and so it can be used for evo-devo dynamically sufficient analysis, and in that regression coefficients of phenotype to genotype are uncorrelated with residuals whereas the derivative analogues need not be.

The total effects of the environment on the phenotype measure the total plasticity of the phenotype, considering downstream effects over development. This is given by the block matrix of *total effects of a mutant's environment* *on her phenotype*

$$\left.\frac{d\mathbf{x}^\intercal}{d\boldsymbol{\epsilon}}\right|_{\mathbf{y}=\bar{\mathbf{y}}} = \left.\left(\frac{\delta\mathbf{x}^\intercal}{\delta\boldsymbol{\epsilon}}\frac{d\mathbf{x}^\intercal}{d\mathbf{x}}\right)\right|_{\mathbf{y}=\bar{\mathbf{y}}} \in \mathbb{R}^{N_a N_e \times N_a N_p}. \qquad \text{(Layer 4, Eq. S3)}$$

Thus, the total plasticity of the phenotype can be interpreted as a developmentally immediate pulse of plastic change in the phenotype followed by the triggered developmental feedback of the phenotype.

The total effects of social partners' genotype or phenotype on the phenotype measure the total *social* developmental bias of the phenotype. The block matrix of *total effects of social partners' phenotype or genotype on a mutant's phenotype* is

$$\left.\frac{d\mathbf{x}^\intercal}{d\bar{\boldsymbol{\zeta}}}\right|_{\mathbf{y}=\bar{\mathbf{y}}} = \left.\left(\frac{\delta\mathbf{x}^\intercal}{\delta\bar{\boldsymbol{\zeta}}}\frac{d\mathbf{x}^\intercal}{d\mathbf{x}}\right)\right|_{\mathbf{y}=\bar{\mathbf{y}}} \qquad \text{(Layer 4, Eq. S4)}$$

for $\bar{\boldsymbol{\zeta}} \in \{\bar{\mathbf{x}}, \bar{\mathbf{y}}\}$. This matrix can be interpreted as measuring total social developmental bias of the phenotype from phenotype or genotype, as well as the total effects on the phenotype of extra-genetic inheritance, and the total indirect genetic effects. In particular, the matrix of total social developmental bias of the phenotype from phenotype, $d\mathbf{x}^\intercal/d\bar{\mathbf{x}}|_{\mathbf{y}=\bar{\mathbf{y}}}$, is a mechanistic version of the matrix of interaction coefficients in the indirect genetic effects literature (i.e., $\boldsymbol{\Psi}$ in Eq. 17 of Moore *et al.* 1997, which is defined as a matrix of regression coefficients). From Layer 4, Eq. S4, the total social developmental bias of the phenotype can be interpreted as a developmentally immediate pulse of phenotype change caused by a change in social partners' traits followed by the triggered developmental feedback of the mutant's phenotype.

The total effects on the genotype are simple since genotypic traits are developmentally independent by assumption. The block matrix of *total effects of a mutant's genotype on itself* is

$$\left.\frac{d\mathbf{y}^\intercal}{d\mathbf{y}}\right|_{\mathbf{y}=\bar{\mathbf{y}}} = \mathbf{I} \in \mathbb{R}^{N_a N_g \times N_a N_g}, \qquad \text{(Layer 4, Eq. S5)}$$

and the block matrix of *total effects of a vector* $\boldsymbol{\zeta} \in \{\mathbf{x}, \boldsymbol{\epsilon}, \bar{\mathbf{x}}, \bar{\mathbf{y}}, \bar{\mathbf{z}}, \bar{\boldsymbol{\epsilon}}, \bar{\mathbf{m}}\}$ *on a mutant's genotype* is

$$\left.\frac{d\mathbf{y}^\intercal}{d\boldsymbol{\zeta}}\right|_{\mathbf{y}=\bar{\mathbf{y}}} = \mathbf{0},$$

(section S5.2, Eq. S5.2.13).

We can use some of the previous total-effect matrices to construct the following total-effect matrices involving the geno-phenotype. The block matrix of *total effects of a mutant's phenotype on her geno-phenotype* is

$$\begin{aligned}\left.\frac{d\mathbf{z}^\intercal}{d\mathbf{x}}\right|_{\mathbf{y}=\bar{\mathbf{y}}} &\equiv \left.\begin{pmatrix}\frac{d\mathbf{x}^\intercal}{d\mathbf{x}} & \frac{d\mathbf{y}^\intercal}{d\mathbf{x}}\end{pmatrix}\right|_{\mathbf{y}=\bar{\mathbf{y}}} \\ &= \left.\begin{pmatrix}\frac{d\mathbf{x}^\intercal}{d\mathbf{x}} & \mathbf{0}\end{pmatrix}\right|_{\mathbf{y}=\bar{\mathbf{y}}} \in \mathbb{R}^{N_a N_p \times N_a (N_p+N_g)},\end{aligned} \qquad \text{(Layer 4, Eq. S6)}$$

measuring total developmental bias of the geno-phenotype from the phenotype. The block matrix of *total effects of the genotype on her geno-phenotype* is

$$\begin{aligned}\left.\frac{d\mathbf{z}^\intercal}{d\mathbf{y}}\right|_{\mathbf{y}=\bar{\mathbf{y}}} &\equiv \left.\begin{pmatrix}\frac{d\mathbf{x}^\intercal}{d\mathbf{y}} & \frac{d\mathbf{y}^\intercal}{d\mathbf{y}}\end{pmatrix}\right|_{\mathbf{y}=\bar{\mathbf{y}}} \\ &= \left.\begin{pmatrix}\frac{d\mathbf{x}^\intercal}{d\mathbf{y}} & \mathbf{I}\end{pmatrix}\right|_{\mathbf{y}=\bar{\mathbf{y}}} \in \mathbb{R}^{N_a N_g \times N_a (N_p+N_g)},\end{aligned} \qquad \text{(Layer 4, Eq. S7)}$$

measuring total developmental bias of the geno-phenotype from the genotype. This matrix $d\mathbf{z}^\intercal/d\mathbf{y}|_{\mathbf{y}=\bar{\mathbf{y}}}$ is singular because any matrix with fewer rows than columns is singular (Horn and Johnson, 2013, p. 14). This singularity is important when we consider mechanistic additive genetic covariances (Layer 6, section 5.1). Now, the block matrix of *total effects of a mutant's geno-phenotype on her geno-phenotype* is

$$\begin{aligned}\left.\frac{d\mathbf{z}^\intercal}{d\mathbf{z}}\right|_{\mathbf{y}=\bar{\mathbf{y}}} &\equiv \left.\begin{pmatrix}\frac{d\mathbf{x}^\intercal}{d\mathbf{x}} & \frac{d\mathbf{y}^\intercal}{d\mathbf{x}} \\ \frac{d\mathbf{x}^\intercal}{d\mathbf{y}} & \frac{d\mathbf{y}^\intercal}{d\mathbf{y}}\end{pmatrix}\right|_{\mathbf{y}=\bar{\mathbf{y}}} = \left.\begin{pmatrix}\frac{d\mathbf{x}^\intercal}{d\mathbf{x}} & \mathbf{0} \\ \frac{d\mathbf{x}^\intercal}{d\mathbf{y}} & \mathbf{I}\end{pmatrix}\right|_{\mathbf{y}=\bar{\mathbf{y}}} \\ &= \left.\left(2\mathbf{I} - \frac{\delta\mathbf{z}^\intercal}{\delta\mathbf{z}}\right)^{-1}\right|_{\mathbf{y}=\bar{\mathbf{y}}} = \sum_{a=1}^{N_a}\left(\frac{\delta\mathbf{z}^\intercal}{\delta\mathbf{z}} - \mathbf{I}\right)^{a-1} \qquad \text{(Layer 4, Eq. S8)} \\ &\in \mathbb{R}^{N_a (N_p+N_g) \times N_a (N_p+N_g)},\end{aligned}$$

which can be interpreted as measuring the developmental feedback of the geno-phenotype (section S5.4, Eq. S5.4.4). Since $d\mathbf{z}^\intercal/d\mathbf{z}|_{\mathbf{y}=\bar{\mathbf{y}}}$ is square and block lower triangular, and since $d\mathbf{x}^\intercal/d\mathbf{x}|_{\mathbf{y}=\bar{\mathbf{y}}}$ is invertible (section S5.1, Eq. S5.1.15), we have that $d\mathbf{z}^\intercal/d\mathbf{z}|_{\mathbf{y}=\bar{\mathbf{y}}}$ is invertible.

Moreover, the total effects of the phenotype and genotype on the environment quantify total niche construction. Total niche construction by the phenotype is quantified by the block matrix of *total effects of a mutant's phenotype on her environment*

$$\begin{aligned}\left.\frac{d\boldsymbol{\epsilon}^\intercal}{d\mathbf{x}}\right|_{\mathbf{y}=\bar{\mathbf{y}}} &= \left.\left(\frac{d\mathbf{x}^\intercal}{d\mathbf{x}}\frac{\partial\boldsymbol{\epsilon}^\intercal}{\partial\mathbf{x}}\right)\right|_{\mathbf{y}=\bar{\mathbf{y}}} \\ &= \left.\left(\frac{d\mathbf{z}^\intercal}{d\mathbf{x}}\frac{\partial\boldsymbol{\epsilon}^\intercal}{\partial\mathbf{z}}\right)\right|_{\mathbf{y}=\bar{\mathbf{y}}} \in \mathbb{R}^{N_a N_p \times N_a N_e},\end{aligned} \qquad \text{(Layer 4, Eq. S9)}$$

which can be interpreted as showing that developmental feedback of the phenotype occurs first and then direct niche-constructing effects by the phenotype follow. Similarly, total niche construction by the genotype is quantified by the block matrix of *total effects of a mutant's genotype on her environment*

$$\begin{aligned}\left.\frac{d\boldsymbol{\epsilon}^\intercal}{d\mathbf{y}}\right|_{\mathbf{y}=\bar{\mathbf{y}}} &= \left.\left(\frac{d\mathbf{x}^\intercal}{d\mathbf{y}}\frac{\partial\boldsymbol{\epsilon}^\intercal}{\partial\mathbf{x}} + \frac{\partial\boldsymbol{\epsilon}^\intercal}{\partial\mathbf{y}}\right)\right|_{\mathbf{y}=\bar{\mathbf{y}}} \\ &= \left.\left(\frac{d\mathbf{z}^\intercal}{d\mathbf{y}}\frac{\partial\boldsymbol{\epsilon}^\intercal}{\partial\mathbf{z}}\right)\right|_{\mathbf{y}=\bar{\mathbf{y}}} \in \mathbb{R}^{N_a N_g \times N_a N_e},\end{aligned} \qquad \text{(Layer 4, Eq. S10)}$$

which depends on direct niche construction by the genotype and on total developmental bias of the phenotype from the genotype followed by niche construction by the phenotype. The analogous relationship holds for total niche construction by the geno-phenotype, quantified by the block matrix of *total effects of a mutant's geno-phenotype on her environment*

$$\left.\frac{\mathrm{d}\boldsymbol{\epsilon}^\intercal}{\mathrm{d}\mathbf{z}}\right|_{\mathbf{y}=\bar{\mathbf{y}}} = \left.\left(\frac{\mathrm{d}\mathbf{z}^\intercal}{\mathrm{d}\mathbf{z}}\frac{\partial\boldsymbol{\epsilon}^\intercal}{\partial\mathbf{z}}\right)\right|_{\mathbf{y}=\bar{\mathbf{y}}} \in \mathbb{R}^{N_\mathrm{a}(N_\mathrm{p}+N_\mathrm{g})\times N_\mathrm{a}N_\mathrm{e}}, \quad \text{(Layer 4, Eq. S11)}$$

which depends on the developmental feedback of the geno-phenotype and direct niche construction by the geno-phenotype.

The total effects of the environment on itself quantify environmental feedback. The block matrix of *total effects of a mutant's environment on her environment* is

$$\left.\frac{\mathrm{d}\boldsymbol{\epsilon}^\intercal}{\mathrm{d}\boldsymbol{\epsilon}}\right|_{\mathbf{y}=\bar{\mathbf{y}}} = \left.\left(\frac{\partial\boldsymbol{\epsilon}^\intercal}{\partial\boldsymbol{\epsilon}}+\frac{\mathrm{d}\mathbf{x}^\intercal}{\mathrm{d}\boldsymbol{\epsilon}}\frac{\partial\boldsymbol{\epsilon}^\intercal}{\partial\mathbf{x}}\right)\right|_{\mathbf{y}=\bar{\mathbf{y}}} \in \mathbb{R}^{N_\mathrm{a}N_\mathrm{e}\times N_\mathrm{a}N_\mathrm{e}}, \quad \text{(Layer 4, Eq. S12)}$$

which is always invertible (section S5.3, Eq. S5.3.5). This matrix can be interpreted as measuring *environmental feedback*, which depends on direct mutual environmental dependence, total plasticity of the phenotype, and direct niche construction by the phenotype.

We can also use some of the following previous total-effect matrices to construct the following total-effect matrices involving the geno-envo-phenotype. The block matrix of *total effects of a mutant's phenotype on her geno-envo-phenotype* is

$$\begin{aligned}\left.\frac{\mathrm{d}\mathbf{m}^\intercal}{\mathrm{d}\mathbf{x}}\right|_{\mathbf{y}=\bar{\mathbf{y}}} &\equiv \left.\begin{pmatrix}\frac{\mathrm{d}\mathbf{x}^\intercal}{\mathrm{d}\mathbf{x}} & \frac{\mathrm{d}\mathbf{y}^\intercal}{\mathrm{d}\mathbf{x}} & \frac{\mathrm{d}\boldsymbol{\epsilon}^\intercal}{\mathrm{d}\mathbf{x}}\end{pmatrix}\right|_{\mathbf{y}=\bar{\mathbf{y}}} \\ &= \left.\begin{pmatrix}\frac{\mathrm{d}\mathbf{x}^\intercal}{\mathrm{d}\mathbf{x}} & \mathbf{0} & \frac{\mathrm{d}\boldsymbol{\epsilon}^\intercal}{\mathrm{d}\mathbf{x}}\end{pmatrix}\right|_{\mathbf{y}=\bar{\mathbf{y}}} \quad \text{(Layer 4, Eq. S13)} \\ &\in \mathbb{R}^{N_\mathrm{a}N_\mathrm{p}\times N_\mathrm{a}(N_\mathrm{p}+N_\mathrm{g}+N_\mathrm{e})},\end{aligned}$$

measuring total developmental bias of the geno-envo-phenotype from the phenotype. The block matrix of *total effects of a mutant's genotype on her geno-envo-phenotype* is

$$\begin{aligned}\left.\frac{\mathrm{d}\mathbf{m}^\intercal}{\mathrm{d}\mathbf{y}}\right|_{\mathbf{y}=\bar{\mathbf{y}}} &\equiv \left.\begin{pmatrix}\frac{\mathrm{d}\mathbf{x}^\intercal}{\mathrm{d}\mathbf{y}} & \frac{\mathrm{d}\mathbf{y}^\intercal}{\mathrm{d}\mathbf{y}} & \frac{\mathrm{d}\boldsymbol{\epsilon}^\intercal}{\mathrm{d}\mathbf{y}}\end{pmatrix}\right|_{\mathbf{y}=\bar{\mathbf{y}}} \\ &= \left.\begin{pmatrix}\frac{\mathrm{d}\mathbf{x}^\intercal}{\mathrm{d}\mathbf{y}} & \mathbf{I} & \frac{\mathrm{d}\boldsymbol{\epsilon}^\intercal}{\mathrm{d}\mathbf{y}}\end{pmatrix}\right|_{\mathbf{y}=\bar{\mathbf{y}}} \quad \text{(Layer 4, Eq. S14)} \\ &\in \mathbb{R}^{N_\mathrm{a}N_\mathrm{g}\times N_\mathrm{a}(N_\mathrm{p}+N_\mathrm{g}+N_\mathrm{e})},\end{aligned}$$

measuring total developmental bias of the geno-envo-phenotype from the genotype, and which is singular because it has fewer rows than columns.

The block matrix of *total effects of a mutant's environment on her geno-envo-phenotype* is

$$\begin{aligned}\left.\frac{\mathrm{d}\mathbf{m}^\intercal}{\mathrm{d}\boldsymbol{\epsilon}}\right|_{\mathbf{y}=\bar{\mathbf{y}}} &= \left.\begin{pmatrix}\frac{\mathrm{d}\mathbf{x}^\intercal}{\mathrm{d}\boldsymbol{\epsilon}} & \frac{\mathrm{d}\mathbf{y}^\intercal}{\mathrm{d}\boldsymbol{\epsilon}} & \frac{\mathrm{d}\boldsymbol{\epsilon}^\intercal}{\mathrm{d}\boldsymbol{\epsilon}}\end{pmatrix}\right|_{\mathbf{y}=\bar{\mathbf{y}}} \\ &= \left.\begin{pmatrix}\frac{\mathrm{d}\mathbf{x}^\intercal}{\mathrm{d}\boldsymbol{\epsilon}} & \mathbf{0} & \frac{\mathrm{d}\boldsymbol{\epsilon}^\intercal}{\mathrm{d}\boldsymbol{\epsilon}}\end{pmatrix}\right|_{\mathbf{y}=\bar{\mathbf{y}}} \quad \text{(Layer 4, Eq. S15)} \\ &\in \mathbb{R}^{N_\mathrm{a}N_\mathrm{e}\times N_\mathrm{a}(N_\mathrm{p}+N_\mathrm{g}+N_\mathrm{e})},\end{aligned}$$

measuring total plasticity of the geno-envo-phenotype. The block matrix of *total effects of a mutant's geno-phenotype on her geno-envo-phenotype* is

$$\left.\frac{\mathrm{d}\mathbf{m}^\intercal}{\mathrm{d}\mathbf{z}}\right|_{\mathbf{y}=\bar{\mathbf{y}}} \equiv \left.\begin{pmatrix}\frac{\mathrm{d}\mathbf{m}^\intercal}{\mathrm{d}\mathbf{x}} \\ \frac{\mathrm{d}\mathbf{m}^\intercal}{\mathrm{d}\mathbf{y}}\end{pmatrix}\right|_{\mathbf{y}=\bar{\mathbf{y}}} = \left.\begin{pmatrix}\frac{\mathrm{d}\mathbf{x}^\intercal}{\mathrm{d}\mathbf{x}} & \mathbf{0} & \frac{\mathrm{d}\boldsymbol{\epsilon}^\intercal}{\mathrm{d}\mathbf{x}} \\ \frac{\mathrm{d}\mathbf{x}^\intercal}{\mathrm{d}\mathbf{y}} & \mathbf{I} & \frac{\mathrm{d}\boldsymbol{\epsilon}^\intercal}{\mathrm{d}\mathbf{y}}\end{pmatrix}\right|_{\mathbf{y}=\bar{\mathbf{y}}} \quad \text{(Layer 4, Eq. S16)}$$
$$\in \mathbb{R}^{N_\mathrm{a}(N_\mathrm{p}+N_\mathrm{g})\times N_\mathrm{a}(N_\mathrm{p}+N_\mathrm{g}+N_\mathrm{e})},$$

measuring total developmental bias of the geno-envo-phenotype from the geno-phenotype. The block matrix of *total effects of a mutant's geno-envo-phenotype on her geno-envo-phenotype* is

$$\left.\frac{\mathrm{d}\mathbf{m}^\intercal}{\mathrm{d}\mathbf{m}}\right|_{\mathbf{y}=\bar{\mathbf{y}}} = \left.\begin{pmatrix}\frac{\mathrm{d}\mathbf{m}^\intercal}{\mathrm{d}\mathbf{x}} \\ \frac{\mathrm{d}\mathbf{m}^\intercal}{\mathrm{d}\mathbf{y}} \\ \frac{\mathrm{d}\mathbf{m}^\intercal}{\mathrm{d}\boldsymbol{\epsilon}}\end{pmatrix}\right|_{\mathbf{y}=\bar{\mathbf{y}}} = \left.\begin{pmatrix}\frac{\mathrm{d}\mathbf{x}^\intercal}{\mathrm{d}\mathbf{x}} & \mathbf{0} & \frac{\mathrm{d}\boldsymbol{\epsilon}^\intercal}{\mathrm{d}\mathbf{x}} \\ \frac{\mathrm{d}\mathbf{x}^\intercal}{\mathrm{d}\mathbf{y}} & \mathbf{I} & \frac{\mathrm{d}\boldsymbol{\epsilon}^\intercal}{\mathrm{d}\mathbf{y}} \\ \frac{\mathrm{d}\mathbf{x}^\intercal}{\mathrm{d}\boldsymbol{\epsilon}} & \mathbf{0} & \frac{\mathrm{d}\boldsymbol{\epsilon}^\intercal}{\mathrm{d}\boldsymbol{\epsilon}}\end{pmatrix}\right|_{\mathbf{y}=\bar{\mathbf{y}}} \quad \text{(Layer 4, Eq. S17)}$$
$$\in \mathbb{R}^{N_\mathrm{a}(N_\mathrm{p}+N_\mathrm{g}+N_\mathrm{e})\times N_\mathrm{a}(N_\mathrm{p}+N_\mathrm{g}+N_\mathrm{e})},$$

measuring developmental feedback of the geno-envo-phenotype, and which we show is invertible (section S5.5). Obtaining a compact form for $\mathrm{d}\mathbf{m}^\intercal/\mathrm{d}\mathbf{m}|_{\mathbf{y}=\bar{\mathbf{y}}}$ analogous to Layer 4, Eq. S8 seemingly needs $(\mathrm{d}\boldsymbol{\epsilon}^\intercal/\mathrm{d}\boldsymbol{\epsilon}|_{\mathbf{y}=\bar{\mathbf{y}}})^{-1}$ which appears to yield relatively complex expressions so we leave this for future analysis.

We will see that the evolutionary dynamics of the phenotype depends on a matrix measuring "joint" total developmental bias of the phenotype. This matrix is the transpose of the matrix of *total social effects of a focal individual's genotype or phenotype on hers and her partners' phenotypes*

$$\left.\frac{\mathrm{d}(\mathbf{x}+\check{\mathbf{x}})}{\mathrm{d}\boldsymbol{\zeta}^\intercal}\right|_{\mathbf{y}=\bar{\mathbf{y}}} = \left.\left(\frac{\mathrm{d}\mathbf{x}}{\mathrm{d}\boldsymbol{\zeta}^\intercal}+\frac{\mathrm{d}\mathbf{x}}{\mathrm{d}\bar{\boldsymbol{\zeta}}^\intercal}\right)\right|_{\mathbf{y}=\bar{\mathbf{y}}}, \quad \text{(Layer 4, Eq. S18)}$$

for $\boldsymbol{\zeta} \in \{\mathbf{x},\mathbf{y}\}$ where we denote by $\check{\mathbf{x}}$ the phenotype that a resident develops in the context of mutants (a donor perspective for the mutant). Thus, this matrix can be interpreted as measuring joint total developmental bias of the phenotype. Note that the second term on the right-hand side of Layer 4, Eq. S18 is the total effects of social partners' phenotype or genotype on a focal mutant (a recipient perspective for the mutant). Thus, the joint total developmental bias of the phenotype as described by Layer 4, Eq. S18 can be equivalently interpreted either from a donor or a recipient perspective.

Having written expressions for the above total-effect matrices, we can now write the total selection gradients, which measure total directional selection, that is, directional selection considering all the pathways in which a trait can affect fitness in Fig. 1 (see also Morrissey 2014). This contrasts with Lande's (1979) selection gradient, which corresponds to the direct selection gradient measuring the direct effect of a variable on fitness in Fig. 1. In sections S5.1-S5.5, we show that the total selection gradient of vector $\boldsymbol{\zeta} \in \{\mathbf{x},\mathbf{y},\mathbf{z},\boldsymbol{\epsilon},\mathbf{m}\}$ is

$$\left.\frac{\mathrm{d}w}{\mathrm{d}\boldsymbol{\zeta}}\right|_{\mathbf{y}=\bar{\mathbf{y}}} = \left.\left(\frac{\mathrm{d}\mathbf{m}^\intercal}{\mathrm{d}\boldsymbol{\zeta}}\frac{\partial w}{\partial\mathbf{m}}\right)\right|_{\mathbf{y}=\bar{\mathbf{y}}}, \quad \text{(Layer 4, Eq. S19)}$$

which has the form of the chain rule in matrix calculus notation. Hence, the total selection gradient of $\boldsymbol{\zeta}$ depends on the total effects of $\boldsymbol{\zeta}$ on the geno-envo-phenotype and direct directional selection on the geno-envo-phenotype. Consequently, the total directional selection on $\boldsymbol{\zeta}$ is the directional selection on the geno-envo-phenotype transformed by the total effects of $\boldsymbol{\zeta}$ on the geno-envo-phenotype considering the downstream developmental effects. Layer 4, Eq. S19 has the same form of previous expressions by Caswell (e.g., Caswell, 1982, Eq. 4 and Caswell, 2001, Eq. 9.38), except that it is in terms of traits (including traits constructed according to developmental constraints) rather than vital rates (i.e., Caswell's equations have the entries of the Leslie matrix in Eq. S2.1.7 in the place of $\mathbf{m}$). Layer 4, Eq. S19 also recovers the form of Morrissey's (2014) extended selection gradient. Total selection gradients take the following particular forms.

The total selection gradient of the phenotype is

$$\begin{aligned}\left.\frac{\mathrm{d}w}{\mathrm{d}\mathbf{x}}\right|_{\mathbf{y}=\bar{\mathbf{y}}} &= \left.\left(\frac{\mathrm{d}\mathbf{x}^\intercal}{\mathrm{d}\mathbf{x}}\frac{\partial w}{\partial \mathbf{x}} + \frac{\mathrm{d}\boldsymbol{\epsilon}^\intercal}{\mathrm{d}\mathbf{x}}\frac{\partial w}{\partial \boldsymbol{\epsilon}}\right)\right|_{\mathbf{y}=\bar{\mathbf{y}}} \qquad \text{(Layer 4, Eq. S20)} \\ &= \left.\left(\frac{\mathrm{d}\mathbf{x}^\intercal}{\mathrm{d}\mathbf{x}}\frac{\delta w}{\delta \mathbf{x}}\right)\right|_{\mathbf{y}=\bar{\mathbf{y}}} \\ &= \left.\left(\frac{\mathrm{d}\mathbf{z}^\intercal}{\mathrm{d}\mathbf{x}}\frac{\delta w}{\delta \mathbf{z}}\right)\right|_{\mathbf{y}=\bar{\mathbf{y}}} \\ &= \left.\left(\frac{\mathrm{d}\mathbf{m}^\intercal}{\mathrm{d}\mathbf{x}}\frac{\partial w}{\partial \mathbf{m}}\right)\right|_{\mathbf{y}=\bar{\mathbf{y}}}.\end{aligned}$$

This gradient depends on direct directional selection on the phenotype and direct directional selection on the environment (Layer 2, Eq. S1). It also depends on developmental feedback of the phenotype (Layer 4, Eq. S1) and total niche construction by the phenotype, which also depends on developmental feedback of the phenotype (Layer 4, Eq. S9). Consequently, the total selection gradient of the phenotype can be interpreted as measuring total (directional) phenotypic selection in the fitness landscape modified by developmental feedback of the phenotype and by the interaction of total niche construction and environmental sensitivity of selection. Additionally, the total selection gradient of the phenotype at an admissible stable evolutionary equilibrium equals generation time $T$ times the vector of costate variables, which are key variables used for solving optimal control problems (section S4, Eq. S4.3; see also Metz *et al.* 2016). In optimal control problems with continuous time, costate variables must be found by solving differential equations with boundary conditions at the terminal time while simultaneously solving differential equations describing state dynamics with boundary conditions at the initial time (Bryson, Jr. and Ho, 1975; Sydsæter *et al.*, 2008). This poses a two-point boundary value problem, which is typically challenging to solve. Our discrete-age approach allows us to obtain closed-form formulas for the total selection gradient of the phenotype (Layer 4, Eq. S20), thus providing closed-form formulas for costate variables and avoiding having to solve a two-point boundary value problem.

The total selection gradient of the genotype is

$$\begin{aligned}\left.\frac{\mathrm{d}w}{\mathrm{d}\mathbf{y}}\right|_{\mathbf{y}=\bar{\mathbf{y}}} &= \left.\left(\frac{\mathrm{d}\mathbf{x}^\intercal}{\mathrm{d}\mathbf{y}}\frac{\partial w}{\partial \mathbf{x}} + \frac{\partial w}{\partial \mathbf{y}} + \frac{\mathrm{d}\boldsymbol{\epsilon}^\intercal}{\mathrm{d}\mathbf{y}}\frac{\partial w}{\partial \boldsymbol{\epsilon}}\right)\right|_{\mathbf{y}=\bar{\mathbf{y}}} \\ &\qquad \text{(Layer 4, Eq. S21)} \\ &= \left.\left(\frac{\mathrm{d}\mathbf{x}^\intercal}{\mathrm{d}\mathbf{y}}\frac{\delta w}{\delta \mathbf{x}} + \frac{\delta w}{\delta \mathbf{y}}\right)\right|_{\mathbf{y}=\bar{\mathbf{y}}} \\ &= \left.\left(\frac{\mathrm{d}\mathbf{z}^\intercal}{\mathrm{d}\mathbf{y}}\frac{\delta w}{\delta \mathbf{z}}\right)\right|_{\mathbf{y}=\bar{\mathbf{y}}} \\ &= \left.\left(\frac{\mathrm{d}\mathbf{m}^\intercal}{\mathrm{d}\mathbf{y}}\frac{\partial w}{\partial \mathbf{m}}\right)\right|_{\mathbf{y}=\bar{\mathbf{y}}} \\ &= \left.\left(\frac{\delta\mathbf{x}^\intercal}{\delta\mathbf{y}}\frac{\mathrm{d} w}{\mathrm{d} \mathbf{x}} + \frac{\delta w}{\delta \mathbf{y}}\right)\right|_{\mathbf{y}=\bar{\mathbf{y}}}.\end{aligned}$$

This gradient not only depends on direct directional selection on the phenotype and the environment, but also on direct directional selection on the genotype (Layer 2, Eq. S1). It also depends on the mechanistic analogue of Fisher's (1918) additive effects of allelic substitution or of Wagner's (1984, 1989) developmental matrix (Layer 4, Eq. S2) and on total niche construction by the genotype, which also depends on the developmental matrix (Layer 4, Eq. S10). The fifth line of Layer 4, Eq. S21 has the form of previous expressions for the total selection gradient of controls in continuous age in terms of partial derivatives of the Hamiltonian involving costate variables for which closed-form formulas have been lacking (e.g., Day and Taylor 1997, Eq. 4, Day and Taylor 2000, Eq. 6, and Avila *et al.* 2021, Eq. 23; see also our Eq. S4.4). Our discrete-age approach allowed us to obtain closed-form formulas for the total selection gradient of states (Layer 4, Eq. S20), thus providing closed-form formulas for the total selection gradient of controls.

To derive equations describing the evolutionary dynamics of the geno-envo-phenotype, we make use of the total selection gradient of the environment, although such gradient is not necessary to obtain equations describing the evolutionary dynamics of the geno-phenotype. The total selection gradient of the environment is

$$\begin{aligned}\left.\frac{\mathrm{d}w}{\mathrm{d}\boldsymbol{\epsilon}}\right|_{\mathbf{y}=\bar{\mathbf{y}}} &= \left.\left(\frac{\mathrm{d}\mathbf{x}^\intercal}{\mathrm{d}\boldsymbol{\epsilon}}\frac{\partial w}{\partial \mathbf{x}} + \frac{\mathrm{d}\boldsymbol{\epsilon}^\intercal}{\mathrm{d}\boldsymbol{\epsilon}}\frac{\partial w}{\partial \boldsymbol{\epsilon}}\right)\right|_{\mathbf{y}=\bar{\mathbf{y}}} \qquad \text{(Layer 4, Eq. S22)} \\ &= \left.\left(\frac{\mathrm{d}\mathbf{x}^\intercal}{\mathrm{d}\boldsymbol{\epsilon}}\frac{\delta w}{\delta \mathbf{x}} + \frac{\delta w}{\delta \boldsymbol{\epsilon}}\right)\right|_{\mathbf{y}=\bar{\mathbf{y}}} \\ &= \left.\left(\frac{\mathrm{d}\mathbf{m}^\intercal}{\mathrm{d}\boldsymbol{\epsilon}}\frac{\partial w}{\partial \mathbf{m}}\right)\right|_{\mathbf{y}=\bar{\mathbf{y}}} \\ &= \left.\left(\frac{\delta\mathbf{x}^\intercal}{\delta\boldsymbol{\epsilon}}\frac{\mathrm{d} w}{\mathrm{d} \mathbf{x}} + \frac{\delta w}{\delta \boldsymbol{\epsilon}}\right)\right|_{\mathbf{y}=\bar{\mathbf{y}}}.\end{aligned}$$

This gradient depends on total plasticity of the phenotype and on environmental feedback, which in turn depends on total plasticity of the phenotype and niche construction by the phenotype (Layer 4, Eq. S12). Consequently, the total selection gradient of the environment can be understood as measuring total (directional) environmental selection in a fitness landscape modified by environmental feedback and by the interaction of total plasticity of the phenotype and direct directional selection on the phenotype.

We can combine the expressions for the total selection gradients above to obtain the total selection gradient of the geno-phenotype and the geno-envo-phenotype. The total selection gradient of the geno-phenotype is

$$\begin{aligned}\left.\frac{\mathrm{d}w}{\mathrm{d}\mathbf{z}}\right|_{\mathbf{y}=\bar{\mathbf{y}}} &= \left.\left(\frac{\mathrm{d}\mathbf{z}^\intercal}{\mathrm{d}\mathbf{z}}\frac{\partial w}{\partial \mathbf{z}} + \frac{\mathrm{d}\boldsymbol{\epsilon}^\intercal}{\mathrm{d}\mathbf{z}}\frac{\partial w}{\partial \boldsymbol{\epsilon}}\right)\right|_{\mathbf{y}=\bar{\mathbf{y}}} \\ &= \left.\left(\frac{\mathrm{d}\mathbf{z}^\intercal}{\mathrm{d}\mathbf{z}}\frac{\delta w}{\delta \mathbf{z}}\right)\right|_{\mathbf{y}=\bar{\mathbf{y}}} \\ &= \left.\left(\frac{\mathrm{d}\mathbf{m}^\intercal}{\mathrm{d}\mathbf{z}}\frac{\partial w}{\partial \mathbf{m}}\right)\right|_{\mathbf{y}=\bar{\mathbf{y}}}.\end{aligned} \quad \text{(Layer 4, Eq. S23)}$$

Thus, the total selection gradient of the geno-phenotype can be interpreted as measuring total (directional) geno-phenotypic selection in a fitness landscape modified by developmental feedback of the geno-phenotype and by the interaction of total niche construction by the geno-phenotype and environmental sensitivity of selection. In turn, the total selection gradient of the geno-envo-phenotype is

$$\left.\frac{\mathrm{d}w}{\mathrm{d}\mathbf{m}}\right|_{\mathbf{y}=\bar{\mathbf{y}}} = \left.\left(\frac{\mathrm{d}\mathbf{m}^\intercal}{\mathrm{d}\mathbf{m}}\frac{\partial w}{\partial \mathbf{m}}\right)\right|_{\mathbf{y}=\bar{\mathbf{y}}}, \quad \text{(Layer 4, Eq. S24)}$$

which can be interpreted as measuring total (directional) geno-envo-phenotypic selection in a fitness landscape modified by developmental feedback of the geno-envo-phenotype.

#### S3.5 Layer 5: stabilized effects

We now move on to write the equations for the next layer of the evo-devo process, that of (socio-devo) stabilized-effect matrices. Stabilized-effect matrices measure the total effects of a variable on another one considering downstream developmental effects, after the effects of social development have stabilized in the population. Stabilized-effect matrices arise in the derivation of the evolutionary dynamics of the phenotype and environment as a result of social development. If development is not social (i.e., $\mathrm{d}\mathbf{x}^\intercal/\mathrm{d}\bar{\mathbf{z}}|_{\mathbf{y}=\bar{\mathbf{y}}} = \mathbf{0}$), then all stabilized-effect matrices ($\mathrm{s}\boldsymbol{\zeta}^\intercal/\mathrm{s}\boldsymbol{\xi}|_{\mathbf{y}=\bar{\mathbf{y}}}$) reduce to the corresponding total-effect matrices ($\mathrm{d}\boldsymbol{\zeta}^\intercal/\mathrm{d}\boldsymbol{\xi}|_{\mathbf{y}=\bar{\mathbf{y}}}$), except one ($\mathrm{s}\mathbf{x}^\intercal/\mathrm{s}\bar{\mathbf{x}}|_{\mathbf{y}=\bar{\mathbf{y}}}$) that reduces to the identity matrix.

The stabilized effects of social partners' phenotypes on a focal individual's phenotype measure *social feedback*. This is given by the transpose of the matrix of *stabilized effects of social partners' phenotypes on a focal individual's phenotype*

$$\begin{aligned}\left.\frac{\mathrm{s}\mathbf{x}}{\mathrm{s}\bar{\mathbf{x}}^\intercal}\right|_{\mathbf{y}=\bar{\mathbf{y}}} &= \left(\mathbf{I} - \left.\frac{\mathrm{d}\check{\mathbf{x}}}{\mathrm{d}\mathbf{x}^\intercal}\right|_{\mathbf{y}=\bar{\mathbf{y}}}\right)^{-1} = \left(\mathbf{I} - \left.\frac{\mathrm{d}\mathbf{x}}{\mathrm{d}\bar{\mathbf{x}}^\intercal}\right|_{\mathbf{y}=\bar{\mathbf{y}}}\right)^{-1} \\ &= \left.\sum_{\theta=1}^{\infty}\left(\frac{\mathrm{d}\mathbf{x}}{\mathrm{d}\bar{\mathbf{x}}^\intercal}\right)^{\theta-1}\right|_{\mathbf{y}=\bar{\mathbf{y}}} \in \mathbb{R}^{N_\mathrm{a}N_\mathrm{p}\times N_\mathrm{a}N_\mathrm{p}},\end{aligned} \quad \text{(Layer 5, Eq. S1)}$$

where the last equality follows by the geometric series of matrices. The matrix $\mathrm{s}\mathbf{x}/\mathrm{s}\bar{\mathbf{x}}^\intercal|_{\mathbf{y}=\bar{\mathbf{y}}}$ is invertible by our assumption that all the eigenvalues of $\mathrm{d}\mathbf{x}/\mathrm{d}\bar{\mathbf{x}}^\intercal|_{\mathbf{y}=\bar{\mathbf{y}}}$ have absolute value strictly less than one, to guarantee that the resident is socio-devo stable. The matrix $\mathrm{s}\mathbf{x}/\mathrm{s}\bar{\mathbf{x}}^\intercal|_{\mathbf{y}=\bar{\mathbf{y}}}$ can be interpreted as the total effects of social partners' phenotypes on a focal individual's phenotype after socio-devo stabilization (Eq. S2.1.1); or vice versa, of a focal individual's phenotype on social partners' phenotypes. Thus, the matrix $\mathrm{s}\mathbf{x}/\mathrm{s}\bar{\mathbf{x}}^\intercal|_{\mathbf{y}=\bar{\mathbf{y}}}$ describes social feedback arising from social development. This matrix corresponds to an analogous matrix found in the indirect genetic effects literature (Moore *et al.*, 1997, Eq. 19b and subsequent text). If development is not social from the phenotype (i.e., $\mathrm{d}\mathbf{x}^\intercal/\mathrm{d}\bar{\mathbf{x}}|_{\mathbf{y}=\bar{\mathbf{y}}} = \mathbf{0}$), then the matrix $\mathrm{s}\mathbf{x}/\mathrm{s}\bar{\mathbf{x}}^\intercal|_{\mathbf{y}=\bar{\mathbf{y}}}$ is the identity matrix. This is the only stabilized-effect matrix that does not reduce to the corresponding total-effect matrix when development is not social.

The stabilized effects of a focal individual's phenotype or genotype on her phenotype measure stabilized developmental bias. We define the transpose of the matrix of *stabilized effects of a focal individual's phenotype or genotype on her phenotype* as

$$\left.\frac{\mathrm{s}\mathbf{x}}{\mathrm{s}\boldsymbol{\zeta}^\intercal}\right|_{\mathbf{y}=\bar{\mathbf{y}}} = \left.\left(\frac{\mathrm{s}\mathbf{x}}{\mathrm{s}\bar{\mathbf{x}}^\intercal}\frac{\mathrm{d}(\mathbf{x}+\check{\mathbf{x}})}{\mathrm{d}\boldsymbol{\zeta}^\intercal}\right)\right|_{\mathbf{y}=\bar{\mathbf{y}}}, \quad \text{(Layer 5, Eq. S2a)}$$

for $\boldsymbol{\zeta} \in \{\mathbf{x}, \mathbf{y}\}$. This matrix can be interpreted as measuring stabilized developmental bias of the phenotype from $\boldsymbol{\zeta}$, where a focal individual's genotype or phenotype first affects the development of her own and social partners' phenotype which then feeds back to affect the individual's phenotype. Stabilized developmental bias is "joint" in that it includes both the effects of the focal individual on herself and on social partners. If development is not social (i.e., $\mathrm{d}\mathbf{x}^\intercal/\mathrm{d}\bar{\mathbf{z}}|_{\mathbf{y}=\bar{\mathbf{y}}} = \mathbf{0}$), then a stabilized developmental bias matrix ($\mathrm{s}\mathbf{x}/\mathrm{s}\boldsymbol{\zeta}^\intercal|_{\mathbf{y}=\bar{\mathbf{y}}}$) reduces to the corresponding total developmental bias matrix ($\mathrm{d}\mathbf{x}/\mathrm{d}\boldsymbol{\zeta}^\intercal|_{\mathbf{y}=\bar{\mathbf{y}}}$).

The stabilized effects of the environment on the phenotype measure stabilized plasticity. The transpose of the matrix of *stabilized effects of a focal individual's environment on the phenotype* is

$$\left.\frac{\mathrm{s}\mathbf{x}}{\mathrm{s}\boldsymbol{\epsilon}^\intercal}\right|_{\mathbf{y}=\bar{\mathbf{y}}} = \left.\left(\frac{\mathrm{s}\mathbf{x}}{\mathrm{s}\bar{\mathbf{x}}^\intercal}\frac{\mathrm{d}\mathbf{x}}{\mathrm{d}\boldsymbol{\epsilon}^\intercal}\right)\right|_{\mathbf{y}=\bar{\mathbf{y}}} \in \mathbb{R}^{N_\mathrm{a}N_\mathrm{p}\times N_\mathrm{a}N_\mathrm{e}}. \quad \text{(Layer 5, Eq. S2b)}$$

This matrix can be interpreted as measuring stabilized plasticity of the phenotype, where the environment first causes total plasticity in a focal individual and then the focal individual causes stabilized social effects on social partners. Stabilized plasticity does not depend on the joint effects of the environment. If development is not social (i.e., $\mathrm{d}\mathbf{x}^\intercal/\mathrm{d}\bar{\mathbf{z}}|_{\mathbf{y}=\bar{\mathbf{y}}} = \mathbf{0}$), then stabilized plasticity reduces to total plasticity.

The stabilized effects on the genotype are simple since genotypic traits are developmentally independent by assumption. The transpose of the matrix of *stabilized effects of a focal individual's phenotype or environment on the genotype* is

$$\left.\frac{\mathrm{s}\mathbf{y}}{\mathrm{s}\boldsymbol{\zeta}^\intercal}\right|_{\mathbf{y}=\bar{\mathbf{y}}} = \left.\frac{\mathrm{d}\mathbf{y}}{\mathrm{d}\boldsymbol{\zeta}^\intercal}\right|_{\mathbf{y}=\bar{\mathbf{y}}} = \mathbf{0}, \quad \text{(Layer 5, Eq. S3a)}$$

for $\boldsymbol{\zeta} \in \{\mathbf{x}, \boldsymbol{\epsilon}\}$. The transpose of the matrix of *stabilized ef-*

*fects of a focal individual's genotype on the genotype* is

$$\left.\frac{\mathrm{s}\mathbf{y}}{\mathrm{s}\mathbf{y}^\intercal}\right|_{\mathbf{y}=\bar{\mathbf{y}}} = \left.\frac{\mathrm{d}\mathbf{y}}{\mathrm{d}\mathbf{y}^\intercal}\right|_{\mathbf{y}=\bar{\mathbf{y}}} = \mathbf{I} \quad \in \mathbb{R}^{N_\mathrm{a}N_\mathrm{g}\times N_\mathrm{a}N_\mathrm{g}}. \tag{Layer 5, Eq. S3b}$$

We can use some of the previous stabilized-effect matrices to construct the following stabilized-effect matrices involving the geno-phenotype. The transpose of the matrix of *stabilized effects of a focal individual's genotype on the geno-phenotype* is

$$\left.\frac{\mathrm{s}\mathbf{z}}{\mathrm{s}\mathbf{y}^\intercal}\right|_{\mathbf{y}=\bar{\mathbf{y}}} \equiv \left.\left(\frac{\mathrm{s}\mathbf{x}}{\mathrm{s}\mathbf{y}^\intercal}; \frac{\mathrm{s}\mathbf{y}}{\mathrm{s}\mathbf{y}^\intercal}\right)\right|_{\mathbf{y}=\bar{\mathbf{y}}} \in \mathbb{R}^{N_\mathrm{a}(N_\mathrm{p}+N_\mathrm{g})\times N_\mathrm{a}N_\mathrm{g}}, \tag{Layer 5, Eq. S4a}$$

measuring stabilized developmental bias of the geno-phenotype from the genotype. The transpose of the matrix of *stabilized effects of a focal individual's environment on the geno-phenotype* is

$$\left.\frac{\mathrm{s}\mathbf{z}}{\mathrm{s}\boldsymbol{\epsilon}^\intercal}\right|_{\mathbf{y}=\bar{\mathbf{y}}} \equiv \left.\left(\frac{\mathrm{s}\mathbf{x}}{\mathrm{s}\boldsymbol{\epsilon}^\intercal}; \frac{\mathrm{s}\mathbf{y}}{\mathrm{s}\boldsymbol{\epsilon}^\intercal}\right)\right|_{\mathbf{y}=\bar{\mathbf{y}}} \in \mathbb{R}^{N_\mathrm{a}(N_\mathrm{p}+N_\mathrm{g})\times N_\mathrm{a}N_\mathrm{e}}, \tag{Layer 5, Eq. S4b}$$

measuring stabilized plasticity of the geno-phenotype. The transpose of the matrix of *stabilized effects of a focal individual's geno-phenotype on the geno-phenotype* is

$$\left.\frac{\mathrm{s}\mathbf{z}}{\mathrm{s}\mathbf{z}^\intercal}\right|_{\mathbf{y}=\bar{\mathbf{y}}} \equiv \left.\begin{pmatrix} \frac{\mathrm{s}\mathbf{x}}{\mathrm{s}\mathbf{x}^\intercal} & \frac{\mathrm{s}\mathbf{x}}{\mathrm{s}\mathbf{y}^\intercal} \\ \frac{\mathrm{s}\mathbf{y}}{\mathrm{s}\mathbf{x}^\intercal} & \frac{\mathrm{s}\mathbf{y}}{\mathrm{s}\mathbf{y}^\intercal} \end{pmatrix}\right|_{\mathbf{y}=\bar{\mathbf{y}}} = \left.\begin{pmatrix} \frac{\mathrm{s}\mathbf{x}}{\mathrm{s}\mathbf{x}^\intercal} & \frac{\mathrm{s}\mathbf{x}}{\mathrm{s}\mathbf{y}^\intercal} \\ \mathbf{0} & \mathbf{I} \end{pmatrix}\right|_{\mathbf{y}=\bar{\mathbf{y}}} \tag{Layer 5, Eq. S5}$$

$$\in \mathbb{R}^{N_\mathrm{a}(N_\mathrm{p}+N_\mathrm{g})\times N_\mathrm{a}(N_\mathrm{p}+N_\mathrm{g})},$$

measuring stabilized developmental feedback of the geno-phenotype.

The stabilized effects of the phenotype or genotype on the environment measure stabilized niche construction. Although the matrix

$$\left.\frac{\mathrm{s}\boldsymbol{\epsilon}}{\mathrm{s}\mathbf{x}^\intercal}\right|_{\mathbf{y}=\bar{\mathbf{y}}}$$

appears in some of the matrices we construct, it is irrelevant as it disappears in the matrix products we encounter. The following matrix does not disappear. The transpose of the matrix of *stabilized effects of a focal individual's genotype on the environment* is

$$\left.\frac{\mathrm{s}\boldsymbol{\epsilon}}{\mathrm{s}\mathbf{y}^\intercal}\right|_{\mathbf{y}=\bar{\mathbf{y}}} = \left.\left(\frac{\partial(\boldsymbol{\epsilon}+\check{\boldsymbol{\epsilon}})}{\partial\mathbf{z}^\intercal}\frac{\mathrm{s}\mathbf{z}}{\mathrm{s}\mathbf{y}^\intercal}\right)\right|_{\mathbf{y}=\bar{\mathbf{y}}} \in \mathbb{R}^{N_\mathrm{a}N_\mathrm{e}\times N_\mathrm{a}N_\mathrm{g}}, \tag{Layer 5, Eq. S6a}$$

which is formed by stabilized developmental bias of the geno-phenotype from genotype followed by joint direct niche construction by the geno-phenotype. This matrix can be interpreted as measuring stabilized niche construction by the genotype. If development is not social (i.e., $\mathrm{d}\mathbf{x}^\intercal/\mathrm{d}\bar{\mathbf{z}}|_{\mathbf{y}=\bar{\mathbf{y}}} = \mathbf{0}$), then stabilized niche construction by the genotype reduces to total niche construction by the genotype (see Layer 4, Eq. S10 and Layer 2, Eq. S10).

The stabilized effects of the environment on itself measure stabilized environmental feedback. The transpose of the matrix of *stabilized effects of a focal individual's environment on the environment* is

$$\left.\frac{\mathrm{s}\boldsymbol{\epsilon}}{\mathrm{s}\boldsymbol{\epsilon}^\intercal}\right|_{\mathbf{y}=\bar{\mathbf{y}}} = \left.\left(\frac{\partial(\boldsymbol{\epsilon}+\check{\boldsymbol{\epsilon}})}{\partial\mathbf{z}^\intercal}\frac{\mathrm{s}\mathbf{z}}{\mathrm{s}\boldsymbol{\epsilon}^\intercal} + \frac{\partial\boldsymbol{\epsilon}}{\partial\boldsymbol{\epsilon}^\intercal}\right)\right|_{\mathbf{y}=\bar{\mathbf{y}}} \in \mathbb{R}^{N_\mathrm{a}N_\mathrm{e}\times N_\mathrm{a}N_\mathrm{e}}, \tag{Layer 5, Eq. S6b}$$

which depends on stabilized plasticity of the geno-phenotype, joint direct niche construction by the geno-phenotype, and direct mutual environmental dependence.

We can also use some of the following previous stabilized-effect matrices to construct the following stabilized-effect matrices comprising the geno-envo-phenotype. The transpose of the matrix of *stabilized effects of a focal individual's genotype on the geno-envo-phenotype* is

$$\left.\frac{\mathrm{s}\mathbf{m}}{\mathrm{s}\mathbf{y}^\intercal}\right|_{\mathbf{y}=\bar{\mathbf{y}}} \equiv \left.\left(\frac{\mathrm{s}\mathbf{x}}{\mathrm{s}\mathbf{y}^\intercal}; \frac{\mathrm{s}\mathbf{y}}{\mathrm{s}\mathbf{y}^\intercal}; \frac{\mathrm{s}\boldsymbol{\epsilon}}{\mathrm{s}\mathbf{y}^\intercal}\right)\right|_{\mathbf{y}=\bar{\mathbf{y}}} \tag{Layer 5, Eq. S7a}$$

$$\in \mathbb{R}^{N_\mathrm{a}(N_\mathrm{p}+N_\mathrm{g}+N_\mathrm{e})\times N_\mathrm{a}N_\mathrm{g}},$$

measuring stabilized developmental bias of the geno-envo-phenotype from the genotype. The transpose of the matrix of *stabilized effects of a focal individual's environment on the geno-envo-phenotype* is

$$\left.\frac{\mathrm{s}\mathbf{m}}{\mathrm{s}\boldsymbol{\epsilon}^\intercal}\right|_{\mathbf{y}=\bar{\mathbf{y}}} \equiv \left.\left(\frac{\mathrm{s}\mathbf{x}}{\mathrm{s}\boldsymbol{\epsilon}^\intercal}; \frac{\mathrm{s}\mathbf{y}}{\mathrm{s}\boldsymbol{\epsilon}^\intercal}; \frac{\mathrm{s}\boldsymbol{\epsilon}}{\mathrm{s}\boldsymbol{\epsilon}^\intercal}\right)\right|_{\mathbf{y}=\bar{\mathbf{y}}} \tag{Layer 5, Eq. S7b}$$

$$\in \mathbb{R}^{N_\mathrm{a}(N_\mathrm{p}+N_\mathrm{g}+N_\mathrm{e})\times N_\mathrm{a}N_\mathrm{e}},$$

measuring stabilized plasticity of the geno-envo-phenotype. Finally, the transpose of the matrix of *stabilized effects of a focal individual's geno-envo-phenotype on the geno-envo-phenotype* is

$$\left.\frac{\mathrm{s}\mathbf{m}}{\mathrm{s}\mathbf{m}^\intercal}\right|_{\mathbf{y}=\bar{\mathbf{y}}} \equiv \left.\begin{pmatrix} \frac{\mathrm{s}\mathbf{x}}{\mathrm{s}\mathbf{x}^\intercal} & \frac{\mathrm{s}\mathbf{x}}{\mathrm{s}\mathbf{y}^\intercal} & \frac{\mathrm{s}\mathbf{x}}{\mathrm{s}\boldsymbol{\epsilon}^\intercal} \\ \frac{\mathrm{s}\mathbf{y}}{\mathrm{s}\mathbf{x}^\intercal} & \frac{\mathrm{s}\mathbf{y}}{\mathrm{s}\mathbf{y}^\intercal} & \frac{\mathrm{s}\mathbf{y}}{\mathrm{s}\boldsymbol{\epsilon}^\intercal} \\ \frac{\mathrm{s}\boldsymbol{\epsilon}}{\mathrm{s}\mathbf{x}^\intercal} & \frac{\mathrm{s}\boldsymbol{\epsilon}}{\mathrm{s}\mathbf{y}^\intercal} & \frac{\mathrm{s}\boldsymbol{\epsilon}}{\mathrm{s}\boldsymbol{\epsilon}^\intercal} \end{pmatrix}\right|_{\mathbf{y}=\bar{\mathbf{y}}}$$

$$= \left.\begin{pmatrix} \frac{\mathrm{s}\mathbf{x}}{\mathrm{s}\mathbf{x}^\intercal} & \frac{\mathrm{s}\mathbf{x}}{\mathrm{s}\mathbf{y}^\intercal} & \frac{\mathrm{s}\mathbf{x}}{\mathrm{s}\boldsymbol{\epsilon}^\intercal} \\ \mathbf{0} & \mathbf{I} & \mathbf{0} \\ \frac{\mathrm{s}\boldsymbol{\epsilon}}{\mathrm{s}\mathbf{x}^\intercal} & \frac{\mathrm{s}\boldsymbol{\epsilon}}{\mathrm{s}\mathbf{y}^\intercal} & \frac{\mathrm{s}\boldsymbol{\epsilon}}{\mathrm{s}\boldsymbol{\epsilon}^\intercal} \end{pmatrix}\right|_{\mathbf{y}=\bar{\mathbf{y}}} \tag{Layer 5, Eq. S8}$$

$$\in \mathbb{R}^{N_\mathrm{a}(N_\mathrm{p}+N_\mathrm{g}+N_\mathrm{e})\times N_\mathrm{a}(N_\mathrm{p}+N_\mathrm{g}+N_\mathrm{e})},$$

measuring stabilized developmental feedback of the geno-envo-phenotype.

### S4 Connection to dynamic optimization

In this section we connect evo-devo dynamics to dynamic optimization. Specifically, we first show that an admissible locally stable equilibrium of the evolutionary dynamics of genotypic traits that modulate the developmental dynamics of phenotypic traits locally solves a general life history problem. Then, we show that such locally stable equilibrium satisfies the key elements of Pontryagin's maximum principle in discrete time, which is a theorem that specifies necessary conditions for control variables to solve an optimal control problem.

Life-history models often consider genetically controlled traits (controls) that depend on an underlying variable (e.g., age) together with traits (states) constructed via dynamic (e.g., developmental) constraints over the underlying variable. When such a model is simple enough, analytical solution (i.e., identification of evolutionarily stable strategies) is possible using optimal control or dynamic programming methods (Sydsæter *et al.*, 2008). A key tool from optimal control theory that enables finding such analytical solutions (i.e., optimal controls) is Pontryagin's maximum principle. The maximum principle is a theorem that essentially transforms the dynamic optimization problem into a simpler problem of maximizing a function called the Hamiltonian, which depends on control, state, and costate (or adjoint) variables. The problem is then to maximize the Hamiltonian with respect to the controls, while state and costate variables can be found from associated dynamic equations. We now show that our results imply the key elements of Pontryagin's maximum principle for a standard life-history problem.

First, we state the optimization problem and show that it is locally solved by the evo-devo dynamics. Let $\mathbf{y}$ and $\mathbf{x}$ respectively denote the control and state variables over age, and assume that there are no environmental traits. Let survivorship be a state variable, denoted by $x_{\ell a} = \ell_a$, so it satisfies the developmental constraint $x_{\ell,a+1} = g_{\ell a}(\mathbf{z}_a, \bar{\mathbf{z}}) = x_{\ell a} p_a(\mathbf{z}_a, \bar{\mathbf{z}})$ with initial condition $x_{\ell 1} = \bar{x}_{\ell 1} = 1$. Thus, using Eq. 8, we can write the expected lifetime number of offspring of a mutant with pair $\mathbf{z} = (\mathbf{x}; \mathbf{y})$ in the context of a resident with pair $\bar{\mathbf{z}} = (\bar{\mathbf{x}}; \bar{\mathbf{y}})$ as

$$R_0(\mathbf{z}, \bar{\mathbf{z}}) = \sum_{a=1}^{N_\mathrm{a}} x_{\ell a} f_a(\mathbf{z}_a, \bar{\mathbf{z}}). \tag{Eq. S4.1a}$$

Consider the optimization problem of finding an optimal pair $\mathbf{z}^* = (\mathbf{x}^*; \mathbf{y}^*)$ such that

$$\mathbf{y}^* \in \arg\max_{\mathbf{y}} R_0(\mathbf{z}, \mathbf{z}^*), \tag{Eq. S4.1b}$$

subject to the dynamic constraint

$$\mathbf{x}_{a+1} = \mathbf{g}_a(\mathbf{z}_a, \bar{\mathbf{z}}), \tag{Eq. S4.1c}$$

for $a \in \{1, \ldots, N_\mathrm{a}\}$, with $\mathbf{x}_1 = \bar{\mathbf{x}}_1$ given and $\mathbf{x}_{N_\mathrm{a}}$ free. Hence, $\mathbf{z}^*$ is a best response to itself under the best response function $R_0$, where $\mathbf{y}^*$ is an optimal control and $\mathbf{x}^*$ is its associated optimal state. The optimization problem in Eq. S4.1 is a standard life-history problem generalized to include social interactions. Then, it follows that since there is no exogenous environmental change, an admissible locally stable evolutionary equilibrium $\mathbf{z}^*$ of Layer 7, Eq. 9c for $\boldsymbol{\zeta} = \mathbf{z}$ locally solves the problem in Eq. S4.1.

Second, we define the costate variables for the problem in Eq. S4.1 and show that they are proportional to the total selection gradient of states evaluated at an admissible locally stable evolutionary equilibrium. The costate variable of the $i$-th state variable at age $a$ for problem Eq. S4.1 is defined as

$$k_{x_{ia}} \equiv \left.\frac{\mathrm{d}R_0}{\mathrm{d}x_{ia}}\right|_{\mathbf{z}=\bar{\mathbf{z}}=\mathbf{z}^*} \tag{Eq. S4.2}$$

(section 9.6 of Sydsæter *et al.* 2008). Hence, from Eq. S2.3.12b, we have that the costate for the $i$-th state variable at age $a$ is

$$k_{x_{ia}} = T \left.\frac{\mathrm{d}w}{\mathrm{d}x_{ia}}\right|_{\mathbf{z}=\bar{\mathbf{z}}=\mathbf{z}^*}. \tag{Eq. S4.3}$$

That is, costate variables are proportional to the total selection gradient of state variables at an admissible locally stable evolutionary equilibrium $\mathbf{z}^*$. The total selection gradient of states thus generalizes the costate notion to the situation where controls and states are outside of evolutionary equilibrium for the life-history problem of $R_0$ maximization (Metz *et al.*, 2016). We have obtained various equations (Layer 4, Eq. S20) that enable direct calculation of such generalized costates in age structured models with $R_0$ maximization. Moreover, we have obtained an equation that relates such generalized costates to the evolutionary dynamics (fifth line of Layer 4, Eq. S21). Since we are assuming that there are no environmental traits, total immediate effect matrices reduce to direct effect matrices. Thus, the fifth line of Layer 4, Eq. S21 shows that such generalized costates affect the evolutionary dynamics indirectly by being transformed by the direct effects of controls on states, $\partial\mathbf{x}^\intercal/\partial\mathbf{y}$.

Third, we show that total maximization of $R_0$ is equivalent to direct maximization of the Hamiltonian, which is the central feature of Pontryagin's maximum principle. We have that the total selection gradient of controls can be written in terms of the total selection gradients of states (fifth line of Layer 4, Eq. S21), so for the controls at age $a$ we have

$$\left.\frac{\mathrm{d}w}{\mathrm{d}\mathbf{y}_a}\right|_{\mathbf{y}=\bar{\mathbf{y}}} = \left.\left(\frac{\partial\mathbf{x}^\intercal}{\partial\mathbf{y}_a}\frac{\mathrm{d}w}{\mathrm{d}\mathbf{x}} + \frac{\partial w}{\partial\mathbf{y}_a}\right)\right|_{\mathbf{y}=\bar{\mathbf{y}}},$$

where we substituted total immediate derivatives for partial derivatives because we are assuming that there are no environmental traits. Using Eq. S2.3.12 yields

$$\left.\frac{\mathrm{d}R_0}{\mathrm{d}\mathbf{y}_a}\right|_{\mathbf{y}=\bar{\mathbf{y}}} = \left.\left(\frac{\partial\mathbf{x}^\intercal}{\partial\mathbf{y}_a}\frac{\mathrm{d}R_0}{\mathrm{d}\mathbf{x}} + \frac{\partial R_0}{\partial\mathbf{y}_a}\right)\right|_{\mathbf{y}=\bar{\mathbf{y}}}.$$

From Eq. S5.2.10 and Eq. S4.1a given that the partial derivative ignores the dynamic constraint (Eq. S4.1c), it follows that

$$\left.\frac{\mathrm{d}R_0}{\mathrm{d}\mathbf{y}_a}\right|_{\mathbf{y}=\bar{\mathbf{y}}} = \left.\left(\frac{\partial\mathbf{x}_{a+1}^\intercal}{\partial\mathbf{y}_a}\frac{\mathrm{d}R_0}{\mathrm{d}\mathbf{x}_{a+1}} + \frac{\partial(x_{\ell a} f_a)}{\partial\mathbf{y}_a}\right)\right|_{\mathbf{y}=\bar{\mathbf{y}}}.$$

Using Eq. S4.2 and Eq. S4.1c and evaluating at optimal controls yields

$$\left.\frac{\mathrm{d}R_0}{\mathrm{d}\mathbf{y}_a}\right|_{\mathbf{y}=\bar{\mathbf{y}}=\mathbf{y}^*} = \left.\left(\frac{\partial\mathbf{g}_a^\intercal}{\partial\mathbf{y}_a}\mathbf{k}_{\mathbf{x}_{a+1}} + \frac{\partial(x_{\ell a} f_a)}{\partial\mathbf{y}_a}\right)\right|_{\mathbf{y}=\bar{\mathbf{y}}=\mathbf{y}^*}. \tag{Eq. S4.4}$$

This suggests to define

$$\mathcal{H}_a \equiv \mathbf{g}_a^\top \mathbf{k}_{\mathbf{x}_{a+1}} + x_{\ell a} f_a, \qquad \text{(Eq. S4.5)}$$

which recovers the Hamiltonian of Pontryagin's maximum principle in discrete time (section 12.5 of Sydsæter *et al.* 2008) for the objective function Eq. S4.1a. Then, the total derivative of the objective function with respect to the controls at a given age equals the partial derivative of the Hamiltonian when both derivatives are evaluated at optimal controls:

$$\left.\frac{\mathrm{d}R_0}{\mathrm{d}\mathbf{y}_a}\right|_{\mathbf{y}=\bar{\mathbf{y}}=\mathbf{y}^*} = \left.\frac{\partial \mathcal{H}_a}{\partial \mathbf{y}_a}\right|_{\mathbf{y}=\bar{\mathbf{y}}=\mathbf{y}^*}.$$

This is the essence of Pontryagin's maximum principle: the signs of the left-hand side derivatives are the same as the signs of the derivatives on the right-hand side, which are simpler to compute (although one must then compute costate variables).

Fourth, we show that the formulas we found for the costate variables (Eq. S4.2) imply the costate equations of Pontryagin's maximum principle for discrete time. Such costate equations are dynamic equations that allow one to calculate the costate variables. Using Layer 4, Eq. S20 and Eq. S2.3.12, we have that

$$\left.\frac{\mathrm{d}R_0}{\mathrm{d}\mathbf{x}_a}\right|_{\mathbf{y}=\bar{\mathbf{y}}} = \left.\left(\frac{\mathrm{d}\mathbf{x}^\top}{\mathrm{d}\mathbf{x}_a}\frac{\partial R_0}{\partial \mathbf{x}}\right)\right|_{\mathbf{y}=\bar{\mathbf{y}}}.$$

Expanding the matrix multiplication on the right-hand side, this is

$$\begin{aligned}\left.\frac{\mathrm{d}R_0}{\mathrm{d}\mathbf{x}_a}\right|_{\mathbf{y}=\bar{\mathbf{y}}} &= \left.\left(\sum_{j=1}^{N_\mathrm{a}}\frac{\mathrm{d}\mathbf{x}_j^\top}{\mathrm{d}\mathbf{x}_a}\frac{\partial R_0}{\partial \mathbf{x}_j}\right)\right|_{\mathbf{y}=\bar{\mathbf{y}}} \\ &= \left.\left(\frac{\partial R_0}{\partial \mathbf{x}_a} + \sum_{j=a+1}^{N_\mathrm{a}}\frac{\mathrm{d}\mathbf{x}_j^\top}{\mathrm{d}\mathbf{x}_a}\frac{\partial R_0}{\partial \mathbf{x}_j}\right)\right|_{\mathbf{y}=\bar{\mathbf{y}}},\end{aligned}$$

where we used Eq. S5.1.15. Using the expression of the total effect of states on themselves as a product (12) yields

$$\left.\frac{\mathrm{d}R_0}{\mathrm{d}\mathbf{x}_a}\right|_{\mathbf{y}=\bar{\mathbf{y}}} = \left.\left(\frac{\partial R_0}{\partial \mathbf{x}_a} + \sum_{j=a+1}^{N_\mathrm{a}}\frac{\partial\mathbf{x}_{a+1}^\top}{\partial \mathbf{x}_a}\frac{\mathrm{d}\mathbf{x}_j^\top}{\mathrm{d}\mathbf{x}_{a+1}}\frac{\partial R_0}{\partial \mathbf{x}_j}\right)\right|_{\mathbf{y}=\bar{\mathbf{y}}}.$$

Doing the sum over $j$ yields

$$\begin{aligned}\left.\frac{\mathrm{d}R_0}{\mathrm{d}\mathbf{x}_a}\right|_{\mathbf{y}=\bar{\mathbf{y}}} &= \left.\left(\frac{\partial R_0}{\partial \mathbf{x}_a} + \frac{\partial\mathbf{x}_{a+1}^\top}{\partial \mathbf{x}_a}\sum_{j=a+1}^{N_\mathrm{a}}\frac{\mathrm{d}\mathbf{x}_j^\top}{\mathrm{d}\mathbf{x}_{a+1}}\frac{\partial R_0}{\partial \mathbf{x}_j}\right)\right|_{\mathbf{y}=\bar{\mathbf{y}}} \\ &= \left.\left(\frac{\partial R_0}{\partial \mathbf{x}_a} + \frac{\partial\mathbf{x}_{a+1}^\top}{\partial \mathbf{x}_a}\frac{\mathrm{d}\mathbf{x}^\top}{\mathrm{d}\mathbf{x}_{a+1}}\frac{\partial R_0}{\partial \mathbf{x}}\right)\right|_{\mathbf{y}=\bar{\mathbf{y}}}.\end{aligned}$$

Using the second line of Layer 4, Eq. S20 and Eq. S2.3.12 again yields

$$\left.\frac{\mathrm{d}R_0}{\mathrm{d}\mathbf{x}_a}\right|_{\mathbf{y}=\bar{\mathbf{y}}} = \left.\left(\frac{\partial R_0}{\partial \mathbf{x}_a} + \frac{\partial\mathbf{x}_{a+1}^\top}{\partial \mathbf{x}_a}\frac{\mathrm{d}R_0}{\mathrm{d}\mathbf{x}_{a+1}}\right)\right|_{\mathbf{y}=\bar{\mathbf{y}}}. \qquad \text{(Eq. S4.6)}$$

This equals the partial derivative of the Hamiltonian with respect to the states at age $a$. Indeed, using Eq. S4.5 we have

$$\left.\frac{\partial \mathcal{H}_a}{\partial \mathbf{x}_a}\right|_{\mathbf{y}=\bar{\mathbf{y}}} = \left.\left(\frac{\partial\mathbf{x}_{a+1}^\top}{\partial \mathbf{x}_a}\frac{\mathrm{d}R_0}{\mathrm{d}\mathbf{x}_{a+1}} + \frac{\partial R_0}{\partial \mathbf{x}_a}\right)\right|_{\mathbf{y}=\bar{\mathbf{y}}}.$$

Substituting this in Eq. S4.6 and evaluating at optimal controls yields

$$\mathbf{k}_a = \left.\frac{\partial \mathcal{H}_a}{\partial \mathbf{x}_a}\right|_{\mathbf{y}=\bar{\mathbf{y}}=\mathbf{y}^*}.$$

This is the costate equation of Pontryagin's maximum principle in discrete time (Eq. 4 in section 12.5 of Sydsæter *et al.* 2008).

### S5 Derivation of results

This section contains the derivations of all our results.

#### S5.1 Total selection gradient of the phenotype

Here we derive the total selection gradient of the phenotype $\mathrm{d}\lambda/\mathrm{d}\mathbf{x}|_{\mathbf{y}=\bar{\mathbf{y}}}$, which is part of and simpler to derive than the total selection gradient of the genotype $\mathrm{d}\lambda/\mathrm{d}\mathbf{y}|_{\mathbf{y}=\bar{\mathbf{y}}}$.

##### S5.1.1 Total selection gradient of the phenotype in terms of direct fitness effects

We start by considering the total selection gradient of the $i$-th phenotype at age $a$. By this, we mean the total selection gradient of a perturbation of $x_{ia}$ taken as initial condition of the recurrence equation (1) when applied at the ages $\{a,\dots,n\}$. Consequently, a perturbation in a phenotype at a given age does not affect phenotypes at earlier ages, in short, due to *the arrow of developmental time*. By letting $\zeta$ in Eq. S2.3.9 be $x_{ia}$, we have

$$\left.\frac{\mathrm{d}\lambda}{\mathrm{d}x_{ia}}\right|_{\mathbf{y}=\bar{\mathbf{y}}} = \left.\frac{\mathrm{d}w}{\mathrm{d}x_{ia}}\right|_{\mathbf{y}=\bar{\mathbf{y}}} = \sum_{j=1}^{N_\mathrm{a}}\left.\frac{\mathrm{d}w_j}{\mathrm{d}x_{ia}}\right|_{\mathbf{y}=\bar{\mathbf{y}}}. \qquad \text{(Eq. S5.1.1)}$$

Note that the total derivatives of a mutant's relative fitness at age $j$ in Eq. S5.1.1 are with respect to the individual's phenotype at possibly another age $a$. From Eq. S2.3.7, we have that a mutant's relative fitness at age $j$, $w_j(\mathbf{z}_j, \mathbf{h}_j(\mathbf{z}_j, \bar{\mathbf{z}}, \tau), \bar{\mathbf{m}})$, depends on the individual's phenotype at the current age (recall $\mathbf{z}_j = (\mathbf{x}_j;\mathbf{y}_j)$), but from the developmental constraint (1) the phenotype at a given age depends on the phenotype at previous ages. We must then calculate the total derivatives of fitness in Eq. S5.1.1 in terms of direct (i.e., partial) derivatives, thus separating the effects of phenotypes at the current age from those of phenotypes at other ages.

To do this, we start by applying the chain rule, and since we assume that genotypic traits are developmentally independent (hence, they do not depend on the phenotype, so $\mathrm{d}\mathbf{y}_j/\mathrm{d}x_{ia} = \mathbf{0}$ for all $i \in \{1,\dots,N_\mathrm{p}\}$ and all $a, j \in \{1,\dots,N_\mathrm{a}\}$), we obtain

$$\left.\frac{\mathrm{d}w_j}{\mathrm{d}x_{ia}}\right|_{\mathbf{y}=\bar{\mathbf{y}}} = \left.\left(\sum_{k=1}^{N_\mathrm{p}}\frac{\partial w_j}{\partial x_{kj}}\frac{\mathrm{d}x_{kj}}{\mathrm{d}x_{ia}} + \sum_{k=1}^{N_\mathrm{p}}\sum_{r=1}^{N_\mathrm{e}}\frac{\partial w_j}{\partial \epsilon_{rj}}\frac{\partial \epsilon_{rj}}{\partial x_{kj}}\frac{\mathrm{d}x_{kj}}{\mathrm{d}x_{ia}}\right)\right|_{\mathbf{y}=\bar{\mathbf{y}}}.$$

Applying matrix calculus notation (Appendix A), this is

$$\left.\frac{\mathrm{d}w_j}{\mathrm{d}x_{ia}}\right|_{\mathbf{y}=\bar{\mathbf{y}}} = \left.\left(\frac{\mathrm{d}\mathbf{x}_j^\top}{\mathrm{d}x_{ia}}\frac{\partial w_j}{\partial \mathbf{x}_j} + \sum_{k=1}^{N_\mathrm{p}}\frac{\partial \boldsymbol{\epsilon}_j^\top}{\partial x_{kj}}\frac{\partial w_j}{\partial \boldsymbol{\epsilon}_j}\frac{\mathrm{d}x_{kj}}{\mathrm{d}x_{ia}}\right)\right|_{\mathbf{y}=\bar{\mathbf{y}}}.$$

Applying matrix calculus notation again yields

$$\left.\frac{\mathrm{d}w_j}{\mathrm{d}x_{ia}}\right|_{\mathbf{y}=\bar{\mathbf{y}}} = \left.\left(\frac{\mathrm{d}\mathbf{x}_j^\intercal}{\mathrm{d}x_{ia}}\frac{\partial w_j}{\partial \mathbf{x}_j} + \frac{\mathrm{d}\mathbf{x}_j^\intercal}{\mathrm{d}x_{ia}}\frac{\partial \boldsymbol{\epsilon}_j^\intercal}{\partial \mathbf{x}_j}\frac{\partial w_j}{\partial \boldsymbol{\epsilon}_j}\right)\right|_{\mathbf{y}=\bar{\mathbf{y}}}.$$

Factorizing, we have

$$\left.\frac{\mathrm{d}w_j}{\mathrm{d}x_{ia}}\right|_{\mathbf{y}=\bar{\mathbf{y}}} = \left.\left[\frac{\mathrm{d}\mathbf{x}_j^\intercal}{\mathrm{d}x_{ia}}\left(\frac{\partial w_j}{\partial \mathbf{x}_j} + \frac{\partial \boldsymbol{\epsilon}_j^\intercal}{\partial \mathbf{x}_j}\frac{\partial w_j}{\partial \boldsymbol{\epsilon}_j}\right)\right]\right|_{\mathbf{y}=\bar{\mathbf{y}}}. \quad \text{(Eq. S5.1.2)}$$

Eq. S5.1.2 now contains only partial derivatives of age-specific fitness.

We now write Eq. S5.1.2 in terms of partial derivatives of lifetime fitness. Consider the *direct selection gradient of the phenotype at age j* defined as

$$\left.\frac{\partial w}{\partial \mathbf{x}_j}\right|_{\mathbf{y}=\bar{\mathbf{y}}} \equiv \left.\left(\frac{\partial w}{\partial x_{1j}},\ldots,\frac{\partial w}{\partial x_{N_\mathrm{p}j}}\right)^\intercal\right|_{\mathbf{y}=\bar{\mathbf{y}}} \in \mathbb{R}^{N_\mathrm{p}\times 1}.$$

Such selection gradient of the phenotype at age $j$ forms the selection gradient of the phenotype at all ages (Layer 2, Eq. S1). Similarly, the *direct selection gradient of the environment at age j* is

$$\left.\frac{\partial w}{\partial \boldsymbol{\epsilon}_j}\right|_{\mathbf{y}=\bar{\mathbf{y}}} \equiv \left.\left(\frac{\partial w}{\partial \epsilon_{1j}},\ldots,\frac{\partial w}{\partial \epsilon_{N_\mathrm{e}j}}\right)^\intercal\right|_{\mathbf{y}=\bar{\mathbf{y}}} \in \mathbb{R}^{N_\mathrm{e}\times 1},$$

and the matrix of *direct effects of a mutant's phenotype at age j on her environment at age j* is

$$\left.\frac{\partial \boldsymbol{\epsilon}_j^\intercal}{\partial \mathbf{x}_j}\right|_{\mathbf{y}=\bar{\mathbf{y}}} \equiv \left.\begin{pmatrix} \frac{\partial \epsilon_{1j}}{\partial x_{1j}} & \cdots & \frac{\partial \epsilon_{N_\mathrm{e}j}}{\partial x_{1j}} \\ \vdots & \ddots & \vdots \\ \frac{\partial \epsilon_{1j}}{\partial x_{N_\mathrm{p}j}} & \cdots & \frac{\partial \epsilon_{N_\mathrm{e}j}}{\partial x_{N_\mathrm{p}j}} \end{pmatrix}\right|_{\mathbf{y}=\bar{\mathbf{y}}} \in \mathbb{R}^{N_\mathrm{p}\times N_\mathrm{e}}.$$

From Eq. S2.3.8, $w$ only depends directly on $\mathbf{x}_j$, $\mathbf{y}_j$, and $\boldsymbol{\epsilon}_j$ through $w_j$. So,

$$\frac{\partial w_j}{\partial \mathbf{x}_j} = \frac{\partial w}{\partial \mathbf{x}_j} \quad \text{(Eq. S5.1.3a)}$$

$$\frac{\partial w_j}{\partial \mathbf{y}_j} = \frac{\partial w}{\partial \mathbf{y}_j} \quad \text{(Eq. S5.1.3b)}$$

$$\frac{\partial w_j}{\partial \boldsymbol{\epsilon}_j} = \frac{\partial w}{\partial \boldsymbol{\epsilon}_j}, \quad \text{(Eq. S5.1.3c)}$$

which substituted in Eq. S5.1.2 yields

$$\begin{aligned}\left.\frac{\mathrm{d}w_j}{\mathrm{d}x_{ia}}\right|_{\mathbf{y}=\bar{\mathbf{y}}} &= \left.\left[\frac{\mathrm{d}\mathbf{x}_j^\intercal}{\mathrm{d}x_{ia}}\left(\frac{\partial w}{\partial \mathbf{x}_j} + \frac{\partial \boldsymbol{\epsilon}_j^\intercal}{\partial \mathbf{x}_j}\frac{\partial w}{\partial \boldsymbol{\epsilon}_j}\right)\right]\right|_{\mathbf{y}=\bar{\mathbf{y}}} \\ &= \left.\left(\frac{\mathrm{d}\mathbf{x}_j^\intercal}{\mathrm{d}x_{ia}}\frac{\delta w}{\delta \mathbf{x}_j}\right)\right|_{\mathbf{y}=\bar{\mathbf{y}}}, \quad \text{(Eq. S5.1.4)}\end{aligned}$$

where the *total immediate selection gradient of the phenotype at age j* is

$$\left.\frac{\delta w}{\delta \mathbf{x}_j}\right|_{\mathbf{y}=\bar{\mathbf{y}}} = \left.\left(\frac{\partial w}{\partial \mathbf{x}_j} + \frac{\partial \boldsymbol{\epsilon}_j^\intercal}{\partial \mathbf{x}_j}\frac{\partial w}{\partial \boldsymbol{\epsilon}_j}\right)\right|_{\mathbf{y}=\bar{\mathbf{y}}} \in \mathbb{R}^{N_\mathrm{p}\times 1}. \quad \text{(Eq. S5.1.5)}$$

Consider now the total immediate selection gradient of the phenotype at all ages. The block column vector of *total immediate effects of a mutant's phenotype on fitness* is

$$\left.\frac{\delta w}{\delta \mathbf{x}}\right|_{\mathbf{y}=\bar{\mathbf{y}}} \equiv \left.\left(\frac{\delta w}{\delta \mathbf{x}_1};\cdots;\frac{\delta w}{\delta \mathbf{x}_{N_\mathrm{a}}}\right)\right|_{\mathbf{y}=\bar{\mathbf{y}}} \in \mathbb{R}^{N_\mathrm{a}N_\mathrm{p}\times 1}.$$

Using Layer 2, Eq. S2d, we have that

$$\frac{\partial \boldsymbol{\epsilon}^\intercal}{\partial \mathbf{x}}\frac{\partial w}{\partial \boldsymbol{\epsilon}} = \left(\sum_{k=1}^{N_\mathrm{a}}\frac{\partial \boldsymbol{\epsilon}_k^\intercal}{\partial \mathbf{x}_j}\frac{\partial w}{\partial \boldsymbol{\epsilon}_k}\right) = \left(\frac{\partial \boldsymbol{\epsilon}_j^\intercal}{\partial \mathbf{x}_j}\frac{\partial w}{\partial \boldsymbol{\epsilon}_j}\right) \quad \text{(Eq. S5.1.6)}$$

is a block column vector whose $j$-th entry equals the rightmost term in Eq. S5.1.5. Thus, from Eq. S5.1.5, Layer 2, Eq. S1, and Eq. S5.1.6, it follows that the total immediate selection gradient of the phenotype is given by Layer 3, Eq. S1.

Now, we write the total selection gradient of $x_{ia}$ in terms of the total immediate selection gradient of the phenotype. Substituting Eq. S5.1.4 in Eq. S5.1.1 yields

$$\left.\frac{\mathrm{d}w}{\mathrm{d}x_{ia}}\right|_{\mathbf{y}=\bar{\mathbf{y}}} = \sum_{j=1}^{N_\mathrm{a}}\left.\left(\frac{\mathrm{d}\mathbf{x}_j^\intercal}{\mathrm{d}x_{ia}}\frac{\delta w}{\delta \mathbf{x}_j}\right)\right|_{\mathbf{y}=\bar{\mathbf{y}}} = \left.\left(\frac{\mathrm{d}\mathbf{x}^\intercal}{\mathrm{d}x_{ia}}\frac{\delta w}{\delta \mathbf{x}}\right)\right|_{\mathbf{y}=\bar{\mathbf{y}}},$$

where we use the block row vector

$$\frac{\mathrm{d}\mathbf{x}^\intercal}{\mathrm{d}x_{ia}} = \left(\frac{\mathrm{d}\mathbf{x}_1^\intercal}{\mathrm{d}x_{ia}},\ldots,\frac{\mathrm{d}\mathbf{x}_{N_\mathrm{a}}^\intercal}{\mathrm{d}x_{ia}}\right) \in \mathbb{R}^{1\times N_\mathrm{a}N_\mathrm{p}}.$$

Therefore, the total selection gradient of all phenotypes across all ages is

$$\left.\frac{\mathrm{d}w}{\mathrm{d}\mathbf{x}}\right|_{\mathbf{y}=\bar{\mathbf{y}}} = \left.\left(\frac{\mathrm{d}\mathbf{x}^\intercal}{\mathrm{d}\mathbf{x}}\frac{\delta w}{\delta \mathbf{x}}\right)\right|_{\mathbf{y}=\bar{\mathbf{y}}} \in \mathbb{R}^{N_\mathrm{a}N_\mathrm{p}\times 1}, \quad \text{(Eq. S5.1.7)}$$

where the total immediate selection gradient of the phenotype is given by Layer 3, Eq. S1 and the block matrix of *total effects of a mutant's phenotype on her phenotype* is

$$\left.\frac{\mathrm{d}\mathbf{x}^\intercal}{\mathrm{d}\mathbf{x}}\right|_{\mathbf{y}=\bar{\mathbf{y}}} \equiv \left.\begin{pmatrix} \frac{\mathrm{d}\mathbf{x}_1^\intercal}{\mathrm{d}\mathbf{x}_1} & \cdots & \frac{\mathrm{d}\mathbf{x}_{N_\mathrm{a}}^\intercal}{\partial \mathbf{x}_1} \\ \vdots & \ddots & \vdots \\ \frac{\mathrm{d}\mathbf{x}_1^\intercal}{\mathrm{d}\mathbf{x}_{N_\mathrm{a}}} & \cdots & \frac{\mathrm{d}\mathbf{x}_{N_\mathrm{a}}^\intercal}{\mathrm{d}\mathbf{x}_{N_\mathrm{a}}} \end{pmatrix}\right|_{\mathbf{y}=\bar{\mathbf{y}}} \in \mathbb{R}^{N_\mathrm{a}N_\mathrm{p}\times N_\mathrm{a}N_\mathrm{p}}.$$

Using Layer 3, Eq. S1, expression Eq. S5.1.7 is now in terms of partial derivatives of fitness, partial derivatives of the environment, and total effects of a mutant's phenotype on her phenotype, $\mathrm{d}\mathbf{x}^\intercal/\mathrm{d}\mathbf{x}$, which we now proceed to write in terms of partial derivatives only.

##### S5.1.2 Matrix of total effects of a mutant's phenotype on her phenotype

From the developmental constraint (1) for the $k$-th phenotype at age $j \in \{2,\ldots,N_\mathrm{a}\}$ we have that $x_{kj} = g_{k,j-1}(\mathbf{z}_{j-1},\mathbf{h}_{j-1}(\mathbf{z}_{j-1},\bar{\mathbf{z}},\tau),\bar{\mathbf{z}})$, so using the chain rule and since genotypic traits are developmentally independent we obtain

$$\left.\frac{\mathrm{d}x_{kj}}{\mathrm{d}x_{ia}}\right|_{\mathbf{y}=\bar{\mathbf{y}}} = \Bigg(\sum_{l=1}^{N_\mathrm{p}}\frac{\partial g_{k,j-1}}{\partial x_{l,j-1}}\frac{\mathrm{d}x_{l,j-1}}{\mathrm{d}x_{ia}}$$

$$+\sum_{l=1}^{N_\mathrm{p}}\sum_{r=1}^{N_\mathrm{e}} \frac{\partial g_{k,j-1}}{\partial \epsilon_{r,j-1}} \frac{\partial \epsilon_{r,j-1}}{\partial x_{l,j-1}} \frac{\mathrm{d}x_{l,j-1}}{\mathrm{d}x_{ia}}\Bigg)\Bigg|_{\mathbf{y}=\bar{\mathbf{y}}}.$$

Applying matrix calculus notation (Appendix A), this is

$$\left.\frac{\mathrm{d}x_{kj}}{\mathrm{d}x_{ia}}\right|_{\mathbf{y}=\bar{\mathbf{y}}} = \left.\left(\frac{\mathrm{d}\mathbf{x}_{j-1}^\intercal}{\mathrm{d}x_{ia}} \frac{\partial g_{k,j-1}}{\partial \mathbf{x}_{j-1}} + \sum_{l=1}^{N_\mathrm{p}} \frac{\partial \boldsymbol{\epsilon}_{j-1}^\intercal}{\partial x_{l,j-1}} \frac{\partial g_{k,j-1}}{\partial \boldsymbol{\epsilon}_{j-1}} \frac{\mathrm{d}x_{l,j-1}}{\mathrm{d}x_{ia}}\right)\right|_{\mathbf{y}=\bar{\mathbf{y}}}.$$

Applying matrix calculus notation again yields

$$\left.\frac{\mathrm{d}x_{kj}}{\mathrm{d}x_{ia}}\right|_{\mathbf{y}=\bar{\mathbf{y}}} = \left.\left(\frac{\mathrm{d}\mathbf{x}_{j-1}^\intercal}{\mathrm{d}x_{ia}} \frac{\partial g_{k,j-1}}{\partial \mathbf{x}_{j-1}} + \frac{\mathrm{d}\mathbf{x}_{j-1}^\intercal}{\mathrm{d}x_{ia}} \frac{\partial \boldsymbol{\epsilon}_{j-1}^\intercal}{\partial \mathbf{x}_{j-1}} \frac{\partial g_{k,j-1}}{\partial \boldsymbol{\epsilon}_{j-1}}\right)\right|_{\mathbf{y}=\bar{\mathbf{y}}}.$$

Factorizing, we have

$$\left.\frac{\mathrm{d}x_{kj}}{\mathrm{d}x_{ia}}\right|_{\mathbf{y}=\bar{\mathbf{y}}} = \left.\left[\frac{\mathrm{d}\mathbf{x}_{j-1}^\intercal}{\mathrm{d}x_{ia}}\left(\frac{\partial g_{k,j-1}}{\partial \mathbf{x}_{j-1}} + \frac{\partial \boldsymbol{\epsilon}_{j-1}^\intercal}{\partial \mathbf{x}_{j-1}} \frac{\partial g_{k,j-1}}{\partial \boldsymbol{\epsilon}_{j-1}}\right)\right]\right|_{\mathbf{y}=\bar{\mathbf{y}}}.$$

Rewriting $g_{k,j-1}$ as $x_{kj}$ yields

$$\left.\frac{\mathrm{d}x_{kj}}{\mathrm{d}x_{ia}}\right|_{\mathbf{y}=\bar{\mathbf{y}}} = \left.\left[\frac{\mathrm{d}\mathbf{x}_{j-1}^\intercal}{\mathrm{d}x_{ia}}\left(\frac{\partial x_{kj}}{\partial \mathbf{x}_{j-1}} + \frac{\partial \boldsymbol{\epsilon}_{j-1}^\intercal}{\partial \mathbf{x}_{j-1}} \frac{\partial x_{kj}}{\partial \boldsymbol{\epsilon}_{j-1}}\right)\right]\right|_{\mathbf{y}=\bar{\mathbf{y}}}.$$

Hence,

$$\left.\frac{\mathrm{d}\mathbf{x}_{j}^\intercal}{\mathrm{d}x_{ia}}\right|_{\mathbf{y}=\bar{\mathbf{y}}} = \left.\left[\frac{\mathrm{d}\mathbf{x}_{j-1}^\intercal}{\mathrm{d}x_{ia}}\left(\frac{\partial \mathbf{x}_{j}^\intercal}{\partial \mathbf{x}_{j-1}} + \frac{\partial \boldsymbol{\epsilon}_{j-1}^\intercal}{\partial \mathbf{x}_{j-1}} \frac{\partial \mathbf{x}_{j}^\intercal}{\partial \boldsymbol{\epsilon}_{j-1}}\right)\right]\right|_{\mathbf{y}=\bar{\mathbf{y}}}, \tag{Eq. S5.1.8}$$

where we use the matrix of *direct effects of a mutant's phenotype at age j on her phenotype at age j + 1*

$$\left.\frac{\partial \mathbf{x}_{j+1}^\intercal}{\partial \mathbf{x}_j}\right|_{\mathbf{y}=\bar{\mathbf{y}}} \equiv \left.\begin{pmatrix} \frac{\partial x_{1,j+1}}{\partial x_{1j}} & \cdots & \frac{\partial x_{N_\mathrm{p},j+1}}{\partial x_{1j}} \\ \vdots & \ddots & \vdots \\ \frac{\partial x_{1,j+1}}{\partial x_{N_\mathrm{p}j}} & \cdots & \frac{\partial x_{N_\mathrm{p},j+1}}{\partial x_{N_\mathrm{p}j}} \end{pmatrix}\right|_{\mathbf{y}=\bar{\mathbf{y}}} \in \mathbb{R}^{N_\mathrm{p}\times N_\mathrm{p}},$$

and the matrix of *direct effects of a mutant's environment at age j on her phenotype at age j + 1*

$$\left.\frac{\partial \mathbf{x}_{j+1}^\intercal}{\partial \boldsymbol{\epsilon}_j}\right|_{\mathbf{y}=\bar{\mathbf{y}}} \equiv \left.\begin{pmatrix} \frac{\partial x_{1,j+1}}{\partial \epsilon_{1j}} & \cdots & \frac{\partial x_{N_\mathrm{p},j+1}}{\partial \epsilon_{1j}} \\ \vdots & \ddots & \vdots \\ \frac{\partial x_{1,j+1}}{\partial \epsilon_{N_\mathrm{e}j}} & \cdots & \frac{\partial x_{N_\mathrm{p},j+1}}{\partial \epsilon_{N_\mathrm{e}j}} \end{pmatrix}\right|_{\mathbf{y}=\bar{\mathbf{y}}} \in \mathbb{R}^{N_\mathrm{e}\times N_\mathrm{p}}.$$

We can write Eq. S5.1.8 more succinctly as

$$\left.\frac{\mathrm{d}\mathbf{x}_{j}^\intercal}{\mathrm{d}x_{ia}}\right|_{\mathbf{y}=\bar{\mathbf{y}}} = \left.\left(\frac{\mathrm{d}\mathbf{x}_{j-1}^\intercal}{\mathrm{d}x_{ia}} \frac{\delta \mathbf{x}_{j}^\intercal}{\delta \mathbf{x}_{j-1}}\right)\right|_{\mathbf{y}=\bar{\mathbf{y}}}, \tag{Eq. S5.1.9}$$

where we use the matrix of *total immediate effects of a mutant's phenotype at age j on her phenotype at age j + 1*

$$\left.\frac{\delta \mathbf{x}_{j+1}^\intercal}{\delta \mathbf{x}_j}\right|_{\mathbf{y}=\bar{\mathbf{y}}} = \left.\left(\frac{\partial \mathbf{x}_{j+1}^\intercal}{\partial \mathbf{x}_j} + \frac{\partial \boldsymbol{\epsilon}_{j}^\intercal}{\partial \mathbf{x}_j}\frac{\partial \mathbf{x}_{j+1}^\intercal}{\partial \boldsymbol{\epsilon}_j}\right)\right|_{\mathbf{y}=\bar{\mathbf{y}}} \in \mathbb{R}^{N_\mathrm{p}\times N_\mathrm{p}}. \tag{Eq. S5.1.10}$$

The block matrix of *total immediate effects a mutant's phenotype on her phenotype* is

$$\left.\frac{\delta \mathbf{x}^\intercal}{\delta \mathbf{x}}\right|_{\mathbf{y}=\bar{\mathbf{y}}} \equiv \left.\begin{pmatrix} \frac{\delta \mathbf{x}_1^\intercal}{\delta \mathbf{x}_1} & \cdots & \frac{\delta \mathbf{x}_{N_\mathrm{a}}^\intercal}{\delta \mathbf{x}_1} \\ \vdots & \ddots & \vdots \\ \frac{\delta \mathbf{x}_1^\intercal}{\delta \mathbf{x}_{N_\mathrm{a}}} & \cdots & \frac{\delta \mathbf{x}_{N_\mathrm{a}}^\intercal}{\delta \mathbf{x}_{N_\mathrm{a}}} \end{pmatrix}\right|_{\mathbf{y}=\bar{\mathbf{y}}} = \left.\begin{pmatrix} \mathbf{I} & \frac{\delta \mathbf{x}_2^\intercal}{\delta \mathbf{x}_1} & \cdots & \mathbf{0} & \mathbf{0} \\ \mathbf{0} & \mathbf{I} & \cdots & \mathbf{0} & \mathbf{0} \\ \vdots & \vdots & \ddots & \vdots & \vdots \\ \mathbf{0} & \mathbf{0} & \cdots & \mathbf{I} & \frac{\delta \mathbf{x}_{N_\mathrm{a}}^\intercal}{\delta \mathbf{x}_{N_\mathrm{a}-1}} \\ \mathbf{0} & \mathbf{0} & \cdots & \mathbf{0} & \mathbf{I} \end{pmatrix}\right|_{\mathbf{y}=\bar{\mathbf{y}}} \tag{Eq. S5.1.11}$$

$$\in \mathbb{R}^{N_\mathrm{a}N_\mathrm{p}\times N_\mathrm{a}N_\mathrm{p}}.$$

The equality Eq. S5.1.11 follows because total immediate effects of a mutant's phenotype on her phenotype are only non-zero at the next age (from the developmental constraint in Eq. 1) or when a variable is differentiated with respect to itself. Using Layer 2, Eq. S2d and Layer 2, Eq. S2c, we have that

$$\frac{\partial \boldsymbol{\epsilon}^\intercal}{\partial \mathbf{x}}\frac{\partial \mathbf{x}^\intercal}{\partial \boldsymbol{\epsilon}} = \left(\sum_{k=1}^{N_\mathrm{a}} \frac{\partial \boldsymbol{\epsilon}_k^\intercal}{\partial \mathbf{x}_a}\frac{\partial \mathbf{x}_j^\intercal}{\partial \boldsymbol{\epsilon}_k}\right) = \left(\begin{cases} \frac{\partial \boldsymbol{\epsilon}_a^\intercal}{\partial \mathbf{x}_a}\frac{\partial \mathbf{x}_j^\intercal}{\partial \boldsymbol{\epsilon}_a} & \text{for } j = a+1 \\ \mathbf{0} & \text{for } j \neq a+1 \end{cases}\right), \tag{Eq. S5.1.12}$$

which equals the rightmost term in Eq. S5.1.10 for $j = a+1$. Thus, from Eq. S5.1.10, Layer 2, Eq. S2a, Eq. S5.1.11, and Eq. S5.1.12, it follows that the block matrix of total immediate effects of a mutant's phenotype on her phenotype satisfies Layer 3, Eq. S3.

Eq. S5.1.9 gives the matrix of total effects of the $i$-th phenotype of a mutant at age $a$ on her phenotype at age $j$. Then, it follows that the matrix of total effects of all the phenotypes of a mutant at age $a$ on her phenotype at age $j$ is

$$\left.\frac{\mathrm{d}\mathbf{x}_{j}^\intercal}{\mathrm{d}\mathbf{x}_{a}}\right|_{\mathbf{y}=\bar{\mathbf{y}}} = \left.\left(\frac{\mathrm{d}\mathbf{x}_{j-1}^\intercal}{\mathrm{d}\mathbf{x}_{a}} \frac{\delta \mathbf{x}_{j}^\intercal}{\delta \mathbf{x}_{j-1}}\right)\right|_{\mathbf{y}=\bar{\mathbf{y}}}. \tag{Eq. S5.1.13}$$

Eq. S5.1.13 is a recurrence equation for $\mathrm{d}\mathbf{x}_j^\intercal/\mathrm{d}\mathbf{x}_a$ over age $j \in \{2,\ldots,N_\mathrm{a}\}$. Because of the arrow of developmental time (due to the developmental constraint (1)), perturbations in an individual's late phenotype do not affect the individual's early phenotype (i.e., $\mathrm{d}\mathbf{x}_j^\intercal/\mathrm{d}\mathbf{x}_a = \mathbf{0}$ for $j < a$ and $j \in \{1,\ldots,N_\mathrm{a}-1\}$)[1]. Additionally, from the arrow of developmental time (Eq. 1), a perturbation in an individual's phenotype at a given age does not affect any other of the individual's phenotypes at the *same* age (i.e., $\mathrm{d}\mathbf{x}_a^\intercal/\mathrm{d}\mathbf{x}_a = \mathbf{I}$

[1] More specifically, we take the derivative $\mathrm{d}\mathbf{x}_j^\intercal/\mathrm{d}x_{ia}$ as referring to the effect on $\mathbf{x}_j^\intercal$ of a perturbation of the initial condition $\mathbf{x}_a$ of the difference equation (1) applied at the ages $\{a,\ldots,n\}$. Hence, if $j < a$, $\mathbf{x}_j^\intercal$ is unmodified by a change in the initial condition of (1) applied at the ages $\{a,\ldots,n\}$.

where $\mathbf{I}$ is the identity matrix). Hence, expanding the recurrence in Eq. S5.1.13, we obtain for $j \in \{1, \ldots, N_\mathrm{a}\}$ that

$$\left.\frac{\mathrm{d}\mathbf{x}_j^\intercal}{\mathrm{d}\mathbf{x}_a}\right|_{\mathbf{y}=\bar{\mathbf{y}}} = \begin{cases} \left.\left(\frac{\mathrm{d}\mathbf{x}_a^\intercal}{\mathrm{d}\mathbf{x}_a}\frac{\delta\mathbf{x}_{a+1}^\intercal}{\delta\mathbf{x}_a}\cdots\frac{\delta\mathbf{x}_j^\intercal}{\delta\mathbf{x}_{j-1}}\right)\right|_{\mathbf{y}=\bar{\mathbf{y}}} & \text{for } j > a \\ \left.\frac{\mathrm{d}\mathbf{x}_a^\intercal}{\mathrm{d}\mathbf{x}_a}\right|_{\mathbf{y}=\bar{\mathbf{y}}} & \text{for } j = a \\ \mathbf{0} & \text{for } j < a \end{cases}$$

$$= \begin{cases} \left.\left(\frac{\delta\mathbf{x}_{a+1}^\intercal}{\delta\mathbf{x}_a}\cdots\frac{\delta\mathbf{x}_j^\intercal}{\delta\mathbf{x}_{j-1}}\right)\right|_{\mathbf{y}=\bar{\mathbf{y}}} & \text{for } j > a \\ \mathbf{I} & \text{for } j = a \\ \mathbf{0} & \text{for } j < a. \end{cases}$$
(Eq. S5.1.14)

Thus, the block matrix of *total effects of a mutant's phenotype on her phenotype* is

$$\left.\frac{\mathrm{d}\mathbf{x}^\intercal}{\mathrm{d}\mathbf{x}}\right|_{\mathbf{y}=\bar{\mathbf{y}}} = \left.\begin{pmatrix} \frac{\mathrm{d}\mathbf{x}_1^\intercal}{\mathrm{d}\mathbf{x}_1} & \cdots & \frac{\mathrm{d}\mathbf{x}_{N_\mathrm{a}}^\intercal}{\mathrm{d}\mathbf{x}_1} \\ \vdots & \ddots & \vdots \\ \frac{\mathrm{d}\mathbf{x}_1^\intercal}{\mathrm{d}\mathbf{x}_{N_\mathrm{a}}} & \cdots & \frac{\mathrm{d}\mathbf{x}_{N_\mathrm{a}}^\intercal}{\mathrm{d}\mathbf{x}_{N_\mathrm{a}}} \end{pmatrix}\right|_{\mathbf{y}=\bar{\mathbf{y}}}$$

$$= \left.\begin{pmatrix} \mathbf{I} & \frac{\mathrm{d}\mathbf{x}_2^\intercal}{\mathrm{d}\mathbf{x}_1} & \cdots & \frac{\mathrm{d}\mathbf{x}_{N_\mathrm{a}-1}^\intercal}{\mathrm{d}\mathbf{x}_1} & \frac{\mathrm{d}\mathbf{x}_{N_\mathrm{a}}^\intercal}{\mathrm{d}\mathbf{x}_1} \\ \mathbf{0} & \mathbf{I} & \cdots & \frac{\mathrm{d}\mathbf{x}_{N_\mathrm{a}-1}^\intercal}{\mathrm{d}\mathbf{x}_2} & \frac{\mathrm{d}\mathbf{x}_{N_\mathrm{a}}^\intercal}{\mathrm{d}\mathbf{x}_2} \\ \vdots & \vdots & \ddots & \vdots & \vdots \\ \mathbf{0} & \mathbf{0} & \cdots & \mathbf{I} & \frac{\mathrm{d}\mathbf{x}_{N_\mathrm{a}}^\intercal}{\mathrm{d}\mathbf{x}_{N_\mathrm{a}-1}} \\ \mathbf{0} & \mathbf{0} & \cdots & \mathbf{0} & \mathbf{I} \end{pmatrix}\right|_{\mathbf{y}=\bar{\mathbf{y}}}$$
(Eq. S5.1.15)

$$\in \mathbb{R}^{N_\mathrm{a}N_\mathrm{p}\times N_\mathrm{a}N_\mathrm{p}},$$

which is block upper triangular and its $aj$-th block entry is given by Eq. (12). Eq. S5.1.15 and Eq. (12) write the matrix of total effects of a mutant's phenotype on her phenotype in terms of partial derivatives, given Eq. S5.1.10, as we sought.

From Eq. S5.1.15, it follows that the matrix of total effects of a mutant's phenotype on her phenotype $\mathrm{d}\mathbf{x}^\intercal/\mathrm{d}\mathbf{x}|_{\mathbf{y}=\bar{\mathbf{y}}}$ is invertible. Indeed, since $\mathrm{d}\mathbf{x}^\intercal/\mathrm{d}\mathbf{x}|_{\mathbf{y}=\bar{\mathbf{y}}}$ is square and block upper triangular, then its determinant is

$$\det\left(\left.\frac{\mathrm{d}\mathbf{x}^\intercal}{\mathrm{d}\mathbf{x}}\right|_{\mathbf{y}=\bar{\mathbf{y}}}\right) = \det\left(\left.\frac{\mathrm{d}\mathbf{x}_1^\intercal}{\mathrm{d}\mathbf{x}_1}\right|_{\mathbf{y}=\bar{\mathbf{y}}}\right)\cdots\det\left(\left.\frac{\mathrm{d}\mathbf{x}_{N_\mathrm{a}}^\intercal}{\mathrm{d}\mathbf{x}_{N_\mathrm{a}}}\right|_{\mathbf{y}=\bar{\mathbf{y}}}\right)$$

(Horn and Johnson, 2013, p. 32). Since $\mathrm{d}\mathbf{x}_a^\intercal/\mathrm{d}\mathbf{x}_a|_{\mathbf{y}=\bar{\mathbf{y}}} = \mathbf{I}$, then $\det(\mathrm{d}\mathbf{x}_a^\intercal/\mathrm{d}\mathbf{x}_a|_{\mathbf{y}=\bar{\mathbf{y}}}) = 1$ for all $a \in \{1, \ldots, N_\mathrm{a}\}$. Hence, $\det(\mathrm{d}\mathbf{x}^\intercal/\mathrm{d}\mathbf{x}|_{\mathbf{y}=\bar{\mathbf{y}}}) \neq 0$, so $\mathrm{d}\mathbf{x}^\intercal/\mathrm{d}\mathbf{x}|_{\mathbf{y}=\bar{\mathbf{y}}}$ is invertible.

We now obtain a more compact expression for the matrix of total effects of a mutant's phenotype on her phenotype in terms of partial derivatives. From Eq. S5.1.11, it follows that

$$\left.\frac{\delta\mathbf{x}^\intercal}{\delta\mathbf{x}}\right|_{\mathbf{y}=\bar{\mathbf{y}}} - \mathbf{I} = \left.\begin{pmatrix} \mathbf{0} & \frac{\delta\mathbf{x}_2^\intercal}{\delta\mathbf{x}_1} & \cdots & \mathbf{0} & \mathbf{0} \\ \mathbf{0} & \mathbf{0} & \cdots & \mathbf{0} & \mathbf{0} \\ \vdots & \vdots & \ddots & \vdots & \vdots \\ \mathbf{0} & \mathbf{0} & \cdots & \mathbf{0} & \frac{\delta\mathbf{x}_{N_\mathrm{a}}^\intercal}{\delta\mathbf{x}_{N_\mathrm{a}-1}} \\ \mathbf{0} & \mathbf{0} & \cdots & \mathbf{0} & \mathbf{0} \end{pmatrix}\right|_{\mathbf{y}=\bar{\mathbf{y}}},$$
(Eq. S5.1.16)

which is block 1-superdiagonal (i.e., only the entries in its first block super diagonal are non-zero). By definition of matrix power, we have that $(\delta\mathbf{x}^\intercal/\delta\mathbf{x}-\mathbf{I})^0 = \mathbf{I}$. Now, from Eq. S5.1.16, we have that

$$\frac{\delta\mathbf{x}^\intercal}{\delta\mathbf{x}} - \mathbf{I} = \left(\begin{cases} \frac{\delta\mathbf{x}_j^\intercal}{\delta\mathbf{x}_a} & \text{if } j = a+1 \\ \mathbf{0} & \text{otherwise} \end{cases}\right).$$

Using Eq. S5.1.16, taking the second power yields

$$\left(\frac{\delta\mathbf{x}^\intercal}{\delta\mathbf{x}} - \mathbf{I}\right)^2 = \left(\frac{\delta\mathbf{x}^\intercal}{\delta\mathbf{x}} - \mathbf{I}\right)\left(\frac{\delta\mathbf{x}^\intercal}{\delta\mathbf{x}} - \mathbf{I}\right) = \left(\begin{cases} \frac{\delta\mathbf{x}_{a+1}^\intercal}{\delta\mathbf{x}_a}\frac{\delta\mathbf{x}_j^\intercal}{\delta\mathbf{x}_{a+1}} & \text{if } j = a+2 \\ \mathbf{0} & \text{otherwise} \end{cases}\right),$$

which is block 2-superdiagonal. This suggests the inductive hypothesis that

$$\left(\frac{\delta\mathbf{x}^\intercal}{\delta\mathbf{x}} - \mathbf{I}\right)^i = \left(\begin{cases} \overset{\curvearrowleft}{\prod_{k=a}^{j-1}}\frac{\delta\mathbf{x}_{k+1}^\intercal}{\delta\mathbf{x}_k} & \text{if } j = a+i \\ \mathbf{0} & \text{otherwise} \end{cases}\right) \quad \text{(Eq. S5.1.17)}$$

holds for some $i \in \{0, 1, \ldots\}$, which is a block $i$-superdiagonal matrix. If this is the case, then we have that

$$\left(\frac{\delta\mathbf{x}^\intercal}{\delta\mathbf{x}} - \mathbf{I}\right)^{i+1} = \left(\frac{\delta\mathbf{x}^\intercal}{\delta\mathbf{x}} - \mathbf{I}\right)^i\left(\frac{\delta\mathbf{x}^\intercal}{\delta\mathbf{x}} - \mathbf{I}\right) = \left(\begin{cases} \overset{\curvearrowleft}{\prod_{k=a}^{a+i-1}}\frac{\delta\mathbf{x}_{k+1}^\intercal}{\delta\mathbf{x}_k}\frac{\delta\mathbf{x}_j^\intercal}{\delta\mathbf{x}_{a+i}} & \text{if } j = a+i+1 \\ \mathbf{0} & \text{otherwise} \end{cases}\right) = \left(\begin{cases} \overset{\curvearrowleft}{\prod_{k=a}^{j-1}}\frac{\delta\mathbf{x}_{k+1}^\intercal}{\delta\mathbf{x}_k} & \text{if } j = a+i+1 \\ \mathbf{0} & \text{otherwise} \end{cases}\right).$$

This proves by induction that Eq. S5.1.17 holds for every $i \in \{0, 1, \ldots\}$, which together with Eq. (12) proves that

$$\left(\frac{\delta\mathbf{x}^\intercal}{\delta\mathbf{x}} - \mathbf{I}\right)^i = \left(\begin{cases} \frac{\mathrm{d}\mathbf{x}_j^\intercal}{\mathrm{d}\mathbf{x}_a} & \text{if } j = a+i \\ \mathbf{0} & \text{otherwise} \end{cases}\right)$$

holds for all $i \in \{0, 1, \ldots, N_\mathrm{a}\}$. Evaluating this result at various $i$, note that

$$\left(\frac{\delta\mathbf{x}^\intercal}{\delta\mathbf{x}} - \mathbf{I}\right)^0 = \left(\begin{cases} \frac{\mathrm{d}\mathbf{x}_j^\intercal}{\mathrm{d}\mathbf{x}_a} & \text{if } j = a \\ \mathbf{0} & \text{otherwise} \end{cases}\right) = \left(\begin{cases} \mathbf{I} & \text{if } j = a \\ \mathbf{0} & \text{otherwise} \end{cases}\right)$$

is a block matrix of zeros except in its block main diagonal which coincides with the block main diagonal of Eq. S5.1.15. Similarly,

$$\left(\frac{\delta \mathbf{x}^\intercal}{\delta \mathbf{x}} - \mathbf{I}\right)^1 = \left(\begin{cases} \dfrac{\mathrm{d}\mathbf{x}_{a+1}^\intercal}{\mathrm{d}\mathbf{x}_a} & \text{if } j = a+1 \\ \mathbf{0} & \text{otherwise} \end{cases}\right)$$

is a block matrix of zeros except in its first block super diagonal which coincides with the first block super diagonal of Eq. S5.1.15. Indeed,

$$\left(\frac{\delta \mathbf{x}^\intercal}{\delta \mathbf{x}} - \mathbf{I}\right)^i = \left(\begin{cases} \dfrac{\mathrm{d}\mathbf{x}_{a+i}^\intercal}{\mathrm{d}\mathbf{x}_a} & \text{if } j = a+i \\ \mathbf{0} & \text{otherwise} \end{cases}\right)$$

is a block matrix of zeros except in its $i$-th block super diagonal which coincides with the $i$-th block super diagonal of Eq. S5.1.15 for all $i \in \{1, \dots, N_\mathrm{a} - 1\}$. Therefore, since any non-zero entry of the matrix $(\delta\mathbf{x}^\intercal/\delta\mathbf{x} - \mathbf{I})^i$ corresponds to a zero entry for the matrix $(\delta\mathbf{x}^\intercal/\delta\mathbf{x} - \mathbf{I})^j$ for any $i \neq j$ with $i, j \in \{0, \dots, N_\mathrm{a} - 1\}$, it follows that

$$\frac{\mathrm{d}\mathbf{x}^\intercal}{\mathrm{d}\mathbf{x}} = \sum_{i=0}^{N_\mathrm{a}-1} \left(\frac{\delta \mathbf{x}^\intercal}{\delta \mathbf{x}} - \mathbf{I}\right)^i. \qquad \text{(Eq. S5.1.18)}$$

From the geometric series of matrices we have that

$$\begin{aligned}\sum_{i=0}^{N_\mathrm{a}-1} \left(\frac{\delta \mathbf{x}^\intercal}{\delta \mathbf{x}} - \mathbf{I}\right)^i &= \left[\mathbf{I} - \left(\frac{\delta \mathbf{x}^\intercal}{\delta \mathbf{x}} - \mathbf{I}\right)\right]^{-1} \left[\mathbf{I} - \left(\frac{\delta \mathbf{x}^\intercal}{\delta \mathbf{x}} - \mathbf{I}\right)^{N_\mathrm{a}}\right] \\ &= \left(2\mathbf{I} - \frac{\delta \mathbf{x}^\intercal}{\delta \mathbf{x}}\right)^{-1}. \qquad \text{(Eq. S5.1.19)}\end{aligned}$$

The last equality follows because $\delta\mathbf{x}^\intercal/\delta\mathbf{x} - \mathbf{I}$ is strictly block triangular with block dimension $N_\mathrm{a}$ and so $\delta\mathbf{x}^\intercal/\delta\mathbf{x} - \mathbf{I}$ is nilpotent with index smaller than or equal to $N_\mathrm{a}$, which implies that $(\delta\mathbf{x}^\intercal/\delta\mathbf{x} - \mathbf{I})^{N_\mathrm{a}} = \mathbf{0}$. From Eq. S5.1.11, the matrix $2\mathbf{I} - \delta\mathbf{x}^\intercal/\delta\mathbf{x}$ is block upper triangular with only identity matrices in its block main diagonal, so all the eigenvalues of $2\mathbf{I} - \delta\mathbf{x}^\intercal/\delta\mathbf{x}$ equal one and the matrix is invertible; thus, the inverse matrix in Eq. S5.1.19 exists. Finally, using Eq. S5.1.19 in Eq. S5.1.18 yields Layer 4, Eq. S1, which is a compact expression for the matrix of total effects of a mutant's phenotype on her phenotype in terms of partial derivatives only, once Layer 3, Eq. S3 is used.

##### S5.1.3 Conclusion

**Form 1.** Using Eq. S5.1.7 and Layer 3, Eq. S1 for $\boldsymbol{\zeta} = \mathbf{x}$, we have that the total selection gradient of the phenotype is

$$\left.\frac{\mathrm{d}w}{\mathrm{d}\mathbf{x}}\right|_{\mathbf{y}=\bar{\mathbf{y}}} = \left.\left[\frac{\mathrm{d}\mathbf{x}^\intercal}{\mathrm{d}\mathbf{x}}\left(\frac{\partial w}{\partial \mathbf{x}} + \frac{\partial \boldsymbol{\epsilon}^\intercal}{\partial \mathbf{x}}\frac{\partial w}{\partial \boldsymbol{\epsilon}}\right)\right]\right|_{\mathbf{y}=\bar{\mathbf{y}}}.$$

Thus, using Layer 4, Eq. S9 yields the first line of Layer 4, Eq. S20.

**Form 2.** Using Eq. S5.1.7, the total selection gradient of the phenotype is given by the second line of Layer 4, Eq. S20.

**Form 3.** Using Eq. S5.1.7, Layer 3, Eq. S1 for $\boldsymbol{\zeta} = \mathbf{z}$, and Layer 4, Eq. S6, we have that the total selection gradient of the phenotype is given by the third line of Layer 4, Eq. S20, where the *total immediate selection gradient of the geno-phenotype* is

$$\left.\frac{\delta w}{\delta \mathbf{z}}\right|_{\mathbf{y}=\bar{\mathbf{y}}} \equiv \left.\begin{pmatrix} \dfrac{\delta w}{\delta \mathbf{x}} \\ \dfrac{\delta w}{\delta \mathbf{y}} \end{pmatrix}\right|_{\mathbf{y}=\bar{\mathbf{y}}} \in \mathbb{R}^{N_\mathrm{a}(N_\mathrm{p}+N_\mathrm{g})\times 1}. \qquad \text{(Eq. S5.1.20)}$$

**Form 4.** Finally, using the first line of Layer 4, Eq. S20 and Layer 4, Eq. S13, we obtain the fourth line of Layer 4, Eq. S20.

#### S5.2 Total selection gradient of the genotype

Here we derive the total selection gradient of the genotype following an analogous procedure to the one used in section S5.1 for the total selection gradient of the phenotype.

##### S5.2.1 Total selection gradient of the genotype in terms of direct fitness effects

As before, we start by considering the total selection gradient entry for the $i$-th genotypic trait at age $a$. By this, we mean the total selection gradient of a perturbation of $y_{ia}$ taken as initial condition of the developmental constraint (1) when applied at the ages $\{a, \dots, n\}$. Consequently, a genotypic perturbation at a given age does not affect the phenotype at earlier ages due to the arrow of developmental time. By letting $\zeta$ in Eq. S2.3.9 be $y_{ia}$, we have

$$\left.\frac{\mathrm{d}\lambda}{\mathrm{d}y_{ia}}\right|_{\mathbf{y}=\bar{\mathbf{y}}} = \left.\frac{\mathrm{d}w}{\mathrm{d}y_{ia}}\right|_{\mathbf{y}=\bar{\mathbf{y}}} = \sum_{j=1}^{N_\mathrm{a}} \left.\frac{\mathrm{d}w_j}{\mathrm{d}y_{ia}}\right|_{\mathbf{y}=\bar{\mathbf{y}}}. \qquad \text{(Eq. S5.2.1)}$$

The total derivatives of a mutant's relative fitness at age $j$ in Eq. S5.2.1 are with respect to the individual's genotypic trait at possibly another age $a$. We now seek to express such selection gradient entry in terms of partial derivatives only.

From Eq. S2.3.7, we have $w_j(\mathbf{z}_j, \mathbf{h}_j(\mathbf{z}_j, \bar{\mathbf{z}}, \tau), \bar{\mathbf{m}})$ with $\mathbf{z}_j = (\mathbf{x}_j; \mathbf{y}_j)$, so applying the chain rule, we obtain

$$\begin{aligned}\left.\frac{\mathrm{d}w_j}{\mathrm{d}y_{ia}}\right|_{\mathbf{y}=\bar{\mathbf{y}}} = \Bigg( &\sum_{k=1}^{N_\mathrm{p}} \frac{\partial w_j}{\partial x_{kj}}\frac{\mathrm{d}x_{kj}}{\mathrm{d}y_{ia}} + \sum_{k=1}^{N_\mathrm{g}} \frac{\partial w_j}{\partial y_{kj}}\frac{\mathrm{d}y_{kj}}{\mathrm{d}y_{ia}} \\ &+ \sum_{k=1}^{N_\mathrm{p}}\sum_{r=1}^{N_\mathrm{e}} \frac{\partial w_j}{\partial \epsilon_{rj}}\frac{\partial \epsilon_{rj}}{\partial x_{kj}}\frac{\mathrm{d}x_{kj}}{\mathrm{d}y_{ia}} \\ &+ \sum_{k=1}^{N_\mathrm{g}}\sum_{r=1}^{N_\mathrm{e}} \frac{\partial w_j}{\partial \epsilon_{rj}}\frac{\partial \epsilon_{rj}}{\partial y_{kj}}\frac{\mathrm{d}y_{kj}}{\mathrm{d}y_{ia}}\Bigg)\Bigg|_{\mathbf{y}=\bar{\mathbf{y}}}.\end{aligned}$$

Applying matrix calculus notation (Appendix A), this is

$$\begin{aligned}\left.\frac{\mathrm{d}w_j}{\mathrm{d}y_{ia}}\right|_{\mathbf{y}=\bar{\mathbf{y}}} = \Bigg( &\frac{\mathrm{d}\mathbf{x}_j^\intercal}{\mathrm{d}y_{ia}}\frac{\partial w_j}{\partial \mathbf{x}_j} + \frac{\mathrm{d}\mathbf{y}_j^\intercal}{\mathrm{d}y_{ia}}\frac{\partial w_j}{\partial \mathbf{y}_j} + \sum_{k=1}^{N_\mathrm{p}} \frac{\partial \boldsymbol{\epsilon}_j^\intercal}{\partial x_{kj}}\frac{\partial w_j}{\partial \boldsymbol{\epsilon}_j}\frac{\mathrm{d}x_{kj}}{\mathrm{d}y_{ia}} \\ &+ \sum_{k=1}^{N_\mathrm{g}} \frac{\partial \boldsymbol{\epsilon}_j^\intercal}{\partial y_{kj}}\frac{\partial w_j}{\partial \boldsymbol{\epsilon}_j}\frac{\mathrm{d}y_{kj}}{\mathrm{d}y_{ia}}\Bigg)\Bigg|_{\mathbf{y}=\bar{\mathbf{y}}}.\end{aligned}$$

Applying matrix calculus notation again yields

$$\left.\frac{\mathrm{d}w_j}{\mathrm{d}y_{ia}}\right|_{\mathbf{y}=\bar{\mathbf{y}}} = \left(\frac{\mathrm{d}\mathbf{x}_j^\intercal}{\mathrm{d}y_{ia}}\frac{\partial w_j}{\partial \mathbf{x}_j} + \frac{\mathrm{d}\mathbf{y}_j^\intercal}{\mathrm{d}y_{ia}}\frac{\partial w_j}{\partial \mathbf{y}_j} + \frac{\mathrm{d}\mathbf{x}_j^\intercal}{\mathrm{d}y_{ia}}\frac{\partial \boldsymbol{\epsilon}_j^\intercal}{\partial \mathbf{x}_j}\frac{\partial w_j}{\partial \boldsymbol{\epsilon}_j} + \left.\frac{\mathrm{d}\mathbf{y}_j^\intercal}{\mathrm{d}y_{ia}}\frac{\partial \boldsymbol{\epsilon}_j^\intercal}{\partial \mathbf{y}_j}\frac{\partial w_j}{\partial \boldsymbol{\epsilon}_j}\right)\right|_{\mathbf{y}=\bar{\mathbf{y}}}.$$

Factorizing, we have

$$\left.\frac{\mathrm{d}w_j}{\mathrm{d}y_{ia}}\right|_{\mathbf{y}=\bar{\mathbf{y}}} = \left[\frac{\mathrm{d}\mathbf{x}_j^\intercal}{\mathrm{d}y_{ia}}\left(\frac{\partial w_j}{\partial \mathbf{x}_j} + \frac{\partial \boldsymbol{\epsilon}_j^\intercal}{\partial \mathbf{x}_j}\frac{\partial w_j}{\partial \boldsymbol{\epsilon}_j}\right) + \left.\frac{\mathrm{d}\mathbf{y}_j^\intercal}{\mathrm{d}y_{ia}}\left(\frac{\partial w_j}{\partial \mathbf{y}_j} + \frac{\partial \boldsymbol{\epsilon}_j^\intercal}{\partial \mathbf{y}_j}\frac{\partial w_j}{\partial \boldsymbol{\epsilon}_j}\right)\right]\right|_{\mathbf{y}=\bar{\mathbf{y}}}. \quad \text{(Eq. S5.2.2)}$$

We now write Eq. S5.2.2 in terms of partial derivatives of lifetime fitness. Consider the *direct selection gradient of the genotype at age j*

$$\left.\frac{\partial w}{\partial \mathbf{y}_j}\right|_{\mathbf{y}=\bar{\mathbf{y}}} \equiv \left.\left(\frac{\partial w}{\partial y_{1j}},\ldots,\frac{\partial w}{\partial y_{N_\mathrm{g}j}}\right)^\intercal\right|_{\mathbf{y}=\bar{\mathbf{y}}} \in \mathbb{R}^{N_\mathrm{g}\times 1},$$

and the matrix of *direct effects of a mutant's genotype at age j on her environment at age j*

$$\left.\frac{\partial \boldsymbol{\epsilon}_j^\intercal}{\partial \mathbf{y}_j}\right|_{\mathbf{y}=\bar{\mathbf{y}}} \equiv \left.\begin{pmatrix} \frac{\partial \epsilon_{1j}}{\partial y_{1j}} & \cdots & \frac{\partial \epsilon_{N_\mathrm{e}j}}{\partial y_{1j}} \\ \vdots & \ddots & \vdots \\ \frac{\partial \epsilon_{1j}}{\partial y_{N_\mathrm{g}j}} & \cdots & \frac{\partial \epsilon_{N_\mathrm{e}j}^\intercal}{\partial y_{N_\mathrm{g}j}} \end{pmatrix}\right|_{\mathbf{y}=\bar{\mathbf{y}}} \in \mathbb{R}^{N_\mathrm{g}\times N_\mathrm{e}}.$$

Using Eq. S5.1.3 and Eq. S5.1.5 in Eq. S5.2.2 yields

$$\begin{aligned}\left.\frac{\mathrm{d}w_j}{\mathrm{d}y_{ia}}\right|_{\mathbf{y}=\bar{\mathbf{y}}} &= \left[\frac{\mathrm{d}\mathbf{x}_j^\intercal}{\mathrm{d}y_{ia}}\left(\frac{\partial w}{\partial \mathbf{x}_j} + \frac{\partial \boldsymbol{\epsilon}_j^\intercal}{\partial \mathbf{x}_j}\frac{\partial w}{\partial \boldsymbol{\epsilon}_j}\right) + \left.\frac{\mathrm{d}\mathbf{y}_j^\intercal}{\mathrm{d}y_{ia}}\left(\frac{\partial w}{\partial \mathbf{y}_j} + \frac{\partial \boldsymbol{\epsilon}_j^\intercal}{\partial \mathbf{y}_j}\frac{\partial w}{\partial \boldsymbol{\epsilon}_j}\right)\right]\right|_{\mathbf{y}=\bar{\mathbf{y}}} \\ &= \left.\left(\frac{\mathrm{d}\mathbf{x}_j^\intercal}{\mathrm{d}y_{ia}}\frac{\delta w}{\delta \mathbf{x}_j} + \frac{\mathrm{d}\mathbf{y}_j^\intercal}{\mathrm{d}y_{ia}}\frac{\delta w}{\delta \mathbf{y}_j}\right)\right|_{\mathbf{y}=\bar{\mathbf{y}}}, \quad \text{(Eq. S5.2.3)}\end{aligned}$$

where we use the *total immediate selection gradient of the genotype at age j* or, equivalently, the *total immediate effects of a mutant's genotype at age j on fitness*

$$\left.\frac{\delta w}{\delta \mathbf{y}_j}\right|_{\mathbf{y}=\bar{\mathbf{y}}} = \left.\left(\frac{\partial w}{\partial \mathbf{y}_j} + \frac{\partial \boldsymbol{\epsilon}_j^\intercal}{\partial \mathbf{y}_j}\frac{\partial w}{\partial \boldsymbol{\epsilon}_j}\right)\right|_{\mathbf{y}=\bar{\mathbf{y}}} \in \mathbb{R}^{N_\mathrm{g}\times 1}. \quad \text{(Eq. S5.2.4)}$$

Consider now the *total immediate selection gradient of the genotype* for all ages

$$\left.\frac{\delta w}{\delta \mathbf{y}}\right|_{\mathbf{y}=\bar{\mathbf{y}}} \equiv \left.\left(\frac{\delta w}{\delta \mathbf{y}_1};\cdots;\frac{\delta w}{\delta \mathbf{y}_{N_\mathrm{a}}}\right)\right|_{\mathbf{y}=\bar{\mathbf{y}}} \in \mathbb{R}^{N_\mathrm{a}N_\mathrm{g}\times 1}.$$

Using Layer 2, Eq. S2d, we have that

$$\frac{\partial \boldsymbol{\epsilon}^\intercal}{\partial \mathbf{y}}\frac{\partial w}{\partial \boldsymbol{\epsilon}} = \left(\sum_{k=1}^{N_\mathrm{a}}\frac{\partial \boldsymbol{\epsilon}_k^\intercal}{\partial \mathbf{y}_j}\frac{\partial w}{\partial \boldsymbol{\epsilon}_k}\right) = \left(\frac{\partial \boldsymbol{\epsilon}_j^\intercal}{\partial \mathbf{y}_j}\frac{\partial w}{\partial \boldsymbol{\epsilon}_j}\right) \quad \text{(Eq. S5.2.5)}$$

is a block column vector whose $j$-th entry is the rightmost term in Eq. S5.2.4. Thus, from Eq. S5.2.4, Layer 2, Eq. S1, and Eq. S5.2.5, it follows that the total immediate selection gradient of the genotype satisfies Layer 3, Eq. S1.

Now, we write the total selection gradient of $y_{ia}$ in terms of the total immediate selection gradient of the genotype. Substituting Eq. S5.2.3 in Eq. S5.2.1 yields

$$\begin{aligned}\left.\frac{\mathrm{d}w}{\mathrm{d}y_{ia}}\right|_{\mathbf{y}=\bar{\mathbf{y}}} &= \left.\sum_{j=1}^{N_\mathrm{a}}\left(\frac{\mathrm{d}\mathbf{x}_j^\intercal}{\mathrm{d}y_{ia}}\frac{\delta w}{\delta \mathbf{x}_j} + \frac{\mathrm{d}\mathbf{y}_j^\intercal}{\mathrm{d}y_{ia}}\frac{\delta w}{\delta \mathbf{y}_j}\right)\right|_{\mathbf{y}=\bar{\mathbf{y}}} \\ &= \left.\left(\frac{\mathrm{d}\mathbf{x}^\intercal}{\mathrm{d}y_{ia}}\frac{\delta w}{\delta \mathbf{x}} + \frac{\mathrm{d}\mathbf{y}^\intercal}{\mathrm{d}y_{ia}}\frac{\delta w}{\delta \mathbf{y}}\right)\right|_{\mathbf{y}=\bar{\mathbf{y}}},\end{aligned}$$

where we use the block row vectors

$$\frac{\mathrm{d}\mathbf{x}^\intercal}{\mathrm{d}y_{ia}} \equiv \left(\frac{\mathrm{d}\mathbf{x}_1^\intercal}{\mathrm{d}y_{ia}},\ldots,\frac{\mathrm{d}\mathbf{x}_{N_\mathrm{a}}^\intercal}{\mathrm{d}y_{ia}}\right) \in \mathbb{R}^{1\times N_\mathrm{a}N_\mathrm{p}}$$

$$\frac{\mathrm{d}\mathbf{y}^\intercal}{\mathrm{d}y_{ia}} \equiv \left(\frac{\mathrm{d}\mathbf{y}_1^\intercal}{\mathrm{d}y_{ia}},\ldots,\frac{\mathrm{d}\mathbf{y}_{N_\mathrm{a}}^\intercal}{\mathrm{d}y_{ia}}\right) \in \mathbb{R}^{1\times N_\mathrm{a}N_\mathrm{g}}.$$

Therefore, the total selection gradient of the genotype for all genotypic traits across all ages is

$$\left.\frac{\mathrm{d}w}{\mathrm{d}\mathbf{y}}\right|_{\mathbf{y}=\bar{\mathbf{y}}} = \left.\left(\frac{\mathrm{d}\mathbf{x}^\intercal}{\mathrm{d}\mathbf{y}}\frac{\delta w}{\delta \mathbf{x}} + \frac{\mathrm{d}\mathbf{y}^\intercal}{\mathrm{d}\mathbf{y}}\frac{\delta w}{\delta \mathbf{y}}\right)\right|_{\mathbf{y}=\bar{\mathbf{y}}} \in \mathbb{R}^{N_\mathrm{a}N_\mathrm{p}\times 1}, \quad \text{(Eq. S5.2.6)}$$

where we use the block matrix of *total effects of a mutant's genotype on her phenotype*

$$\left.\frac{\mathrm{d}\mathbf{x}^\intercal}{\mathrm{d}\mathbf{y}}\right|_{\mathbf{y}=\bar{\mathbf{y}}} \equiv \left.\begin{pmatrix} \frac{\mathrm{d}\mathbf{x}_1^\intercal}{\mathrm{d}\mathbf{y}_1} & \cdots & \frac{\mathrm{d}\mathbf{x}_{N_\mathrm{a}}^\intercal}{\partial \mathbf{y}_1} \\ \vdots & \ddots & \vdots \\ \frac{\mathrm{d}\mathbf{x}_1^\intercal}{\mathrm{d}\mathbf{y}_{N_\mathrm{a}}} & \cdots & \frac{\mathrm{d}\mathbf{x}_{N_\mathrm{a}}^\intercal}{\mathrm{d}\mathbf{y}_{N_\mathrm{a}}} \end{pmatrix}\right|_{\mathbf{y}=\bar{\mathbf{y}}} \in \mathbb{R}^{N_\mathrm{a}N_\mathrm{g}\times N_\mathrm{a}N_\mathrm{p}},$$

and the block matrix of *total effects of a mutant's genotype on her genotype*

$$\left.\frac{\mathrm{d}\mathbf{y}^\intercal}{\mathrm{d}\mathbf{y}}\right|_{\mathbf{y}=\bar{\mathbf{y}}} \equiv \left.\begin{pmatrix} \frac{\mathrm{d}\mathbf{y}_1^\intercal}{\mathrm{d}\mathbf{y}_1} & \cdots & \frac{\mathrm{d}\mathbf{y}_{N_\mathrm{a}}^\intercal}{\partial \mathbf{y}_1} \\ \vdots & \ddots & \vdots \\ \frac{\mathrm{d}\mathbf{y}_1^\intercal}{\mathrm{d}\mathbf{y}_{N_\mathrm{a}}} & \cdots & \frac{\mathrm{d}\mathbf{y}_{N_\mathrm{a}}^\intercal}{\mathrm{d}\mathbf{y}_{N_\mathrm{a}}} \end{pmatrix}\right|_{\mathbf{y}=\bar{\mathbf{y}}} \in \mathbb{R}^{N_\mathrm{a}N_\mathrm{g}\times N_\mathrm{a}N_\mathrm{g}}.$$

Eq. S5.2.6 is now in terms of partial derivatives of fitness, partial derivatives of the environment, total effects of a mutant's genotype on her phenotype, $\mathrm{d}\mathbf{x}^\intercal/\mathrm{d}\mathbf{y}$, and total effects of a mutant's genotype on her genotype, $\mathrm{d}\mathbf{y}^\intercal/\mathrm{d}\mathbf{y}$, once Layer 3, Eq. S1 is used. We now proceed to write $\mathrm{d}\mathbf{x}^\intercal/\mathrm{d}\mathbf{y}$ and $\mathrm{d}\mathbf{y}^\intercal/\mathrm{d}\mathbf{y}$ in terms of partial derivatives only.

##### S5.2.2 Matrix of total effects of a mutant's genotype on her phenotype and her genotype

From the developmental constraint (1) for the $k$-th phenotype at age $j \in \{2,\ldots,N_\mathrm{a}\}$ we have that $x_{kj} = g_{k,j-1}(\mathbf{z}_{j-1},\mathbf{h}_{j-1}(\mathbf{z}_{j-1},\bar{\mathbf{z}},\tau),\bar{\mathbf{z}})$, so using the chain rule we obtain

$$\left.\frac{\mathrm{d}x_{kj}}{\mathrm{d}y_{ia}}\right|_{\mathbf{y}=\bar{\mathbf{y}}} = \left(\sum_{l=1}^{N_\mathrm{p}}\frac{\partial g_{k,j-1}}{\partial x_{l,j-1}}\frac{\mathrm{d}x_{l,j-1}}{\mathrm{d}y_{ia}} + \sum_{l=1}^{N_\mathrm{g}}\frac{\partial g_{k,j-1}}{\partial y_{l,j-1}}\frac{\mathrm{d}y_{l,j-1}}{\mathrm{d}y_{ia}}\right.$$

$$+\sum_{l=1}^{N_\mathrm{p}}\sum_{r=1}^{N_\mathrm{e}}\frac{\partial g_{k,j-1}}{\partial \epsilon_{r,j-1}}\frac{\partial \epsilon_{r,j-1}}{\partial x_{l,j-1}}\frac{\mathrm{d}x_{l,j-1}}{\mathrm{d}y_{ia}}$$
$$+\sum_{l=1}^{N_\mathrm{g}}\sum_{r=1}^{N_\mathrm{e}}\frac{\partial g_{k,j-1}}{\partial \epsilon_{r,j-1}}\frac{\partial \epsilon_{r,j-1}}{\partial y_{l,j-1}}\frac{\mathrm{d}y_{l,j-1}}{\mathrm{d}y_{ia}}\Bigg)\Bigg|_{\mathbf{y}=\bar{\mathbf{y}}}.$$

Applying matrix calculus notation (Appendix A), this is

$$\frac{\mathrm{d}x_{kj}}{\mathrm{d}y_{ia}}\bigg|_{\mathbf{y}=\bar{\mathbf{y}}}=\Bigg(\frac{\mathrm{d}\mathbf{x}_{j-1}^\intercal}{\mathrm{d}y_{ia}}\frac{\partial g_{k,j-1}}{\partial \mathbf{x}_{j-1}}+\frac{\mathrm{d}\mathbf{y}_{j-1}^\intercal}{\mathrm{d}y_{ia}}\frac{\partial g_{k,j-1}}{\partial \mathbf{y}_{j-1}}$$
$$+\sum_{l=1}^{N_\mathrm{p}}\frac{\partial \boldsymbol{\epsilon}_{j-1}^\intercal}{\partial x_{l,j-1}}\frac{\partial g_{k,j-1}}{\partial \boldsymbol{\epsilon}_{j-1}}\frac{\mathrm{d}x_{l,j-1}}{\mathrm{d}y_{ia}}$$
$$+\sum_{l=1}^{N_\mathrm{g}}\frac{\partial \boldsymbol{\epsilon}_{j-1}^\intercal}{\partial y_{l,j-1}}\frac{\partial g_{k,j-1}}{\partial \boldsymbol{\epsilon}_{j-1}}\frac{\mathrm{d}y_{l,j-1}}{\mathrm{d}y_{ia}}\Bigg)\Bigg|_{\mathbf{y}=\bar{\mathbf{y}}}.$$

Applying matrix calculus notation again yields

$$\frac{\mathrm{d}x_{kj}}{\mathrm{d}y_{ia}}\bigg|_{\mathbf{y}=\bar{\mathbf{y}}}=\Bigg(\frac{\mathrm{d}\mathbf{x}_{j-1}^\intercal}{\mathrm{d}y_{ia}}\frac{\partial g_{k,j-1}}{\partial \mathbf{x}_{j-1}}+\frac{\mathrm{d}\mathbf{y}_{j-1}^\intercal}{\mathrm{d}y_{ia}}\frac{\partial g_{k,j-1}}{\partial \mathbf{y}_{j-1}}$$
$$+\frac{\mathrm{d}\mathbf{x}_{j-1}^\intercal}{\mathrm{d}y_{ia}}\frac{\partial \boldsymbol{\epsilon}_{j-1}^\intercal}{\partial \mathbf{x}_{j-1}}\frac{\partial g_{k,j-1}}{\partial \boldsymbol{\epsilon}_{j-1}}$$
$$+\frac{\mathrm{d}\mathbf{y}_{j-1}^\intercal}{\mathrm{d}y_{ia}}\frac{\partial \boldsymbol{\epsilon}_{j-1}^\intercal}{\partial \mathbf{y}_{j-1}}\frac{\partial g_{k,j-1}}{\partial \boldsymbol{\epsilon}_{j-1}}\Bigg)\Bigg|_{\mathbf{y}=\bar{\mathbf{y}}}.$$

Factorizing, we have

$$\frac{\mathrm{d}x_{kj}}{\mathrm{d}y_{ia}}\bigg|_{\mathbf{y}=\bar{\mathbf{y}}}=\Bigg[\frac{\mathrm{d}\mathbf{x}_{j-1}^\intercal}{\mathrm{d}y_{ia}}\left(\frac{\partial g_{k,j-1}}{\partial \mathbf{x}_{j-1}}+\frac{\partial \boldsymbol{\epsilon}_{j-1}^\intercal}{\partial \mathbf{x}_{j-1}}\frac{\partial g_{k,j-1}}{\partial \boldsymbol{\epsilon}_{j-1}}\right)$$
$$+\frac{\mathrm{d}\mathbf{y}_{j-1}^\intercal}{\mathrm{d}y_{ia}}\left(\frac{\partial g_{k,j-1}}{\partial \mathbf{y}_{j-1}}+\frac{\partial \boldsymbol{\epsilon}_{j-1}^\intercal}{\partial \mathbf{y}_{j-1}}\frac{\partial g_{k,j-1}}{\partial \boldsymbol{\epsilon}_{j-1}}\right)\Bigg]\Bigg|_{\mathbf{y}=\bar{\mathbf{y}}}.$$

Rewriting $g_{k,j-1}$ as $x_{kj}$ yields

$$\frac{\mathrm{d}x_{kj}}{\mathrm{d}y_{ia}}\bigg|_{\mathbf{y}=\bar{\mathbf{y}}}=\Bigg[\frac{\mathrm{d}\mathbf{x}_{j-1}^\intercal}{\mathrm{d}y_{ia}}\left(\frac{\partial x_{kj}}{\partial \mathbf{x}_{j-1}}+\frac{\partial \boldsymbol{\epsilon}_{j-1}^\intercal}{\partial \mathbf{x}_{j-1}}\frac{\partial x_{kj}}{\partial \boldsymbol{\epsilon}_{j-1}}\right)$$
$$+\frac{\mathrm{d}\mathbf{y}_{j-1}^\intercal}{\mathrm{d}y_{ia}}\left(\frac{\partial x_{kj}}{\partial \mathbf{y}_{j-1}}+\frac{\partial \boldsymbol{\epsilon}_{j-1}^\intercal}{\partial \mathbf{y}_{j-1}}\frac{\partial x_{kj}}{\partial \boldsymbol{\epsilon}_{j-1}}\right)\Bigg]\Bigg|_{\mathbf{y}=\bar{\mathbf{y}}}.$$

Hence,

$$\frac{\mathrm{d}\mathbf{x}_j^\intercal}{\mathrm{d}y_{ia}}\bigg|_{\mathbf{y}=\bar{\mathbf{y}}}=\Bigg[\frac{\mathrm{d}\mathbf{x}_{j-1}^\intercal}{\mathrm{d}y_{ia}}\left(\frac{\partial \mathbf{x}_j^\intercal}{\partial \mathbf{x}_{j-1}}+\frac{\partial \boldsymbol{\epsilon}_{j-1}^\intercal}{\partial \mathbf{x}_{j-1}}\frac{\partial \mathbf{x}_j^\intercal}{\partial \boldsymbol{\epsilon}_{j-1}}\right)$$
$$+\frac{\mathrm{d}\mathbf{y}_{j-1}^\intercal}{\mathrm{d}y_{ia}}\left(\frac{\partial \mathbf{x}_j^\intercal}{\partial \mathbf{y}_{j-1}}+\frac{\partial \boldsymbol{\epsilon}_{j-1}^\intercal}{\partial \mathbf{y}_{j-1}}\frac{\partial \mathbf{x}_j^\intercal}{\partial \boldsymbol{\epsilon}_{j-1}}\right)\Bigg]\Bigg|_{\mathbf{y}=\bar{\mathbf{y}}},\tag{Eq. S5.2.7}$$

where we use the matrix of *direct effects of a mutant's genotypic trait values at age* $j$ *on her phenotype at age* $j+1$

$$\frac{\partial \mathbf{x}_{j+1}^\intercal}{\partial \mathbf{y}_j}\bigg|_{\mathbf{y}=\bar{\mathbf{y}}}\equiv\begin{pmatrix}\frac{\partial x_{1,j+1}}{\partial y_{1j}} & \cdots & \frac{\partial x_{N_\mathrm{p},j+1}}{\partial y_{1j}}\\ \vdots & \ddots & \vdots\\ \frac{\partial x_{1,j+1}}{\partial y_{N_\mathrm{g}j}} & \cdots & \frac{\partial x_{N_\mathrm{p},j+1}}{\partial y_{N_\mathrm{g}j}}\end{pmatrix}\Bigg|_{\mathbf{y}=\bar{\mathbf{y}}}\in\mathbb{R}^{N_\mathrm{g}\times N_\mathrm{p}}.$$

We can write Eq. S5.2.7 more succinctly as

$$\frac{\mathrm{d}\mathbf{x}_j^\intercal}{\mathrm{d}y_{ia}}\bigg|_{\mathbf{y}=\bar{\mathbf{y}}}=\left(\frac{\mathrm{d}\mathbf{x}_{j-1}^\intercal}{\mathrm{d}y_{ia}}\frac{\delta \mathbf{x}_j^\intercal}{\delta \mathbf{x}_{j-1}}+\frac{\mathrm{d}\mathbf{y}_{j-1}^\intercal}{\mathrm{d}y_{ia}}\frac{\delta \mathbf{x}_j^\intercal}{\delta \mathbf{y}_{j-1}}\right)\Bigg|_{\mathbf{y}=\bar{\mathbf{y}}},\tag{Eq. S5.2.8}$$

where we use the matrix of *total immediate effects of a mutant's genotypic trait values at age* $j$ *on her phenotype at age* $j+1$

$$\frac{\delta \mathbf{x}_{j+1}^\intercal}{\delta \mathbf{y}_j}\bigg|_{\mathbf{y}=\bar{\mathbf{y}}}=\left(\frac{\partial \mathbf{x}_{j+1}^\intercal}{\partial \mathbf{y}_j}+\frac{\partial \boldsymbol{\epsilon}_j^\intercal}{\partial \mathbf{y}_j}\frac{\partial \mathbf{x}_{j+1}^\intercal}{\partial \boldsymbol{\epsilon}_j}\right)\Bigg|_{\mathbf{y}=\bar{\mathbf{y}}}\in\mathbb{R}^{N_\mathrm{g}\times N_\mathrm{p}}.\tag{Eq. S5.2.9}$$

We also define the corresponding matrix across all ages. Specifically, the block matrix of *total immediate effects of a mutant's genotype on her phenotype* is

$$\frac{\delta \mathbf{x}^\intercal}{\delta \mathbf{y}}\bigg|_{\mathbf{y}=\bar{\mathbf{y}}}\equiv\begin{pmatrix}\frac{\delta \mathbf{x}_1^\intercal}{\delta \mathbf{y}_1} & \cdots & \frac{\delta \mathbf{x}_{N_\mathrm{a}}^\intercal}{\delta \mathbf{y}_1}\\ \vdots & \ddots & \vdots\\ \frac{\delta \mathbf{x}_1^\intercal}{\delta \mathbf{y}_{N_\mathrm{a}}} & \cdots & \frac{\delta \mathbf{x}_{N_\mathrm{a}}^\intercal}{\delta \mathbf{y}_{N_\mathrm{a}}}\end{pmatrix}\Bigg|_{\mathbf{y}=\bar{\mathbf{y}}}$$
$$=\begin{pmatrix}\mathbf{0} & \frac{\delta \mathbf{x}_2^\intercal}{\delta \mathbf{y}_1} & \cdots & \mathbf{0} & \mathbf{0}\\ \mathbf{0} & \mathbf{0} & \cdots & \mathbf{0} & \mathbf{0}\\ \vdots & \vdots & \ddots & \vdots & \vdots\\ \mathbf{0} & \mathbf{0} & \cdots & \mathbf{0} & \frac{\delta \mathbf{x}_{N_\mathrm{a}}^\intercal}{\delta \mathbf{y}_{N_\mathrm{a}-1}}\\ \mathbf{0} & \mathbf{0} & \cdots & \mathbf{0} & \mathbf{0}\end{pmatrix}\Bigg|_{\mathbf{y}=\bar{\mathbf{y}}}\tag{Eq. S5.2.10}$$
$$\in\mathbb{R}^{N_\mathrm{a}N_\mathrm{g}\times N_\mathrm{a}N_\mathrm{p}}.$$

The equality in Eq. S5.2.10 follows because the total immediate effects of a mutant's genotypic trait values on her phenotype are only non-zero at the next age (from the developmental constraint in Eq. 1). Using Layer 2, Eq. S2d and Layer 2, Eq. S2c, we have that

$$\frac{\partial \boldsymbol{\epsilon}^\intercal}{\partial \mathbf{y}}\frac{\partial \mathbf{x}^\intercal}{\partial \boldsymbol{\epsilon}}=\left(\sum_{k=1}^{N_\mathrm{a}}\frac{\partial \boldsymbol{\epsilon}_k^\intercal}{\partial \mathbf{y}_a}\frac{\partial \mathbf{x}_j^\intercal}{\partial \boldsymbol{\epsilon}_k}\right)=\left(\begin{cases}\frac{\partial \boldsymbol{\epsilon}_a^\intercal}{\partial \mathbf{y}_a}\frac{\partial \mathbf{x}_j^\intercal}{\partial \boldsymbol{\epsilon}_a} & \text{for } j=a+1\\ \mathbf{0} & \text{for } j\neq a+1\end{cases}\right),\tag{Eq. S5.2.11}$$

which equals the rightmost term in Eq. S5.2.9 for $j=a+1$. Thus, from Eq. S5.2.9–Eq. S5.2.11, it follows that the block matrix of total immediate effects of a mutant's genotype on her phenotype satisfies Layer 3, Eq. S3.

Eq. S5.2.8 gives the matrix of total effects of a mutant's $i$-th genotypic trait value at age $a$ on her phenotype at age $j$. Then, it follows that the matrix of total effects of a mutant's genotypic traits for all genotypic traits at age $a$ on her phenotype at age $j$ is

$$\frac{\mathrm{d}\mathbf{x}_j^\intercal}{\mathrm{d}\mathbf{y}_a}\bigg|_{\mathbf{y}=\bar{\mathbf{y}}}=\left(\frac{\mathrm{d}\mathbf{x}_{j-1}^\intercal}{\mathrm{d}\mathbf{y}_a}\frac{\delta \mathbf{x}_j^\intercal}{\delta \mathbf{x}_{j-1}}+\frac{\mathrm{d}\mathbf{y}_{j-1}^\intercal}{\mathrm{d}\mathbf{y}_a}\frac{\delta \mathbf{x}_j^\intercal}{\delta \mathbf{y}_{j-1}}\right)\Bigg|_{\mathbf{y}=\bar{\mathbf{y}}}.\tag{Eq. S5.2.12}$$

Eq. S5.2.12 is a recurrence equation for $\mathrm{d}\mathbf{x}_j^\mathsf{T}/\mathrm{d}\mathbf{y}_a$ over age $j \in \{2,\ldots,N_\mathrm{a}\}$. Since a given entry of the operator $\mathrm{d}/\mathrm{d}\mathbf{y}$ takes the total derivative with respect to a given $y_{ia}$ while keeping all the other genotypic traits constant and genotypic traits are developmentally independent, a perturbation of an individual's genotypic trait value t a given age does not affect any other of the individual's genotypic trait value at the same or other ages (i.e., $\mathrm{d}\mathbf{y}_a^\mathsf{T}/\mathrm{d}\mathbf{y}_a = \mathbf{I}$ and $\mathrm{d}\mathbf{y}_j^\mathsf{T}/\mathrm{d}\mathbf{y}_a = \mathbf{0}$ for $j \neq a$). Thus, the matrix of total effects of a mutant's genotype on her genotype is

$$\frac{\mathrm{d}\mathbf{y}^\mathsf{T}}{\mathrm{d}\mathbf{y}} = \begin{pmatrix} \frac{\mathrm{d}\mathbf{y}_1^\mathsf{T}}{\mathrm{d}\mathbf{y}_1} & \cdots & \frac{\mathrm{d}\mathbf{y}_{N_\mathrm{a}}^\mathsf{T}}{\mathrm{d}\mathbf{y}_1} \\ \vdots & \ddots & \vdots \\ \frac{\mathrm{d}\mathbf{y}_1^\mathsf{T}}{\mathrm{d}\mathbf{y}_{N_\mathrm{a}}} & \cdots & \frac{\mathrm{d}\mathbf{y}_{N_\mathrm{a}}^\mathsf{T}}{\mathrm{d}\mathbf{y}_{N_\mathrm{a}}} \end{pmatrix} = \begin{pmatrix} \mathbf{I} & \mathbf{0} & \cdots & \mathbf{0} & \mathbf{0} \\ \mathbf{0} & \mathbf{I} & \cdots & \mathbf{0} & \mathbf{0} \\ \vdots & \vdots & \ddots & \vdots & \vdots \\ \mathbf{0} & \mathbf{0} & \cdots & \mathbf{I} & \mathbf{0} \\ \mathbf{0} & \mathbf{0} & \cdots & \mathbf{0} & \mathbf{I} \end{pmatrix} = \mathbf{I} \in \mathbb{R}^{N_\mathrm{a}N_\mathrm{g} \times N_\mathrm{a}N_\mathrm{g}}. \tag{Eq. S5.2.13}$$

Moreover, because of the arrow of developmental time (due to the developmental constraint in Eq. 1), perturbations in an individual's late genotypic trait values do not affect the individual's early phenotype (i.e., $\mathrm{d}\mathbf{x}_j^\mathsf{T}/\mathrm{d}\mathbf{y}_a = \mathbf{0}$ for $j < a$ and $j \in \{1,\ldots,N_\mathrm{a}-1\}$)[2]. Additionally, from the arrow of developmental time (Eq. 1), a perturbation in an individual's genotypic trait values at a given age does not affect any of the individual's phenotypes at the *same* age (i.e., $\mathrm{d}\mathbf{x}_j^\mathsf{T}/\mathrm{d}\mathbf{y}_a = \mathbf{0}$ for $j = a$). Consequently, Eq. S5.2.12 for $j \in \{1,\ldots,N_\mathrm{a}\}$ reduces to

$$\left.\frac{\mathrm{d}\mathbf{x}_j^\mathsf{T}}{\mathrm{d}\mathbf{y}_a}\right|_{\mathbf{y}=\bar{\mathbf{y}}} = \begin{cases} \left.\left(\frac{\mathrm{d}\mathbf{x}_{j-1}^\mathsf{T}}{\mathrm{d}\mathbf{y}_a}\frac{\delta\mathbf{x}_j^\mathsf{T}}{\delta\mathbf{x}_{j-1}} + \underbrace{\frac{\mathrm{d}\mathbf{y}_{j-1}^\mathsf{T}}{\mathrm{d}\mathbf{y}_a}}_{\mathbf{0},\text{ by Eq. S5.2.13}}\frac{\delta\mathbf{x}_j^\mathsf{T}}{\delta\mathbf{y}_{j-1}}\right)\right|_{\mathbf{y}=\bar{\mathbf{y}}} & \text{for } j-1>a \\ \left.\left(\underbrace{\frac{\mathrm{d}\mathbf{x}_{j-1}^\mathsf{T}}{\mathrm{d}\mathbf{y}_a}}_{\mathbf{0},\text{ by (1)}}\frac{\delta\mathbf{x}_j^\mathsf{T}}{\delta\mathbf{x}_{j-1}} + \underbrace{\frac{\mathrm{d}\mathbf{y}_{j-1}^\mathsf{T}}{\mathrm{d}\mathbf{y}_a}}_{\mathbf{I},\text{ by Eq. S5.2.13}}\frac{\delta\mathbf{x}_j^\mathsf{T}}{\delta\mathbf{y}_{j-1}}\right)\right|_{\mathbf{y}=\bar{\mathbf{y}}} & \text{for } j-1=a \\ \left.\left(\underbrace{\frac{\mathrm{d}\mathbf{x}_{j-1}^\mathsf{T}}{\mathrm{d}\mathbf{y}_a}}_{\mathbf{0},\text{ by (1)}}\frac{\delta\mathbf{x}_j^\mathsf{T}}{\delta\mathbf{x}_{j-1}} + \underbrace{\frac{\mathrm{d}\mathbf{y}_{j-1}^\mathsf{T}}{\mathrm{d}\mathbf{y}_a}}_{\mathbf{0},\text{ by Eq. S5.2.13}}\frac{\delta\mathbf{x}_j^\mathsf{T}}{\delta\mathbf{y}_{j-1}}\right)\right|_{\mathbf{y}=\bar{\mathbf{y}}} & \text{for } j-1<a. \end{cases}$$

That is,

$$\left.\frac{\mathrm{d}\mathbf{x}_j^\mathsf{T}}{\mathrm{d}\mathbf{y}_a}\right|_{\mathbf{y}=\bar{\mathbf{y}}} = \begin{cases} \left.\left(\frac{\mathrm{d}\mathbf{x}_{j-1}^\mathsf{T}}{\mathrm{d}\mathbf{y}_a}\frac{\delta\mathbf{x}_j^\mathsf{T}}{\delta\mathbf{x}_{j-1}}\right)\right|_{\mathbf{y}=\bar{\mathbf{y}}} & \text{for } j-1>a \\ \left.\frac{\delta\mathbf{x}_j^\mathsf{T}}{\delta\mathbf{y}_{j-1}}\right|_{\mathbf{y}=\bar{\mathbf{y}}} & \text{for } j-1=a \\ \mathbf{0} & \text{for } j-1<a. \end{cases}$$

[2]Again, we take the derivative $\mathrm{d}\mathbf{x}_j^\mathsf{T}/\mathrm{d}y_{ia}$ as referring to the effect on $\mathbf{x}_j^\mathsf{T}$ of a perturbation of the initial condition $\mathbf{y}_a$ of the difference equation (1) applied at the ages $\{a,\ldots,n\}$. Hence, if $j < a$, $\mathbf{x}_j^\mathsf{T}$ is unmodified by a change in the initial condition of (1) applied at the ages $\{a,\ldots,n\}$.

Expanding this recurrence yields

$$\left.\frac{\mathrm{d}\mathbf{x}_j^\mathsf{T}}{\mathrm{d}\mathbf{y}_a}\right|_{\mathbf{y}=\bar{\mathbf{y}}} = \begin{cases} \left.\left(\frac{\mathrm{d}\mathbf{x}_{a+1}^\mathsf{T}}{\mathrm{d}\mathbf{y}_a}\frac{\delta\mathbf{x}_{a+2}^\mathsf{T}}{\delta\mathbf{x}_{a+1}}\cdots\frac{\delta\mathbf{x}_j^\mathsf{T}}{\delta\mathbf{x}_{j-1}}\right)\right|_{\mathbf{y}=\bar{\mathbf{y}}} & \text{for } j-1>a \\ \left.\frac{\delta\mathbf{x}_{a+1}^\mathsf{T}}{\delta\mathbf{y}_a}\right|_{\mathbf{y}=\bar{\mathbf{y}}} & \text{for } j-1=a \\ \mathbf{0} & \text{for } j-1<a. \end{cases} \tag{Eq. S5.2.14}$$

Evaluating Eq. S5.2.14 at $j = a+1$ yields

$$\left.\frac{\mathrm{d}\mathbf{x}_{a+1}^\mathsf{T}}{\mathrm{d}\mathbf{y}_a}\right|_{\mathbf{y}=\bar{\mathbf{y}}} = \left.\frac{\delta\mathbf{x}_{a+1}^\mathsf{T}}{\delta\mathbf{y}_a}\right|_{\mathbf{y}=\bar{\mathbf{y}}},$$

which substituted back in the top line of Eq. S5.2.14 yields

$$\left.\frac{\mathrm{d}\mathbf{x}_j^\mathsf{T}}{\mathrm{d}\mathbf{y}_a}\right|_{\mathbf{y}=\bar{\mathbf{y}}} = \begin{cases} \left.\left(\frac{\delta\mathbf{x}_{a+1}^\mathsf{T}}{\delta\mathbf{y}_a}\frac{\delta\mathbf{x}_{a+2}^\mathsf{T}}{\delta\mathbf{x}_{a+1}}\cdots\frac{\delta\mathbf{x}_j^\mathsf{T}}{\delta\mathbf{x}_{j-1}}\right)\right|_{\mathbf{y}=\bar{\mathbf{y}}} & \text{for } j-1>a \\ \left.\frac{\delta\mathbf{x}_{a+1}^\mathsf{T}}{\delta\mathbf{y}_a}\right|_{\mathbf{y}=\bar{\mathbf{y}}} & \text{for } j-1=a \\ \mathbf{0} & \text{for } j-1<a. \end{cases} \tag{Eq. S5.2.15}$$

Hence, the block matrix of *total effects of a mutant's genotype on her phenotype* is

$$\left.\frac{\mathrm{d}\mathbf{x}^\mathsf{T}}{\mathrm{d}\mathbf{y}}\right|_{\mathbf{y}=\bar{\mathbf{y}}} = \left.\begin{pmatrix} \frac{\mathrm{d}\mathbf{x}_1^\mathsf{T}}{\mathrm{d}\mathbf{y}_1} & \cdots & \frac{\mathrm{d}\mathbf{x}_{N_\mathrm{a}}^\mathsf{T}}{\mathrm{d}\mathbf{y}_1} \\ \vdots & \ddots & \vdots \\ \frac{\mathrm{d}\mathbf{x}_1^\mathsf{T}}{\mathrm{d}\mathbf{y}_{N_\mathrm{a}}} & \cdots & \frac{\mathrm{d}\mathbf{x}_{N_\mathrm{a}}^\mathsf{T}}{\mathrm{d}\mathbf{y}_{N_\mathrm{a}}} \end{pmatrix}\right|_{\mathbf{y}=\bar{\mathbf{y}}} = \left.\begin{pmatrix} \mathbf{0} & \frac{\mathrm{d}\mathbf{x}_2^\mathsf{T}}{\mathrm{d}\mathbf{y}_1} & \cdots & \frac{\mathrm{d}\mathbf{x}_{N_\mathrm{a}-1}^\mathsf{T}}{\mathrm{d}\mathbf{y}_1} & \frac{\mathrm{d}\mathbf{x}_{N_\mathrm{a}}^\mathsf{T}}{\mathrm{d}\mathbf{y}_1} \\ \mathbf{0} & \mathbf{0} & \cdots & \frac{\mathrm{d}\mathbf{x}_{N_\mathrm{a}-1}^\mathsf{T}}{\mathrm{d}\mathbf{y}_2} & \frac{\mathrm{d}\mathbf{x}_{N_\mathrm{a}}^\mathsf{T}}{\mathrm{d}\mathbf{y}_2} \\ \vdots & \vdots & \ddots & \vdots & \vdots \\ \mathbf{0} & \mathbf{0} & \cdots & \mathbf{0} & \frac{\mathrm{d}\mathbf{x}_{N_\mathrm{a}}^\mathsf{T}}{\mathrm{d}\mathbf{y}_{N_\mathrm{a}-1}} \\ \mathbf{0} & \mathbf{0} & \cdots & \mathbf{0} & \mathbf{0} \end{pmatrix}\right|_{\mathbf{y}=\bar{\mathbf{y}}} \in \mathbb{R}^{N_\mathrm{a}N_\mathrm{g} \times N_\mathrm{a}N_\mathrm{p}}, \tag{Eq. S5.2.16}$$

whose $aj$-th block entry is given by

$$\frac{\mathrm{d}\mathbf{x}_j^\mathsf{T}}{\mathrm{d}\mathbf{y}_a} = \begin{cases} \frac{\delta\mathbf{x}_{a+1}^\mathsf{T}}{\delta\mathbf{y}_a}\frac{\mathrm{d}\mathbf{x}_j^\mathsf{T}}{\mathrm{d}\mathbf{x}_{a+1}} & \text{for } j>a \\ \mathbf{0} & \text{for } j \leq a \end{cases} = \begin{cases} \frac{\delta\mathbf{x}_{a+1}^\mathsf{T}}{\delta\mathbf{y}_a}\overset{\curvearrowright}{\prod_{k=a+1}^{j-1}}\frac{\delta\mathbf{x}_{k+1}^\mathsf{T}}{\delta\mathbf{x}_k} & \text{for } j>a \\ \mathbf{0} & \text{for } j \leq a \end{cases} = \begin{cases} \frac{\delta\mathbf{x}_{a+1}^\mathsf{T}}{\delta\mathbf{y}_a}\frac{\delta\mathbf{x}_{a+2}^\mathsf{T}}{\delta\mathbf{x}_{a+1}}\cdots\frac{\delta\mathbf{x}_j^\mathsf{T}}{\delta\mathbf{x}_{j-1}} & \text{for } j>a \\ \mathbf{0} & \text{for } j \leq a, \end{cases} \tag{Eq. S5.2.17}$$

where we use Eq. (12) and adopt the empty-product convention that

$$\frac{\mathrm{d}\mathbf{x}_{a+1}^{\intercal}}{\mathrm{d}\mathbf{x}_{a+1}} = \prod_{k=a+1}^{\overset{\frown}{a}} \frac{\delta \mathbf{x}_{k+1}^{\intercal}}{\delta \mathbf{x}_k} = \mathbf{I}.$$

Eq. S5.2.16 and Eq. S5.2.17 write the matrix of total effects of a mutant's genotype on her phenotype in terms of partial derivatives, given Eq. S5.2.9, as we sought.

We now obtain a more compact expression for the matrix of total effects of a mutant's genotype on her phenotype in terms of partial derivatives. To do this, we note a relationship between the matrix of total effects of a mutant's genotype on her phenotype with the matrix of total effects of a mutant's phenotype on her phenotype. Note that the $aj$-th block entry of $(\delta\mathbf{x}^{\intercal}/\delta\mathbf{y})(\mathrm{d}\mathbf{x}^{\intercal}/\mathrm{d}\mathbf{x})$ is

$$\begin{aligned}\left(\frac{\delta \mathbf{x}^{\intercal}}{\delta \mathbf{y}} \frac{\mathrm{d}\mathbf{x}^{\intercal}}{\mathrm{d}\mathbf{x}}\right)_{aj} &= \sum_{k=1}^{N_{\mathrm{a}}} \frac{\delta \mathbf{x}_k^{\intercal}}{\delta \mathbf{y}_a} \frac{\mathrm{d}\mathbf{x}_j^{\intercal}}{\mathrm{d}\mathbf{x}_k} \\ &= \frac{\delta \mathbf{x}_{a+1}^{\intercal}}{\delta \mathbf{y}_a} \frac{\mathrm{d}\mathbf{x}_j^{\intercal}}{\mathrm{d}\mathbf{x}_{a+1}} \\ &= \frac{\mathrm{d}\mathbf{x}_j^{\intercal}}{\mathrm{d}\mathbf{y}_a},\end{aligned}$$

where we use Eq. S5.2.10 in the second equality and Eq. S5.2.17 in the third equality, noting that $\mathrm{d}\mathbf{x}_j^{\intercal}/\mathrm{d}\mathbf{x}_{a+1} = \mathbf{0}$ and $\mathrm{d}\mathbf{x}_j^{\intercal}/\mathrm{d}\mathbf{y}_a = \mathbf{0}$ for $j \leq a$. Hence, Layer 4, Eq. S2 follows, which is a compact expression for the matrix of total effects of a mutant's genotype on her phenotype in terms of partial derivatives only, once Layer 4, Eq. S1 and Layer 3, Eq. S3 are used.

##### S5.2.3 Conclusion

**Form 1.** Using Eq. S5.2.6, Eq. S5.2.13, and Layer 3, Eq. S1 for $\boldsymbol{\zeta} \in \{\mathbf{x}, \mathbf{y}\}$, we have that the total selection gradient of the genotype is

$$\left.\frac{\mathrm{d}w}{\mathrm{d}\mathbf{y}}\right|_{\mathbf{y}=\bar{\mathbf{y}}} = \left.\left[\frac{\mathrm{d}\mathbf{x}^{\intercal}}{\mathrm{d}\mathbf{y}}\left(\frac{\partial w}{\partial \mathbf{x}} + \frac{\partial \boldsymbol{\epsilon}^{\intercal}}{\partial \mathbf{x}}\frac{\partial w}{\partial \boldsymbol{\epsilon}}\right) + \frac{\partial w}{\partial \mathbf{y}} + \frac{\partial \boldsymbol{\epsilon}^{\intercal}}{\partial \mathbf{y}}\frac{\partial w}{\partial \boldsymbol{\epsilon}}\right]\right|_{\mathbf{y}=\bar{\mathbf{y}}}.$$

Thus, using Layer 4, Eq. S10 yields the first line of Layer 4, Eq. S21.

**Form 2.** Using Eq. S5.2.6 and Eq. S5.2.13, the total selection gradient of the genotype is given by the second line of Layer 4, Eq. S21.

**Form 3.** Using Eq. S5.2.6, Eq. S5.1.20, and Layer 4, Eq. S7, we have that the total selection gradient of the genotype is given by the third line of Layer 4, Eq. S21.

**Form 4.** Using the first line of Layer 4, Eq. S21 and Layer 4, Eq. S14, we obtain the fourth line of Layer 4, Eq. S21.

**Form 5.** Finally, we can rearrange total genotypic selection (Layer 4, Eq. S21) in terms of total selection on the phenotype. Using Layer 4, Eq. S2 in the second line of Layer 4, Eq. S21, and then using the second line of Layer 4, Eq. S20, we have that the total selection gradient of the genotype is given by the fifth line of Layer 4, Eq. S21.

#### S5.3 Total selection gradient of the environment

Here proceed analogously to derive the total selection gradient of the environment, which allows us to write an equation describing the evolutionary dynamics of the geno-envo-phenotype.

##### S5.3.1 Total selection gradient of the environment in terms of direct fitness effects

As before, we start by considering the total selection gradient entry for the $i$-th environmental trait at age $a$. By this, we mean the total selection gradient of a perturbation of $\epsilon_{ia}$ taken as initial condition of the developmental constraint (1) when applied at the ages $\{a,\ldots,n\}$. Consequently, an environmental perturbation at a given age does not affect the phenotype at earlier ages due to the arrow of developmental time. By letting $\zeta$ in Eq. S2.3.9 be $\epsilon_{ia}$, we have

$$\left.\frac{\mathrm{d}\lambda}{\mathrm{d}\epsilon_{ia}}\right|_{\mathbf{y}=\bar{\mathbf{y}}} = \left.\frac{\mathrm{d}w}{\mathrm{d}\epsilon_{ia}}\right|_{\mathbf{y}=\bar{\mathbf{y}}} = \sum_{j=1}^{N_{\mathrm{a}}} \left.\frac{\mathrm{d}w_j}{\mathrm{d}\epsilon_{ia}}\right|_{\mathbf{y}=\bar{\mathbf{y}}}. \qquad \text{(Eq. S5.3.1)}$$

The total derivatives of a mutant's relative fitness at age $j$ in Eq. S5.3.1 are with respect to the individual's environmental traits at possibly another age $a$. We now seek to express such selection gradient in terms of partial derivatives only.

From Eq. S2.3.7, we have $w_j(\mathbf{z}_j, \boldsymbol{\epsilon}_j, \bar{\mathbf{m}})$ with $\mathbf{z}_j = (\mathbf{x}_j; \mathbf{y}_j)$, so applying the chain rule and, since we assume that genotypic traits are developmentally independent (hence, genotypic trait values do not depend on the environment, so $\mathrm{d}\mathbf{y}_j/\mathrm{d}\epsilon_{ia} = \mathbf{0}$ for all $i \in \{1,\ldots,N_{\mathrm{p}}\}$ and all $a, j \in \{1,\ldots,N_{\mathrm{a}}\}$), we obtain

$$\begin{aligned}\left.\frac{\mathrm{d}w_j}{\mathrm{d}\epsilon_{ia}}\right|_{\mathbf{y}=\bar{\mathbf{y}}} &= \left.\left(\sum_{k=1}^{N_{\mathrm{p}}} \frac{\partial w_j}{\partial x_{kj}} \frac{\mathrm{d}x_{kj}}{\mathrm{d}\epsilon_{ia}} + \sum_{k=1}^{N_{\mathrm{e}}} \frac{\partial w_j}{\partial \epsilon_{kj}} \frac{\mathrm{d}\epsilon_{kj}}{\mathrm{d}\epsilon_{ia}}\right)\right|_{\mathbf{y}=\bar{\mathbf{y}}} \\ &= \left.\left(\frac{\mathrm{d}\mathbf{x}_j^{\intercal}}{\mathrm{d}\epsilon_{ia}} \frac{\partial w_j}{\partial \mathbf{x}_j} + \frac{\mathrm{d}\boldsymbol{\epsilon}_j^{\intercal}}{\mathrm{d}\epsilon_{ia}} \frac{\partial w_j}{\partial \boldsymbol{\epsilon}_j}\right)\right|_{\mathbf{y}=\bar{\mathbf{y}}}.\end{aligned}$$

In the last equality we applied matrix calculus notation (Appendix A). Using Eq. S5.1.3 we have

$$\left.\frac{\mathrm{d}w_j}{\mathrm{d}\epsilon_{ia}}\right|_{\mathbf{y}=\bar{\mathbf{y}}} = \left.\left(\frac{\mathrm{d}\mathbf{x}_j^{\intercal}}{\mathrm{d}\epsilon_{ia}} \frac{\partial w}{\partial \mathbf{x}_j} + \frac{\mathrm{d}\boldsymbol{\epsilon}_j^{\intercal}}{\mathrm{d}\epsilon_{ia}} \frac{\partial w}{\partial \boldsymbol{\epsilon}_j}\right)\right|_{\mathbf{y}=\bar{\mathbf{y}}}. \qquad \text{(Eq. S5.3.2)}$$

Substituting Eq. S5.3.2 in Eq. S5.3.1 yields

$$\begin{aligned}\left.\frac{\mathrm{d}w}{\mathrm{d}\epsilon_{ia}}\right|_{\mathbf{y}=\bar{\mathbf{y}}} &= \sum_{j=1}^{N_{\mathrm{a}}} \left.\left(\frac{\mathrm{d}\mathbf{x}_j^{\intercal}}{\mathrm{d}\epsilon_{ia}} \frac{\partial w}{\partial \mathbf{x}_j} + \frac{\mathrm{d}\boldsymbol{\epsilon}_j^{\intercal}}{\mathrm{d}\epsilon_{ia}} \frac{\partial w}{\partial \boldsymbol{\epsilon}_j}\right)\right|_{\mathbf{y}=\bar{\mathbf{y}}} \\ &= \left.\left(\frac{\mathrm{d}\mathbf{x}^{\intercal}}{\mathrm{d}\epsilon_{ia}} \frac{\partial w}{\partial \mathbf{x}} + \frac{\mathrm{d}\boldsymbol{\epsilon}^{\intercal}}{\mathrm{d}\epsilon_{ia}} \frac{\partial w}{\partial \boldsymbol{\epsilon}}\right)\right|_{\mathbf{y}=\bar{\mathbf{y}}}.\end{aligned}$$

Therefore, the total selection gradient of all environmental traits across all ages is

$$\left.\frac{\mathrm{d}w}{\mathrm{d}\boldsymbol{\epsilon}}\right|_{\mathbf{y}=\bar{\mathbf{y}}} = \left.\left(\frac{\mathrm{d}\mathbf{x}^{\intercal}}{\mathrm{d}\boldsymbol{\epsilon}} \frac{\partial w}{\partial \mathbf{x}} + \frac{\mathrm{d}\boldsymbol{\epsilon}^{\intercal}}{\mathrm{d}\boldsymbol{\epsilon}} \frac{\partial w}{\partial \boldsymbol{\epsilon}}\right)\right|_{\mathbf{y}=\bar{\mathbf{y}}} \in \mathbb{R}^{N_{\mathrm{a}}N_{\mathrm{e}}\times 1}, \qquad \text{(Eq. S5.3.3)}$$

where we use the block matrix of *total effects of a mutant's environment on her phenotype*

$$\left.\frac{\mathrm{d}\mathbf{x}^\intercal}{\mathrm{d}\boldsymbol{\epsilon}}\right|_{\mathbf{y}=\bar{\mathbf{y}}} \equiv \left.\begin{pmatrix} \frac{\mathrm{d}\mathbf{x}_1^\intercal}{\mathrm{d}\boldsymbol{\epsilon}_1} & \cdots & \frac{\mathrm{d}\mathbf{x}_{N_\mathrm{a}}^\intercal}{\partial\boldsymbol{\epsilon}_1} \\ \vdots & \ddots & \vdots \\ \frac{\mathrm{d}\mathbf{x}_1^\intercal}{\mathrm{d}\boldsymbol{\epsilon}_{N_\mathrm{a}}} & \cdots & \frac{\mathrm{d}\mathbf{x}_{N_\mathrm{a}}^\intercal}{\mathrm{d}\boldsymbol{\epsilon}_{N_\mathrm{a}}} \end{pmatrix}\right|_{\mathbf{y}=\bar{\mathbf{y}}} \in \mathbb{R}^{N_\mathrm{a}N_\mathrm{p}\times N_\mathrm{a}N_\mathrm{e}}$$

and the block matrix of *total effects of a mutant's environment on her environment*

$$\left.\frac{\mathrm{d}\boldsymbol{\epsilon}^\intercal}{\mathrm{d}\boldsymbol{\epsilon}}\right|_{\mathbf{y}=\bar{\mathbf{y}}} \equiv \left.\begin{pmatrix} \frac{\mathrm{d}\boldsymbol{\epsilon}_1^\intercal}{\mathrm{d}\boldsymbol{\epsilon}_1} & \cdots & \frac{\mathrm{d}\boldsymbol{\epsilon}_{N_\mathrm{a}}^\intercal}{\partial\boldsymbol{\epsilon}_1} \\ \vdots & \ddots & \vdots \\ \frac{\mathrm{d}\boldsymbol{\epsilon}_1^\intercal}{\mathrm{d}\boldsymbol{\epsilon}_{N_\mathrm{a}}} & \cdots & \frac{\mathrm{d}\boldsymbol{\epsilon}_{N_\mathrm{a}}^\intercal}{\mathrm{d}\boldsymbol{\epsilon}_{N_\mathrm{a}}} \end{pmatrix}\right|_{\mathbf{y}=\bar{\mathbf{y}}} \in \mathbb{R}^{N_\mathrm{a}N_\mathrm{e}\times N_\mathrm{a}N_\mathrm{e}}.$$

Eq. S5.3.3 is now in terms of partial derivatives of fitness, total effects of a mutant's environment on her phenotype, $\mathrm{d}\mathbf{x}^\intercal/\mathrm{d}\boldsymbol{\epsilon}$, and total effects of a mutant's environment on her environment, $\mathrm{d}\boldsymbol{\epsilon}^\intercal/\mathrm{d}\boldsymbol{\epsilon}$. We now proceed to write $\mathrm{d}\mathbf{x}^\intercal/\mathrm{d}\boldsymbol{\epsilon}$ and $\mathrm{d}\boldsymbol{\epsilon}^\intercal/\mathrm{d}\boldsymbol{\epsilon}$ in terms of partial derivatives only.

##### S5.3.2 Matrix of total effects of a mutant's environment on her environment

From the environmental constraint (2) for the $k$-th environmental trait at age $j \in \{1,\ldots,N_\mathrm{a}\}$ we have that $\epsilon_{kj} = h_{kj}(\mathbf{z}_j,\bar{\mathbf{z}},\tau)$, so using the chain rule since genotypic traits are developmentally independent yields

$$\left.\frac{\mathrm{d}\epsilon_{kj}}{\mathrm{d}\epsilon_{ia}}\right|_{\mathbf{y}=\bar{\mathbf{y}}} = \begin{cases} \left.\left(\sum_{l=1}^{N_\mathrm{p}} \frac{\partial h_{kj}}{\partial x_{lj}}\frac{\mathrm{d}x_{lj}}{\mathrm{d}\epsilon_{ia}}\right)\right|_{\mathbf{y}=\bar{\mathbf{y}}} & \text{for } j>a \\ \left.\frac{\partial\epsilon_{kj}}{\partial\epsilon_{ia}}\right|_{\mathbf{y}=\bar{\mathbf{y}}} & \text{for } j=a \\ 0 & \text{for } j<a \end{cases}$$

$$= \begin{cases} \left.\left(\frac{\mathrm{d}\mathbf{x}_j^\intercal}{\mathrm{d}\epsilon_{ia}}\frac{\partial\epsilon_{kj}}{\partial\mathbf{x}_j}\right)\right|_{\mathbf{y}=\bar{\mathbf{y}}} & \text{for } j>a \\ \left.\frac{\partial\epsilon_{kj}}{\partial\epsilon_{ia}}\right|_{\mathbf{y}=\bar{\mathbf{y}}} & \text{for } j=a \\ 0 & \text{for } j<a. \end{cases}$$

In the last equality we used matrix calculus notation and rewrote $h_{kj}$ as $\epsilon_{kj}$. Since we assume that environmental traits are mutually independent, we have that $\partial\epsilon_{ka}/\partial\epsilon_{ia} = 1$ if $i=k$ or $\partial\epsilon_{ka}/\partial\epsilon_{ia} = 0$ otherwise; however, we leave the partial derivatives $\partial\epsilon_{ka}/\partial\epsilon_{ia}$ unevaluated as it is conceptually useful. Hence,

$$\left.\frac{\mathrm{d}\boldsymbol{\epsilon}_j^\intercal}{\mathrm{d}\epsilon_{ia}}\right|_{\mathbf{y}=\bar{\mathbf{y}}} = \begin{cases} \left.\left(\frac{\mathrm{d}\mathbf{x}_j^\intercal}{\mathrm{d}\epsilon_{ia}}\frac{\partial\boldsymbol{\epsilon}_j^\intercal}{\partial\mathbf{x}_j}\right)\right|_{\mathbf{y}=\bar{\mathbf{y}}} & \text{for } j>a \\ \left.\frac{\partial\boldsymbol{\epsilon}_j^\intercal}{\partial\epsilon_{ia}}\right|_{\mathbf{y}=\bar{\mathbf{y}}} & \text{for } j=a \\ \mathbf{0} & \text{for } j<a. \end{cases}$$

Then, the matrix of total effects of a mutant's environment at age $a$ on her environment at age $j$ is

$$\left.\frac{\mathrm{d}\boldsymbol{\epsilon}_j^\intercal}{\mathrm{d}\boldsymbol{\epsilon}_a}\right|_{\mathbf{y}=\bar{\mathbf{y}}} = \begin{cases} \left.\left(\frac{\mathrm{d}\mathbf{x}_j^\intercal}{\mathrm{d}\boldsymbol{\epsilon}_a}\frac{\partial\boldsymbol{\epsilon}_j^\intercal}{\partial\mathbf{x}_j}\right)\right|_{\mathbf{y}=\bar{\mathbf{y}}} & \text{for } j>a \\ \left.\frac{\partial\boldsymbol{\epsilon}_j^\intercal}{\partial\boldsymbol{\epsilon}_a}\right|_{\mathbf{y}=\bar{\mathbf{y}}} & \text{for } j=a \\ \mathbf{0} & \text{for } j<a. \end{cases} \qquad \text{(Eq. S5.3.4)}$$

Hence, the block matrix of *total effects of a mutant's environment on her environment* is

$$\left.\frac{\mathrm{d}\boldsymbol{\epsilon}^\intercal}{\mathrm{d}\boldsymbol{\epsilon}}\right|_{\mathbf{y}=\bar{\mathbf{y}}} \equiv \left.\begin{pmatrix} \frac{\mathrm{d}\boldsymbol{\epsilon}_1^\intercal}{\mathrm{d}\boldsymbol{\epsilon}_1} & \cdots & \frac{\mathrm{d}\boldsymbol{\epsilon}_{N_\mathrm{a}}^\intercal}{\mathrm{d}\boldsymbol{\epsilon}_1} \\ \vdots & \ddots & \vdots \\ \frac{\mathrm{d}\boldsymbol{\epsilon}_1^\intercal}{\mathrm{d}\boldsymbol{\epsilon}_{N_\mathrm{a}}} & \cdots & \frac{\mathrm{d}\boldsymbol{\epsilon}_{N_\mathrm{a}}^\intercal}{\mathrm{d}\boldsymbol{\epsilon}_{N_\mathrm{a}}} \end{pmatrix}\right|_{\mathbf{y}=\bar{\mathbf{y}}}$$

$$= \left.\begin{pmatrix} \frac{\partial\boldsymbol{\epsilon}_1^\intercal}{\partial\boldsymbol{\epsilon}_1} & \frac{\mathrm{d}\boldsymbol{\epsilon}_2^\intercal}{\mathrm{d}\boldsymbol{\epsilon}_1} & \cdots & \frac{\mathrm{d}\boldsymbol{\epsilon}_{N_\mathrm{a}-1}^\intercal}{\mathrm{d}\boldsymbol{\epsilon}_1} & \frac{\mathrm{d}\boldsymbol{\epsilon}_{N_\mathrm{a}}^\intercal}{\mathrm{d}\boldsymbol{\epsilon}_1} \\ \mathbf{0} & \frac{\partial\boldsymbol{\epsilon}_2^\intercal}{\partial\boldsymbol{\epsilon}_2} & \cdots & \frac{\mathrm{d}\boldsymbol{\epsilon}_{N_\mathrm{a}-1}^\intercal}{\mathrm{d}\boldsymbol{\epsilon}_2} & \frac{\mathrm{d}\boldsymbol{\epsilon}_{N_\mathrm{a}}^\intercal}{\mathrm{d}\boldsymbol{\epsilon}_2} \\ \vdots & \vdots & \ddots & \vdots & \vdots \\ \mathbf{0} & \mathbf{0} & \cdots & \frac{\partial\boldsymbol{\epsilon}_{N_\mathrm{a}-1}^\intercal}{\partial\boldsymbol{\epsilon}_{N_\mathrm{a}-1}} & \frac{\mathrm{d}\boldsymbol{\epsilon}_{N_\mathrm{a}}^\intercal}{\mathrm{d}\boldsymbol{\epsilon}_{N_\mathrm{a}-1}} \\ \mathbf{0} & \mathbf{0} & \cdots & \mathbf{0} & \frac{\partial\boldsymbol{\epsilon}_{N_\mathrm{a}}^\intercal}{\partial\boldsymbol{\epsilon}_{N_\mathrm{a}}} \end{pmatrix}\right|_{\mathbf{y}=\bar{\mathbf{y}}} \qquad \text{(Eq. S5.3.5)}$$

$$\in \mathbb{R}^{N_\mathrm{a}N_\mathrm{e}\times N_\mathrm{a}N_\mathrm{e}}.$$

Note that the $aj$-th block entry of $(\mathrm{d}\mathbf{x}^\intercal/\mathrm{d}\boldsymbol{\epsilon})(\partial\boldsymbol{\epsilon}^\intercal/\partial\mathbf{x})$ for $j>a$ is

$$\left(\frac{\mathrm{d}\mathbf{x}^\intercal}{\mathrm{d}\boldsymbol{\epsilon}}\frac{\partial\boldsymbol{\epsilon}^\intercal}{\partial\mathbf{x}}\right)_{aj} = \sum_{k=1}^{N_\mathrm{a}} \frac{\mathrm{d}\mathbf{x}_k^\intercal}{\mathrm{d}\boldsymbol{\epsilon}_a}\frac{\partial\boldsymbol{\epsilon}_j^\intercal}{\partial\mathbf{x}_k} = \frac{\mathrm{d}\mathbf{x}_j^\intercal}{\mathrm{d}\boldsymbol{\epsilon}_a}\frac{\partial\boldsymbol{\epsilon}_j^\intercal}{\partial\mathbf{x}_j},$$

where we use Layer 2, Eq. S2d in the second equality. Note also that since environmental traits are mutually independent, $\partial\boldsymbol{\epsilon}_j^\intercal/\partial\boldsymbol{\epsilon}_a = \mathbf{0}$ for $j \neq a$ from the environmental constraint (2). Finally, note that because of the arrow of developmental time, $\partial\mathbf{x}_j^\intercal/\partial\boldsymbol{\epsilon}_a = \mathbf{0}$ for $j<a$ due to the developmental constraint (1). Hence, Layer 4, Eq. S12 follows, which is a compact expression for the matrix of total effects of a mutant's environment on itself in terms of partial derivatives and the total effects of a mutant's environment on her phenotype, which we now write in terms of partial derivatives only.

##### S5.3.3 Matrix of total effects of a mutant's environment on her phenotype

From the developmental constraint (1) for the $k$-th phenotype at age $j \in \{2,\ldots,N_\mathrm{a}\}$ we have that $x_{kj} = g_{k,j-1}(\mathbf{z}_{j-1},\boldsymbol{\epsilon}_{j-1},\bar{\mathbf{z}})$, so using the chain rule and since genotypic traits are developmentally independent yields

$$\left.\frac{\mathrm{d}x_{kj}}{\mathrm{d}\epsilon_{ia}}\right|_{\mathbf{y}=\bar{\mathbf{y}}} = \left.\left(\sum_{l=1}^{N_\mathrm{p}} \frac{\partial g_{k,j-1}}{\partial x_{l,j-1}}\frac{\mathrm{d}x_{l,j-1}}{\mathrm{d}\epsilon_{ia}} + \sum_{l=1}^{N_\mathrm{e}} \frac{\partial g_{k,j-1}}{\partial\epsilon_{l,j-1}}\frac{\mathrm{d}\epsilon_{l,j-1}}{\mathrm{d}\epsilon_{ia}}\right)\right|_{\mathbf{y}=\bar{\mathbf{y}}}$$

$$= \left(\frac{\mathrm{d}\mathbf{x}_{j-1}^{\intercal}}{\mathrm{d}\epsilon_{ia}}\frac{\partial x_{kj}}{\partial \mathbf{x}_{j-1}} + \frac{\mathrm{d}\boldsymbol{\epsilon}_{j-1}^{\intercal}}{\mathrm{d}\epsilon_{ia}}\frac{\partial x_{kj}}{\partial \boldsymbol{\epsilon}_{j-1}}\right)\Bigg|_{\mathbf{y}=\bar{\mathbf{y}}}.$$

In the last equality we used matrix calculus notation and rewrote $g_{k,j-1}$ as $x_{kj}$. Hence,

$$\left.\frac{\mathrm{d}\mathbf{x}_j^{\intercal}}{\mathrm{d}\epsilon_{ia}}\right|_{\mathbf{y}=\bar{\mathbf{y}}} = \left(\frac{\mathrm{d}\mathbf{x}_{j-1}^{\intercal}}{\mathrm{d}\epsilon_{ia}}\frac{\partial \mathbf{x}_j^{\intercal}}{\partial \mathbf{x}_{j-1}} + \frac{\mathrm{d}\boldsymbol{\epsilon}_{j-1}^{\intercal}}{\mathrm{d}\epsilon_{ia}}\frac{\partial \mathbf{x}_j^{\intercal}}{\partial \boldsymbol{\epsilon}_{j-1}}\right)\Bigg|_{\mathbf{y}=\bar{\mathbf{y}}}.$$

Then, the matrix of total effects of a mutant's environment at age $a$ on her phenotype at age $j$ is

$$\left.\frac{\mathrm{d}\mathbf{x}_j^{\intercal}}{\mathrm{d}\boldsymbol{\epsilon}_a}\right|_{\mathbf{y}=\bar{\mathbf{y}}} = \left(\frac{\mathrm{d}\mathbf{x}_{j-1}^{\intercal}}{\mathrm{d}\boldsymbol{\epsilon}_a}\frac{\partial \mathbf{x}_j^{\intercal}}{\partial \mathbf{x}_{j-1}} + \frac{\mathrm{d}\boldsymbol{\epsilon}_{j-1}^{\intercal}}{\mathrm{d}\boldsymbol{\epsilon}_a}\frac{\partial \mathbf{x}_j^{\intercal}}{\partial \boldsymbol{\epsilon}_{j-1}}\right)\Bigg|_{\mathbf{y}=\bar{\mathbf{y}}}.$$

Using Eq. S5.3.4 yields

$$\left.\frac{\mathrm{d}\mathbf{x}_j^{\intercal}}{\mathrm{d}\boldsymbol{\epsilon}_a}\right|_{\mathbf{y}=\bar{\mathbf{y}}} =$$

$$\begin{cases} \left(\frac{\mathrm{d}\mathbf{x}_{j-1}^{\intercal}}{\mathrm{d}\boldsymbol{\epsilon}_a}\frac{\partial \mathbf{x}_j^{\intercal}}{\partial \mathbf{x}_{j-1}} + \frac{\mathrm{d}\mathbf{x}_{j-1}^{\intercal}}{\mathrm{d}\boldsymbol{\epsilon}_a}\frac{\partial \boldsymbol{\epsilon}_{j-1}^{\intercal}}{\partial \mathbf{x}_{j-1}}\frac{\partial \mathbf{x}_j^{\intercal}}{\partial \boldsymbol{\epsilon}_{j-1}}\right)\Bigg|_{\mathbf{y}=\bar{\mathbf{y}}} & \text{for } j-1>a \\ \left(\underbrace{\frac{\mathrm{d}\mathbf{x}_a^{\intercal}}{\mathrm{d}\boldsymbol{\epsilon}_a}}_{\mathbf{0},\text{ by (1)}}\frac{\partial \mathbf{x}_{a+1}^{\intercal}}{\partial \mathbf{x}_a} + \frac{\partial \boldsymbol{\epsilon}_a^{\intercal}}{\partial \boldsymbol{\epsilon}_a}\frac{\partial \mathbf{x}_{a+1}^{\intercal}}{\partial \boldsymbol{\epsilon}_a}\right)\Bigg|_{\mathbf{y}=\bar{\mathbf{y}}} & \text{for } j-1=a \\ \left(\underbrace{\frac{\mathrm{d}\mathbf{x}_{j-1}^{\intercal}}{\mathrm{d}\boldsymbol{\epsilon}_a}}_{\mathbf{0},\text{ by (1)}}\frac{\partial \mathbf{x}_j^{\intercal}}{\partial \mathbf{x}_{j-1}}\right)\Bigg|_{\mathbf{y}=\bar{\mathbf{y}}} & \text{for } j-1>a \end{cases}$$

$$= \begin{cases} \left[\frac{\mathrm{d}\mathbf{x}_{j-1}^{\intercal}}{\mathrm{d}\boldsymbol{\epsilon}_a}\left(\frac{\partial \mathbf{x}_j^{\intercal}}{\partial \mathbf{x}_{j-1}} + \frac{\partial \boldsymbol{\epsilon}_{j-1}^{\intercal}}{\partial \mathbf{x}_{j-1}}\frac{\partial \mathbf{x}_j^{\intercal}}{\partial \boldsymbol{\epsilon}_{j-1}}\right)\right]\Bigg|_{\mathbf{y}=\bar{\mathbf{y}}} & \text{for } j-1>a \\ \left(\frac{\partial \boldsymbol{\epsilon}_a^{\intercal}}{\partial \boldsymbol{\epsilon}_a}\frac{\partial \mathbf{x}_j^{\intercal}}{\partial \boldsymbol{\epsilon}_{j-1}}\right)\Bigg|_{\mathbf{y}=\bar{\mathbf{y}}} & \text{for } j-1=a \\ \mathbf{0} & \text{for } j-1>a. \end{cases}$$

Using Eq. S5.1.10, this reduces to

$$\left.\frac{\mathrm{d}\mathbf{x}_j^{\intercal}}{\mathrm{d}\boldsymbol{\epsilon}_a}\right|_{\mathbf{y}=\bar{\mathbf{y}}} = \begin{cases} \left(\frac{\mathrm{d}\mathbf{x}_{j-1}^{\intercal}}{\mathrm{d}\boldsymbol{\epsilon}_a}\frac{\delta \mathbf{x}_j^{\intercal}}{\delta \mathbf{x}_{j-1}}\right)\Bigg|_{\mathbf{y}=\bar{\mathbf{y}}} & \text{for } j-1>a \\ \left(\frac{\partial \boldsymbol{\epsilon}_a^{\intercal}}{\partial \boldsymbol{\epsilon}_a}\frac{\partial \mathbf{x}_{a+1}^{\intercal}}{\partial \boldsymbol{\epsilon}_a}\right)\Bigg|_{\mathbf{y}=\bar{\mathbf{y}}} & \text{for } j-1=a \\ \mathbf{0} & \text{for } j-1>a. \end{cases}$$

Expanding this recurrence yields

$$\left.\frac{\mathrm{d}\mathbf{x}_j^{\intercal}}{\mathrm{d}\boldsymbol{\epsilon}_a}\right|_{\mathbf{y}=\bar{\mathbf{y}}} = \begin{cases} \left(\frac{\mathrm{d}\mathbf{x}_{a+1}^{\intercal}}{\mathrm{d}\boldsymbol{\epsilon}_a}\frac{\delta \mathbf{x}_{a+2}^{\intercal}}{\delta \mathbf{x}_{a+1}}\cdots\frac{\delta \mathbf{x}_j^{\intercal}}{\delta \mathbf{x}_{j-1}}\right)\Bigg|_{\mathbf{y}=\bar{\mathbf{y}}} & \text{for } j-1>a \\ \left(\frac{\partial \boldsymbol{\epsilon}_a^{\intercal}}{\partial \boldsymbol{\epsilon}_a}\frac{\partial \mathbf{x}_{a+1}^{\intercal}}{\partial \boldsymbol{\epsilon}_a}\right)\Bigg|_{\mathbf{y}=\bar{\mathbf{y}}} & \text{for } j-1=a \\ \mathbf{0} & \text{for } j-1>a, \end{cases}$$

which using Eq. (12) yields

$$\left.\frac{\mathrm{d}\mathbf{x}_j^{\intercal}}{\mathrm{d}\boldsymbol{\epsilon}_a}\right|_{\mathbf{y}=\bar{\mathbf{y}}} = \begin{cases} \left(\frac{\partial \boldsymbol{\epsilon}_a^{\intercal}}{\partial \boldsymbol{\epsilon}_a}\frac{\partial \mathbf{x}_{a+1}^{\intercal}}{\partial \boldsymbol{\epsilon}_a}\frac{\mathrm{d}\mathbf{x}_j^{\intercal}}{\mathrm{d}\mathbf{x}_{a+1}}\right)\Bigg|_{\mathbf{y}=\bar{\mathbf{y}}} & \text{for } j-1>a \\ \left(\frac{\partial \boldsymbol{\epsilon}_a^{\intercal}}{\partial \boldsymbol{\epsilon}_a}\frac{\partial \mathbf{x}_{a+1}^{\intercal}}{\partial \boldsymbol{\epsilon}_a}\right)\Bigg|_{\mathbf{y}=\bar{\mathbf{y}}} & \text{for } j-1=a \\ \mathbf{0} & \text{for } j-1>a. \end{cases} \quad \text{(Eq. S5.3.6)}$$

It will be useful to denote the matrix of *total immediate effects of a mutant's environment at age j on her phenotype at age j* for $j>0$ as

$$\left.\frac{\delta \mathbf{x}_j^{\intercal}}{\delta \boldsymbol{\epsilon}_{j-1}}\right|_{\mathbf{y}=\bar{\mathbf{y}}} = \left.\frac{\partial \boldsymbol{\epsilon}_{j-1}^{\intercal}}{\partial \boldsymbol{\epsilon}_{j-1}}\frac{\partial \mathbf{x}_j^{\intercal}}{\partial \boldsymbol{\epsilon}_{j-1}}\right|_{\mathbf{y}=\bar{\mathbf{y}}} \in \mathbb{R}^{N_\mathrm{e}\times N_\mathrm{p}}. \quad \text{(Eq. S5.3.7)}$$

The matrix of *direct effects of a mutant's environment on itself* is given by Layer 2, Eq. S3. In turn, the block matrix of *total immediate effects of a mutant's environment on her phenotype* is

$$\left.\frac{\delta \mathbf{x}^{\intercal}}{\delta \boldsymbol{\epsilon}}\right|_{\mathbf{y}=\bar{\mathbf{y}}} \equiv \left.\begin{pmatrix} \frac{\delta \mathbf{x}_1^{\intercal}}{\delta \boldsymbol{\epsilon}_1} & \cdots & \frac{\delta \mathbf{x}_{N_\mathrm{a}}^{\intercal}}{\delta \boldsymbol{\epsilon}_1} \\ \vdots & \ddots & \vdots \\ \frac{\delta \mathbf{x}_1^{\intercal}}{\delta \boldsymbol{\epsilon}_{N_\mathrm{a}}} & \cdots & \frac{\delta \mathbf{x}_{N_\mathrm{a}}^{\intercal}}{\delta \boldsymbol{\epsilon}_{N_\mathrm{a}}} \end{pmatrix}\right|_{\mathbf{y}=\bar{\mathbf{y}}}$$

$$= \left.\begin{pmatrix} \mathbf{0} & \frac{\delta \mathbf{x}_2^{\intercal}}{\delta \boldsymbol{\epsilon}_1} & \cdots & \mathbf{0} & \mathbf{0} \\ \mathbf{0} & \mathbf{0} & \cdots & \mathbf{0} & \mathbf{0} \\ \vdots & \vdots & \ddots & \vdots & \vdots \\ \mathbf{0} & \mathbf{0} & \cdots & \mathbf{0} & \frac{\delta \mathbf{x}_{N_\mathrm{a}}^{\intercal}}{\delta \boldsymbol{\epsilon}_{N_\mathrm{a}-1}} \\ \mathbf{0} & \mathbf{0} & \cdots & \mathbf{0} & \mathbf{0} \end{pmatrix}\right|_{\mathbf{y}=\bar{\mathbf{y}}}$$

$$\in \mathbb{R}^{N_\mathrm{a}N_\mathrm{e}\times N_\mathrm{a}N_\mathrm{p}}, \quad \text{(Eq. S5.3.8)}$$

so Layer 3, Eq. S4 follows from Eq. S5.3.7, Layer 2, Eq. S3, and Layer 2, Eq. S2c.

Using Eq. S5.3.7, Eq. S5.3.6 becomes

$$\left.\frac{\mathrm{d}\mathbf{x}_j^{\intercal}}{\mathrm{d}\boldsymbol{\epsilon}_a}\right|_{\mathbf{y}=\bar{\mathbf{y}}} = \begin{cases} \left(\frac{\delta \mathbf{x}_{a+1}^{\intercal}}{\delta \boldsymbol{\epsilon}_a}\frac{\mathrm{d}\mathbf{x}_j^{\intercal}}{\mathrm{d}\mathbf{x}_{a+1}}\right)\Bigg|_{\mathbf{y}=\bar{\mathbf{y}}} & \text{for } j-1>a \\ \left.\frac{\delta \mathbf{x}_{a+1}^{\intercal}}{\delta \boldsymbol{\epsilon}_a}\right|_{\mathbf{y}=\bar{\mathbf{y}}} & \text{for } j-1=a \\ \mathbf{0} & \text{for } j-1>a. \end{cases}$$

Note that the $aj$-th entry of $(\delta\mathbf{x}^{\intercal}/\delta\boldsymbol{\epsilon})(\mathrm{d}\mathbf{x}^{\intercal}/\mathrm{d}\mathbf{x})$ is

$$\left(\frac{\delta \mathbf{x}^{\intercal}}{\delta \boldsymbol{\epsilon}}\right)_{aj} = \sum_{k=1}^{N_\mathrm{a}}\frac{\delta \mathbf{x}_k^{\intercal}}{\delta \boldsymbol{\epsilon}_a}\frac{\mathrm{d}\mathbf{x}_j^{\intercal}}{\mathrm{d}\mathbf{x}_k} = \frac{\delta \mathbf{x}_{a+1}^{\intercal}}{\delta \boldsymbol{\epsilon}_a}\frac{\mathrm{d}\mathbf{x}_j^{\intercal}}{\mathrm{d}\mathbf{x}_{a+1}} = \frac{\mathrm{d}\mathbf{x}_j^{\intercal}}{\mathrm{d}\boldsymbol{\epsilon}_a}, \quad \text{(Eq. S5.3.9)}$$

where we use Eq. S5.3.8 in the second equality. Hence, Layer 4, Eq. S3 follows, where the block matrix of *total ef-*

*fects of a mutant's environment on her phenotype* is

$$\left.\frac{\mathrm{d}\mathbf{x}^{\mathsf{T}}}{\mathrm{d}\boldsymbol{\epsilon}}\right|_{\mathbf{y}=\bar{\mathbf{y}}} = \left.\begin{pmatrix} \frac{\mathrm{d}\mathbf{x}_1^{\mathsf{T}}}{\mathrm{d}\boldsymbol{\epsilon}_1} & \cdots & \frac{\mathrm{d}\mathbf{x}_{N_\mathrm{a}}^{\mathsf{T}}}{\mathrm{d}\boldsymbol{\epsilon}_1} \\ \vdots & \ddots & \vdots \\ \frac{\mathrm{d}\mathbf{x}_1^{\mathsf{T}}}{\mathrm{d}\boldsymbol{\epsilon}_{N_\mathrm{a}}} & \cdots & \frac{\mathrm{d}\mathbf{x}_{N_\mathrm{a}}^{\mathsf{T}}}{\mathrm{d}\boldsymbol{\epsilon}_{N_\mathrm{a}}} \end{pmatrix}\right|_{\mathbf{y}=\bar{\mathbf{y}}}$$

$$= \left.\begin{pmatrix} \mathbf{0} & \frac{\mathrm{d}\mathbf{x}_2^{\mathsf{T}}}{\mathrm{d}\boldsymbol{\epsilon}_1} & \cdots & \frac{\mathrm{d}\mathbf{x}_{N_\mathrm{a}-1}^{\mathsf{T}}}{\mathrm{d}\boldsymbol{\epsilon}_1} & \frac{\mathrm{d}\mathbf{x}_{N_\mathrm{a}}^{\mathsf{T}}}{\mathrm{d}\boldsymbol{\epsilon}_1} \\ \mathbf{0} & \mathbf{0} & \cdots & \frac{\mathrm{d}\mathbf{x}_{N_\mathrm{a}-1}^{\mathsf{T}}}{\mathrm{d}\boldsymbol{\epsilon}_2} & \frac{\mathrm{d}\mathbf{x}_{N_\mathrm{a}}^{\mathsf{T}}}{\mathrm{d}\boldsymbol{\epsilon}_2} \\ \vdots & \vdots & \ddots & \vdots & \vdots \\ \mathbf{0} & \mathbf{0} & \cdots & \mathbf{0} & \frac{\mathrm{d}\mathbf{x}_{N_\mathrm{a}}^{\mathsf{T}}}{\mathrm{d}\boldsymbol{\epsilon}_{N_\mathrm{a}-1}} \\ \mathbf{0} & \mathbf{0} & \cdots & \mathbf{0} & \mathbf{0} \end{pmatrix}\right|_{\mathbf{y}=\bar{\mathbf{y}}} \quad \text{(Eq. S5.3.10)}$$

$$\in \mathbb{R}^{N_\mathrm{a}N_\mathrm{e}\times N_\mathrm{a}N_\mathrm{p}}.$$

Layer 4, Eq. S3, Eq. S5.3.8, and Layer 4, Eq. S1 write the matrix of total effects of a mutant's environment on her phenotype in terms of partial derivatives. This is a compact expression for the matrix of total effects of a mutant's environment on her phenotype in terms of partial derivatives only.

##### S5.3.4 Conclusion

**Form 1.** Eq. S5.3.3 gives the total selection gradient of the environment as in the first line of Layer 4, Eq. S22.

**Form 2.** Using Eq. S5.3.3 and Layer 4, Eq. S12 yields

$$\left.\frac{\mathrm{d}w}{\mathrm{d}\boldsymbol{\epsilon}}\right|_{\mathbf{y}=\bar{\mathbf{y}}} = \left.\left[\frac{\mathrm{d}\mathbf{x}^{\mathsf{T}}}{\mathrm{d}\boldsymbol{\epsilon}}\frac{\partial w}{\partial \mathbf{x}} + \left(\frac{\partial\boldsymbol{\epsilon}^{\mathsf{T}}}{\partial\boldsymbol{\epsilon}} + \frac{\mathrm{d}\mathbf{x}^{\mathsf{T}}}{\mathrm{d}\boldsymbol{\epsilon}}\frac{\partial\boldsymbol{\epsilon}^{\mathsf{T}}}{\partial\mathbf{x}}\right)\frac{\partial w}{\partial\boldsymbol{\epsilon}}\right]\right|_{\mathbf{y}=\bar{\mathbf{y}}}.$$

Collecting for $\mathrm{d}\mathbf{x}^{\mathsf{T}}/\mathrm{d}\boldsymbol{\epsilon}$ and using Layer 3, Eq. S1 for $\boldsymbol{\zeta} = \mathbf{x}$ as well as Layer 3, Eq. S2, we have that the total selection gradient of the environment is given by the second line of Layer 4, Eq. S22.

**Form 3.** Using the first line of Layer 4, Eq. S22 and Layer 4, Eq. S15, we obtain the third line of Layer 4, Eq. S22.

**Form 4.** Finally, we can rearrange total selection on the environment in terms of total selection on the phenotype. Using Layer 4, Eq. S3 in the second line of Layer 4, Eq. S22, and then using the second line of Layer 4, Eq. S20, we have that the total selection gradient of the environment is given by the fourth line of Layer 4, Eq. S22.

#### S5.4 Total selection gradient of the geno-phenotype

We have that the mutant geno-phenotype is $\mathbf{z} = (\mathbf{x};\mathbf{y})$. We first define the (direct), total immediate, and total selection gradients of the geno-phenotype and write the total selection gradient of the geno-phenotype in terms of the total immediate selection gradient of the geno-phenotype and of the partial selection gradient of the geno-envo-phenotype.

##### S5.4.1 Total selection gradient of the geno-phenotype in terms of direct fitness effects

We have the *selection gradient of the geno-phenotype*

$$\left.\frac{\partial w}{\partial \mathbf{z}}\right|_{\mathbf{y}=\bar{\mathbf{y}}} \equiv \left.\left(\frac{\partial w}{\partial \mathbf{x}};\frac{\partial w}{\partial \mathbf{y}}\right)\right|_{\mathbf{y}=\bar{\mathbf{y}}} \in \mathbb{R}^{N_\mathrm{a}(N_\mathrm{p}+N_\mathrm{g})\times 1},$$

the *total immediate selection gradient of the geno-phenotype*

$$\left.\frac{\delta w}{\delta \mathbf{z}}\right|_{\mathbf{y}=\bar{\mathbf{y}}} \equiv \left.\left(\frac{\delta w}{\delta \mathbf{x}};\frac{\delta w}{\delta \mathbf{y}}\right)\right|_{\mathbf{y}=\bar{\mathbf{y}}} \in \mathbb{R}^{N_\mathrm{a}(N_\mathrm{p}+N_\mathrm{g})\times 1},$$

and the *total selection gradient of the geno-phenotype*

$$\left.\frac{\mathrm{d}w}{\mathrm{d}\mathbf{z}}\right|_{\mathbf{y}=\bar{\mathbf{y}}} \equiv \left.\left(\frac{\mathrm{d}w}{\mathrm{d}\mathbf{x}};\frac{\mathrm{d}w}{\mathrm{d}\mathbf{y}}\right)\right|_{\mathbf{y}=\bar{\mathbf{y}}} \in \mathbb{R}^{N_\mathrm{a}(N_\mathrm{p}+N_\mathrm{g})\times 1}.$$

Now, we write the total immediate selection gradient of the geno-phenotype as a linear combination of the selection gradients of the geno-phenotype and environment. Using Layer 3, Eq. S1 for $\boldsymbol{\zeta} \in \{\mathbf{x},\mathbf{y}\}$, we have that the total immediate selection gradient of the geno-phenotype is

$$\left.\frac{\delta w}{\delta \mathbf{z}}\right|_{\mathbf{y}=\bar{\mathbf{y}}} \equiv \left.\begin{pmatrix}\frac{\delta w}{\delta \mathbf{x}}\\ \frac{\delta w}{\delta \mathbf{y}}\end{pmatrix}\right|_{\mathbf{y}=\bar{\mathbf{y}}} = \left.\begin{pmatrix}\frac{\partial w}{\partial \mathbf{x}} + \frac{\partial\boldsymbol{\epsilon}^{\mathsf{T}}}{\partial \mathbf{x}}\frac{\partial w}{\partial\boldsymbol{\epsilon}}\\ \frac{\partial w}{\partial \mathbf{y}} + \frac{\partial\boldsymbol{\epsilon}^{\mathsf{T}}}{\partial \mathbf{y}}\frac{\partial w}{\partial\boldsymbol{\epsilon}}\end{pmatrix}\right|_{\mathbf{y}=\bar{\mathbf{y}}}$$

$$= \left.\left[\begin{pmatrix}\frac{\partial w}{\partial \mathbf{x}}\\ \frac{\partial w}{\partial \mathbf{y}}\end{pmatrix} + \begin{pmatrix}\frac{\partial\boldsymbol{\epsilon}^{\mathsf{T}}}{\partial \mathbf{x}}\frac{\partial w}{\partial\boldsymbol{\epsilon}}\\ \frac{\partial\boldsymbol{\epsilon}^{\mathsf{T}}}{\partial \mathbf{y}}\frac{\partial w}{\partial\boldsymbol{\epsilon}}\end{pmatrix}\right]\right|_{\mathbf{y}=\bar{\mathbf{y}}}. \quad \text{(Eq. S5.4.1)}$$

Using Layer 2, Eq. S7, we have that

$$\left.\left(\frac{\partial\boldsymbol{\epsilon}^{\mathsf{T}}}{\partial \mathbf{z}}\frac{\partial w}{\partial\boldsymbol{\epsilon}}\right)\right|_{\mathbf{y}=\bar{\mathbf{y}}} = \left.\left[\begin{pmatrix}\frac{\partial\boldsymbol{\epsilon}^{\mathsf{T}}}{\partial \mathbf{x}}\\ \frac{\partial\boldsymbol{\epsilon}^{\mathsf{T}}}{\partial \mathbf{y}}\end{pmatrix}\frac{\partial w}{\partial\boldsymbol{\epsilon}}\right]\right|_{\mathbf{y}=\bar{\mathbf{y}}} = \left.\begin{pmatrix}\frac{\partial\boldsymbol{\epsilon}^{\mathsf{T}}}{\partial \mathbf{x}}\frac{\partial w}{\partial\boldsymbol{\epsilon}}\\ \frac{\partial\boldsymbol{\epsilon}^{\mathsf{T}}}{\partial \mathbf{y}}\frac{\partial w}{\partial\boldsymbol{\epsilon}}\end{pmatrix}\right|_{\mathbf{y}=\bar{\mathbf{y}}}.$$

Therefore, Eq. S5.4.1 becomes Layer 3, Eq. S1 for $\boldsymbol{\zeta} = \mathbf{z}$.

**Form 2.** Now we bring together the total selection gradients of the phenotype and genotype to write the total selection gradient of the geno-phenotype as a linear transformation of the total immediate selection gradient of the geno-phenotype.

Using the third lines of Layer 4, Eq. S20 and Layer 4, Eq. S21, we have

$$\left.\frac{\mathrm{d}w}{\mathrm{d}\mathbf{z}}\right|_{\mathbf{y}=\bar{\mathbf{y}}} \equiv \left.\begin{pmatrix}\frac{\mathrm{d}w}{\mathrm{d}\mathbf{x}}\\ \frac{\mathrm{d}w}{\mathrm{d}\mathbf{y}}\end{pmatrix}\right|_{\mathbf{y}=\bar{\mathbf{y}}} = \left.\begin{pmatrix}\frac{\mathrm{d}\mathbf{z}^{\mathsf{T}}}{\mathrm{d}\mathbf{x}}\frac{\delta w}{\delta \mathbf{z}}\\ \frac{\mathrm{d}\mathbf{z}^{\mathsf{T}}}{\mathrm{d}\mathbf{y}}\frac{\delta w}{\delta \mathbf{z}}\end{pmatrix}\right|_{\mathbf{y}=\bar{\mathbf{y}}}$$

$$= \left.\left[\begin{pmatrix}\frac{\mathrm{d}\mathbf{z}^{\mathsf{T}}}{\mathrm{d}\mathbf{x}}\\ \frac{\mathrm{d}\mathbf{z}^{\mathsf{T}}}{\mathrm{d}\mathbf{y}}\end{pmatrix}\frac{\delta w}{\delta \mathbf{z}}\right]\right|_{\mathbf{y}=\bar{\mathbf{y}}} = \left.\left(\frac{\mathrm{d}\mathbf{z}^{\mathsf{T}}}{\mathrm{d}\mathbf{z}}\frac{\delta w}{\delta \mathbf{z}}\right)\right|_{\mathbf{y}=\bar{\mathbf{y}}},$$

which is the second line of Layer 4, Eq. S23.

**Form 3.** Now we use the expressions of the total selection gradients of the phenotype and genotype as linear transformations of the geno-envo-phenotype to write the total selection gradient of the geno-phenotype. Using the fourth lines of Layer 4, Eq. S20 and Layer 4, Eq. S21, we have

$$\left.\frac{\mathrm{d}w}{\mathrm{d}\mathbf{z}}\right|_{\mathbf{y}=\bar{\mathbf{y}}} \equiv \left.\begin{pmatrix} \frac{\mathrm{d}w}{\mathrm{d}\mathbf{x}} \\ \frac{\mathrm{d}w}{\mathrm{d}\mathbf{y}} \end{pmatrix}\right|_{\mathbf{y}=\bar{\mathbf{y}}} = \left.\begin{pmatrix} \frac{\mathrm{d}\mathbf{m}^\intercal}{\mathrm{d}\mathbf{x}}\frac{\partial w}{\partial \mathbf{m}} \\ \frac{\mathrm{d}\mathbf{m}^\intercal}{\mathrm{d}\mathbf{y}}\frac{\partial w}{\partial \mathbf{m}} \end{pmatrix}\right|_{\mathbf{y}=\bar{\mathbf{y}}}$$

$$= \left.\left[\begin{pmatrix} \frac{\mathrm{d}\mathbf{m}^\intercal}{\mathrm{d}\mathbf{x}} \\ \frac{\mathrm{d}\mathbf{m}^\intercal}{\mathrm{d}\mathbf{y}} \end{pmatrix}\frac{\partial w}{\partial \mathbf{m}}\right]\right|_{\mathbf{y}=\bar{\mathbf{y}}} = \left.\left(\frac{\mathrm{d}\mathbf{m}^\intercal}{\mathrm{d}\mathbf{z}}\frac{\partial w}{\partial \mathbf{m}}\right)\right|_{\mathbf{y}=\bar{\mathbf{y}}},$$

which is the third line of Layer 4, Eq. S23.

**Form 1.** Now, we obtain the total selection gradient of the geno-phenotype as a linear combination of selection gradients of the geno-phenotype and environment. Using Layer 3, Eq. S1 for $\boldsymbol{\zeta} = \mathbf{z}$, the second line of Layer 4, Eq. S23 becomes

$$\left.\frac{\mathrm{d}w}{\mathrm{d}\mathbf{z}}\right|_{\mathbf{y}=\bar{\mathbf{y}}} = \left.\left[\frac{\mathrm{d}\mathbf{z}^\intercal}{\mathrm{d}\mathbf{z}}\left(\frac{\partial w}{\partial \mathbf{z}} + \frac{\partial \boldsymbol{\epsilon}^\intercal}{\partial \mathbf{z}}\frac{\partial w}{\partial \boldsymbol{\epsilon}}\right)\right]\right|_{\mathbf{y}=\bar{\mathbf{y}}}. \qquad \text{(Eq. S5.4.2)}$$

We define the block matrix of total effects of a mutant's geno-phenotype on her environment as

$$\left.\frac{\mathrm{d}\boldsymbol{\epsilon}^\intercal}{\mathrm{d}\mathbf{z}}\right|_{\mathbf{y}=\bar{\mathbf{y}}} \equiv \left.\begin{pmatrix} \frac{\mathrm{d}\boldsymbol{\epsilon}^\intercal}{\mathrm{d}\mathbf{x}} \\ \frac{\mathrm{d}\boldsymbol{\epsilon}^\intercal}{\mathrm{d}\mathbf{y}} \end{pmatrix}\right|_{\mathbf{y}=\bar{\mathbf{y}}} \in \mathbb{R}^{N_\mathrm{a}(N_\mathrm{p}+N_\mathrm{g})\times N_\mathrm{a}N_\mathrm{e}},$$

which using Layer 4, Eq. S9 and Layer 4, Eq. S10 yields

$$\left.\frac{\mathrm{d}\boldsymbol{\epsilon}^\intercal}{\mathrm{d}\mathbf{z}}\right|_{\mathbf{y}=\bar{\mathbf{y}}} = \left.\begin{pmatrix} \frac{\mathrm{d}\mathbf{z}^\intercal}{\mathrm{d}\mathbf{x}}\frac{\partial \boldsymbol{\epsilon}^\intercal}{\partial \mathbf{z}} \\ \frac{\mathrm{d}\mathbf{z}^\intercal}{\mathrm{d}\mathbf{y}}\frac{\partial \boldsymbol{\epsilon}^\intercal}{\partial \mathbf{z}} \end{pmatrix}\right|_{\mathbf{y}=\bar{\mathbf{y}}} = \left.\left[\begin{pmatrix} \frac{\mathrm{d}\mathbf{z}^\intercal}{\mathrm{d}\mathbf{x}} \\ \frac{\mathrm{d}\mathbf{z}^\intercal}{\mathrm{d}\mathbf{y}} \end{pmatrix}\frac{\partial \boldsymbol{\epsilon}^\intercal}{\partial \mathbf{z}}\right]\right|_{\mathbf{y}=\bar{\mathbf{y}}}$$

$$= \left.\left(\frac{\mathrm{d}\mathbf{z}^\intercal}{\mathrm{d}\mathbf{z}}\frac{\partial \boldsymbol{\epsilon}^\intercal}{\partial \mathbf{z}}\right)\right|_{\mathbf{y}=\bar{\mathbf{y}}},$$

which is Layer 4, Eq. S11, where in the second equality we factorized and in the third equality we used Layer 4, Eq. S8. Using this in Eq. S5.4.2, the first line of Layer 4, Eq. S23 follows.

##### S5.4.2 Matrix of total effects of a mutant's geno-phenotype on her geno-phenotype

Here we obtain a compact expression for $\mathrm{d}\mathbf{z}^\intercal/\mathrm{d}\mathbf{z}|_{\mathbf{y}=\bar{\mathbf{y}}}$. Before doing so, let us obtain the block matrix of *total immediate effects of a mutant's geno-phenotype on her geno-phenotype*

$$\frac{\delta \mathbf{z}^\intercal}{\delta \mathbf{z}} \equiv \begin{pmatrix} \frac{\delta \mathbf{x}^\intercal}{\delta \mathbf{x}} & \frac{\delta \mathbf{y}^\intercal}{\delta \mathbf{x}} \\ \frac{\delta \mathbf{x}^\intercal}{\delta \mathbf{y}} & \frac{\delta \mathbf{y}^\intercal}{\delta \mathbf{y}} \end{pmatrix} = \begin{pmatrix} \frac{\delta \mathbf{x}^\intercal}{\delta \mathbf{x}} & \mathbf{0} \\ \frac{\delta \mathbf{x}^\intercal}{\delta \mathbf{y}} & \mathbf{I} \end{pmatrix} \qquad \text{(Eq. S5.4.3)}$$
$$\in \mathbb{R}^{N_\mathrm{a}(N_\mathrm{p}+N_\mathrm{g})\times N_\mathrm{a}(N_\mathrm{p}+N_\mathrm{g})},$$

where the equality follows from the assumption that genotypic traits are developmentally independent. Using Layer 2, Eq. S6, Layer 2, Eq. S7, and Layer 2, Eq. S9 we have that

$$\frac{\partial \mathbf{z}^\intercal}{\partial \mathbf{z}} + \frac{\partial \boldsymbol{\epsilon}^\intercal}{\partial \mathbf{z}}\frac{\partial \mathbf{z}^\intercal}{\partial \boldsymbol{\epsilon}} = \begin{pmatrix} \frac{\partial \mathbf{x}^\intercal}{\partial \mathbf{x}} & \mathbf{0} \\ \frac{\partial \mathbf{x}^\intercal}{\partial \mathbf{y}} & \mathbf{I} \end{pmatrix} + \begin{pmatrix} \frac{\partial \boldsymbol{\epsilon}^\intercal}{\partial \mathbf{x}} \\ \frac{\partial \boldsymbol{\epsilon}^\intercal}{\partial \mathbf{y}} \end{pmatrix}\begin{pmatrix} \frac{\partial \mathbf{x}^\intercal}{\partial \boldsymbol{\epsilon}} & \mathbf{0} \end{pmatrix}$$

$$= \begin{pmatrix} \frac{\partial \mathbf{x}^\intercal}{\partial \mathbf{x}} & \mathbf{0} \\ \frac{\partial \mathbf{x}^\intercal}{\partial \mathbf{y}} & \mathbf{I} \end{pmatrix} + \begin{pmatrix} \frac{\partial \boldsymbol{\epsilon}^\intercal}{\partial \mathbf{x}}\frac{\partial \mathbf{x}^\intercal}{\partial \boldsymbol{\epsilon}} & \mathbf{0} \\ \frac{\partial \boldsymbol{\epsilon}^\intercal}{\partial \mathbf{y}}\frac{\partial \mathbf{x}^\intercal}{\partial \boldsymbol{\epsilon}} & \mathbf{0} \end{pmatrix}$$

$$= \begin{pmatrix} \frac{\partial \mathbf{x}^\intercal}{\partial \mathbf{x}} + \frac{\partial \boldsymbol{\epsilon}^\intercal}{\partial \mathbf{x}}\frac{\partial \mathbf{x}^\intercal}{\partial \boldsymbol{\epsilon}} & \mathbf{0} \\ \frac{\partial \mathbf{x}^\intercal}{\partial \mathbf{y}} + \frac{\partial \boldsymbol{\epsilon}^\intercal}{\partial \mathbf{y}}\frac{\partial \mathbf{x}^\intercal}{\partial \boldsymbol{\epsilon}} & \mathbf{I} \end{pmatrix},$$

which equals the right-hand side of Eq. S5.4.3 so Layer 3, Eq. S5 holds.

Now, motivated by Layer 4, Eq. S1 and the equation for total effects in path analysis (Greene, 1977), suppose that

$$\frac{\mathrm{d}\mathbf{z}^\intercal}{\mathrm{d}\mathbf{z}} = (\mathbf{I} - \mathbf{E}_\mathbf{z})^{-1},$$

for some matrix $\mathbf{E}_\mathbf{z}$ to be determined. Then,

$$\mathbf{E}_\mathbf{z} = \mathbf{I} - \left(\frac{\mathrm{d}\mathbf{z}^\intercal}{\mathrm{d}\mathbf{z}}\right)^{-1}. \qquad \text{(Eq. S5.4.4)}$$

Using Layer 4, Eq. S8 and a formula for the inverse of a $2\times 2$ block matrix (Horn and Johnson, 2013, Eq. 0.7.3.1), we have

$$\left(\frac{\mathrm{d}\mathbf{z}^\intercal}{\mathrm{d}\mathbf{z}}\right)^{-1} = \begin{pmatrix} \left(\frac{\mathrm{d}\mathbf{x}^\intercal}{\mathrm{d}\mathbf{x}}\right)^{-1} & \mathbf{0} \\ -\frac{\mathrm{d}\mathbf{x}^\intercal}{\mathrm{d}\mathbf{y}}\left(\frac{\mathrm{d}\mathbf{x}^\intercal}{\mathrm{d}\mathbf{x}}\right)^{-1} & \mathbf{I} \end{pmatrix}.$$

Using Layer 4, Eq. S2 yields

$$\left(\frac{\mathrm{d}\mathbf{z}^\intercal}{\mathrm{d}\mathbf{z}}\right)^{-1} = \begin{pmatrix} \left(\frac{\mathrm{d}\mathbf{x}^\intercal}{\mathrm{d}\mathbf{x}}\right)^{-1} & \mathbf{0} \\ -\frac{\delta \mathbf{x}^\intercal}{\delta \mathbf{y}}\frac{\mathrm{d}\mathbf{x}^\intercal}{\mathrm{d}\mathbf{x}}\left(\frac{\mathrm{d}\mathbf{x}^\intercal}{\mathrm{d}\mathbf{x}}\right)^{-1} & \mathbf{I} \end{pmatrix}.$$

Simplifying and using Layer 4, Eq. S1 yields

$$\left(\frac{\mathrm{d}\mathbf{z}^\intercal}{\mathrm{d}\mathbf{z}}\right)^{-1} = \begin{pmatrix} 2\mathbf{I} - \frac{\delta \mathbf{x}^\intercal}{\delta \mathbf{x}} & \mathbf{0} \\ -\frac{\delta \mathbf{x}^\intercal}{\delta \mathbf{y}} & \mathbf{I} \end{pmatrix}.$$

Substituting in Eq. S5.4.4 and simplifying yields

$$\mathbf{E}_\mathbf{z} = \mathbf{I} - \begin{pmatrix} 2\mathbf{I} - \frac{\delta \mathbf{x}^\intercal}{\delta \mathbf{x}} & \mathbf{0} \\ -\frac{\delta \mathbf{x}^\intercal}{\delta \mathbf{y}} & \mathbf{I} \end{pmatrix} = \begin{pmatrix} -\mathbf{I} + \frac{\delta \mathbf{x}^\intercal}{\delta \mathbf{x}} & \mathbf{0} \\ \frac{\delta \mathbf{x}^\intercal}{\delta \mathbf{y}} & \mathbf{0} \end{pmatrix}$$

$$= \begin{pmatrix} \frac{\delta \mathbf{x}^\intercal}{\delta \mathbf{x}} & \mathbf{0} \\ \frac{\delta \mathbf{x}^\intercal}{\delta \mathbf{y}} & \mathbf{I} \end{pmatrix} - \mathbf{I}.$$

Hence,

$$\mathbf{E}_\mathbf{z} = \frac{\delta \mathbf{z}^\intercal}{\delta \mathbf{z}} - \mathbf{I},$$

and so Layer 4, Eq. S8 holds.

#### S5.5 Total selection gradient of the geno-envo-phenotype

We have that the mutant geno-envo-phenotype is $\mathbf{m} = (\mathbf{x};\mathbf{y};\boldsymbol{\epsilon})$. We now define the direct, total immediate, and total selection gradients of the geno-envo-phenotype and write the total selection gradient of the geno-envo-phenotype in terms of the partial selection gradient of the geno-envo-phenotype.

We have the *selection gradient of the geno-envo-phenotype*

$$\left.\frac{\partial w}{\partial \mathbf{m}}\right|_{\mathbf{y}=\bar{\mathbf{y}}} \equiv \left.\left(\frac{\partial w}{\partial \mathbf{x}};\frac{\partial w}{\partial \mathbf{y}};\frac{\partial w}{\partial \boldsymbol{\epsilon}}\right)\right|_{\mathbf{y}=\bar{\mathbf{y}}} \in \mathbb{R}^{N_\mathrm{a}(N_\mathrm{p}+N_\mathrm{g}+N_\mathrm{e})\times 1},$$

the *total immediate selection gradient of the geno-envo-phenotype*

$$\left.\frac{\delta w}{\delta \mathbf{m}}\right|_{\mathbf{y}=\bar{\mathbf{y}}} = \left.\left(\frac{\delta w}{\delta \mathbf{x}};\frac{\delta w}{\delta \mathbf{y}};\frac{\delta w}{\delta \boldsymbol{\epsilon}}\right)\right|_{\mathbf{y}=\bar{\mathbf{y}}} \in \mathbb{R}^{N_\mathrm{a}(N_\mathrm{p}+N_\mathrm{g}+N_\mathrm{e})\times 1},$$

and the *total selection gradient of the geno-envo-phenotype*

$$\left.\frac{\mathrm{d}w}{\mathrm{d}\mathbf{m}}\right|_{\mathbf{y}=\bar{\mathbf{y}}} = \left.\left(\frac{\mathrm{d}w}{\mathrm{d}\mathbf{x}};\frac{\mathrm{d}w}{\mathrm{d}\mathbf{y}};\frac{\mathrm{d}w}{\mathrm{d}\boldsymbol{\epsilon}}\right)\right|_{\mathbf{y}=\bar{\mathbf{y}}} \in \mathbb{R}^{N_\mathrm{a}(N_\mathrm{p}+N_\mathrm{g}+N_\mathrm{e})\times 1}.$$

Now we use the expressions of the total selection gradients of the phenotype, genotype, and environment as linear transformations of the geno-envo-phenotype to write the total selection gradient of the geno-envo-phenotype. Using the fourth lines of Layer 4, Eq. S20 and Layer 4, Eq. S21 and the third line of Layer 4, Eq. S22, we have

$$\left.\frac{\mathrm{d}w}{\mathrm{d}\mathbf{m}}\right|_{\mathbf{y}=\bar{\mathbf{y}}} \equiv \left.\begin{pmatrix}\frac{\mathrm{d}w}{\mathrm{d}\mathbf{x}}\\ \frac{\mathrm{d}w}{\mathrm{d}\mathbf{y}}\\ \frac{\mathrm{d}w}{\mathrm{d}\boldsymbol{\epsilon}}\end{pmatrix}\right|_{\mathbf{y}=\bar{\mathbf{y}}} = \left.\begin{pmatrix}\frac{\mathrm{d}\mathbf{m}^\intercal}{\mathrm{d}\mathbf{x}}\frac{\partial w}{\partial \mathbf{m}}\\ \frac{\mathrm{d}\mathbf{m}^\intercal}{\mathrm{d}\mathbf{y}}\frac{\partial w}{\partial \mathbf{m}}\\ \frac{\mathrm{d}\mathbf{m}^\intercal}{\mathrm{d}\boldsymbol{\epsilon}}\frac{\partial w}{\partial \mathbf{m}}\end{pmatrix}\right|_{\mathbf{y}=\bar{\mathbf{y}}}$$
$$= \left.\left[\begin{pmatrix}\frac{\mathrm{d}\mathbf{m}^\intercal}{\mathrm{d}\mathbf{x}}\\ \frac{\mathrm{d}\mathbf{m}^\intercal}{\mathrm{d}\mathbf{y}}\\ \frac{\mathrm{d}\mathbf{m}^\intercal}{\mathrm{d}\boldsymbol{\epsilon}}\end{pmatrix}\frac{\partial w}{\partial \mathbf{m}}\right]\right|_{\mathbf{y}=\bar{\mathbf{y}}} = \left.\left(\frac{\mathrm{d}\mathbf{m}^\intercal}{\mathrm{d}\mathbf{m}}\frac{\partial w}{\partial \mathbf{m}}\right)\right|_{\mathbf{y}=\bar{\mathbf{y}}},$$

which is Layer 4, Eq. S24.

To see that $\mathrm{d}\mathbf{m}^\intercal/\mathrm{d}\mathbf{m}|_{\mathbf{y}=\bar{\mathbf{y}}}$ is non-singular, we factorize it as follows. We define the block matrix

$$\left.\frac{\gamma\mathbf{m}^\intercal}{\gamma\mathbf{m}}\right|_{\mathbf{y}=\bar{\mathbf{y}}} = \left.\begin{pmatrix}\mathbf{I} & \mathbf{0} & \frac{\partial\boldsymbol{\epsilon}^\intercal}{\partial\mathbf{x}}\\ \mathbf{0} & \mathbf{I} & \frac{\partial\boldsymbol{\epsilon}^\intercal}{\partial\mathbf{y}}\\ \mathbf{0} & \mathbf{0} & \frac{\partial\boldsymbol{\epsilon}^\intercal}{\partial\boldsymbol{\epsilon}}\end{pmatrix}\right|_{\mathbf{y}=\bar{\mathbf{y}}}$$
$$\in \mathbb{R}^{N_\mathrm{a}(N_\mathrm{p}+N_\mathrm{g}+N_\mathrm{e})\times N_\mathrm{a}(N_\mathrm{p}+N_\mathrm{g}+N_\mathrm{e})},$$

which is non-singular since it is square, block upper triangular, and $\partial\boldsymbol{\epsilon}^\intercal/\partial\boldsymbol{\epsilon} = \mathbf{I}$ (Layer 2, Eq. S3). We also define the block matrix of

$$\left.\frac{\beta\mathbf{m}^\intercal}{\beta\mathbf{m}}\right|_{\mathbf{y}=\bar{\mathbf{y}}} = \left.\begin{pmatrix}\frac{\mathrm{d}\mathbf{x}^\intercal}{\mathrm{d}\mathbf{x}} & \mathbf{0} & \mathbf{0}\\ \frac{\mathrm{d}\mathbf{x}^\intercal}{\mathrm{d}\mathbf{y}} & \mathbf{I} & \mathbf{0}\\ \frac{\mathrm{d}\mathbf{x}^\intercal}{\mathrm{d}\boldsymbol{\epsilon}} & \mathbf{0} & \mathbf{I}\end{pmatrix}\right|_{\mathbf{y}=\bar{\mathbf{y}}}$$
$$\in \mathbb{R}^{N_\mathrm{a}(N_\mathrm{p}+N_\mathrm{g}+N_\mathrm{e})\times N_\mathrm{a}(N_\mathrm{p}+N_\mathrm{g}+N_\mathrm{e})},$$

which is non-singular since it is square, block lower triangular, and $\mathrm{d}\mathbf{x}^\intercal/\mathrm{d}\mathbf{x}$ is non-singular (Eq. S5.1.15). Note that

$$\left.\left(\frac{\beta\mathbf{m}^\intercal}{\beta\mathbf{m}}\frac{\gamma\mathbf{m}^\intercal}{\gamma\mathbf{m}}\right)\right|_{\mathbf{y}=\bar{\mathbf{y}}} = \left.\left[\begin{pmatrix}\frac{\mathrm{d}\mathbf{x}^\intercal}{\mathrm{d}\mathbf{x}} & \mathbf{0} & \mathbf{0}\\ \frac{\mathrm{d}\mathbf{x}^\intercal}{\mathrm{d}\mathbf{y}} & \mathbf{I} & \mathbf{0}\\ \frac{\mathrm{d}\mathbf{x}^\intercal}{\mathrm{d}\boldsymbol{\epsilon}} & \mathbf{0} & \mathbf{I}\end{pmatrix}\begin{pmatrix}\mathbf{I} & \mathbf{0} & \frac{\partial\boldsymbol{\epsilon}^\intercal}{\partial\mathbf{x}}\\ \mathbf{0} & \mathbf{I} & \frac{\partial\boldsymbol{\epsilon}^\intercal}{\partial\mathbf{y}}\\ \mathbf{0} & \mathbf{0} & \frac{\partial\boldsymbol{\epsilon}^\intercal}{\partial\boldsymbol{\epsilon}}\end{pmatrix}\right]\right|_{\mathbf{y}=\bar{\mathbf{y}}}$$
$$= \left.\begin{pmatrix}\frac{\mathrm{d}\mathbf{x}^\intercal}{\mathrm{d}\mathbf{x}} & \mathbf{0} & \frac{\mathrm{d}\mathbf{x}^\intercal}{\mathrm{d}\mathbf{x}}\frac{\partial\boldsymbol{\epsilon}^\intercal}{\partial\mathbf{x}}\\ \frac{\mathrm{d}\mathbf{x}^\intercal}{\mathrm{d}\mathbf{y}} & \mathbf{I} & \frac{\mathrm{d}\mathbf{x}^\intercal}{\mathrm{d}\mathbf{y}}\frac{\partial\boldsymbol{\epsilon}^\intercal}{\partial\mathbf{x}} + \frac{\partial\boldsymbol{\epsilon}^\intercal}{\partial\mathbf{y}}\\ \frac{\mathrm{d}\mathbf{x}^\intercal}{\mathrm{d}\boldsymbol{\epsilon}} & \mathbf{0} & \frac{\mathrm{d}\mathbf{x}^\intercal}{\mathrm{d}\boldsymbol{\epsilon}}\frac{\partial\boldsymbol{\epsilon}^\intercal}{\partial\mathbf{x}} + \frac{\partial\boldsymbol{\epsilon}^\intercal}{\partial\boldsymbol{\epsilon}}\end{pmatrix}\right|_{\mathbf{y}=\bar{\mathbf{y}}}$$
$$= \left.\begin{pmatrix}\frac{\mathrm{d}\mathbf{x}^\intercal}{\mathrm{d}\mathbf{x}} & \mathbf{0} & \frac{\mathrm{d}\boldsymbol{\epsilon}^\intercal}{\mathrm{d}\mathbf{x}}\\ \frac{\mathrm{d}\mathbf{x}^\intercal}{\mathrm{d}\mathbf{y}} & \mathbf{I} & \frac{\mathrm{d}\boldsymbol{\epsilon}^\intercal}{\mathrm{d}\mathbf{y}}\\ \frac{\mathrm{d}\mathbf{x}^\intercal}{\mathrm{d}\boldsymbol{\epsilon}} & \mathbf{0} & \frac{\mathrm{d}\boldsymbol{\epsilon}^\intercal}{\mathrm{d}\boldsymbol{\epsilon}}\end{pmatrix}\right|_{\mathbf{y}=\bar{\mathbf{y}}},$$

where the last equality follows from Layer 4, Eq. S9, Layer 4, Eq. S10, and Layer 4, Eq. S12. Using Layer 4, Eq. S17, we thus have that

$$\left.\frac{\mathrm{d}\mathbf{m}^\intercal}{\mathrm{d}\mathbf{m}}\right|_{\mathbf{y}=\bar{\mathbf{y}}} = \left.\left(\frac{\beta\mathbf{m}^\intercal}{\beta\mathbf{m}}\frac{\gamma\mathbf{m}^\intercal}{\gamma\mathbf{m}}\right)\right|_{\mathbf{y}=\bar{\mathbf{y}}}.$$

Hence, $\mathrm{d}\mathbf{m}^\intercal/\mathrm{d}\mathbf{m}|_{\mathbf{y}=\bar{\mathbf{y}}}$ is non-singular since $\beta\mathbf{m}^\intercal/\beta\mathbf{m}|_{\mathbf{y}=\bar{\mathbf{y}}}$ and $\gamma\mathbf{m}^\intercal/\gamma\mathbf{m}|_{\mathbf{y}=\bar{\mathbf{y}}}$ are square and non-singular.

#### S5.6 Evolutionary dynamics of the phenotype

Here we derive equations describing the evolutionary dynamics of the resident phenotype.

From Eq. S2.2.2 and Eq. S2.3.9, we have that the evolutionary dynamics of the resident genotype satisfy the canonical equation

$$\frac{\Delta\bar{\mathbf{y}}}{\Delta\tau} \approx \iota\mathbf{H}_\mathbf{y}\left.\frac{\mathrm{d}w}{\mathrm{d}\mathbf{y}}\right|_{\mathbf{y}=\bar{\mathbf{y}}}, \qquad \text{(Eq. S5.6.1)}$$

whereas the developmental dynamics of the resident phenotype satisfy the developmental constraint

$$\bar{\mathbf{x}}_{a+1} = \mathbf{g}_a(\bar{\mathbf{m}}_a,\bar{\mathbf{z}})$$

for all $a \in \{1,\ldots,N_\mathrm{a}-1\}$ with $\bar{\mathbf{x}}_1$ constant, and where the resident environment is given by

$$\bar{\boldsymbol{\epsilon}}_a = \mathbf{h}_a(\bar{\mathbf{z}}_a, \bar{\mathbf{z}}, \tau)$$

for all $a \in \{1,\ldots,N_\mathrm{a}\}$.

Let $\bar{\mathbf{z}}(\tau)$ be the resident geno-phenotype at evolutionary time $\tau$, specifically at the point where the socio-devo stable resident is at carrying capacity, marked in Fig. 3. The $i$-th mutant phenotype at age $j+1$ at such evolutionary time $\tau$ is $x_{i,j+1} = g_{ij}(\mathbf{z}_j(\tau), \mathbf{h}_j(\mathbf{z}_j(\tau), \bar{\mathbf{z}}(\tau), \tau), \bar{\mathbf{z}}(\tau))$. Then, evolutionary change in the $i$-th resident phenotype at age $a \in \{2,\ldots,N_\mathrm{a}\}$ is

$$\begin{aligned}\frac{\Delta \bar{x}_{ia}}{\Delta \tau} = \frac{1}{\Delta\tau}\Bigg[ g_{i,a-1}\Big(&\mathbf{z}_{a-1}(\tau+\Delta\tau),\\ &\mathbf{h}_{a-1}(\mathbf{z}_{a-1}(\tau+\Delta\tau), \bar{\mathbf{z}}(\tau+\Delta\tau), \tau+\Delta\tau),\\ &\bar{\mathbf{z}}(\tau+\Delta\tau)\Big)\\ &- g_{i,a-1}(\mathbf{z}_{a-1}(\tau), \mathbf{h}_{a-1}(\mathbf{z}_{a-1}(\tau), \bar{\mathbf{z}}(\tau), \tau), \bar{\mathbf{z}}(\tau))\Bigg]\Bigg|_{\mathbf{y}=\bar{\mathbf{y}}}.\end{aligned}$$

Taking the limit as $\Delta\tau \to 0$, this becomes

$$\frac{\mathrm{d}\bar{x}_{ia}}{\mathrm{d}\tau} = \frac{\mathrm{d}g_{i,a-1}(\mathbf{z}_{a-1}(\tau), \mathbf{h}_{a-1}(\mathbf{z}_{a-1}(\tau), \bar{\mathbf{z}}(\tau), \tau), \bar{\mathbf{z}}(\tau))}{\mathrm{d}\tau}\Bigg|_{\mathbf{y}=\bar{\mathbf{y}}}.$$

Applying the chain rule, we obtain

$$\begin{aligned}\frac{\mathrm{d}\bar{x}_{ia}}{\mathrm{d}\tau} = &\\ \Bigg(&\sum_{j=1}^{N_\mathrm{p}} \frac{\partial g_{i,a-1}}{\partial x_{j,a-1}} \frac{\mathrm{d}x_{j,a-1}}{\mathrm{d}\tau} + \sum_{j=1}^{N_\mathrm{g}} \frac{\partial g_{i,a-1}}{\partial y_{j,a-1}} \frac{\mathrm{d}y_{j,a-1}}{\mathrm{d}\tau}\\ &+ \sum_{j=1}^{N_\mathrm{p}} \sum_{r=1}^{N_\mathrm{e}} \frac{\partial g_{i,a-1}}{\partial \epsilon_{r,a-1}} \frac{\partial \epsilon_{r,a-1}}{\partial x_{j,a-1}} \frac{\mathrm{d}x_{j,a-1}}{\mathrm{d}\tau}\\ &+ \sum_{j=1}^{N_\mathrm{g}} \sum_{r=1}^{N_\mathrm{e}} \frac{\partial g_{i,a-1}}{\partial \epsilon_{r,a-1}} \frac{\partial \epsilon_{r,a-1}}{\partial y_{j,a-1}} \frac{\mathrm{d}y_{j,a-1}}{\mathrm{d}\tau}\\ &+ \sum_{k=1}^{N_\mathrm{a}} \sum_{j=1}^{N_\mathrm{p}} \sum_{r=1}^{N_\mathrm{e}} \frac{\partial g_{i,a-1}}{\partial \epsilon_{r,a-1}} \frac{\partial \epsilon_{r,a-1}}{\partial \bar{x}_{jk}} \frac{\mathrm{d}\bar{x}_{jk}}{\mathrm{d}\tau}\\ &+ \sum_{k=1}^{N_\mathrm{a}} \sum_{j=1}^{N_\mathrm{g}} \sum_{r=1}^{N_\mathrm{e}} \frac{\partial g_{i,a-1}}{\partial \epsilon_{r,a-1}} \frac{\partial \epsilon_{r,a-1}}{\partial \bar{y}_{jk}} \frac{\mathrm{d}\bar{y}_{jk}}{\mathrm{d}\tau}\\ &+ \sum_{r=1}^{N_\mathrm{e}} \frac{\partial g_{i,a-1}}{\partial \epsilon_{r,a-1}} \frac{\partial \epsilon_{r,a-1}}{\partial \tau}\\ &+ \sum_{k=1}^{N_\mathrm{a}} \sum_{j=1}^{N_\mathrm{p}} \frac{\partial g_{i,a-1}}{\partial \bar{x}_{jk}} \frac{\mathrm{d}\bar{x}_{jk}}{\mathrm{d}\tau} + \sum_{k=1}^{N_\mathrm{a}} \sum_{j=1}^{N_\mathrm{g}} \frac{\partial g_{i,a-1}}{\partial \bar{y}_{jk}} \frac{\mathrm{d}\bar{y}_{jk}}{\mathrm{d}\tau}\Bigg)\Bigg|_{\mathbf{y}=\bar{\mathbf{y}}}.\end{aligned}$$

Applying matrix calculus notation (Appendix A), this is

$$\begin{aligned}\frac{\mathrm{d}\bar{x}_{ia}}{\mathrm{d}\tau} = &\\ \Bigg(&\frac{\partial g_{i,a-1}}{\partial \mathbf{x}_{a-1}^\intercal} \frac{\mathrm{d}\mathbf{x}_{a-1}}{\mathrm{d}\tau} + \frac{\partial g_{i,a-1}}{\partial \mathbf{y}_{a-1}^\intercal} \frac{\mathrm{d}\mathbf{y}_{a-1}}{\mathrm{d}\tau}\\ &+ \sum_{j=1}^{N_\mathrm{p}} \frac{\partial g_{i,a-1}}{\partial \boldsymbol{\epsilon}_{a-1}^\intercal} \frac{\partial \boldsymbol{\epsilon}_{a-1}}{\partial x_{j,a-1}} \frac{\mathrm{d}x_{j,a-1}}{\mathrm{d}\tau}\\ &+ \sum_{j=1}^{N_\mathrm{g}} \frac{\partial g_{i,a-1}}{\partial \boldsymbol{\epsilon}_{a-1}^\intercal} \frac{\partial \boldsymbol{\epsilon}_{a-1}}{\partial y_{j,a-1}} \frac{\mathrm{d}y_{j,a-1}}{\mathrm{d}\tau}\\ &+ \sum_{k=1}^{N_\mathrm{a}} \sum_{j=1}^{N_\mathrm{p}} \frac{\partial g_{i,a-1}}{\partial \boldsymbol{\epsilon}_{a-1}^\intercal} \frac{\partial \boldsymbol{\epsilon}_{a-1}}{\partial \bar{x}_{jk}} \frac{\mathrm{d}\bar{x}_{jk}}{\mathrm{d}\tau}\\ &+ \sum_{k=1}^{N_\mathrm{a}} \sum_{j=1}^{N_\mathrm{g}} \frac{\partial g_{i,a-1}}{\partial \boldsymbol{\epsilon}_{a-1}^\intercal} \frac{\partial \boldsymbol{\epsilon}_{a-1}}{\partial \bar{y}_{jk}} \frac{\mathrm{d}\bar{y}_{jk}}{\mathrm{d}\tau}\\ &+ \frac{\partial g_{i,a-1}}{\partial \boldsymbol{\epsilon}_{a-1}^\intercal} \frac{\partial \boldsymbol{\epsilon}_{a-1}}{\partial \tau}\\ &+ \sum_{k=1}^{N_\mathrm{a}} \frac{\partial g_{i,a-1}}{\partial \bar{\mathbf{x}}_k^\intercal} \frac{\mathrm{d}\bar{\mathbf{x}}_k}{\mathrm{d}\tau} + \sum_{k=1}^{N_\mathrm{a}} \frac{\partial g_{i,a-1}}{\partial \bar{\mathbf{y}}_k^\intercal} \frac{\mathrm{d}\bar{\mathbf{y}}_k}{\mathrm{d}\tau}\Bigg)\Bigg|_{\mathbf{y}=\bar{\mathbf{y}}}.\end{aligned}$$

Applying matrix calculus notation again yields

$$\begin{aligned}\frac{\mathrm{d}\bar{x}_{ia}}{\mathrm{d}\tau} = &\\ \Bigg(&\frac{\partial g_{i,a-1}}{\partial \mathbf{x}_{a-1}^\intercal} \frac{\mathrm{d}\mathbf{x}_{a-1}}{\mathrm{d}\tau} + \frac{\partial g_{i,a-1}}{\partial \mathbf{y}_{a-1}^\intercal} \frac{\mathrm{d}\mathbf{y}_{a-1}}{\mathrm{d}\tau}\\ &+ \frac{\partial g_{i,a-1}}{\partial \boldsymbol{\epsilon}_{a-1}^\intercal} \frac{\partial \boldsymbol{\epsilon}_{a-1}}{\partial \mathbf{x}_{a-1}^\intercal} \frac{\mathrm{d}\mathbf{x}_{a-1}}{\mathrm{d}\tau} + \frac{\partial g_{i,a-1}}{\partial \boldsymbol{\epsilon}_{a-1}^\intercal} \frac{\partial \boldsymbol{\epsilon}_{a-1}}{\partial \mathbf{y}_{a-1}^\intercal} \frac{\mathrm{d}\mathbf{y}_{a-1}}{\mathrm{d}\tau}\\ &+ \frac{\partial g_{i,a-1}}{\partial \boldsymbol{\epsilon}_{a-1}^\intercal} \frac{\partial \boldsymbol{\epsilon}_{a-1}}{\partial \bar{\mathbf{x}}^\intercal} \frac{\mathrm{d}\bar{\mathbf{x}}}{\mathrm{d}\tau} + \frac{\partial g_{i,a-1}}{\partial \boldsymbol{\epsilon}_{a-1}^\intercal} \frac{\partial \boldsymbol{\epsilon}_{a-1}}{\partial \bar{\mathbf{y}}^\intercal} \frac{\mathrm{d}\bar{\mathbf{y}}}{\mathrm{d}\tau}\\ &+ \frac{\partial g_{i,a-1}}{\partial \boldsymbol{\epsilon}_{a-1}^\intercal} \frac{\partial \boldsymbol{\epsilon}_{a-1}}{\partial \tau}\\ &+ \frac{\partial g_{i,a-1}}{\partial \bar{\mathbf{x}}^\intercal} \frac{\mathrm{d}\bar{\mathbf{x}}}{\mathrm{d}\tau} + \frac{\partial g_{i,a-1}}{\partial \bar{\mathbf{y}}^\intercal} \frac{\mathrm{d}\bar{\mathbf{y}}}{\mathrm{d}\tau}\Bigg)\Bigg|_{\mathbf{y}=\bar{\mathbf{y}}}.\end{aligned}$$

Factorizing, we have

$$\begin{aligned}\frac{\mathrm{d}\bar{x}_{ia}}{\mathrm{d}\tau} = &\\ \Bigg[&\left(\frac{\partial g_{i,a-1}}{\partial \mathbf{x}_{a-1}^\intercal} + \frac{\partial g_{i,a-1}}{\partial \boldsymbol{\epsilon}_{a-1}^\intercal} \frac{\partial \boldsymbol{\epsilon}_{a-1}}{\partial \mathbf{x}_{a-1}^\intercal}\right) \frac{\mathrm{d}\mathbf{x}_{a-1}}{\mathrm{d}\tau}\\ &+ \left(\frac{\partial g_{i,a-1}}{\partial \mathbf{y}_{a-1}^\intercal} + \frac{\partial g_{i,a-1}}{\partial \boldsymbol{\epsilon}_{a-1}^\intercal} \frac{\partial \boldsymbol{\epsilon}_{a-1}}{\partial \mathbf{y}_{a-1}^\intercal}\right) \frac{\mathrm{d}\mathbf{y}_{a-1}}{\mathrm{d}\tau}\\ &+ \left(\frac{\partial g_{i,a-1}}{\partial \bar{\mathbf{x}}^\intercal} + \frac{\partial g_{i,a-1}}{\partial \boldsymbol{\epsilon}_{a-1}^\intercal} \frac{\partial \boldsymbol{\epsilon}_{a-1}}{\partial \bar{\mathbf{x}}^\intercal}\right) \frac{\mathrm{d}\bar{\mathbf{x}}}{\mathrm{d}\tau}\\ &+ \left(\frac{\partial g_{i,a-1}}{\partial \bar{\mathbf{y}}^\intercal} + \frac{\partial g_{i,a-1}}{\partial \boldsymbol{\epsilon}_{a-1}^\intercal} \frac{\partial \boldsymbol{\epsilon}_{a-1}}{\partial \bar{\mathbf{y}}^\intercal}\right) \frac{\mathrm{d}\bar{\mathbf{y}}}{\mathrm{d}\tau} + \frac{\partial g_{i,a-1}}{\partial \boldsymbol{\epsilon}_{a-1}^\intercal} \frac{\partial \boldsymbol{\epsilon}_{a-1}}{\partial \tau}\Bigg]\Bigg|_{\mathbf{y}=\bar{\mathbf{y}}}.\end{aligned}$$

Rewriting $g_{i,a-1}$ as $x_{ia}$ yields

$$\begin{aligned}\frac{\mathrm{d}\bar{x}_{ia}}{\mathrm{d}\tau} = &\\ \Bigg[&\left(\frac{\partial x_{ia}}{\partial \mathbf{x}_{a-1}^\intercal} + \frac{\partial x_{ia}}{\partial \boldsymbol{\epsilon}_{a-1}^\intercal} \frac{\partial \boldsymbol{\epsilon}_{a-1}}{\partial \mathbf{x}_{a-1}^\intercal}\right) \frac{\mathrm{d}\mathbf{x}_{a-1}}{\mathrm{d}\tau}\\ &+ \left(\frac{\partial x_{ia}}{\partial \mathbf{y}_{a-1}^\intercal} + \frac{\partial x_{ia}}{\partial \boldsymbol{\epsilon}_{a-1}^\intercal} \frac{\partial \boldsymbol{\epsilon}_{a-1}}{\partial \mathbf{y}_{a-1}^\intercal}\right) \frac{\mathrm{d}\mathbf{y}_{a-1}}{\mathrm{d}\tau}\\ &+ \left(\frac{\partial x_{ia}}{\partial \bar{\mathbf{x}}^\intercal} + \frac{\partial x_{ia}}{\partial \boldsymbol{\epsilon}_{a-1}^\intercal} \frac{\partial \boldsymbol{\epsilon}_{a-1}}{\partial \bar{\mathbf{x}}^\intercal}\right) \frac{\mathrm{d}\bar{\mathbf{x}}}{\mathrm{d}\tau}\\ &+ \left(\frac{\partial x_{ia}}{\partial \bar{\mathbf{y}}^\intercal} + \frac{\partial x_{ia}}{\partial \boldsymbol{\epsilon}_{a-1}^\intercal} \frac{\partial \boldsymbol{\epsilon}_{a-1}}{\partial \bar{\mathbf{y}}^\intercal}\right) \frac{\mathrm{d}\bar{\mathbf{y}}}{\mathrm{d}\tau} + \frac{\partial x_{ia}}{\partial \boldsymbol{\epsilon}_{a-1}^\intercal} \frac{\partial \boldsymbol{\epsilon}_{a-1}}{\partial \tau}\Bigg]\Bigg|_{\mathbf{y}=\bar{\mathbf{y}}}.\end{aligned}$$

Hence, for all resident phenotypes at age $a \in \{2,\ldots,N_\mathrm{a}\}$, we have

$$\frac{\mathrm{d}\bar{\mathbf{x}}_a}{\mathrm{d}\tau} = \Bigg[\left(\frac{\partial \mathbf{x}_a}{\partial \mathbf{x}_{a-1}^\intercal} + \frac{\partial \mathbf{x}_a}{\partial \boldsymbol{\epsilon}_{a-1}^\intercal}\frac{\partial \boldsymbol{\epsilon}_{a-1}}{\partial \mathbf{x}_{a-1}^\intercal}\right)\frac{\mathrm{d}\mathbf{x}_{a-1}}{\mathrm{d}\tau} + \left(\frac{\partial \mathbf{x}_a}{\partial \mathbf{y}_{a-1}^\intercal} + \frac{\partial \mathbf{x}_a}{\partial \boldsymbol{\epsilon}_{a-1}^\intercal}\frac{\partial \boldsymbol{\epsilon}_{a-1}}{\partial \mathbf{y}_{a-1}^\intercal}\right)\frac{\mathrm{d}\mathbf{y}_{a-1}}{\mathrm{d}\tau} + \left(\frac{\partial \mathbf{x}_a}{\partial \bar{\mathbf{x}}^\intercal} + \frac{\partial \mathbf{x}_a}{\partial \boldsymbol{\epsilon}_{a-1}^\intercal}\frac{\partial \boldsymbol{\epsilon}_{a-1}}{\partial \bar{\mathbf{x}}^\intercal}\right)\frac{\mathrm{d}\bar{\mathbf{x}}}{\mathrm{d}\tau} + \left(\frac{\partial \mathbf{x}_a}{\partial \bar{\mathbf{y}}^\intercal} + \frac{\partial \mathbf{x}_a}{\partial \boldsymbol{\epsilon}_{a-1}^\intercal}\frac{\partial \boldsymbol{\epsilon}_{a-1}}{\partial \bar{\mathbf{y}}^\intercal}\right)\frac{\mathrm{d}\bar{\mathbf{y}}}{\mathrm{d}\tau} + \frac{\partial \mathbf{x}_a}{\partial \boldsymbol{\epsilon}_{a-1}^\intercal}\frac{\partial \boldsymbol{\epsilon}_{a-1}}{\partial \tau}\Bigg]\Bigg|_{\mathbf{y}=\bar{\mathbf{y}}}. \tag{Eq. S5.6.2}$$

Here we used the following series of definitions. The matrix of *direct effects of social partner's phenotype at age a on the mutant's phenotype at age j* is

$$\left.\frac{\partial \mathbf{x}_j^\intercal}{\partial \bar{\mathbf{x}}_a}\right|_{\mathbf{y}=\bar{\mathbf{y}}} \equiv \left.\begin{pmatrix} \frac{\partial x_{1j}}{\partial \bar{x}_{1a}} & \cdots & \frac{\partial x_{N_\mathrm{p}j}}{\partial \bar{x}_{1a}} \\ \vdots & \ddots & \vdots \\ \frac{\partial x_{1j}}{\partial \bar{x}_{N_\mathrm{p}a}} & \cdots & \frac{\partial x_{N_\mathrm{p}j}}{\partial \bar{x}_{N_\mathrm{p}a}} \end{pmatrix}\right|_{\mathbf{y}=\bar{\mathbf{y}}} \in \mathbb{R}^{N_\mathrm{p}\times N_\mathrm{p}},$$

and the block matrix of direct effects of social partners' phenotype on a mutant's phenotype is given by Layer 2, Eq. S4 with $\bar{\boldsymbol{\zeta}} = \bar{\mathbf{x}}$. The matrix $\partial \mathbf{x}_a^\intercal/\partial \bar{\mathbf{x}}$ is the $a$-th block column of $\partial \mathbf{x}^\intercal/\partial \bar{\mathbf{x}}$.

Similarly, the matrix of *direct effects of social partners' genotypic trait values at age a on a mutant's phenotype at age j* is

$$\left.\frac{\partial \mathbf{x}_j^\intercal}{\partial \bar{\mathbf{y}}_a}\right|_{\mathbf{y}=\bar{\mathbf{y}}} \equiv \left.\begin{pmatrix} \frac{\partial x_{1j}}{\partial \bar{y}_{1a}} & \cdots & \frac{\partial x_{N_\mathrm{p}j}}{\partial \bar{y}_{1a}} \\ \vdots & \ddots & \vdots \\ \frac{\partial x_{1j}}{\partial \bar{y}_{N_\mathrm{g}a}} & \cdots & \frac{\partial x_{N_\mathrm{p}j}}{\partial \bar{y}_{N_\mathrm{g}a}} \end{pmatrix}\right|_{\mathbf{y}=\bar{\mathbf{y}}} \in \mathbb{R}^{N_\mathrm{g}\times N_\mathrm{p}},$$

and the block matrix of direct effects of social partners' genotype on a mutant's phenotype is given by Layer 2, Eq. S4 with $\bar{\boldsymbol{\zeta}} = \bar{\mathbf{y}}$. The matrix $\partial \mathbf{x}_a^\intercal/\partial \bar{\mathbf{y}}$ is the $a$-th block column of $\partial \mathbf{x}^\intercal/\partial \bar{\mathbf{y}}$.

In turn, the matrix of *direct effects of social partners' phenotype at age a on a mutant's environment at age j* is

$$\left.\frac{\partial \boldsymbol{\epsilon}_j^\intercal}{\partial \bar{\mathbf{x}}_a}\right|_{\mathbf{y}=\bar{\mathbf{y}}} \equiv \left.\begin{pmatrix} \frac{\partial \epsilon_{1j}}{\partial \bar{x}_{1a}} & \cdots & \frac{\partial \epsilon_{N_\mathrm{e}j}}{\partial \bar{x}_{1a}} \\ \vdots & \ddots & \vdots \\ \frac{\partial \epsilon_{1j}}{\partial \bar{x}_{N_\mathrm{p}a}} & \cdots & \frac{\partial \epsilon_{N_\mathrm{e}j}}{\partial \bar{x}_{N_\mathrm{p}a}} \end{pmatrix}\right|_{\mathbf{y}=\bar{\mathbf{y}}} \in \mathbb{R}^{N_\mathrm{e}\times N_\mathrm{p}},$$

and the block matrix of direct effects of social partners' phenotype on a mutant's environment is given by Layer 2, Eq. S5 with $\bar{\boldsymbol{\zeta}} = \bar{\mathbf{x}}$. The matrix $\partial \boldsymbol{\epsilon}_a^\intercal/\partial \bar{\mathbf{x}}$ is the $a$-th block column of $\partial \boldsymbol{\epsilon}^\intercal/\partial \bar{\mathbf{x}}$.

Similarly, the matrix of *direct effects of social partners' genotypic trait values at age a on a mutant's environment at age j* is

$$\left.\frac{\partial \boldsymbol{\epsilon}_j^\intercal}{\partial \bar{\mathbf{y}}_a}\right|_{\mathbf{y}=\bar{\mathbf{y}}} \equiv \left.\begin{pmatrix} \frac{\partial \epsilon_{1j}}{\partial \bar{y}_{1a}} & \cdots & \frac{\partial \epsilon_{N_\mathrm{e}j}}{\partial \bar{y}_{1a}} \\ \vdots & \ddots & \vdots \\ \frac{\partial \epsilon_{1j}}{\partial \bar{y}_{N_\mathrm{g}a}} & \cdots & \frac{\partial \epsilon_{N_\mathrm{e}j}}{\partial \bar{y}_{N_\mathrm{g}a}} \end{pmatrix}\right|_{\mathbf{y}=\bar{\mathbf{y}}} \in \mathbb{R}^{N_\mathrm{e}\times N_\mathrm{g}},$$

and the block matrix of *direct effects of social partners' genotype on a mutant's environment* is given by Layer 2, Eq. S5 with $\bar{\boldsymbol{\zeta}} = \bar{\mathbf{y}}$. The matrix $\partial \boldsymbol{\epsilon}_a^\intercal/\partial \bar{\mathbf{y}}$ is the $a$-th block column of $\partial \boldsymbol{\epsilon}^\intercal/\partial \bar{\mathbf{y}}$.

Having made these definitions explicit, we now write Eq. S5.6.2 as

$$\frac{\mathrm{d}\bar{\mathbf{x}}_a}{\mathrm{d}\tau} = \left(\frac{\delta \mathbf{x}_a}{\delta \mathbf{x}_{a-1}^\intercal}\frac{\mathrm{d}\mathbf{x}_{a-1}}{\mathrm{d}\tau} + \frac{\delta \mathbf{x}_a}{\delta \mathbf{y}_{a-1}^\intercal}\frac{\mathrm{d}\mathbf{y}_{a-1}}{\mathrm{d}\tau} + \frac{\delta \mathbf{x}_a}{\delta \bar{\mathbf{x}}^\intercal}\frac{\mathrm{d}\bar{\mathbf{x}}}{\mathrm{d}\tau} + \frac{\delta \mathbf{x}_a}{\delta \bar{\mathbf{y}}^\intercal}\frac{\mathrm{d}\bar{\mathbf{y}}}{\mathrm{d}\tau} + \frac{\delta \mathbf{x}_a}{\delta \boldsymbol{\epsilon}_{a-1}^\intercal}\frac{\partial \boldsymbol{\epsilon}_{a-1}}{\partial \tau}\right)\Bigg|_{\mathbf{y}=\bar{\mathbf{y}}}, \tag{Eq. S5.6.3}$$

where we used the transpose of the total immediate effects of a mutant's phenotype and genotype on her phenotype (Eq. S5.1.10 and Eq. S5.2.9), and the the matrix of *total immediate effects of social partners' phenotype or genotype at age a on a mutant's phenotype at age j*

$$\left.\frac{\delta \mathbf{x}_j^\intercal}{\delta \bar{\boldsymbol{\zeta}}_a}\right|_{\mathbf{y}=\bar{\mathbf{y}}} = \begin{cases} \left.\left(\frac{\partial \mathbf{x}_j^\intercal}{\partial \bar{\boldsymbol{\zeta}}_a} + \frac{\partial \boldsymbol{\epsilon}_{j-1}^\intercal}{\partial \bar{\boldsymbol{\zeta}}_a}\frac{\partial \mathbf{x}_j^\intercal}{\partial \boldsymbol{\epsilon}_{j-1}}\right)\right|_{\mathbf{y}=\bar{\mathbf{y}}} & \text{for } j>1 \\ \mathbf{0} & \text{for } j=1, \end{cases} \tag{Eq. S5.6.4}$$

for $\bar{\boldsymbol{\zeta}} \in \{\bar{\mathbf{x}}, \bar{\mathbf{y}}\}$ since the initial phenotype $\mathbf{x}_1$ is constant by assumption. We also define the corresponding matrix of *total immediate effects of social partners' phenotype on a mutant's phenotype* as

$$\left.\frac{\delta \mathbf{x}^\intercal}{\delta \bar{\boldsymbol{\zeta}}}\right|_{\mathbf{y}=\bar{\mathbf{y}}} \equiv \left.\begin{pmatrix} \frac{\delta \mathbf{x}_1^\intercal}{\delta \bar{\boldsymbol{\zeta}}_1} & \cdots & \frac{\delta \mathbf{x}_{N_\mathrm{a}}^\intercal}{\delta \bar{\boldsymbol{\zeta}}_1} \\ \vdots & \ddots & \vdots \\ \frac{\delta \mathbf{x}_1^\intercal}{\delta \bar{\boldsymbol{\zeta}}_{N_\mathrm{a}}} & \cdots & \frac{\delta \mathbf{x}_{N_\mathrm{a}}^\intercal}{\delta \bar{\boldsymbol{\zeta}}_{N_\mathrm{a}}} \end{pmatrix}\right|_{\mathbf{y}=\bar{\mathbf{y}}} = \left.\begin{pmatrix} \mathbf{0} & \frac{\delta \mathbf{x}_2^\intercal}{\delta \bar{\boldsymbol{\zeta}}_1} & \cdots & \frac{\delta \mathbf{x}_{N_\mathrm{a}}^\intercal}{\delta \bar{\boldsymbol{\zeta}}_1} \\ \mathbf{0} & \frac{\delta \mathbf{x}_2^\intercal}{\delta \bar{\boldsymbol{\zeta}}_1} & \cdots & \frac{\delta \mathbf{x}_{N_\mathrm{a}}^\intercal}{\delta \bar{\boldsymbol{\zeta}}_1} \\ \vdots & \vdots & \ddots & \vdots \\ \mathbf{0} & \frac{\delta \mathbf{x}_2^\intercal}{\delta \bar{\boldsymbol{\zeta}}_{N_\mathrm{a}}} & \cdots & \frac{\delta \mathbf{x}_{N_\mathrm{a}}^\intercal}{\delta \bar{\boldsymbol{\zeta}}_{N_\mathrm{a}}} \end{pmatrix}\right|_{\mathbf{y}=\bar{\mathbf{y}}}, \tag{Eq. S5.6.5}$$

for $\bar{\boldsymbol{\zeta}} \in \{\bar{\mathbf{x}}, \bar{\mathbf{y}}\}$. The matrix $\delta \mathbf{x}_a^\intercal/\delta \bar{\boldsymbol{\zeta}}$ is the $a$-th block column of $\delta \mathbf{x}^\intercal/\delta \bar{\boldsymbol{\zeta}}$. Using Layer 2, Eq. S2c and since the initial phenotype $\mathbf{x}_1$ is constant by assumption, we have that

$$\frac{\partial \boldsymbol{\epsilon}^\intercal}{\partial \bar{\boldsymbol{\zeta}}}\frac{\partial \mathbf{x}^\intercal}{\partial \boldsymbol{\epsilon}} = \left(\sum_{k=1}^{N_\mathrm{a}} \frac{\partial \boldsymbol{\epsilon}_k^\intercal}{\partial \bar{\boldsymbol{\zeta}}_a}\frac{\partial \mathbf{x}_j^\intercal}{\partial \boldsymbol{\epsilon}_k}\right) = \left(\begin{cases} \frac{\partial \boldsymbol{\epsilon}_{j-1}^\intercal}{\partial \bar{\boldsymbol{\zeta}}_a}\frac{\partial \mathbf{x}_j^\intercal}{\partial \boldsymbol{\epsilon}_{j-1}} & \text{for } j>1 \\ \mathbf{0} & \text{for } j=1 \end{cases}\right), \tag{Eq. S5.6.6}$$

for $\bar{\boldsymbol{\zeta}} \in \{\bar{\mathbf{x}}, \bar{\mathbf{y}}\}$, which equals the rightmost term in Eq. S5.6.4. Thus, from Eq. S5.6.4, Eq. S5.6.5, and Eq. S5.6.6, it follows that the block matrix of total immediate effects of social partners' phenotype or genotype on a mutant's phenotype satisfies Layer 3, Eq. S3.

Noting that $\delta\mathbf{x}_a/\delta\bar{\mathbf{z}}^\intercal = (\delta\mathbf{x}_a/\delta\bar{\mathbf{x}}^\intercal, \delta\mathbf{x}_a/\delta\bar{\mathbf{y}}^\intercal)$ and that evaluation of $\mathrm{d}\mathbf{z}_a/\mathrm{d}\tau$ and $\partial\boldsymbol{\epsilon}_a/\partial\tau$ at $\mathbf{y}=\bar{\mathbf{y}}$ is $\mathrm{d}\bar{\mathbf{z}}_a/\mathrm{d}\tau$ and $\partial\bar{\boldsymbol{\epsilon}}_a/\partial\tau$ respectively, Eq. S5.6.3 can be written as

$$\frac{\mathrm{d}\bar{\mathbf{x}}_a}{\mathrm{d}\tau} = \left(\frac{\delta\mathbf{x}_a}{\delta\mathbf{x}_{a-1}^\intercal}\frac{\mathrm{d}\bar{\mathbf{x}}_{a-1}}{\mathrm{d}\tau} + \frac{\delta\mathbf{x}_a}{\delta\mathbf{y}_{a-1}^\intercal}\frac{\mathrm{d}\bar{\mathbf{y}}_{a-1}}{\mathrm{d}\tau} + \frac{\delta\mathbf{x}_a}{\delta\bar{\mathbf{z}}^\intercal}\frac{\mathrm{d}\bar{\mathbf{z}}}{\mathrm{d}\tau} + \frac{\delta\mathbf{x}_a}{\delta\boldsymbol{\epsilon}_{a-1}^\intercal}\frac{\partial\bar{\boldsymbol{\epsilon}}_{a-1}}{\partial\tau}\right)\Bigg|_{\mathbf{y}=\bar{\mathbf{y}}},$$

which is a recursion for $\mathrm{d}\bar{\mathbf{x}}_a/\mathrm{d}\tau$ over $a$. Expanding this recursion two steps yields

$$\begin{aligned}\frac{\mathrm{d}\bar{\mathbf{x}}_a}{\mathrm{d}\tau} = \Bigg\{\frac{\delta\mathbf{x}_a}{\delta\mathbf{x}_{a-1}^\intercal}\Bigg[&\frac{\delta\mathbf{x}_{a-1}}{\delta\mathbf{x}_{a-2}^\intercal}\left(\frac{\delta\mathbf{x}_{a-2}}{\delta\mathbf{x}_{a-3}^\intercal}\frac{\mathrm{d}\bar{\mathbf{x}}_{a-3}}{\mathrm{d}\tau} + \frac{\delta\mathbf{x}_{a-2}}{\delta\mathbf{y}_{a-3}^\intercal}\frac{\mathrm{d}\bar{\mathbf{y}}_{a-3}}{\mathrm{d}\tau} + \frac{\delta\mathbf{x}_{a-2}}{\delta\bar{\mathbf{z}}^\intercal}\frac{\mathrm{d}\bar{\mathbf{z}}}{\mathrm{d}\tau} + \frac{\delta\mathbf{x}_{a-2}}{\delta\boldsymbol{\epsilon}_{a-3}^\intercal}\frac{\partial\bar{\boldsymbol{\epsilon}}_{a-3}}{\partial\tau}\right)\\ &+ \frac{\delta\mathbf{x}_{a-1}}{\delta\mathbf{y}_{a-1}^\intercal}\frac{\mathrm{d}\bar{\mathbf{y}}_{a-2}}{\mathrm{d}\tau} + \frac{\delta\mathbf{x}_{a-1}}{\delta\bar{\mathbf{z}}^\intercal}\frac{\mathrm{d}\bar{\mathbf{z}}}{\mathrm{d}\tau} + \frac{\delta\mathbf{x}_{a-1}}{\delta\boldsymbol{\epsilon}_{a-2}^\intercal}\frac{\partial\bar{\boldsymbol{\epsilon}}_{a-2}}{\partial\tau}\Bigg]\\ &+ \frac{\delta\mathbf{x}_a}{\delta\mathbf{y}_{a-1}^\intercal}\frac{\mathrm{d}\bar{\mathbf{y}}_{a-1}}{\mathrm{d}\tau} + \frac{\delta\mathbf{x}_a}{\delta\bar{\mathbf{z}}^\intercal}\frac{\mathrm{d}\bar{\mathbf{z}}}{\mathrm{d}\tau} + \frac{\delta\mathbf{x}_a}{\delta\boldsymbol{\epsilon}_{a-1}^\intercal}\frac{\partial\bar{\boldsymbol{\epsilon}}_{a-1}}{\partial\tau}\Bigg\}\Bigg|_{\mathbf{y}=\bar{\mathbf{y}}}.\end{aligned}$$

Collecting the derivatives with respect to $\tau$ yields

$$\begin{aligned}\frac{\mathrm{d}\bar{\mathbf{x}}_a}{\mathrm{d}\tau} = \Bigg[\Bigg(&\frac{\delta\mathbf{x}_a}{\delta\mathbf{x}_{a-1}^\intercal}\frac{\delta\mathbf{x}_{a-1}}{\delta\mathbf{x}_{a-2}^\intercal}\frac{\delta\mathbf{x}_{a-2}}{\delta\mathbf{x}_{a-3}^\intercal}\Bigg)\frac{\mathrm{d}\bar{\mathbf{x}}_{a-3}}{\mathrm{d}\tau}\\ &+ \left(\frac{\delta\mathbf{x}_a}{\delta\mathbf{x}_{a-1}^\intercal}\frac{\delta\mathbf{x}_{a-1}}{\delta\mathbf{x}_{a-2}^\intercal}\frac{\delta\mathbf{x}_{a-2}}{\delta\mathbf{y}_{a-3}^\intercal}\right)\frac{\mathrm{d}\bar{\mathbf{y}}_{a-3}}{\mathrm{d}\tau}\\ &+ \left(\frac{\delta\mathbf{x}_a}{\delta\mathbf{x}_{a-1}^\intercal}\frac{\delta\mathbf{x}_{a-1}}{\delta\mathbf{y}_{a-1}^\intercal}\right)\frac{\mathrm{d}\bar{\mathbf{y}}_{a-2}}{\mathrm{d}\tau} + \frac{\delta\mathbf{x}_a}{\delta\mathbf{y}_{a-1}^\intercal}\frac{\mathrm{d}\bar{\mathbf{y}}_{a-1}}{\mathrm{d}\tau}\\ &+ \left(\frac{\delta\mathbf{x}_a}{\delta\mathbf{x}_{a-1}^\intercal}\frac{\delta\mathbf{x}_{a-1}}{\delta\mathbf{x}_{a-2}^\intercal}\frac{\delta\mathbf{x}_{a-2}}{\delta\boldsymbol{\epsilon}_{a-3}^\intercal}\right)\frac{\partial\bar{\boldsymbol{\epsilon}}_{a-3}}{\partial\tau}\\ &+ \left(\frac{\delta\mathbf{x}_a}{\delta\mathbf{x}_{a-1}^\intercal}\frac{\delta\mathbf{x}_{a-1}}{\delta\boldsymbol{\epsilon}_{a-2}^\intercal}\right)\frac{\partial\bar{\boldsymbol{\epsilon}}_{a-2}}{\partial\tau} + \frac{\delta\mathbf{x}_a}{\delta\boldsymbol{\epsilon}_{a-1}^\intercal}\frac{\partial\bar{\boldsymbol{\epsilon}}_{a-1}}{\partial\tau}\\ &+ \left(\frac{\delta\mathbf{x}_a}{\delta\mathbf{x}_{a-1}^\intercal}\frac{\delta\mathbf{x}_{a-1}}{\delta\mathbf{x}_{a-2}^\intercal}\frac{\delta\mathbf{x}_{a-2}}{\delta\bar{\mathbf{z}}^\intercal} + \frac{\delta\mathbf{x}_a}{\delta\mathbf{x}_{a-1}^\intercal}\frac{\delta\mathbf{x}_{a-1}}{\delta\bar{\mathbf{z}}^\intercal} + \frac{\delta\mathbf{x}_a}{\delta\bar{\mathbf{z}}^\intercal}\right)\frac{\mathrm{d}\bar{\mathbf{z}}}{\mathrm{d}\tau}\Bigg]\Bigg|_{\mathbf{y}=\bar{\mathbf{y}}}.\end{aligned}$$

Inspection shows that by expanding the recursion completely and since we assume that initial phenotype does not evolve (i.e., $\mathrm{d}\bar{\mathbf{x}}_1/\mathrm{d}\tau = \mathbf{0}$), the resulting expression can be succinctly written as

$$\begin{aligned}\frac{\mathrm{d}\bar{\mathbf{x}}_a}{\mathrm{d}\tau} = \Bigg(&\sum_{j=1}^{a-1}\overset{\curvearrowleft}{\prod_{k=j+1}^{a-1}}\frac{\delta\mathbf{x}_{k+1}}{\delta\mathbf{x}_k^\intercal}\frac{\delta\mathbf{x}_{j+1}}{\delta\mathbf{y}_j^\intercal}\frac{\mathrm{d}\bar{\mathbf{y}}_j}{\mathrm{d}\tau}\\ &+\sum_{j=1}^{a-1}\overset{\curvearrowleft}{\prod_{k=j+1}^{a-1}}\frac{\delta\mathbf{x}_{k+1}}{\delta\mathbf{x}_k^\intercal}\frac{\delta\mathbf{x}_{j+1}}{\delta\boldsymbol{\epsilon}_j^\intercal}\frac{\partial\bar{\boldsymbol{\epsilon}}_j}{\partial\tau}\\ &+\sum_{j=1}^{a-1}\overset{\curvearrowleft}{\prod_{k=j+1}^{a-1}}\frac{\delta\mathbf{x}_{k+1}}{\delta\mathbf{x}_k^\intercal}\frac{\delta\mathbf{x}_{j+1}}{\delta\bar{\mathbf{z}}^\intercal}\frac{\mathrm{d}\bar{\mathbf{z}}}{\mathrm{d}\tau}\Bigg)\Bigg|_{\mathbf{y}=\bar{\mathbf{y}}},\end{aligned}$$

where the $\curvearrowleft$ denotes left multiplication. Note that the products over $k$ are the transpose of the total effects of a mutant's phenotype at age $j+1$ on her phenotype at age $a$ (Eq. 12). Hence,

$$\begin{aligned}\frac{\mathrm{d}\bar{\mathbf{x}}_a}{\mathrm{d}\tau} = \Bigg(&\sum_{j=1}^{a-1}\frac{\mathrm{d}\mathbf{x}_a}{\mathrm{d}\mathbf{x}_{j+1}^\intercal}\frac{\delta\mathbf{x}_{j+1}}{\delta\mathbf{y}_j^\intercal}\frac{\mathrm{d}\bar{\mathbf{y}}_j}{\mathrm{d}\tau} + \sum_{j=1}^{a-1}\frac{\mathrm{d}\mathbf{x}_a}{\mathrm{d}\mathbf{x}_{j+1}^\intercal}\frac{\delta\mathbf{x}_{j+1}}{\delta\boldsymbol{\epsilon}_j^\intercal}\frac{\partial\bar{\boldsymbol{\epsilon}}_j}{\partial\tau}\\ &+\sum_{j=1}^{a-1}\frac{\mathrm{d}\mathbf{x}_a}{\mathrm{d}\mathbf{x}_{j+1}^\intercal}\frac{\delta\mathbf{x}_{j+1}}{\delta\bar{\mathbf{z}}^\intercal}\frac{\mathrm{d}\bar{\mathbf{z}}}{\mathrm{d}\tau}\Bigg)\Bigg|_{\mathbf{y}=\bar{\mathbf{y}}}. \qquad \text{(Eq. S5.6.7)}\end{aligned}$$

Before simplifying Eq. S5.6.7, we introduce a couple of matrices that are analogous to those already provided, based on Eq. S5.2.17. The matrix of *total effects of social partners' phenotype or genotypic traits at age a on a mutant's phenotype at age j* is

$$\frac{\mathrm{d}\mathbf{x}_j^\intercal}{\mathrm{d}\bar{\boldsymbol{\zeta}}_a}\Bigg|_{\mathbf{y}=\bar{\mathbf{y}}} = \begin{cases}\displaystyle\sum_{l=1}^{N_\mathrm{a}}\left(\frac{\delta\mathbf{x}_l^\intercal}{\delta\bar{\boldsymbol{\zeta}}_a}\frac{\mathrm{d}\mathbf{x}_j^\intercal}{\mathrm{d}\mathbf{x}_l}\right)\Bigg|_{\mathbf{y}=\bar{\mathbf{y}}} & \text{for } j>1\\ \mathbf{0} & \text{for } j=1,\end{cases} \qquad \text{(Eq. S5.6.8)}$$

for $\bar{\boldsymbol{\zeta}} \in \{\bar{\mathbf{x}}, \bar{\mathbf{y}}\}$. The block matrix of *total effects of social partners' phenotype or genotype on a mutant's phenotype* is thus

$$\begin{aligned}\frac{\mathrm{d}\mathbf{x}^\intercal}{\mathrm{d}\bar{\boldsymbol{\zeta}}}\Bigg|_{\mathbf{y}=\bar{\mathbf{y}}} &\equiv \begin{pmatrix}\frac{\mathrm{d}\mathbf{x}_1^\intercal}{\mathrm{d}\bar{\boldsymbol{\zeta}}_1} & \cdots & \frac{\mathrm{d}\mathbf{x}_{N_\mathrm{a}}^\intercal}{\mathrm{d}\bar{\boldsymbol{\zeta}}_1}\\ \vdots & \ddots & \vdots\\ \frac{\mathrm{d}\mathbf{x}_1^\intercal}{\mathrm{d}\bar{\boldsymbol{\zeta}}_{N_\mathrm{a}}} & \cdots & \frac{\mathrm{d}\mathbf{x}_{N_\mathrm{a}}^\intercal}{\mathrm{d}\bar{\boldsymbol{\zeta}}_{N_\mathrm{a}}}\end{pmatrix}\Bigg|_{\mathbf{y}=\bar{\mathbf{y}}}\\ &= \begin{pmatrix}\mathbf{0} & \frac{\mathrm{d}\mathbf{x}_2^\intercal}{\mathrm{d}\bar{\boldsymbol{\zeta}}_1} & \cdots & \frac{\mathrm{d}\mathbf{x}_{N_\mathrm{a}}^\intercal}{\mathrm{d}\bar{\boldsymbol{\zeta}}_1}\\ \mathbf{0} & \frac{\mathrm{d}\mathbf{x}_2^\intercal}{\mathrm{d}\bar{\boldsymbol{\zeta}}_2} & \cdots & \frac{\mathrm{d}\mathbf{x}_{N_\mathrm{a}}^\intercal}{\mathrm{d}\bar{\boldsymbol{\zeta}}_2}\\ \vdots & \vdots & \ddots & \vdots\\ \mathbf{0} & \frac{\mathrm{d}\mathbf{x}_2^\intercal}{\mathrm{d}\bar{\boldsymbol{\zeta}}_{N_\mathrm{a}}} & \cdots & \frac{\mathrm{d}\mathbf{x}_{N_\mathrm{a}}^\intercal}{\mathrm{d}\bar{\boldsymbol{\zeta}}_{N_\mathrm{a}}}\end{pmatrix}\Bigg|_{\mathbf{y}=\bar{\mathbf{y}}}, \qquad \text{(Eq. S5.6.9)}\end{aligned}$$

for $\bar{\boldsymbol{\zeta}} \in \{\bar{\mathbf{x}}, \bar{\mathbf{y}}\}$. Then, from Eq. S5.6.8, the block matrix in Eq. S5.6.9 satisfies Layer 4, Eq. S4.

Using Eq. S5.2.17 and Eq. S5.3.9 and given the property of transpose of a product (i.e., $(\mathbf{AB})^\intercal = \mathbf{B}^\intercal\mathbf{A}^\intercal$), Eq. S5.6.7 can be written more succinctly as

$$\frac{\mathrm{d}\bar{\mathbf{x}}_a}{\mathrm{d}\tau} = \left(\sum_{j=1}^{a-1}\frac{\mathrm{d}\mathbf{x}_a}{\mathrm{d}\mathbf{y}_j^\intercal}\frac{\mathrm{d}\bar{\mathbf{y}}_j}{\mathrm{d}\tau} + \sum_{j=1}^{a-1}\frac{\mathrm{d}\mathbf{x}_a}{\mathrm{d}\boldsymbol{\epsilon}_j^\intercal}\frac{\partial\bar{\boldsymbol{\epsilon}}_j}{\partial\tau} + \sum_{j=1}^{a-1}\frac{\mathrm{d}\mathbf{x}_a}{\mathrm{d}\mathbf{x}_{j+1}^\intercal}\frac{\delta\mathbf{x}_{j+1}}{\delta\bar{\mathbf{z}}^\intercal}\frac{\mathrm{d}\bar{\mathbf{z}}}{\mathrm{d}\tau}\right)\Bigg|_{\mathbf{y}=\bar{\mathbf{y}}}.$$

Note that from Eq. S5.2.16, we have that $\mathrm{d}\mathbf{x}_a/\mathrm{d}\mathbf{y}_j^\intercal = \mathbf{0}$ for $j \geq a$, from Eq. S5.3.10, we have that $\mathrm{d}\mathbf{x}_a/\mathrm{d}\boldsymbol{\epsilon}_j^\intercal = \mathbf{0}$ for $j \geq a$, and from Eq. S5.1.15, we have that $\mathrm{d}\mathbf{x}_a/\mathrm{d}\mathbf{x}_{j+1}^\intercal = \mathbf{0}$ for $j+1 \geq a$. Hence, the same expression holds extending the upper bounds of the sums to the last possible age:

$$\frac{\mathrm{d}\bar{\mathbf{x}}_a}{\mathrm{d}\tau} = \left(\sum_{j=1}^{N_\mathrm{a}}\frac{\mathrm{d}\mathbf{x}_a}{\mathrm{d}\mathbf{y}_j^\intercal}\frac{\mathrm{d}\bar{\mathbf{y}}_j}{\mathrm{d}\tau} + \sum_{j=1}^{N_\mathrm{a}}\frac{\mathrm{d}\mathbf{x}_a}{\mathrm{d}\boldsymbol{\epsilon}_j^\intercal}\frac{\partial\bar{\boldsymbol{\epsilon}}_j}{\partial\tau}\right.$$

$$+\sum_{j=1}^{N_a-1}\frac{\mathrm{d}\mathbf{x}_a}{\mathrm{d}\mathbf{x}_{j+1}^{\intercal}}\frac{\delta\mathbf{x}_{j+1}}{\delta\bar{\mathbf{z}}^{\intercal}}\frac{\mathrm{d}\bar{\mathbf{z}}}{\mathrm{d}\tau}\Bigg)\Bigg|_{\mathbf{y}=\bar{\mathbf{y}}}.$$

Changing the sum index for the rightmost sum yields

$$\frac{\mathrm{d}\bar{\mathbf{x}}_a}{\mathrm{d}\tau}=\left(\sum_{j=1}^{N_a}\frac{\mathrm{d}\mathbf{x}_a}{\mathrm{d}\mathbf{y}_j^{\intercal}}\frac{\mathrm{d}\bar{\mathbf{y}}_j}{\mathrm{d}\tau}+\sum_{j=1}^{N_a}\frac{\mathrm{d}\mathbf{x}_a}{\mathrm{d}\boldsymbol{\epsilon}_j^{\intercal}}\frac{\partial\bar{\boldsymbol{\epsilon}}_j}{\partial\tau}+\sum_{j=2}^{N_a}\frac{\mathrm{d}\mathbf{x}_a}{\mathrm{d}\mathbf{x}_j^{\intercal}}\frac{\delta\mathbf{x}_j}{\delta\bar{\mathbf{z}}^{\intercal}}\frac{\mathrm{d}\bar{\mathbf{z}}}{\mathrm{d}\tau}\right)\Bigg|_{\mathbf{y}=\bar{\mathbf{y}}}.$$

Expanding the matrix calculus notation for the entries of $\bar{\mathbf{z}}$ in the rightmost sum yields

$$\frac{\mathrm{d}\bar{\mathbf{x}}_a}{\mathrm{d}\tau}=\Bigg(\sum_{j=1}^{N_a}\frac{\mathrm{d}\mathbf{x}_a}{\mathrm{d}\mathbf{y}_j^{\intercal}}\frac{\mathrm{d}\bar{\mathbf{y}}_j}{\mathrm{d}\tau}+\sum_{j=1}^{N_a}\frac{\mathrm{d}\mathbf{x}_a}{\mathrm{d}\boldsymbol{\epsilon}_j^{\intercal}}\frac{\partial\bar{\boldsymbol{\epsilon}}_j}{\partial\tau}$$
$$+\sum_{j=2}^{N_a}\frac{\mathrm{d}\mathbf{x}_a}{\mathrm{d}\mathbf{x}_j^{\intercal}}\frac{\delta\mathbf{x}_j}{\delta\bar{\mathbf{x}}^{\intercal}}\frac{\mathrm{d}\bar{\mathbf{x}}}{\mathrm{d}\tau}+\sum_{j=2}^{N_a}\frac{\mathrm{d}\mathbf{x}_a}{\mathrm{d}\mathbf{x}_j^{\intercal}}\frac{\delta\mathbf{x}_j}{\delta\bar{\mathbf{y}}^{\intercal}}\frac{\mathrm{d}\bar{\mathbf{y}}}{\mathrm{d}\tau}\Bigg)\Bigg|_{\mathbf{y}=\bar{\mathbf{y}}}.$$

Expanding again the matrix calculus notation for the entries of $\bar{\mathbf{x}}$ and $\bar{\mathbf{y}}$ in the two rightmost sums yields

$$\frac{\mathrm{d}\bar{\mathbf{x}}_a}{\mathrm{d}\tau}=\Bigg(\sum_{j=1}^{N_a}\frac{\mathrm{d}\mathbf{x}_a}{\mathrm{d}\mathbf{y}_j^{\intercal}}\frac{\mathrm{d}\bar{\mathbf{y}}_j}{\mathrm{d}\tau}+\sum_{j=1}^{N_a}\frac{\mathrm{d}\mathbf{x}_a}{\mathrm{d}\boldsymbol{\epsilon}_j^{\intercal}}\frac{\partial\bar{\boldsymbol{\epsilon}}_j}{\partial\tau}$$
$$+\sum_{l=1}^{N_a}\sum_{j=2}^{N_a}\frac{\mathrm{d}\mathbf{x}_a}{\mathrm{d}\mathbf{x}_j^{\intercal}}\frac{\delta\mathbf{x}_j}{\delta\bar{\mathbf{x}}_l^{\intercal}}\frac{\mathrm{d}\bar{\mathbf{x}}_l}{\mathrm{d}\tau}+\sum_{l=1}^{N_a}\sum_{j=2}^{N_a}\frac{\mathrm{d}\mathbf{x}_a}{\mathrm{d}\mathbf{x}_j^{\intercal}}\frac{\delta\mathbf{x}_j}{\delta\bar{\mathbf{y}}_l^{\intercal}}\frac{\mathrm{d}\bar{\mathbf{y}}_l}{\mathrm{d}\tau}\Bigg)\Bigg|_{\mathbf{y}=\bar{\mathbf{y}}}.$$

Using the transpose of the matrix in Eq. S5.6.8 in the two rightmost terms, noting that $\delta\mathbf{x}_j/\delta\bar{\mathbf{x}}_l^{\intercal}=\mathbf{0}$ and $\delta\mathbf{x}_j/\delta\bar{\mathbf{y}}_l^{\intercal}=\mathbf{0}$ for $j=1$ (from Eq. S5.6.5), yields

$$\frac{\mathrm{d}\bar{\mathbf{x}}_a}{\mathrm{d}\tau}=\Bigg(\sum_{j=1}^{N_a}\frac{\mathrm{d}\mathbf{x}_a}{\mathrm{d}\mathbf{y}_j^{\intercal}}\frac{\mathrm{d}\bar{\mathbf{y}}_j}{\mathrm{d}\tau}+\sum_{j=1}^{N_a}\frac{\mathrm{d}\mathbf{x}_a}{\mathrm{d}\boldsymbol{\epsilon}_j^{\intercal}}\frac{\partial\bar{\boldsymbol{\epsilon}}_j}{\partial\tau}$$
$$+\sum_{l=1}^{N_a}\frac{\mathrm{d}\mathbf{x}_a}{\mathrm{d}\bar{\mathbf{x}}_l^{\intercal}}\frac{\mathrm{d}\bar{\mathbf{x}}_l}{\mathrm{d}\tau}+\sum_{l=1}^{N_a}\frac{\mathrm{d}\mathbf{x}_a}{\mathrm{d}\bar{\mathbf{y}}_l^{\intercal}}\frac{\mathrm{d}\bar{\mathbf{y}}_l}{\mathrm{d}\tau}\Bigg)\Bigg|_{\mathbf{y}=\bar{\mathbf{y}}}.$$

Applying matrix calculus notation to each term yields

$$\frac{\mathrm{d}\bar{\mathbf{x}}_a}{\mathrm{d}\tau}=\left(\frac{\mathrm{d}\mathbf{x}_a}{\mathrm{d}\mathbf{y}^{\intercal}}\frac{\mathrm{d}\bar{\mathbf{y}}}{\mathrm{d}\tau}+\frac{\mathrm{d}\mathbf{x}_a}{\mathrm{d}\boldsymbol{\epsilon}^{\intercal}}\frac{\partial\bar{\boldsymbol{\epsilon}}}{\partial\tau}+\frac{\mathrm{d}\mathbf{x}_a}{\mathrm{d}\bar{\mathbf{x}}^{\intercal}}\frac{\mathrm{d}\bar{\mathbf{x}}}{\mathrm{d}\tau}+\frac{\mathrm{d}\mathbf{x}_a}{\mathrm{d}\bar{\mathbf{y}}^{\intercal}}\frac{\mathrm{d}\bar{\mathbf{y}}}{\mathrm{d}\tau}\right)\Bigg|_{\mathbf{y}=\bar{\mathbf{y}}},$$

for $a\in\{2,\ldots,N_a\}$. Since $\mathrm{d}\bar{\mathbf{x}}_1/\mathrm{d}\tau=\mathbf{0}$, it follows that

$$\frac{\mathrm{d}\bar{\mathbf{x}}}{\mathrm{d}\tau}=\left(\frac{\mathrm{d}\mathbf{x}}{\mathrm{d}\mathbf{y}^{\intercal}}\frac{\mathrm{d}\bar{\mathbf{y}}}{\mathrm{d}\tau}+\frac{\mathrm{d}\mathbf{x}}{\mathrm{d}\boldsymbol{\epsilon}^{\intercal}}\frac{\partial\bar{\boldsymbol{\epsilon}}}{\partial\tau}+\frac{\mathrm{d}\mathbf{x}}{\mathrm{d}\bar{\mathbf{x}}^{\intercal}}\frac{\mathrm{d}\bar{\mathbf{x}}}{\mathrm{d}\tau}+\frac{\mathrm{d}\mathbf{x}}{\mathrm{d}\bar{\mathbf{y}}^{\intercal}}\frac{\mathrm{d}\bar{\mathbf{y}}}{\mathrm{d}\tau}\right)\Bigg|_{\mathbf{y}=\bar{\mathbf{y}}},$$

(Eq. S5.6.10)

which contains our desired $\mathrm{d}\bar{\mathbf{x}}/\mathrm{d}\tau$ on both sides of the equation.

The matrix premultiplying $\mathrm{d}\bar{\mathbf{x}}/\mathrm{d}\tau$ on the right-hand side of Eq. S5.6.10 is $\mathrm{d}\mathbf{x}/\mathrm{d}\bar{\mathbf{x}}^{\intercal}|_{\mathbf{y}=\bar{\mathbf{y}}}$, which is square. We now make use of our assumption that the absolute value of all the eigenvalues of $\mathrm{d}\mathbf{x}/\mathrm{d}\bar{\mathbf{x}}^{\intercal}|_{\mathbf{y}=\bar{\mathbf{y}}}$ is strictly less than one, which guarantees that the resident geno-phenotype is socio-devo stable (Eq. S2.1.3 and following text). Given this property of $\mathrm{d}\mathbf{x}/\mathrm{d}\bar{\mathbf{x}}^{\intercal}|_{\mathbf{y}=\bar{\mathbf{y}}}$, then $\mathbf{I}-\mathrm{d}\mathbf{x}/\mathrm{d}\bar{\mathbf{x}}^{\intercal}|_{\mathbf{y}=\bar{\mathbf{y}}}$ is invertible. Hence, we can define the transpose of the matrix of *stabilized effects of a focal individual's phenotype on a social partners' phenotype* (second equality of Layer 5, Eq. S1). Thus, solving for $\mathrm{d}\bar{\mathbf{x}}/\mathrm{d}\tau$ in Eq. S5.6.10, we finally obtain an equation describing the evolutionary dynamics of the phenotype

$$\frac{\mathrm{d}\bar{\mathbf{x}}}{\mathrm{d}\tau}=\left[\frac{\mathrm{s}\mathbf{x}}{\mathrm{s}\bar{\mathbf{x}}^{\intercal}}\left(\frac{\mathrm{d}\mathbf{x}}{\mathrm{d}\mathbf{y}^{\intercal}}+\frac{\mathrm{d}\mathbf{x}}{\mathrm{d}\bar{\mathbf{y}}^{\intercal}}\right)\frac{\mathrm{d}\bar{\mathbf{y}}}{\mathrm{d}\tau}+\frac{\mathrm{s}\mathbf{x}}{\mathrm{s}\bar{\mathbf{x}}^{\intercal}}\frac{\mathrm{d}\mathbf{x}}{\mathrm{d}\boldsymbol{\epsilon}^{\intercal}}\frac{\partial\bar{\boldsymbol{\epsilon}}}{\partial\tau}\right]\Bigg|_{\mathbf{y}=\bar{\mathbf{y}}}.$$

Let us momentarily write $\mathbf{x}=\tilde{\mathbf{g}}(\mathbf{y},\bar{\mathbf{y}})$ for some continuously differentiable function $\tilde{\mathbf{g}}$ to highlight the dependence of a mutant's phenotype $\mathbf{x}$ on her genotype $\mathbf{y}$ and on the genotype $\bar{\mathbf{y}}$ of resident social partners. Consider the resident phenotype that develops in the context of the mutant genotype, denoted by $\check{\mathbf{x}}=\tilde{\mathbf{g}}(\bar{\mathbf{y}},\mathbf{y})$. Hence,

$$\frac{\mathrm{d}\check{\mathbf{x}}}{\mathrm{d}\mathbf{y}^{\intercal}}\Bigg|_{\mathbf{y}=\bar{\mathbf{y}}}=\frac{\mathrm{d}\tilde{\mathbf{g}}(\bar{\mathbf{y}},\mathbf{y})}{\mathrm{d}\mathbf{y}^{\intercal}}\Bigg|_{\mathbf{y}=\bar{\mathbf{y}}}=\frac{\mathrm{d}\tilde{\mathbf{g}}(\mathbf{y},\bar{\mathbf{y}})}{\mathrm{d}\bar{\mathbf{y}}^{\intercal}}\Bigg|_{\mathbf{y}=\bar{\mathbf{y}}}=\frac{\mathrm{d}\mathbf{x}}{\mathrm{d}\bar{\mathbf{y}}^{\intercal}}\Bigg|_{\mathbf{y}=\bar{\mathbf{y}}},$$

(Eq. S5.6.11)

where the second equality follows by exchanging dummy variables. Then, the transpose of the matrix of *total social effects of a mutant's genotype on her and a partner's phenotypes* is

$$\frac{\mathrm{d}(\mathbf{x}+\check{\mathbf{x}})}{\mathrm{d}\mathbf{y}^{\intercal}}\Bigg|_{\mathbf{y}=\bar{\mathbf{y}}}=\left(\frac{\mathrm{d}\mathbf{x}}{\mathrm{d}\mathbf{y}^{\intercal}}+\frac{\mathrm{d}\check{\mathbf{x}}}{\mathrm{d}\mathbf{y}^{\intercal}}\right)\Bigg|_{\mathbf{y}=\bar{\mathbf{y}}}$$
$$=\left(\frac{\mathrm{d}\mathbf{x}}{\mathrm{d}\mathbf{y}^{\intercal}}+\frac{\mathrm{d}\mathbf{x}}{\mathrm{d}\bar{\mathbf{y}}^{\intercal}}\right)\Bigg|_{\mathbf{y}=\bar{\mathbf{y}}}\in\mathbb{R}^{N_aN_p\times N_aN_g}.$$

(Eq. S5.6.12)

Similarly, let us momentarily write $\mathbf{x}=\tilde{\tilde{\mathbf{g}}}(\mathbf{x},\bar{\mathbf{x}})$ for some continuously differentiable function $\tilde{\tilde{\mathbf{g}}}$ to highlight the dependence of a mutant's phenotype $\mathbf{x}$ on her (developmentally earlier) phenotype $\mathbf{x}$ and on the phenotype $\bar{\mathbf{x}}$ of resident social partners. Consider the resident phenotype that develops in the context of the mutant phenotype, denoted by $\check{\mathbf{x}}=\tilde{\tilde{\mathbf{g}}}(\bar{\mathbf{x}},\mathbf{x})$. Hence,

$$\frac{\mathrm{d}\check{\mathbf{x}}}{\mathrm{d}\mathbf{x}^{\intercal}}\Bigg|_{\mathbf{y}=\bar{\mathbf{y}}}=\frac{\mathrm{d}\tilde{\tilde{\mathbf{g}}}(\bar{\mathbf{x}},\mathbf{x})}{\mathrm{d}\mathbf{x}^{\intercal}}\Bigg|_{\mathbf{y}=\bar{\mathbf{y}}}=\frac{\mathrm{d}\tilde{\tilde{\mathbf{g}}}(\mathbf{x},\bar{\mathbf{x}})}{\mathrm{d}\bar{\mathbf{x}}^{\intercal}}\Bigg|_{\mathbf{y}=\bar{\mathbf{y}}}=\frac{\mathrm{d}\mathbf{x}}{\mathrm{d}\bar{\mathbf{x}}^{\intercal}}\Bigg|_{\mathbf{y}=\bar{\mathbf{y}}},$$

(Eq. S5.6.13)

where the second equality follows by exchanging dummy variables. Then, the transpose of the matrix of *total social effects of a mutant's phenotype on her and a partner's phenotypes* is

$$\frac{\mathrm{d}(\mathbf{x}+\check{\mathbf{x}})}{\mathrm{d}\mathbf{x}^{\intercal}}\Bigg|_{\mathbf{y}=\bar{\mathbf{y}}}=\left(\frac{\mathrm{d}\mathbf{x}}{\mathrm{d}\mathbf{x}^{\intercal}}+\frac{\mathrm{d}\check{\mathbf{x}}}{\mathrm{d}\mathbf{x}^{\intercal}}\right)\Bigg|_{\mathbf{y}=\bar{\mathbf{y}}}$$
$$=\left(\frac{\mathrm{d}\mathbf{x}}{\mathrm{d}\mathbf{x}^{\intercal}}+\frac{\mathrm{d}\mathbf{x}}{\mathrm{d}\bar{\mathbf{x}}^{\intercal}}\right)\Bigg|_{\mathbf{y}=\bar{\mathbf{y}}}\in\mathbb{R}^{N_aN_p\times N_aN_p}.$$

(Eq. S5.6.14)

Thus, from Eq. S5.6.13 and the second equality of Layer 5, Eq. S1, the transpose of the matrix of stabilized effects of a focal individual's phenotype on social partners' phenotype may also be written as

$$\frac{\mathrm{s}\mathbf{x}}{\mathrm{s}\bar{\mathbf{x}}^{\intercal}}\Bigg|_{\mathbf{y}=\bar{\mathbf{y}}}=\left(\mathbf{I}-\frac{\mathrm{d}\check{\mathbf{x}}}{\mathrm{d}\mathbf{x}^{\intercal}}\Bigg|_{\mathbf{y}=\bar{\mathbf{y}}}\right)^{-1}$$
$$=\sum_{\theta=1}^{\infty}\left(\frac{\mathrm{d}\check{\mathbf{x}}}{\mathrm{d}\mathbf{x}^{\intercal}}\right)^{\theta-1}\Bigg|_{\mathbf{y}=\bar{\mathbf{y}}}\in\mathbb{R}^{N_aN_p\times N_aN_p},$$

where the last equality follows from the geometric series of matrices. This equation is the first and third equalities of Layer 5, Eq. S1.

Therefore, using Layer 5, Eq. S2 and Layer 5, Eq. S2b, the evolutionary dynamics of the phenotype are given by

$$\frac{\mathrm{d}\bar{\mathbf{x}}}{\mathrm{d}\tau}=\left(\frac{\mathrm{s}\mathbf{x}}{\mathrm{s}\bar{\mathbf{x}}^{\intercal}}\frac{\mathrm{d}(\mathbf{x}+\check{\mathbf{x}})}{\mathrm{d}\mathbf{y}^{\intercal}}\frac{\mathrm{d}\bar{\mathbf{y}}}{\mathrm{d}\tau}+\frac{\mathrm{s}\mathbf{x}}{\mathrm{s}\bar{\mathbf{x}}^{\intercal}}\frac{\mathrm{d}\mathbf{x}}{\mathrm{d}\boldsymbol{\epsilon}^{\intercal}}\frac{\partial\bar{\boldsymbol{\epsilon}}}{\partial\tau}\right)\Bigg|_{\mathbf{y}=\bar{\mathbf{y}}}$$

$$\approx \left( \iota \frac{\mathrm{s}\mathbf{x}}{\mathrm{s}\mathbf{y}^\intercal} \mathbf{H}_\mathbf{y} \frac{\mathrm{d}w}{\mathrm{d}\mathbf{y}} + \frac{\mathrm{s}\mathbf{x}}{\mathrm{s}\boldsymbol{\epsilon}^\intercal} \frac{\partial \bar{\boldsymbol{\epsilon}}}{\partial \tau} \right) \bigg|_{\mathbf{y}=\bar{\mathbf{y}}}$$

$$= \left( \iota \mathbf{L}_{\mathbf{x}\mathbf{y}} \frac{\mathrm{d}w}{\mathrm{d}\mathbf{y}} + \frac{\mathrm{s}\mathbf{x}}{\mathrm{s}\boldsymbol{\epsilon}^\intercal} \frac{\partial \bar{\boldsymbol{\epsilon}}}{\partial \tau} \right) \bigg|_{\mathbf{y}=\bar{\mathbf{y}}}, \qquad \text{(Eq. S5.6.15)}$$

where the second line follows by using Eq. S5.6.1 in the limit $\Delta\tau \to 0$, and the third line follows from Layer 6, Eq. 13. The first line of Eq. S5.6.15 describing evolutionary change of the phenotype in terms of evolutionary change of the genotype is a generalization of previous equations describing the evolution of a multivariate phenotype in terms of allele frequency change (e.g., the first equation on p. 49 of Engen and Sæther 2021). Eq. S5.6.15 is Layer 7, Eq. 5 for $\boldsymbol{\zeta} = \mathbf{x}$. Using the third line of Layer 4, Eq. S21 and Layer 6, Eq. 11 yields Layer 7, Eq. 4 for $\boldsymbol{\zeta} = \mathbf{x}$, whereas using the fourth line of Layer 4, Eq. S21 and Layer 6, Eq. 12 yields Layer 7, Eq. 1a for $\boldsymbol{\zeta} = \mathbf{x}$.

#### S5.7 Evolutionary dynamics of the geno-phenotype

Here we obtain equations describing the evolutionary dynamics of the resident geno-phenotype, that is, $\mathrm{d}\bar{\mathbf{z}}/\mathrm{d}\tau$.

##### S5.7.1 In terms of total genotypic selection

In this section, we obtain such an equation in terms of total genotypic selection. Since $\mathrm{d}\bar{\mathbf{z}}/\mathrm{d}\tau = (\mathrm{d}\bar{\mathbf{x}}/\mathrm{d}\tau; \mathrm{d}\bar{\mathbf{y}}/\mathrm{d}\tau)$, from Eq. S5.6.15 and Eq. S2.2.2a, we can write the evolutionary dynamics of the resident geno-phenotype $\bar{\mathbf{z}}$ as

$$\frac{\mathrm{d}\bar{\mathbf{z}}}{\mathrm{d}\tau} \approx \left[ \iota \begin{pmatrix} \mathbf{L}_{\mathbf{x}\mathbf{y}} \\ \mathbf{H}_\mathbf{y} \end{pmatrix} \frac{\mathrm{d}w}{\mathrm{d}\mathbf{y}} + \begin{pmatrix} \frac{\mathrm{s}\mathbf{x}}{\mathrm{s}\boldsymbol{\epsilon}^\intercal} \\ \mathbf{0} \end{pmatrix} \frac{\partial \bar{\boldsymbol{\epsilon}}}{\partial \tau} \right] \bigg|_{\mathbf{y}=\bar{\mathbf{y}}}. \qquad \text{(Eq. S5.7.1)}$$

Using Layer 6, Eq. 13 and Layer 5, Eq. S3, this is

$$\frac{\mathrm{d}\bar{\mathbf{z}}}{\mathrm{d}\tau} \approx \left[ \iota \begin{pmatrix} \frac{\mathrm{s}\mathbf{x}}{\mathrm{s}\mathbf{y}^\intercal} \\ \frac{\mathrm{s}\mathbf{y}}{\mathrm{s}\mathbf{y}^\intercal} \end{pmatrix} \mathbf{H}_\mathbf{y} \frac{\mathrm{d}w}{\mathrm{d}\mathbf{y}} + \begin{pmatrix} \frac{\mathrm{s}\mathbf{x}}{\mathrm{s}\boldsymbol{\epsilon}^\intercal} \\ \frac{\mathrm{s}\mathbf{y}}{\mathrm{s}\boldsymbol{\epsilon}^\intercal} \end{pmatrix} \frac{\partial \bar{\boldsymbol{\epsilon}}}{\partial \tau} \right] \bigg|_{\mathbf{y}=\bar{\mathbf{y}}}.$$

Using Layer 5, Eq. S4, this reduces to

$$\frac{\mathrm{d}\bar{\mathbf{z}}}{\mathrm{d}\tau} \approx \left( \iota \frac{\mathrm{s}\mathbf{z}}{\mathrm{s}\mathbf{y}^\intercal} \mathbf{H}_\mathbf{y} \frac{\mathrm{d}w}{\mathrm{d}\mathbf{y}} + \frac{\mathrm{s}\mathbf{z}}{\mathrm{s}\boldsymbol{\epsilon}^\intercal} \frac{\partial \bar{\boldsymbol{\epsilon}}}{\partial \tau} \right) \bigg|_{\mathbf{y}=\bar{\mathbf{y}}}.$$

Using Layer 6, Eq. 13 yields Layer 7, Eq. 5 for $\boldsymbol{\zeta} = \mathbf{z}$. Using the third line of Layer 4, Eq. S21 and Layer 6, Eq. 11 yields Layer 7, Eq. 4 for $\boldsymbol{\zeta} = \mathbf{z}$, whereas using the fourth line of Layer 4, Eq. S21 and Layer 6, Eq. 12 yields Layer 7, Eq. 1a for $\boldsymbol{\zeta} = \mathbf{z}$.

In contrast to other arrangements, the premultiplying matrix $\mathbf{L}_{\mathbf{z}\mathbf{y}}$ is non-singular if $\mathbf{H}_\mathbf{y}$ is non-singular. Indeed, if

$$\frac{\mathrm{s}\mathbf{z}}{\mathrm{s}\mathbf{y}^\intercal}\bigg|_{\mathbf{y}=\bar{\mathbf{y}}} \mathbf{r} = \mathbf{0}$$

for some vector $\mathbf{r}$, then from Layer 5, Eq. S4a and Layer 5, Eq. S3b we have

$$\begin{pmatrix} \frac{\mathrm{s}\mathbf{x}}{\mathrm{s}\mathbf{y}^\intercal} \\ \mathbf{I} \end{pmatrix} \bigg|_{\mathbf{y}=\bar{\mathbf{y}}} \mathbf{r} = \mathbf{0}.$$

Doing the multiplication yields

$$\begin{pmatrix} \frac{\mathrm{s}\mathbf{x}}{\mathrm{s}\mathbf{y}^\intercal}\Big|_{\mathbf{y}=\bar{\mathbf{y}}} \mathbf{r} \\ \mathbf{r} \end{pmatrix} = \mathbf{0},$$

which implies that $\mathbf{r} = \mathbf{0}$, so $\mathrm{s}\mathbf{z}/\mathrm{s}\mathbf{y}^\intercal|_{\mathbf{y}=\bar{\mathbf{y}}}$ is non-singular. Thus, $\mathbf{L}_{\mathbf{z}\mathbf{y}}$ is non-singular if $\mathbf{H}_\mathbf{y}$ is non-singular.

##### S5.7.2 In terms of total selection on the geno-phenotype

In this section, we obtain an equation for the evolutionary dynamics of the resident geno-phenotype in terms of the total selection gradient of the geno-phenotype.

First, using Layer 6, Eq. 2, we define the *mechanistic additive genetic covariance matrix of the unperturbed geno-phenotype* $\hat{\mathbf{z}} \equiv (\bar{\mathbf{x}}; \mathbf{y})$ as

$$\mathbf{H}_{\hat{\mathbf{z}}} \equiv \mathrm{cov}[\mathbf{b}_{\hat{\mathbf{z}}}, \mathbf{b}_{\hat{\mathbf{z}}}] = \left( \frac{\mathrm{d}\hat{\mathbf{z}}}{\mathrm{d}\mathbf{y}^\intercal} \mathbf{H}_\mathbf{y} \frac{\mathrm{d}\hat{\mathbf{z}}^\intercal}{\mathrm{d}\mathbf{y}} \right) \bigg|_{\mathbf{y}=\bar{\mathbf{y}}} \in \mathbb{R}^{N_\mathrm{a}(N_\mathrm{p}+N_\mathrm{g}) \times N_\mathrm{a}(N_\mathrm{p}+N_\mathrm{g})}.$$

By definition of $\hat{\mathbf{z}}$, we have

$$\mathbf{H}_{\hat{\mathbf{z}}} = \left[ \begin{pmatrix} \frac{\mathrm{d}\bar{\mathbf{x}}}{\mathrm{d}\mathbf{y}^\intercal} \\ \frac{\mathrm{d}\mathbf{y}}{\mathrm{d}\mathbf{y}^\intercal} \end{pmatrix} \mathbf{H}_\mathbf{y} \begin{pmatrix} \frac{\mathrm{d}\bar{\mathbf{x}}^\intercal}{\mathrm{d}\mathbf{y}} & \frac{\mathrm{d}\mathbf{y}^\intercal}{\mathrm{d}\mathbf{y}} \end{pmatrix} \right] \bigg|_{\mathbf{y}=\bar{\mathbf{y}}}.$$

From Eq. S2.2.2c, the resident phenotype is independent of mutant genotype, so

$$\mathbf{H}_{\hat{\mathbf{z}}} = \left[ \begin{pmatrix} \mathbf{0} \\ \mathbf{I} \end{pmatrix} \mathbf{H}_\mathbf{y} \begin{pmatrix} \mathbf{0} & \mathbf{I} \end{pmatrix} \right] \bigg|_{\mathbf{y}=\bar{\mathbf{y}}}.$$

Doing the matrix multiplication yields

$$\mathbf{H}_{\hat{\mathbf{z}}} = \left[ \begin{pmatrix} \mathbf{0} \\ \mathbf{I} \end{pmatrix} \begin{pmatrix} \mathbf{0} & \mathbf{H}_\mathbf{y} \end{pmatrix} \right] \bigg|_{\mathbf{y}=\bar{\mathbf{y}}} = \begin{pmatrix} \mathbf{0} & \mathbf{0} \\ \mathbf{0} & \mathbf{H}_\mathbf{y} \end{pmatrix}. \qquad \text{(Eq. S5.7.2)}$$

The matrix $\mathbf{H}_{\hat{\mathbf{z}}}$ is singular because the unperturbed geno-phenotype includes the genotype (i.e., $\mathrm{d}\hat{\mathbf{z}}^\intercal/\mathrm{d}\mathbf{y}|_{\mathbf{y}=\bar{\mathbf{y}}}$ has fewer rows than columns). For this reason, the matrix $\mathbf{H}_{\hat{\mathbf{z}}}$ would still be singular even if the zero block entries in Eq. S5.7.2 were non-zero (i.e., if $\mathrm{d}\bar{\mathbf{x}}^\intercal/\mathrm{d}\mathbf{y}|_{\mathbf{y}=\bar{\mathbf{y}}} \neq \mathbf{0}$).

Now, we write an alternative factorization of $\mathbf{L}_\mathbf{z}$ in terms of $\mathbf{H}_{\hat{\mathbf{z}}}$. Using Layer 4, Eq. S8 and Layer 5, Eq. S5, consider the matrix

$$\left( \frac{\mathrm{s}\mathbf{z}}{\mathrm{s}\mathbf{z}^\intercal} \mathbf{H}_{\hat{\mathbf{z}}} \frac{\mathrm{d}\mathbf{z}^\intercal}{\mathrm{d}\mathbf{z}} \right) \bigg|_{\mathbf{y}=\bar{\mathbf{y}}} = \left[ \begin{pmatrix} \frac{\mathrm{s}\mathbf{x}}{\mathrm{s}\mathbf{x}^\intercal} & \frac{\mathrm{s}\mathbf{x}}{\mathrm{s}\mathbf{y}^\intercal} \\ \mathbf{0} & \mathbf{I} \end{pmatrix} \begin{pmatrix} \mathbf{0} & \mathbf{0} \\ \mathbf{0} & \mathbf{H}_\mathbf{y} \end{pmatrix} \begin{pmatrix} \frac{\mathrm{d}\mathbf{x}^\intercal}{\mathrm{d}\mathbf{x}} & \mathbf{0} \\ \frac{\mathrm{d}\mathbf{x}^\intercal}{\mathrm{d}\mathbf{y}} & \mathbf{I} \end{pmatrix} \right] \bigg|_{\mathbf{y}=\bar{\mathbf{y}}}.$$

Doing the matrix multiplication yields

$$\left( \frac{\mathrm{s}\mathbf{z}}{\mathrm{s}\mathbf{z}^\intercal} \mathbf{H}_{\hat{\mathbf{z}}} \frac{\mathrm{d}\mathbf{z}^\intercal}{\mathrm{d}\mathbf{z}} \right) \bigg|_{\mathbf{y}=\bar{\mathbf{y}}} = \left[ \begin{pmatrix} \frac{\mathrm{s}\mathbf{x}}{\mathrm{s}\mathbf{x}^\intercal} & \frac{\mathrm{s}\mathbf{x}}{\mathrm{s}\mathbf{y}^\intercal} \\ \mathbf{0} & \mathbf{I} \end{pmatrix} \begin{pmatrix} \mathbf{0} & \mathbf{0} \\ \mathbf{H}_\mathbf{y} \frac{\mathrm{d}\mathbf{x}^\intercal}{\mathrm{d}\mathbf{y}} & \mathbf{H}_\mathbf{y} \end{pmatrix} \right] \bigg|_{\mathbf{y}=\bar{\mathbf{y}}}$$

$$= \left.\begin{pmatrix} \frac{\mathrm{s}\mathbf{x}}{\mathrm{s}\mathbf{y}^\intercal}\mathbf{H_y}\frac{\mathrm{d}\mathbf{x}^\intercal}{\mathrm{d}\mathbf{y}} & \frac{\mathrm{s}\mathbf{x}}{\mathrm{s}\mathbf{y}^\intercal}\mathbf{H_y} \\ \mathbf{H_y}\frac{\mathrm{d}\mathbf{x}^\intercal}{\mathrm{d}\mathbf{y}} & \mathbf{H_y} \end{pmatrix}\right|_{\mathbf{y}=\bar{\mathbf{y}}}.$$

Using Layer 5, Eq. S3b, we have

$$\left.\left(\frac{\mathrm{s}\mathbf{z}}{\mathrm{s}\mathbf{z}^\intercal}\mathbf{H}_{\hat{\mathbf{z}}}\frac{\mathrm{d}\mathbf{z}^\intercal}{\mathrm{d}\mathbf{z}}\right)\right|_{\mathbf{y}=\bar{\mathbf{y}}} = \left.\begin{pmatrix} \frac{\mathrm{s}\mathbf{x}}{\mathrm{s}\mathbf{y}^\intercal}\mathbf{H_y}\frac{\mathrm{d}\mathbf{x}^\intercal}{\mathrm{d}\mathbf{y}} & \frac{\mathrm{s}\mathbf{x}}{\mathrm{s}\mathbf{y}^\intercal}\mathbf{H_y}\frac{\mathrm{d}\mathbf{y}^\intercal}{\mathrm{d}\mathbf{y}} \\ \frac{\mathrm{s}\mathbf{y}}{\mathrm{s}\mathbf{y}^\intercal}\mathbf{H_y}\frac{\mathrm{d}\mathbf{x}^\intercal}{\mathrm{d}\mathbf{y}} & \frac{\mathrm{s}\mathbf{y}}{\mathrm{s}\mathbf{y}^\intercal}\mathbf{H_y}\frac{\mathrm{d}\mathbf{y}^\intercal}{\mathrm{d}\mathbf{y}} \end{pmatrix}\right|_{\mathbf{y}=\bar{\mathbf{y}}}.$$

Notice that the matrix on the right-hand side is

$$\left.\left(\frac{\mathrm{s}\mathbf{z}}{\mathrm{s}\mathbf{y}^\intercal}\mathbf{H_y}\frac{\mathrm{d}\mathbf{z}^\intercal}{\mathrm{d}\mathbf{y}}\right)\right|_{\mathbf{y}=\bar{\mathbf{y}}} = \mathbf{L_z}.$$

Hence, we obtain an alternative factorization for $\mathbf{L_z}$ as

$$\mathbf{L_z} = \left.\left(\frac{\mathrm{s}\mathbf{z}}{\mathrm{s}\mathbf{z}^\intercal}\mathbf{H}_{\hat{\mathbf{z}}}\frac{\mathrm{d}\mathbf{z}^\intercal}{\mathrm{d}\mathbf{z}}\right)\right|_{\mathbf{y}=\bar{\mathbf{y}}}.$$

Thus, we can write the selection response of the geno-phenotype (in the form of Layer 7, Eq. 4) as

$$\iota\mathbf{L_z}\left.\frac{\delta w}{\delta \mathbf{z}}\right|_{\mathbf{y}=\bar{\mathbf{y}}} = \iota\left.\left(\frac{\mathrm{s}\mathbf{z}}{\mathrm{s}\mathbf{z}^\intercal}\mathbf{H}_{\hat{\mathbf{z}}}\frac{\mathrm{d}\mathbf{z}^\intercal}{\mathrm{d}\mathbf{z}}\frac{\delta w}{\delta \mathbf{z}}\right)\right|_{\mathbf{y}=\bar{\mathbf{y}}}.$$

Using the relationship between the total and total immediate selection gradients of the geno-phenotype (second line of Layer 4, Eq. S23), this becomes

$$\iota\mathbf{L_z}\left.\frac{\delta w}{\delta \mathbf{z}}\right|_{\mathbf{y}=\bar{\mathbf{y}}} = \iota\left.\left(\frac{\mathrm{s}\mathbf{z}}{\mathrm{s}\mathbf{z}^\intercal}\mathbf{H}_{\hat{\mathbf{z}}}\frac{\mathrm{d}w}{\mathrm{d}\mathbf{z}}\right)\right|_{\mathbf{y}=\bar{\mathbf{y}}}.$$

We can further simplify this equation by noticing the following. Using Layer 6, Eq. 10 and $\hat{\mathbf{z}} = (\bar{\mathbf{x}};\mathbf{y})$, we have that the *mechanistic additive socio-genetic cross-covariance matrix of the geno-phenotype and the unperturbed geno-phenotype* is

$$\mathbf{L}_{\mathbf{z}\hat{\mathbf{z}}} = \left.\left(\frac{\mathrm{s}\mathbf{z}}{\mathrm{s}\mathbf{y}^\intercal}\mathbf{H_y}\frac{\mathrm{d}\hat{\mathbf{z}}^\intercal}{\mathrm{d}\mathbf{y}}\right)\right|_{\mathbf{y}=\bar{\mathbf{y}}} \in \mathbb{R}^{N_\mathrm{a}(N_\mathrm{p}+N_\mathrm{g})\times N_\mathrm{a}(N_\mathrm{p}+N_\mathrm{g})}. \tag{Eq. S5.7.3}$$

Expanding, we have

$$\mathbf{L}_{\mathbf{z}\hat{\mathbf{z}}} = \left.\left[\begin{pmatrix} \frac{\mathrm{s}\mathbf{x}}{\mathrm{s}\mathbf{y}^\intercal} \\ \frac{\mathrm{s}\mathbf{y}}{\mathrm{s}\mathbf{y}^\intercal} \end{pmatrix}\mathbf{H_y}\begin{pmatrix} \frac{\mathrm{d}\bar{\mathbf{x}}^\intercal}{\mathrm{d}\mathbf{y}} & \frac{\mathrm{d}\mathbf{y}^\intercal}{\mathrm{d}\mathbf{y}} \end{pmatrix}\right]\right|_{\mathbf{y}=\bar{\mathbf{y}}}.$$

Using Layer 5, Eq. S3b and since the resident phenotype does not depend on mutant genotype, then

$$\mathbf{L}_{\mathbf{z}\hat{\mathbf{z}}} = \left.\left[\begin{pmatrix} \frac{\mathrm{s}\mathbf{x}}{\mathrm{s}\mathbf{y}^\intercal} \\ \mathbf{I} \end{pmatrix}\mathbf{H_y}\begin{pmatrix} \mathbf{0} & \mathbf{I} \end{pmatrix}\right]\right|_{\mathbf{y}=\bar{\mathbf{y}}}.$$

Doing the matrix multiplication yields

$$\mathbf{L}_{\mathbf{z}\hat{\mathbf{z}}} = \left.\left[\begin{pmatrix} \frac{\mathrm{s}\mathbf{x}}{\mathrm{s}\mathbf{y}^\intercal} \\ \mathbf{I} \end{pmatrix}\begin{pmatrix} \mathbf{0} & \mathbf{H_y} \end{pmatrix}\right]\right|_{\mathbf{y}=\bar{\mathbf{y}}} = \left.\begin{pmatrix} \mathbf{0} & \frac{\mathrm{s}\mathbf{x}}{\mathrm{s}\mathbf{y}^\intercal}\mathbf{H_y} \\ \mathbf{0} & \mathbf{H_y} \end{pmatrix}\right|_{\mathbf{y}=\bar{\mathbf{y}}}. \tag{Eq. S5.7.4}$$

Notice that the last matrix equals

$$\left.\left(\frac{\mathrm{s}\mathbf{z}}{\mathrm{s}\mathbf{z}^\intercal}\mathbf{H}_{\hat{\mathbf{z}}}\right)\right|_{\mathbf{y}=\bar{\mathbf{y}}}.$$

We can then write the evolutionary dynamics of the resident geno-phenotype $\bar{\mathbf{z}}$ in terms of the total selection gradient of the geno-phenotype as

$$\frac{\mathrm{d}\bar{\mathbf{z}}}{\mathrm{d}\tau} \approx \left.\left(\iota\mathbf{L}_{\mathbf{z}\hat{\mathbf{z}}}\frac{\mathrm{d}w}{\mathrm{d}\mathbf{z}} + \frac{\mathrm{s}\mathbf{z}}{\mathrm{s}\boldsymbol{\epsilon}^\intercal}\frac{\partial\bar{\boldsymbol{\epsilon}}}{\partial\tau}\right)\right|_{\mathbf{y}=\bar{\mathbf{y}}}. \tag{Eq. S5.7.5}$$

The cross-covariance matrix $\mathbf{L}_{\mathbf{z}\hat{\mathbf{z}}}$ is singular because $\mathrm{d}\hat{\mathbf{z}}^\intercal/\mathrm{d}\mathbf{y}|_{\mathbf{y}=\bar{\mathbf{y}}}$ has fewer rows than columns since the unperturbed geno-phenotype includes the genotype (Eq. S5.7.3). For this reason, $\mathbf{L}_{\mathbf{z}\hat{\mathbf{z}}}$ would still be singular even if the zero block entries in Eq. S5.7.4 were non-zero (i.e., if $\mathrm{d}\bar{\mathbf{x}}^\intercal/\mathrm{d}\mathbf{y}|_{\mathbf{y}=\bar{\mathbf{y}}} \neq \mathbf{0}$). Then, evolutionary equilibria of the geno-phenotype do not imply absence of total selection on the geno-phenotype, even if exogenous plastic response is absent.

#### S5.8 Evolutionary dynamics of the environment

Here we derive equations describing the evolutionary dynamics of the resident environment.

##### S5.8.1 In terms of joint direct niche construction

In this section, we obtain an equation for the evolutionary dynamics of the resident environment in terms of joint direct niche construction. Let $\bar{\mathbf{z}}(\tau)$ be the resident geno-phenotype at evolutionary time $\tau$, specifically at the point where the socio-devo stable resident is at carrying capacity, marked in Fig. 3. From the environmental constraint (2), the $i$-th environmental trait experienced by a mutant of age $a$ at such evolutionary time $\tau$ is $\epsilon_{ia} = h_{ia}(\mathbf{z}_a(\tau), \bar{\mathbf{z}}(\tau), \tau))$. Then, evolutionary change in the $i$-th environmental trait experienced by residents at age $a \in \{1,\ldots,N_\mathrm{a}\}$ is

$$\frac{\Delta\bar{\epsilon}_{ia}}{\Delta\tau} = \frac{1}{\Delta\tau}\left[h_{ia}(\mathbf{z}_a(\tau+\Delta\tau), \bar{\mathbf{z}}(\tau+\Delta\tau), \tau+\Delta\tau) - h_{ai}(\mathbf{z}_a(\tau), \bar{\mathbf{z}}(\tau), \tau)\right]\Bigg|_{\mathbf{y}=\bar{\mathbf{y}}}.$$

Taking the limit as $\Delta\tau \to 0$, this becomes

$$\frac{\mathrm{d}\bar{\epsilon}_{ia}}{\mathrm{d}\tau} = \left.\frac{\mathrm{d}h_{ia}(\mathbf{z}_a(\tau), \bar{\mathbf{z}}(\tau), \tau)}{\mathrm{d}\tau}\right|_{\mathbf{y}=\bar{\mathbf{y}}}.$$

Applying the chain rule, we obtain

$$\frac{\mathrm{d}\bar{\epsilon}_{ia}}{\mathrm{d}\tau} = \left(\sum_{j=1}^{N_\mathrm{p}}\frac{\partial h_{ia}}{\partial x_{ja}}\frac{\mathrm{d}x_{ja}}{\mathrm{d}\tau} + \sum_{j=1}^{N_\mathrm{g}}\frac{\partial h_{ia}}{\partial y_{ja}}\frac{\mathrm{d}y_{ja}}{\mathrm{d}\tau} + \sum_{k=1}^{N_\mathrm{a}}\sum_{j=1}^{N_\mathrm{p}}\frac{\partial h_{ia}}{\partial \bar{x}_{jk}}\frac{\mathrm{d}\bar{x}_{jk}}{\mathrm{d}\tau} + \sum_{k=1}^{N_\mathrm{a}}\sum_{j=1}^{N_\mathrm{g}}\frac{\partial h_{ia}}{\partial \bar{y}_{jk}}\frac{\mathrm{d}\bar{y}_{jk}}{\mathrm{d}\tau} + \frac{\partial h_{ia}}{\partial\tau}\right)\Bigg|_{\mathbf{y}=\bar{\mathbf{y}}}.$$

Applying matrix calculus notation, this is

$$\frac{\mathrm{d}\bar{\epsilon}_{ia}}{\mathrm{d}\tau} = \left(\frac{\partial h_{ia}}{\partial \mathbf{x}_a^\intercal}\frac{\mathrm{d}\mathbf{x}_a}{\mathrm{d}\tau} + \frac{\partial h_{ia}}{\partial \mathbf{y}_a^\intercal}\frac{\mathrm{d}\mathbf{y}_a}{\mathrm{d}\tau} + \sum_{k=1}^{N_\mathrm{a}}\frac{\partial h_{ia}}{\partial \bar{\mathbf{x}}_k^\intercal}\frac{\mathrm{d}\bar{\mathbf{x}}_k}{\mathrm{d}\tau}\right.$$

$$+\sum_{k=1}^{N_\mathrm{a}}\frac{\partial h_{ia}}{\partial \bar{\mathbf{y}}_k^\intercal}\frac{\mathrm{d}\bar{\mathbf{y}}_k}{\mathrm{d}\tau}+\frac{\partial h_{ia}}{\partial \tau}\bigg)\bigg|_{\mathbf{y}=\bar{\mathbf{y}}}.$$

Applying matrix calculus notation again yields

$$\frac{\mathrm{d}\bar{\epsilon}_{ia}}{\mathrm{d}\tau}=\bigg(\frac{\partial h_{ia}}{\partial \mathbf{x}_a^\intercal}\frac{\mathrm{d}\mathbf{x}_a}{\mathrm{d}\tau}+\frac{\partial h_{ia}}{\partial \mathbf{y}_a^\intercal}\frac{\mathrm{d}\mathbf{y}_a}{\mathrm{d}\tau}+\frac{\partial h_{ia}}{\partial \bar{\mathbf{x}}^\intercal}\frac{\mathrm{d}\bar{\mathbf{x}}}{\mathrm{d}\tau}+\frac{\partial h_{ia}}{\partial \bar{\mathbf{y}}^\intercal}\frac{\mathrm{d}\bar{\mathbf{y}}}{\mathrm{d}\tau}$$
$$+\frac{\partial h_{ia}}{\partial \tau}\bigg)\bigg|_{\mathbf{y}=\bar{\mathbf{y}}}.$$

Rewriting $h_{ia}$ as $\epsilon_{ia}$, we obtain

$$\frac{\mathrm{d}\bar{\epsilon}_{ia}}{\mathrm{d}\tau}=\bigg(\frac{\partial \epsilon_{ia}}{\partial \mathbf{x}_a^\intercal}\frac{\mathrm{d}\mathbf{x}_a}{\mathrm{d}\tau}+\frac{\partial \epsilon_{ia}}{\partial \mathbf{y}_a^\intercal}\frac{\mathrm{d}\mathbf{y}_a}{\mathrm{d}\tau}+\frac{\partial \epsilon_{ia}}{\partial \bar{\mathbf{x}}^\intercal}\frac{\mathrm{d}\bar{\mathbf{x}}}{\mathrm{d}\tau}+\frac{\partial \epsilon_{ia}}{\partial \bar{\mathbf{y}}^\intercal}\frac{\mathrm{d}\bar{\mathbf{y}}}{\mathrm{d}\tau}$$
$$+\frac{\partial \epsilon_{ia}}{\partial \tau}\bigg)\bigg|_{\mathbf{y}=\bar{\mathbf{y}}}.$$

Hence, for all environmental traits at age $a$, we have

$$\frac{\mathrm{d}\bar{\boldsymbol{\epsilon}}_a}{\mathrm{d}\tau}=\left(\frac{\partial \boldsymbol{\epsilon}_a}{\partial \mathbf{x}_a^\intercal}\frac{\mathrm{d}\mathbf{x}_a}{\mathrm{d}\tau}+\frac{\partial \boldsymbol{\epsilon}_a}{\partial \mathbf{y}_a^\intercal}\frac{\mathrm{d}\mathbf{y}_a}{\mathrm{d}\tau}+\frac{\partial \boldsymbol{\epsilon}_a}{\partial \bar{\mathbf{x}}^\intercal}\frac{\mathrm{d}\bar{\mathbf{x}}}{\mathrm{d}\tau}+\frac{\partial \boldsymbol{\epsilon}_a}{\partial \bar{\mathbf{y}}^\intercal}\frac{\mathrm{d}\bar{\mathbf{y}}}{\mathrm{d}\tau}+\frac{\partial \boldsymbol{\epsilon}_a}{\partial \tau}\right)\bigg|_{\mathbf{y}=\bar{\mathbf{y}}}.$$

Note that evaluation of $\mathrm{d}\mathbf{z}_a/\mathrm{d}\tau$ and $\partial\boldsymbol{\epsilon}_a/\partial\tau$ at $\mathbf{y}=\bar{\mathbf{y}}$ is $\mathrm{d}\bar{\mathbf{z}}_a/\mathrm{d}\tau$ and $\partial\bar{\boldsymbol{\epsilon}}_a/\partial\tau$, respectively. Also note that using Layer 2, Eq. S2d and Layer 2, Eq. S2d yields

$$\frac{\partial \boldsymbol{\epsilon}_a}{\partial \mathbf{x}^\intercal}\frac{\mathrm{d}\bar{\mathbf{x}}}{\mathrm{d}\tau}=\sum_{j=1}^{N_\mathrm{a}}\frac{\partial \boldsymbol{\epsilon}_a}{\partial \mathbf{x}_j^\intercal}\frac{\mathrm{d}\bar{\mathbf{x}}_j}{\mathrm{d}\tau}=\frac{\partial \boldsymbol{\epsilon}_a}{\partial \mathbf{x}_a^\intercal}\frac{\mathrm{d}\bar{\mathbf{x}}_a}{\mathrm{d}\tau}$$
$$\frac{\partial \boldsymbol{\epsilon}_a}{\partial \mathbf{y}^\intercal}\frac{\mathrm{d}\bar{\mathbf{y}}}{\mathrm{d}\tau}=\sum_{j=1}^{N_\mathrm{a}}\frac{\partial \boldsymbol{\epsilon}_a}{\partial \mathbf{y}_j^\intercal}\frac{\mathrm{d}\bar{\mathbf{y}}_j}{\mathrm{d}\tau}=\frac{\partial \boldsymbol{\epsilon}_a}{\partial \mathbf{y}_a^\intercal}\frac{\mathrm{d}\bar{\mathbf{y}}_a}{\mathrm{d}\tau}.$$

Then, we have

$$\frac{\mathrm{d}\bar{\boldsymbol{\epsilon}}_a}{\mathrm{d}\tau}=\left(\frac{\partial \boldsymbol{\epsilon}_a}{\partial \mathbf{x}^\intercal}\frac{\mathrm{d}\bar{\mathbf{x}}}{\mathrm{d}\tau}+\frac{\partial \boldsymbol{\epsilon}_a}{\partial \mathbf{y}^\intercal}\frac{\mathrm{d}\bar{\mathbf{y}}}{\mathrm{d}\tau}+\frac{\partial \boldsymbol{\epsilon}_a}{\partial \bar{\mathbf{x}}^\intercal}\frac{\mathrm{d}\bar{\mathbf{x}}}{\mathrm{d}\tau}+\frac{\partial \boldsymbol{\epsilon}_a}{\partial \bar{\mathbf{y}}^\intercal}\frac{\mathrm{d}\bar{\mathbf{y}}}{\mathrm{d}\tau}+\frac{\partial \bar{\boldsymbol{\epsilon}}_a}{\partial \tau}\right)\bigg|_{\mathbf{y}=\bar{\mathbf{y}}}.$$

Now note that $\partial\boldsymbol{\epsilon}_a/\partial\mathbf{z}^\intercal=(\partial\boldsymbol{\epsilon}_a/\partial\mathbf{x}^\intercal,\partial\boldsymbol{\epsilon}_a/\partial\mathbf{y}^\intercal)$, so

$$\frac{\mathrm{d}\bar{\boldsymbol{\epsilon}}_a}{\mathrm{d}\tau}=\left(\frac{\partial \boldsymbol{\epsilon}_a}{\partial \mathbf{z}^\intercal}\frac{\mathrm{d}\bar{\mathbf{z}}}{\mathrm{d}\tau}+\frac{\partial \boldsymbol{\epsilon}_a}{\partial \bar{\mathbf{z}}^\intercal}\frac{\mathrm{d}\bar{\mathbf{z}}}{\mathrm{d}\tau}+\frac{\partial \bar{\boldsymbol{\epsilon}}_a}{\partial \tau}\right)\bigg|_{\mathbf{y}=\bar{\mathbf{y}}}.$$

Hence, for all environmental traits over all ages, we have

$$\frac{\mathrm{d}\bar{\boldsymbol{\epsilon}}}{\mathrm{d}\tau}=\left(\frac{\partial \boldsymbol{\epsilon}}{\partial \mathbf{z}^\intercal}\frac{\mathrm{d}\bar{\mathbf{z}}}{\mathrm{d}\tau}+\frac{\partial \boldsymbol{\epsilon}}{\partial \bar{\mathbf{z}}^\intercal}\frac{\mathrm{d}\bar{\mathbf{z}}}{\mathrm{d}\tau}+\frac{\partial \bar{\boldsymbol{\epsilon}}}{\partial \tau}\right)\bigg|_{\mathbf{y}=\bar{\mathbf{y}}}$$
$$=\left[\left(\frac{\partial \boldsymbol{\epsilon}}{\partial \mathbf{z}^\intercal}+\frac{\partial \boldsymbol{\epsilon}}{\partial \bar{\mathbf{z}}^\intercal}\right)\frac{\mathrm{d}\bar{\mathbf{z}}}{\mathrm{d}\tau}+\frac{\partial \bar{\boldsymbol{\epsilon}}}{\partial \tau}\right]\bigg|_{\mathbf{y}=\bar{\mathbf{y}}},$$

where we use Layer 2, Eq. S7 and the block matrix of direct effects of social partners' geno-phenotype on a mutant's environment (Layer 2, Eq. S8; see also Layer 2, Eq. S5).

Let us momentarily write $\boldsymbol{\epsilon}=\tilde{\mathbf{h}}(\mathbf{z},\bar{\mathbf{z}})$ for some continuously differentiable function $\tilde{\mathbf{h}}$ to highlight the dependence of a mutant's environment $\boldsymbol{\epsilon}$ on her geno-phenotype $\mathbf{z}$ and on the geno-phenotype $\bar{\mathbf{z}}$ of resident social partners. Consider the environment that a resident experiences when she is in the context of mutants, denoted by $\check{\boldsymbol{\epsilon}}=\tilde{\mathbf{h}}(\bar{\mathbf{z}},\mathbf{z})$. Hence,

$$\frac{\partial\check{\boldsymbol{\epsilon}}}{\partial\mathbf{z}^\intercal}\bigg|_{\mathbf{y}=\bar{\mathbf{y}}}=\frac{\partial\tilde{\mathbf{h}}(\bar{\mathbf{z}},\mathbf{z})}{\partial\mathbf{z}^\intercal}\bigg|_{\mathbf{y}=\bar{\mathbf{y}}}=\frac{\partial\tilde{\mathbf{h}}(\mathbf{z},\bar{\mathbf{z}})}{\partial\bar{\mathbf{z}}^\intercal}\bigg|_{\mathbf{y}=\bar{\mathbf{y}}}=\frac{\partial\boldsymbol{\epsilon}}{\partial\bar{\mathbf{z}}^\intercal}\bigg|_{\mathbf{y}=\bar{\mathbf{y}}},\tag{Eq. S5.8.1}$$

where the second equality follows by exchanging dummy variables. Then, the transpose of the matrix of *direct social effects of a mutant's geno-phenotype on her and a partner's environment* is

$$\frac{\partial(\boldsymbol{\epsilon}+\check{\boldsymbol{\epsilon}})}{\partial\mathbf{z}^\intercal}\bigg|_{\mathbf{y}=\bar{\mathbf{y}}}=\left(\frac{\partial\boldsymbol{\epsilon}}{\partial\mathbf{z}^\intercal}+\frac{\partial\check{\boldsymbol{\epsilon}}}{\partial\mathbf{z}^\intercal}\right)\bigg|_{\mathbf{y}=\bar{\mathbf{y}}}=\left(\frac{\partial\boldsymbol{\epsilon}}{\partial\mathbf{z}^\intercal}+\frac{\partial\boldsymbol{\epsilon}}{\partial\bar{\mathbf{z}}^\intercal}\right)\bigg|_{\mathbf{y}=\bar{\mathbf{y}}}$$
$$\in\mathbb{R}^{N_\mathrm{a}N_\mathrm{e}\times N_\mathrm{a}(N_\mathrm{p}+N_\mathrm{g})}.\tag{Eq. S5.8.2}$$

Similarly, the transpose of the matrix of *direct social effects of a mutant's phenotype on her and a partner's environment* is

$$\frac{\partial(\boldsymbol{\epsilon}+\check{\boldsymbol{\epsilon}})}{\partial\mathbf{x}^\intercal}\bigg|_{\mathbf{y}=\bar{\mathbf{y}}}=\left(\frac{\partial\boldsymbol{\epsilon}}{\partial\mathbf{x}^\intercal}+\frac{\partial\check{\boldsymbol{\epsilon}}}{\partial\mathbf{x}^\intercal}\right)\bigg|_{\mathbf{y}=\bar{\mathbf{y}}}=\left(\frac{\partial\boldsymbol{\epsilon}}{\partial\mathbf{x}^\intercal}+\frac{\partial\boldsymbol{\epsilon}}{\partial\bar{\mathbf{x}}^\intercal}\right)\bigg|_{\mathbf{y}=\bar{\mathbf{y}}}$$
$$\in\mathbb{R}^{N_\mathrm{a}N_\mathrm{e}\times N_\mathrm{a}N_\mathrm{p}},\tag{Eq. S5.8.3}$$

and the transpose of the matrix of *direct social effects of a mutant's genotype on her and a partner's environment* is

$$\frac{\partial(\boldsymbol{\epsilon}+\check{\boldsymbol{\epsilon}})}{\partial\mathbf{y}^\intercal}\bigg|_{\mathbf{y}=\bar{\mathbf{y}}}=\left(\frac{\partial\boldsymbol{\epsilon}}{\partial\mathbf{y}^\intercal}+\frac{\partial\check{\boldsymbol{\epsilon}}}{\partial\mathbf{y}^\intercal}\right)\bigg|_{\mathbf{y}=\bar{\mathbf{y}}}=\left(\frac{\partial\boldsymbol{\epsilon}}{\partial\mathbf{y}^\intercal}+\frac{\partial\boldsymbol{\epsilon}}{\partial\bar{\mathbf{y}}^\intercal}\right)\bigg|_{\mathbf{y}=\bar{\mathbf{y}}}$$
$$\in\mathbb{R}^{N_\mathrm{a}N_\mathrm{e}\times N_\mathrm{a}N_\mathrm{g}}.\tag{Eq. S5.8.4}$$

Consequently, the evolutionary dynamics of the resident environment satisfy Layer 7, Eq. 10.

##### S5.8.2 In terms of total genotypic selection

In this section, we obtain an equation for the evolutionary dynamics of the resident environment in terms of total genotypic selection. Using the expression for the evolutionary dynamics of the geno-phenotype (Layer 7, Eq. 5 for $\boldsymbol{\zeta}=\mathbf{z}$) in that for the environment (Layer 7, Eq. 10) yields

$$\frac{\mathrm{d}\bar{\boldsymbol{\epsilon}}}{\mathrm{d}\tau}\approx\left[\frac{\partial(\boldsymbol{\epsilon}+\check{\boldsymbol{\epsilon}})}{\partial\mathbf{z}^\intercal}\left(\iota\mathbf{L}_{\mathbf{zy}}\frac{\mathrm{d}w}{\mathrm{d}\mathbf{y}}+\frac{\mathrm{s}\mathbf{z}}{\mathrm{s}\boldsymbol{\epsilon}^\intercal}\frac{\partial\boldsymbol{\epsilon}}{\partial\tau}\right)+\frac{\partial\boldsymbol{\epsilon}}{\partial\tau}\right]\bigg|_{\mathbf{y}=\bar{\mathbf{y}}}.$$

Using Layer 6, Eq. 13 for $\boldsymbol{\zeta}=\mathbf{z}$ yields

$$\frac{\mathrm{d}\bar{\boldsymbol{\epsilon}}}{\mathrm{d}\tau}\approx\left[\frac{\partial(\boldsymbol{\epsilon}+\check{\boldsymbol{\epsilon}})}{\partial\mathbf{z}^\intercal}\left(\iota\frac{\mathrm{s}\mathbf{z}}{\mathrm{s}\mathbf{y}^\intercal}\mathbf{H}_\mathbf{y}\frac{\mathrm{d}w}{\mathrm{d}\mathbf{y}}+\frac{\mathrm{s}\mathbf{z}}{\mathrm{s}\boldsymbol{\epsilon}^\intercal}\frac{\partial\boldsymbol{\epsilon}}{\partial\tau}\right)+\frac{\partial\boldsymbol{\epsilon}}{\partial\tau}\right]\bigg|_{\mathbf{y}=\bar{\mathbf{y}}}.$$

Collecting for $\partial\boldsymbol{\epsilon}/\partial\tau$ and using Layer 5, Eq. S6 yields

$$\frac{\mathrm{d}\bar{\boldsymbol{\epsilon}}}{\mathrm{d}\tau}\approx\left(\iota\frac{\mathrm{s}\boldsymbol{\epsilon}}{\mathrm{s}\mathbf{y}^\intercal}\mathbf{H}_\mathbf{y}\frac{\mathrm{d}w}{\mathrm{d}\mathbf{y}}+\frac{\mathrm{s}\boldsymbol{\epsilon}}{\mathrm{s}\boldsymbol{\epsilon}^\intercal}\frac{\partial\boldsymbol{\epsilon}}{\partial\tau}\right)\bigg|_{\mathbf{y}=\bar{\mathbf{y}}}.$$

Using Layer 6, Eq. 13 yields Layer 7, Eq. 5 for $\boldsymbol{\zeta}=\boldsymbol{\epsilon}$. Using the third line of Layer 4, Eq. S21 and Layer 6, Eq. 11 yields Layer 7, Eq. 4 for $\boldsymbol{\zeta}=\boldsymbol{\epsilon}$, whereas using the fourth line of Layer 4, Eq. S21 and Layer 6, Eq. 12 yields Layer 7, Eq. 1a for $\boldsymbol{\zeta}=\boldsymbol{\epsilon}$.

#### S5.9 Evolutionary dynamics of the geno-envo-phenotype

Here we obtain equations describing the evolutionary dynamics of the resident geno-envo-phenotype, that is, $\mathrm{d}\bar{\mathbf{m}}/\mathrm{d}\tau$.

##### S5.9.1 In terms of total genotypic selection

In this section, we obtain such an equation in terms of total genotypic selection. Since $\mathrm{d}\bar{\mathbf{m}}/\mathrm{d}\tau = (\mathrm{d}\bar{\mathbf{x}}/\mathrm{d}\tau;\mathrm{d}\bar{\mathbf{y}}/\mathrm{d}\tau;\mathrm{d}\bar{\boldsymbol{\epsilon}}/\mathrm{d}\tau)$, from Eq. S5.6.15, Eq. S2.2.2a, and Layer 7, Eq. 5 for $\boldsymbol{\zeta} = \boldsymbol{\epsilon}$, we can write the evolutionary dynamics of the resident geno-envo-phenotype $\bar{\mathbf{m}}$ as

$$\frac{\mathrm{d}\bar{\mathbf{m}}}{\mathrm{d}\tau} \approx \left[\iota\begin{pmatrix}\mathbf{L_{xy}}\\ \mathbf{H_y}\\ \mathbf{L_{\boldsymbol{\epsilon}y}}\end{pmatrix}\frac{\mathrm{d}w}{\mathrm{d}\mathbf{y}} + \begin{pmatrix}\frac{\mathbf{s}\mathbf{x}}{\mathbf{s}\boldsymbol{\epsilon}^\intercal}\\ \mathbf{0}\\ \frac{\mathbf{s}\boldsymbol{\epsilon}}{\mathbf{s}\boldsymbol{\epsilon}^\intercal}\end{pmatrix}\frac{\partial\bar{\boldsymbol{\epsilon}}}{\partial\tau}\right]\Bigg|_{\mathbf{y}=\bar{\mathbf{y}}}. \qquad \text{(Eq. S5.9.1)}$$

Using Layer 6, Eq. 10 and Layer 5, Eq. S3, this is

$$\frac{\mathrm{d}\bar{\mathbf{m}}}{\mathrm{d}\tau} \approx \left[\iota\begin{pmatrix}\frac{\mathbf{s}\mathbf{x}}{\mathbf{s}\mathbf{y}^\intercal}\\ \frac{\mathbf{s}\mathbf{y}}{\mathbf{s}\mathbf{y}^\intercal}\\ \frac{\mathbf{s}\boldsymbol{\epsilon}}{\mathbf{s}\mathbf{y}^\intercal}\end{pmatrix}\mathbf{H_y}\frac{\mathrm{d}w}{\mathrm{d}\mathbf{y}} + \begin{pmatrix}\frac{\mathbf{s}\mathbf{x}}{\mathbf{s}\boldsymbol{\epsilon}^\intercal}\\ \frac{\mathbf{s}\mathbf{y}}{\mathbf{s}\boldsymbol{\epsilon}^\intercal}\\ \frac{\mathbf{s}\boldsymbol{\epsilon}}{\mathbf{s}\boldsymbol{\epsilon}^\intercal}\end{pmatrix}\frac{\partial\bar{\boldsymbol{\epsilon}}}{\partial\tau}\right]\Bigg|_{\mathbf{y}=\bar{\mathbf{y}}}.$$

Using Layer 5, Eq. S7, this reduces to

$$\frac{\mathrm{d}\bar{\mathbf{m}}}{\mathrm{d}\tau} \approx \left(\iota\frac{\mathbf{s}\mathbf{m}}{\mathbf{s}\mathbf{y}^\intercal}\mathbf{H_y}\frac{\mathrm{d}w}{\mathrm{d}\mathbf{y}} + \frac{\mathbf{s}\mathbf{m}}{\mathbf{s}\boldsymbol{\epsilon}^\intercal}\frac{\partial\bar{\boldsymbol{\epsilon}}}{\partial\tau}\right)\Bigg|_{\mathbf{y}=\bar{\mathbf{y}}}.$$

Using Layer 6, Eq. 13 yields Layer 7, Eq. 5 for $\boldsymbol{\zeta} = \mathbf{m}$.. Using the third line of Layer 4, Eq. S21 and Layer 6, Eq. 11 yields Layer 7, Eq. 4 for $\boldsymbol{\zeta} = \mathbf{m}$, whereas using the fourth line of Layer 4, Eq. S21 and Layer 6, Eq. 12 yields Layer 7, Eq. 1a for $\boldsymbol{\zeta} = \mathbf{m}$.

In contrast to other arrangements, the premultiplying matrix $\mathbf{L_{my}}$ is non-singular if $\mathbf{H_y}$ is non-singular. Indeed, if

$$\frac{\mathbf{s}\mathbf{m}}{\mathbf{s}\mathbf{y}^\intercal}\Bigg|_{\mathbf{y}=\bar{\mathbf{y}}}\mathbf{r} = \mathbf{0}$$

for some vector $\mathbf{r}$, then from Layer 5, Eq. S7a and Layer 5, Eq. S3b we have

$$\begin{pmatrix}\frac{\mathbf{s}\mathbf{x}}{\mathbf{s}\mathbf{y}^\intercal}\\ \mathbf{I}\\ \frac{\mathbf{s}\boldsymbol{\epsilon}}{\mathbf{s}\mathbf{y}^\intercal}\end{pmatrix}\Bigg|_{\mathbf{y}=\bar{\mathbf{y}}}\mathbf{r} = \mathbf{0}.$$

Doing the multiplication yields

$$\begin{pmatrix}\frac{\mathbf{s}\mathbf{x}}{\mathbf{s}\mathbf{y}^\intercal}\Big|_{\mathbf{y}=\bar{\mathbf{y}}}\mathbf{r}\\ \mathbf{r}\\ \frac{\mathbf{s}\boldsymbol{\epsilon}}{\mathbf{s}\mathbf{y}^\intercal}\Big|_{\mathbf{y}=\bar{\mathbf{y}}}\mathbf{r}\end{pmatrix} = \mathbf{0},$$

which implies that $\mathbf{r} = \mathbf{0}$, so $\mathbf{s}\mathbf{m}/\mathbf{s}\mathbf{y}^\intercal|_{\mathbf{y}=\bar{\mathbf{y}}}$ is non-singular. Thus, $\mathbf{L_{my}}$ is non-singular if $\mathbf{H_y}$ is non-singular.

##### S5.9.2 In terms of total selection on the geno-envo-phenotype

In this section, we obtain an equation for the evolutionary dynamics of the resident geno-envo-phenotype in terms of the total selection gradient of the geno-envo-phenotype.

First, using Layer 6, Eq. 2, we define the *mechanistic additive genetic covariance matrix of the unperturbed geno-envo-phenotype* $\hat{\mathbf{m}} = (\bar{\mathbf{x}};\mathbf{y};\bar{\boldsymbol{\epsilon}})$ as

$$\mathbf{H}_{\hat{\mathbf{m}}} \equiv \mathrm{cov}[\mathbf{b}_{\hat{\mathbf{m}}},\mathbf{b}_{\hat{\mathbf{m}}}] = \left(\frac{\mathrm{d}\hat{\mathbf{m}}}{\mathrm{d}\mathbf{y}^\intercal}\mathbf{H_y}\frac{\mathrm{d}\hat{\mathbf{m}}^\intercal}{\mathrm{d}\mathbf{y}}\right)\Bigg|_{\mathbf{y}=\bar{\mathbf{y}}}$$
$$\in \mathbb{R}^{N_\mathrm{a}(N_\mathrm{p}+N_\mathrm{g}+N_\mathrm{e})\times N_\mathrm{a}(N_\mathrm{p}+N_\mathrm{g}+N_\mathrm{e})}.$$

By definition of $\hat{\mathbf{m}}$, we have

$$\mathbf{H}_{\hat{\mathbf{m}}} = \left[\begin{pmatrix}\frac{\mathrm{d}\bar{\mathbf{x}}}{\mathrm{d}\mathbf{y}^\intercal}\\ \frac{\mathrm{d}\mathbf{y}}{\mathrm{d}\mathbf{y}^\intercal}\\ \frac{\mathrm{d}\bar{\boldsymbol{\epsilon}}}{\mathrm{d}\mathbf{y}^\intercal}\end{pmatrix}\mathbf{H_y}\begin{pmatrix}\frac{\mathrm{d}\bar{\mathbf{x}}^\intercal}{\mathrm{d}\mathbf{y}} & \frac{\mathrm{d}\mathbf{y}^\intercal}{\mathrm{d}\mathbf{y}} & \frac{\mathrm{d}\bar{\boldsymbol{\epsilon}}^\intercal}{\mathrm{d}\mathbf{y}}\end{pmatrix}\right]\Bigg|_{\mathbf{y}=\bar{\mathbf{y}}}.$$

From Eq. S2.2.2c and Eq. S2.2.2d, the resident phenotype and environment are independent of the mutant genotype, so

$$\mathbf{H}_{\hat{\mathbf{m}}} = \left[\begin{pmatrix}\mathbf{0}\\ \mathbf{I}\\ \mathbf{0}\end{pmatrix}\mathbf{H_y}\begin{pmatrix}\mathbf{0} & \mathbf{I} & \mathbf{0}\end{pmatrix}\right]\Bigg|_{\mathbf{y}=\bar{\mathbf{y}}}.$$

Doing the matrix multiplication yields

$$\mathbf{H}_{\hat{\mathbf{m}}} = \left[\begin{pmatrix}\mathbf{0}\\ \mathbf{I}\\ \mathbf{0}\end{pmatrix}\begin{pmatrix}\mathbf{0} & \mathbf{H_y} & \mathbf{0}\end{pmatrix}\right]\Bigg|_{\mathbf{y}=\bar{\mathbf{y}}} = \begin{pmatrix}\mathbf{0} & \mathbf{0} & \mathbf{0}\\ \mathbf{0} & \mathbf{H_y} & \mathbf{0}\\ \mathbf{0} & \mathbf{0} & \mathbf{0}\end{pmatrix}. \qquad \text{(Eq. S5.9.2)}$$

The matrix $\mathbf{H}_{\hat{\mathbf{m}}}$ is singular because the unperturbed geno-envo-phenotype includes the genotype (i.e., $\mathrm{d}\hat{\mathbf{m}}^\intercal/\mathrm{d}\mathbf{y}|_{\mathbf{y}=\bar{\mathbf{y}}}$ has fewer rows than columns). For this reason, the matrix $\mathbf{H}_{\hat{\mathbf{m}}}$ would still be singular even if the zero block entries in Eq. S5.9.2 were non-zero (i.e., if $\mathrm{d}\bar{\mathbf{x}}^\intercal/\mathrm{d}\mathbf{y}|_{\mathbf{y}=\bar{\mathbf{y}}} \neq \mathbf{0}$ and $\mathrm{d}\bar{\boldsymbol{\epsilon}}^\intercal/\mathrm{d}\mathbf{y}|_{\mathbf{y}=\bar{\mathbf{y}}} \neq \mathbf{0}$).

Now, we write an alternative factorization of $\mathbf{L_m}$ in terms of $\mathbf{H}_{\hat{\mathbf{m}}}$. Using Layer 4, Eq. S17 and Layer 5, Eq. S8, we have

$$\left(\frac{\mathbf{s}\mathbf{m}}{\mathbf{s}\mathbf{m}^\intercal}\mathbf{H}_{\hat{\mathbf{m}}}\frac{\mathrm{d}\mathbf{m}^\intercal}{\mathrm{d}\mathbf{m}}\right)\Bigg|_{\mathbf{y}=\bar{\mathbf{y}}} = \left[\begin{pmatrix}\frac{\mathbf{s}\mathbf{x}}{\mathbf{s}\mathbf{x}^\intercal} & \frac{\mathbf{s}\mathbf{x}}{\mathbf{s}\mathbf{y}^\intercal} & \frac{\mathbf{s}\mathbf{x}}{\mathbf{s}\boldsymbol{\epsilon}^\intercal}\\ \mathbf{0} & \mathbf{I} & \mathbf{0}\\ \frac{\mathbf{s}\boldsymbol{\epsilon}}{\mathbf{s}\mathbf{x}^\intercal} & \frac{\mathbf{s}\boldsymbol{\epsilon}}{\mathbf{s}\mathbf{y}^\intercal} & \frac{\mathbf{s}\boldsymbol{\epsilon}}{\mathbf{s}\boldsymbol{\epsilon}^\intercal}\end{pmatrix}\begin{pmatrix}\mathbf{0} & \mathbf{0} & \mathbf{0}\\ \mathbf{0} & \mathbf{H_y} & \mathbf{0}\\ \mathbf{0} & \mathbf{0} & \mathbf{0}\end{pmatrix}\begin{pmatrix}\frac{\mathrm{d}\mathbf{x}^\intercal}{\mathrm{d}\mathbf{x}} & \mathbf{0} & \frac{\mathrm{d}\boldsymbol{\epsilon}^\intercal}{\mathrm{d}\mathbf{x}}\\ \frac{\mathrm{d}\mathbf{x}^\intercal}{\mathrm{d}\mathbf{y}} & \mathbf{I} & \frac{\mathrm{d}\boldsymbol{\epsilon}^\intercal}{\mathrm{d}\mathbf{y}}\\ \frac{\mathrm{d}\mathbf{x}^\intercal}{\mathrm{d}\boldsymbol{\epsilon}} & \mathbf{0} & \frac{\mathrm{d}\boldsymbol{\epsilon}^\intercal}{\mathrm{d}\boldsymbol{\epsilon}}\end{pmatrix}\right]\Bigg|_{\mathbf{y}=\bar{\mathbf{y}}}.$$

Doing the matrix multiplication yields

$$\left(\frac{\mathbf{sm}}{\mathbf{sm}^\intercal}\mathbf{H}_{\hat{\mathbf{m}}}\frac{\mathbf{dm}^\intercal}{\mathbf{dm}}\right)\Bigg|_{\mathbf{y}=\bar{\mathbf{y}}}$$

$$= \left[\begin{pmatrix} \frac{\mathbf{sx}}{\mathbf{sx}^\intercal} & \frac{\mathbf{sx}}{\mathbf{sy}^\intercal} & \frac{\mathbf{sx}}{\mathbf{s}\boldsymbol{\epsilon}^\intercal} \\ \mathbf{0} & \mathbf{I} & \mathbf{0} \\ \frac{\mathbf{s}\boldsymbol{\epsilon}}{\mathbf{sx}^\intercal} & \frac{\mathbf{s}\boldsymbol{\epsilon}}{\mathbf{sy}^\intercal} & \frac{\mathbf{s}\boldsymbol{\epsilon}}{\mathbf{s}\boldsymbol{\epsilon}^\intercal} \end{pmatrix} \begin{pmatrix} \mathbf{0} & \mathbf{0} & \mathbf{0} \\ \mathbf{H}_\mathbf{y}\frac{\mathbf{dx}^\intercal}{\mathbf{dy}} & \mathbf{H}_\mathbf{y} & \mathbf{H}_\mathbf{y}\frac{\mathbf{d}\boldsymbol{\epsilon}^\intercal}{\mathbf{dy}} \\ \mathbf{0} & \mathbf{0} & \mathbf{0} \end{pmatrix}\right]\Bigg|_{\mathbf{y}=\bar{\mathbf{y}}}$$

$$= \begin{pmatrix} \frac{\mathbf{sx}}{\mathbf{sy}^\intercal}\mathbf{H}_\mathbf{y}\frac{\mathbf{dx}^\intercal}{\mathbf{dy}} & \frac{\mathbf{sx}}{\mathbf{sy}^\intercal}\mathbf{H}_\mathbf{y} & \frac{\mathbf{sx}}{\mathbf{sy}^\intercal}\mathbf{H}_\mathbf{y}\frac{\mathbf{d}\boldsymbol{\epsilon}^\intercal}{\mathbf{dy}} \\ \mathbf{H}_\mathbf{y}\frac{\mathbf{dx}^\intercal}{\mathbf{dy}} & \mathbf{H}_\mathbf{y} & \mathbf{H}_\mathbf{y}\frac{\mathbf{d}\boldsymbol{\epsilon}^\intercal}{\mathbf{dy}} \\ \frac{\mathbf{s}\boldsymbol{\epsilon}}{\mathbf{sy}^\intercal}\mathbf{H}_\mathbf{y}\frac{\mathbf{dx}^\intercal}{\mathbf{dy}} & \frac{\mathbf{s}\boldsymbol{\epsilon}}{\mathbf{sy}^\intercal}\mathbf{H}_\mathbf{y} & \frac{\mathbf{s}\boldsymbol{\epsilon}}{\mathbf{sy}^\intercal}\mathbf{H}_\mathbf{y}\frac{\mathbf{d}\boldsymbol{\epsilon}^\intercal}{\mathbf{dy}} \end{pmatrix}\Bigg|_{\mathbf{y}=\bar{\mathbf{y}}}$$

Using Layer 5, Eq. S3b, this is

$$\left(\frac{\mathbf{sm}}{\mathbf{sm}^\intercal}\mathbf{H}_{\hat{\mathbf{m}}}\frac{\mathbf{dm}^\intercal}{\mathbf{dm}}\right)\Bigg|_{\mathbf{y}=\bar{\mathbf{y}}}$$

$$= \begin{pmatrix} \frac{\mathbf{sx}}{\mathbf{sy}^\intercal}\mathbf{H}_\mathbf{y}\frac{\mathbf{dx}^\intercal}{\mathbf{dy}} & \frac{\mathbf{sx}}{\mathbf{sy}^\intercal}\mathbf{H}_\mathbf{y}\frac{\mathbf{dy}^\intercal}{\mathbf{dy}} & \frac{\mathbf{sx}}{\mathbf{sy}^\intercal}\mathbf{H}_\mathbf{y}\frac{\mathbf{d}\boldsymbol{\epsilon}^\intercal}{\mathbf{dy}} \\ \frac{\mathbf{sy}}{\mathbf{sy}^\intercal}\mathbf{H}_\mathbf{y}\frac{\mathbf{dx}^\intercal}{\mathbf{dy}} & \frac{\mathbf{sy}}{\mathbf{sy}^\intercal}\mathbf{H}_\mathbf{y}\frac{\mathbf{dy}^\intercal}{\mathbf{dy}} & \frac{\mathbf{sy}}{\mathbf{sy}^\intercal}\mathbf{H}_\mathbf{y}\frac{\mathbf{d}\boldsymbol{\epsilon}^\intercal}{\mathbf{dy}} \\ \frac{\mathbf{s}\boldsymbol{\epsilon}}{\mathbf{sy}^\intercal}\mathbf{H}_\mathbf{y}\frac{\mathbf{dx}^\intercal}{\mathbf{dy}} & \frac{\mathbf{s}\boldsymbol{\epsilon}}{\mathbf{sy}^\intercal}\mathbf{H}_\mathbf{y}\frac{\mathbf{dy}^\intercal}{\mathbf{dy}} & \frac{\mathbf{s}\boldsymbol{\epsilon}}{\mathbf{sy}^\intercal}\mathbf{H}_\mathbf{y}\frac{\mathbf{d}\boldsymbol{\epsilon}^\intercal}{\mathbf{dy}} \end{pmatrix}\Bigg|_{\mathbf{y}=\bar{\mathbf{y}}}$$

Notice that the matrix on the right-hand side is

$$\left(\frac{\mathbf{sm}}{\mathbf{sy}^\intercal}\mathbf{H}_\mathbf{y}\frac{\mathbf{dm}^\intercal}{\mathbf{dy}}\right)\Bigg|_{\mathbf{y}=\bar{\mathbf{y}}} = \mathbf{L}_\mathbf{m}.$$

Hence, we obtain an alternative factorization for $\mathbf{L}_\mathbf{m}$ as

$$\mathbf{L}_\mathbf{m} = \left(\frac{\mathbf{sm}}{\mathbf{sm}^\intercal}\mathbf{H}_{\hat{\mathbf{m}}}\frac{\mathbf{dm}^\intercal}{\mathbf{dm}}\right)\Bigg|_{\mathbf{y}=\bar{\mathbf{y}}}.$$

We can now write the selection response of the geno-envo-phenotype (in the form of Layer 7, Eq. 1a) as

$$\iota\mathbf{L}_\mathbf{m}\frac{\partial w}{\partial\mathbf{m}}\bigg|_{\mathbf{y}=\bar{\mathbf{y}}} = \iota\left(\frac{\mathbf{sm}}{\mathbf{sm}^\intercal}\mathbf{H}_{\hat{\mathbf{m}}}\frac{\mathbf{dm}^\intercal}{\mathbf{dm}}\frac{\partial w}{\partial\mathbf{m}}\right)\bigg|_{\mathbf{y}=\bar{\mathbf{y}}}.$$

Using the relationship between the total and partial selection gradients of the geno-envo-phenotype (Layer 4, Eq. S24), this becomes

$$\iota\mathbf{L}_\mathbf{m}\frac{\partial w}{\partial\mathbf{m}}\bigg|_{\mathbf{y}=\bar{\mathbf{y}}} = \iota\left(\frac{\mathbf{sm}}{\mathbf{sm}^\intercal}\mathbf{H}_{\hat{\mathbf{m}}}\frac{\mathrm{d}w}{\mathbf{dm}}\right)\bigg|_{\mathbf{y}=\bar{\mathbf{y}}}.$$

We can further simplify this equation by noticing the following. Using Layer 6, Eq. 10 and $\hat{\mathbf{m}} = (\bar{\mathbf{x}};\mathbf{y};\bar{\boldsymbol{\epsilon}})$, we have that the *mechanistic additive socio-genetic cross-covariance matrix of the geno-envo-phenotype and the unperturbed geno-envo-phenotype* is

$$\mathbf{L}_{\mathbf{m}\hat{\mathbf{m}}} = \left(\frac{\mathbf{sm}}{\mathbf{sy}^\intercal}\mathbf{H}_\mathbf{y}\frac{\mathbf{d}\hat{\mathbf{m}}^\intercal}{\mathbf{dy}}\right)\bigg|_{\mathbf{y}=\bar{\mathbf{y}}} \in \mathbb{R}^{N_\mathrm{a}(N_\mathrm{p}+N_\mathrm{g}+N_\mathrm{e})\times N_\mathrm{a}(N_\mathrm{p}+N_\mathrm{g}+N_\mathrm{e})}. \quad \text{(Eq. S5.9.3)}$$

Expanding, we have

$$\mathbf{L}_{\mathbf{m}\hat{\mathbf{m}}} = \left[\begin{pmatrix} \frac{\mathbf{sx}}{\mathbf{sy}^\intercal} \\ \frac{\mathbf{sy}}{\mathbf{sy}^\intercal} \\ \frac{\mathbf{s}\boldsymbol{\epsilon}}{\mathbf{sy}^\intercal} \end{pmatrix}\mathbf{H}_\mathbf{y}\begin{pmatrix} \frac{\mathbf{d}\bar{\mathbf{x}}^\intercal}{\mathbf{dy}} & \frac{\mathbf{dy}^\intercal}{\mathbf{dy}} & \frac{\mathbf{d}\bar{\boldsymbol{\epsilon}}^\intercal}{\mathbf{dy}} \end{pmatrix}\right]\Bigg|_{\mathbf{y}=\bar{\mathbf{y}}}.$$

Using Layer 5, Eq. S3b and since the resident phenotype and environment do not depend on the mutant genotype, then

$$\mathbf{L}_{\mathbf{m}\hat{\mathbf{m}}} = \left[\begin{pmatrix} \frac{\mathbf{sx}}{\mathbf{sy}^\intercal} \\ \mathbf{I} \\ \frac{\mathbf{s}\boldsymbol{\epsilon}}{\mathbf{sy}^\intercal} \end{pmatrix}\mathbf{H}_\mathbf{y}\begin{pmatrix} \mathbf{0} & \mathbf{I} & \mathbf{0} \end{pmatrix}\right]\Bigg|_{\mathbf{y}=\bar{\mathbf{y}}}.$$

Doing the matrix multiplication yields

$$\mathbf{L}_{\mathbf{m}\hat{\mathbf{m}}} = \left[\begin{pmatrix} \frac{\mathbf{sx}}{\mathbf{sy}^\intercal} \\ \mathbf{I} \\ \frac{\mathbf{s}\boldsymbol{\epsilon}}{\mathbf{sy}^\intercal} \end{pmatrix}\begin{pmatrix} \mathbf{0} & \mathbf{H}_\mathbf{y} & \mathbf{0} \end{pmatrix}\right]\Bigg|_{\mathbf{y}=\bar{\mathbf{y}}}$$

$$= \begin{pmatrix} \mathbf{0} & \frac{\mathbf{sx}}{\mathbf{sy}^\intercal}\mathbf{H}_\mathbf{y} & \mathbf{0} \\ \mathbf{0} & \mathbf{H}_\mathbf{y} & \mathbf{0} \\ \mathbf{0} & \frac{\mathbf{s}\boldsymbol{\epsilon}}{\mathbf{sy}^\intercal}\mathbf{H}_\mathbf{y} & \mathbf{0} \end{pmatrix}\Bigg|_{\mathbf{y}=\bar{\mathbf{y}}}. \quad \text{(Eq. S5.9.4)}$$

Notice that the last matrix equals

$$\left(\frac{\mathbf{sm}}{\mathbf{sm}^\intercal}\mathbf{H}_{\hat{\mathbf{m}}}\right)\bigg|_{\mathbf{y}=\bar{\mathbf{y}}}.$$

Thus,

$$\mathbf{L}_{\mathbf{m}\hat{\mathbf{m}}} = \left(\frac{\mathbf{sm}}{\mathbf{sm}^\intercal}\mathbf{H}_{\hat{\mathbf{m}}}\right)\bigg|_{\mathbf{y}=\bar{\mathbf{y}}}.$$

We can then write the evolutionary dynamics of the resident geno-envo-phenotype $\bar{\mathbf{m}}$ in terms of the total selection gradient of the geno-envo-phenotype as

$$\frac{\mathbf{d}\bar{\mathbf{m}}}{\mathrm{d}\tau} \approx \left(\iota\mathbf{L}_{\mathbf{m}\hat{\mathbf{m}}}\frac{\mathrm{d}w}{\mathbf{dm}} + \frac{\mathbf{sm}}{\mathbf{s}\boldsymbol{\epsilon}^\intercal}\frac{\partial\bar{\boldsymbol{\epsilon}}}{\partial\tau}\right)\bigg|_{\mathbf{y}=\bar{\mathbf{y}}}. \quad \text{(Eq. S5.9.5)}$$

The cross-covariance matrix $\mathbf{L}_{\mathbf{m}\hat{\mathbf{m}}}$ is singular because $\mathbf{d}\hat{\mathbf{m}}^\intercal/\mathbf{dy}|_{\mathbf{y}=\bar{\mathbf{y}}}$ has fewer rows than columns since the unperturbed geno-envo-phenotype includes the genotype. For this reason, $\mathbf{L}_{\mathbf{m}\hat{\mathbf{m}}}$ would still be singular even if the zero block entries in Eq. S5.9.4 were non-zero (i.e., if $\mathbf{d}\bar{\mathbf{x}}^\intercal/\mathbf{dy}|_{\mathbf{y}=\bar{\mathbf{y}}} \neq \mathbf{0}$ and $\mathbf{d}\bar{\boldsymbol{\epsilon}}^\intercal/\mathbf{dy}|_{\mathbf{y}=\bar{\mathbf{y}}} \neq \mathbf{0}$). Then, evolutionary equilibria of the geno-envo-phenotype do not imply absence of total selection on the geno-envo-phenotype, even if exogenous plastic response is absent.
