## Supplementary material for "A mathematical framework for evo-devo dynamics": Computer Code

### Computer code for:
### A mathematical framework for evo-devo dynamics

Mauricio González – Forero*[1]
1 School of Biology, University of St Andrews, St Andrews, UK
*

This file contains the computer code used to generate the figures given in the main text. This code was prepared in Mathematica 12.1.1.0 and it is made available under a Creative Commons Attribution licence (CC BY).

#### Illustration of socio-devo instability

In[12]:=
```
Clear["Global`*"]
Nθ = 10;
Na = 4;
q = 0.5;
y[a_] := 0.5;
Table[x[1, θ] = 0.1, {θ, 1, Nθ}];
Table[x[a, 1] = 0.1, {a, 1, Na}];
xsol = Table[x[a + 1, θ + 1] = x[a, θ + 1] + y[a] (x[a, θ + 1] + q x[a + 1, θ]^2),
   {a, 1, Na - 1}, {θ, 1, Nθ - 1}];

PlotStyleFunction[θ_, Nθ_] := If[θ == Nθ, {Black, Thickness[0.01]},
  If[θ == 1, {Black, Dashing[0.05], Thickness[0.01]}, {Gray}]]

Export[StringJoin[ToString[NotebookDirectory[]],
   StringJoin["Fig.example.social.SDStabilisation"], ".pdf"],
  Show[Table[ListLinePlot[Table[x[a, θ], {a, 1, Na}],
     PlotStyle → PlotStyleFunction[θ, Nθ], AxesStyle → Large,
     PlotRange → {0, 0.5}, PlotStyle → {Black, Thickness[0.01]},
     PlotMarkers → {"●", Large}], {θ, 1, Nθ}]]];
```

In[22]:=
```
xm[1] = 0.1;
xmm[1] = 0.1;
ym[a_] := 0.6;
(*Obtain SDS resident*)
xb = Table[x[a, Nθ], {a, 1, Na}];
(*Compute mutant in the context of SDS resident*)
Table[xm[a + 1] = xm[a] + ym[a] (xm[a] + q xb[[a + 1]]^2), {a, 1, Na - 1}];
(*Compute mutant in the context of itself*)
Table[xmm[a + 1] = xmm[a] + ym[a] (xmm[a] + q xm[a + 1]^2), {a, 1, Na - 1}];
(*Plot SDS resident, mutant in context of SDS resident,
and mutant in the context of itself*)
Export[StringJoin[ToString[NotebookDirectory[]],
    StringJoin["Fig.example.social.SDNon-Eq"], ".pdf"],
  ListLinePlot[{Table[xb[[a]], {a, 1, Na}], Table[xm[a], {a, 1, Na}],
     Table[xmm[a], {a, 1, Na}]}, PlotStyle →
     {{Black, Dashing[0.05], Thickness[0.01]}, {Gray}, {Black, Thickness[0.01]}},
    AxesStyle → Large, PlotMarkers → {"●", Large}]];
```

### Non-social development

In[29]:=
```
Clear["Global`*"]

(*Choose parameters*)
NA = 4; (*number of ages*)
Pp = 0.7; (*survival probability*)
Finalτ = 200;

ι[Na_, p_] := Sum[p^(a-1), {a, 1, Na}]
T[Na_, p_] := Sum[a (1 - y[a]) x[a] × l[a, p], {a, 1, Na}]
partialwpartialx[Na_, p_] :=
 Table[1/T[Na, p] l[a, p] (1 - y[a]), {a, 1, Na}, {j, 1, 1}]
partialwpartialy[Na_, p_] := Table[-1/T[Na, p] l[a, p] × x[a], {a, 1, Na}, {j, 1, 1}]
partialxpartialy[Na_] := Table[Table[If[j == a + 1, x[a], 0], {j, 1, Na}], {a, 1, Na}]
partialxpartialx[Na_] :=
 Table[Table[If[j == a + 1, 1 + y[a], If[j == a, 1, 0]], {j, 1, Na}], {a, 1, Na}]
totalxtotalx[Na_] :=
 Table[Table[If[j > a, Product[partialxpartialx[Na][[k, k + 1]], {k, a, j - 1}],
    If[j == a, 1, 0]], {j, 1, Na}], {a, 1, Na}]
totalxtotaly[Na_] := partialxpartialy[Na].totalxtotalx[Na];
totalwtotalx[Na_, p_] := totalxtotalx[Na].partialwpartialx[Na, p]
totalwtotaly[Na_, p_] :=
 partialwpartialy[Na, p] + totalxtotaly[Na].partialwpartialx[Na, p]
```

```
l[a_, p_] := p^(a-1)
Hy[Na_, τ_] := Table[
  If[a == j, (ysol[τ]) [[a, 1]] (1 - (ysol[τ]) [[a, 1]]), 0], {a, 1, Na}, {j, 1, Na}]

(*Substitute solutions in total selection gradient*)
Totalwtotalx[Na_, τ_, p_] :=
 totalwtotalx[Na, p] /. Table[y[a] → ysol[τ][[a, 1]], {a, 1, Na}] /.
  Table[x[a] → xsol[τ, a, 1], {a, 1, Na}]
Totalwtotaly[Na_, τ_, p_] :=
 totalwtotaly[Na, p] /. Table[y[a] → ysol[τ][[a, 1]], {a, 1, Na}] /.
  Table[x[a] → xsol[τ, a, 1], {a, 1, Na}]

(*Initial conditions*)
ysol[1] = Table[1/2, {a, 1, 4}, {j, 1, 1}];

Runs[Na_, T_, p_] := Table[{xsol[τ, 1, 1] = 1;
   Table[xsol[τ, a + 1, 1] = (1 + ysol[τ][[a, 1]]) xsol[τ, a, 1], {a, 1, Na - 1}];
   ysol[τ + 1] = ysol[τ] + ι[Na, p] × Hy[Na, τ].Totalwtotaly[Na, τ, p];}, {τ, 1, T}]

(*Plots of runs*)
PlotStyleFunction[τ_] := If[τ == Finalτ, {Black, Thickness[0.01]},
  If[τ == 1, {Black, Dashing[0.05], Thickness[0.01]}, {Gray}]]
PlotRuns[Na_, T_, p_] := {Runs[Na, T, p];,
  Export[StringJoin[ToString[NotebookDirectory[]],
    StringJoin["Fig.example.control.p=", ToString[p]], ".pdf"],
   Show[Table[ListLinePlot[Flatten[ysol[τ]], PlotRange → {-0.1, 1.1},
      PlotStyle → PlotStyleFunction[τ], AxesStyle → Large,
      PlotMarkers → {"●", Large}], {τ, {1, 2, 5, 10, 20, Finalτ}}]]],
  Export[StringJoin[ToString[NotebookDirectory[]],
    StringJoin["Fig.example.state.p=", ToString[p]], ".pdf"],
   Show[Table[ListLinePlot[Table[xsol[τ, a, 1], {a, 1, Na}],
      PlotRange → {0, 5}, PlotStyle → PlotStyleFunction[τ], AxesStyle → Large,
      PlotMarkers → {"●", Large}], {τ, {1, 2, 5, 10, 20, Finalτ}}]]]}

(*Do runs and plots*)
PlotRuns[NA, Finalτ, Pp];

(*Hz plots*)

(*To fix color scale in matrix plot*)
cf = Blend[{{0., RGBColor[0.260487, 0.356, 0.891569]},
     {0.166667, RGBColor[0.230198, 0.499962, 0.848188]},
     {0.333333, RGBColor[0.392401, 0.658762, 0.797589]},
     {0.499999, RGBColor[0.964837, 0.982332, 0.98988]},
     {0.5, RGBColor[1, 1, 1]}, {0.500001, RGBColor[0.95735, 0.957281, 0.896269]},
     {0.666667, RGBColor[0.913252, 0.790646, 0.462837]},
     {0.833333, RGBColor[0.860243, 0.558831, 0.00695811]},
```

```
    {1., RGBColor[1., 0.42, 0.]}}, #1] &;
(*cfScaled=cf@Rescale[#,{-1,1},{0,1}]&;*)
cfScaled = cf@Rescale[#, {-4, 4}, {0, 1}] &;

(*Compute Hx*)
Totalxtotaly[Na_, τ_] :=
 totalxtotaly[Na] /. Table[y[a] → ysol[τ][[a, 1]], {a, 1, Na}] /.
  Table[x[a] → xsol[τ, a, 1], {a, 1, Na}]
Hx[Na_, τ_] := Transpose[Totalxtotaly[Na, τ]].Hy[Na, τ].Totalxtotaly[Na, τ]

(*Compute Hz*)
Totalztotaly[Na_, τ_] :=
 ArrayFlatten[{{Totalxtotaly[Na, τ], IdentityMatrix[Na]}}]
Hz[Na_, τ_] := Transpose[Totalztotaly[Na, τ]].Hy[Na, τ].Totalztotaly[Na, τ]

(*Plots of Hz*)
HzPlot[Na_, p_] := Table[Export[
   StringJoin[ToString[NotebookDirectory[]], StringJoin["Fig.example.Hz.p=",
     ToString[p], ".tau=", ToString[τ]], ".pdf"], MatrixPlot[Hz[Na, τ],
    FrameTicks → {{{{1, 1}, {2, 2}, {3, 3}, {4, 4}, {5, 1}, {6, 2}, {7, 3}, {8, 4}},
       {{1, 1}, {2, 2}, {3, 3}, {4, 4}, {5, 1}, {6, 2}, {7, 3}, {8, 4}}},
      {{{1, 1}, {2, 2}, {3, 3}, {4, 4}, {5, 1}, {6, 2}, {7, 3}, {8, 4}},
       {{1, 1}, {2, 2}, {3, 3}, {4, 4}, {5, 1}, {6, 2}, {7, 3}, {8, 4}}}},
    FrameStyle → Large, PlotLegends → BarLegend[{Automatic, {-4, 4}},
      LabelStyle → Large], ColorFunction → cfScaled,
    ColorFunctionScaling → False]], {τ, {1, 2, 5, 10, 20, Finalτ}}]

(*Do plots of Hz*)

HzPlot[NA, Pp];

(*Plots of the total selection gradients*)

Totalwtotalz[Na_, τ_, p_] :=
 ArrayFlatten[{{Totalwtotalx[Na, τ, p]}, {Totalwtotaly[Na, τ, p]}}]
dwdyxPlot[Na_, p_] := {Export[StringJoin[ToString[NotebookDirectory[]],
    StringJoin["Fig.example.dwdy.p=", ToString[p]], ".pdf"],
   Show[Table[ListLinePlot[Flatten[Totalwtotaly[Na, τ, p]],
      PlotStyle → PlotStyleFunction[τ], AxesStyle → Large,
      PlotRange → {-0.4, 0.1}, PlotStyle → {Black, Thickness[0.01]},
      PlotMarkers → {"●", Large}], {τ, {1, 2, 5, 10, 20, Finalτ}}]]],
  Export[StringJoin[ToString[NotebookDirectory[]],
    StringJoin["Fig.example.dwdx.p=", ToString[p]], ".pdf"],
   Show[Table[ListLinePlot[Flatten[Totalwtotalx[Na, τ, p]],
      PlotStyle → PlotStyleFunction[τ], AxesStyle → Large,
      PlotRange → {0, 0.5}, PlotStyle → {Black, Thickness[0.01]},
      PlotMarkers → {"●", Large}], {τ, {1, 2, 5, 10, 20, Finalτ}}]]]}
```

```
dwdyxPlot[NA, Pp];

(*Plots of the partial selection gradients*)

Partialwpartialx[Na_, τ_, p_] :=
 partialwpartialx[Na, p] /. Table[y[a] → ysol[τ][[a, 1]], {a, 1, Na}] /.
  Table[x[a] → xsol[τ, a, 1], {a, 1, Na}]
Partialwpartialy[Na_, τ_, p_] :=
 partialwpartialy[Na, p] /. Table[y[a] → ysol[τ][[a, 1]], {a, 1, Na}] /.
  Table[x[a] → xsol[τ, a, 1], {a, 1, Na}]

PartialwpartialyxPlot[Na_, p_] :=
 {Export[StringJoin[ToString[NotebookDirectory[]],
    StringJoin["Fig.example.partialwpartialy.p=", ToString[p]], ".pdf"],
   Show[Table[ListLinePlot[Flatten[Partialwpartialy[Na, τ, p]],
      PlotStyle → PlotStyleFunction[τ], AxesStyle → Large,
      PlotRange → {-0.4, 0.1}, PlotStyle → {Black, Thickness[0.01]},
      PlotMarkers → {"●", Large}], {τ, {1, 2, 5, 10, 20, Finalτ}}]]],
  Export[StringJoin[ToString[NotebookDirectory[]],
    StringJoin["Fig.example.partialwpartialx.p=", ToString[p]], ".pdf"],
   Show[Table[ListLinePlot[Flatten[Partialwpartialx[Na, τ, p]],
      PlotStyle → PlotStyleFunction[τ], AxesStyle → Large,
      PlotRange → {0, 0.5}, PlotStyle → {Black, Thickness[0.01]},
      PlotMarkers → {"●", Large}], {τ, {1, 2, 5, 10, 20, Finalτ}}]]]}

PartialwpartialyxPlot[NA, Pp];
```

### Social development

```
In[67]:= Clear["Global`*"]

(*Choose parameters*)
NA = 4; (*number of ages*)
Pp = 0.7; (*survival probability*)
Qp = 0.5; (*rate of social interaction*)
Finalτ = 200;

ι[Na_, p_] := Sum[p^(a-1), {a, 1, Na}]
T[Na_, p_, q_] := Sum[a (1 - y[a]) x[a] × l[a, p] (1 + q)/(1 - q y[a]), {a, 1, Na}]
partialwpartialx[Na_, p_, q_] :=
 Table[1/T[Na, p, q] l[a, p] (1 - y[a]), {a, 1, Na}, {j, 1, 1}]
partialwpartialy[Na_, p_, q_] :=
 Table[- 1/T[Na, p, q] l[a, p] × x[a] (1 + q)/(1 - q y[a]), {a, 1, Na}, {j, 1, 1}]
```

```
partialxpartialy[Na_, q_] :=
 Table[Table[If[j == a + 1, x[a] (1 + q)/(1 - q y[a]), 0], {j, 1, Na}], {a, 1, Na}]
partialxpartialx[Na_] :=
 Table[Table[If[j == a + 1, 1 + y[a], If[j == a, 1, 0]], {j, 1, Na}], {a, 1, Na}]
totalxtotalx[Na_] :=
 Table[Table[If[j > a, Product[partialxpartialx[Na][[k, k + 1]], {k, a, j - 1}],
    If[j == a, 1, 0]], {j, 1, Na}], {a, 1, Na}]
totalxtotaly[Na_, q_] := partialxpartialy[Na, q].totalxtotalx[Na];
totalwtotalx[Na_, p_, q_] := totalxtotalx[Na].partialwpartialx[Na, p, q]
totalwtotaly[Na_, p_, q_] :=
 partialwpartialy[Na, p, q] + totalxtotaly[Na, q].partialwpartialx[Na, p, q]

l[a_, p_] := p^(a-1)
Hy[Na_, τ_] := Table[
  If[a == j, (ysol[τ])[[a, 1]] (1 - (ysol[τ])[[a, 1]]), 0], {a, 1, Na}, {j, 1, Na}]

(*Substitute solutions in total selection gradient*)
Totalwtotalx[Na_, τ_, p_, q_] :=
 totalwtotalx[Na, p, q] /. Table[y[a] → ysol[τ][[a, 1]], {a, 1, Na}] /.
  Table[x[a] → xsol[τ, a, 1], {a, 1, Na}]
Totalwtotaly[Na_, τ_, p_, q_] :=
 totalwtotaly[Na, p, q] /. Table[y[a] → ysol[τ][[a, 1]], {a, 1, Na}] /.
  Table[x[a] → xsol[τ, a, 1], {a, 1, Na}]

(*Initial conditions*)
ysol[1] = Table[1/2, {a, 1, 4}, {j, 1, 1}];

Runs[Na_, T_, p_, q_] := Table[{xsol[τ, 1, 1] = 1;
   Table[xsol[τ, a + 1, 1] = (1 + ysol[τ][[a, 1]])/(1 - q ysol[τ][[a, 1]]) xsol[τ, a, 1], {a, 1, Na - 1}];
   ysol[τ + 1] = ysol[τ] + ι[Na, p] × Hy[Na, τ].Totalwtotaly[Na, τ, p, q];},
  {τ, 1, T}]

(*Plots of runs*)
PlotStyleFunction[τ_, T_] := If[τ == T, {Black, Thickness[0.01]},
  If[τ == 1, {Black, Dashing[0.05], Thickness[0.01]}, {Gray}]]
PlotRuns[Na_, T_, p_, q_] := {Runs[Na, T, p, q];,
  Export[StringJoin[ToString[NotebookDirectory[]], StringJoin[
     "Fig.example.social.control.p=", ToString[p], ".q=", ToString[q]],
    ".pdf"], Show[Table[ListLinePlot[Flatten[ysol[τ]], PlotRange → {-0.1, 1.1},
      PlotStyle → PlotStyleFunction[τ, T], AxesStyle → Large,
      PlotMarkers → {"●", Large}], {τ, {1, 2, 5, 10, 20, Finalτ}}]]],
  Export[StringJoin[ToString[NotebookDirectory[]], StringJoin[
     "Fig.example.social.state.p=", ToString[p], ".q=", ToString[q]], ".pdf"],
   Show[Table[ListLinePlot[Table[xsol[τ, a, 1], {a, 1, Na}], PlotRange → {0, 20},
      PlotStyle → PlotStyleFunction[τ, T], AxesStyle → Large,
```

```
      PlotMarkers → {"●", Large}], {τ, {1, 2, 5, 10, 20, Finalτ}}]]]}

(*Do runs and plots*)
PlotRuns[NA, Finalτ, Pp, Qp];

(*Lz plots*)

(*To fix color scale in matrix plot*)
cf = Blend[{{0., RGBColor[0.260487, 0.356, 0.891569]},
     {0.166667, RGBColor[0.230198, 0.499962, 0.848188]},
     {0.333333, RGBColor[0.392401, 0.658762, 0.797589]},
     {0.499999, RGBColor[0.964837, 0.982332, 0.98988]},
     {0.5, RGBColor[1, 1, 1]}, {0.500001, RGBColor[0.95735, 0.957281, 0.896269]},
     {0.666667, RGBColor[0.913252, 0.790646, 0.462837]},
     {0.833333, RGBColor[0.860243, 0.558831, 0.00695811]},
     {1., RGBColor[1., 0.42, 0.]}}, #1] &;
(*cfScaled=cf@Rescale[#,{-1,1},{0,1}]&;*)
cfScaled = cf@Rescale[#, {-50, 50}, {0, 1}] &;

(*Compute Hx*)
Totalxtotaly[Na_, τ_, q_] :=
 totalxtotaly[Na, q] /. Table[y[a] → ysol[τ][[a, 1]], {a, 1, Na}] /.
  Table[x[a] → xsol[τ, a, 1], {a, 1, Na}]
Hx[Na_, τ_, q_] := Transpose[Totalxtotaly[Na, τ, q]].
  Hy[Na, τ].Totalxtotaly[Na, τ, q]

(*Compute Hz*)
Totalztotaly[Na_, τ_, q_] :=
 ArrayFlatten[{{Totalxtotaly[Na, τ, q], IdentityMatrix[Na]}}]
Hz[Na_, τ_, q_] := Transpose[Totalztotaly[Na, τ, q]].
  Hy[Na, τ].Totalztotaly[Na, τ, q]

(*Compute Lz*)
partialxpartialxbar[Na_, q_] :=
 Table[Table[If[j == a, y[a] q, 0], {j, 1, Na}], {a, 1, Na}]
Partialxpartialxbar[Na_, τ_, q_] :=
 partialxpartialxbar[Na, q] /. Table[y[a] → ysol[τ][[a, 1]], {a, 1, Na}] /.
  Table[x[a] → xsol[τ, a, 1], {a, 1, Na}]
totalxtotalxbar[Na_, q_] := partialxpartialxbar[Na, q].totalxtotalx[Na]
Totalxtotalxbar[Na_, τ_, q_] :=
 totalxtotalxbar[Na, q] /. Table[y[a] → ysol[τ][[a, 1]], {a, 1, Na}] /.
  Table[x[a] → xsol[τ, a, 1], {a, 1, Na}]
sxsxbarTranspose[Na_, τ_, q_] :=
 Inverse[IdentityMatrix[Na] - Transpose[Totalxtotalxbar[Na, τ, q]]]
sxsyTranspose[Na_, τ_, q_] :=
 sxsxbarTranspose[Na, τ, q].Transpose[Totalxtotaly[Na, τ, q]]
szsyTranspose[Na_, τ_, q_] :=
```

```
 ArrayFlatten[{{sxsyTranspose[Na, τ, q]}, {IdentityMatrix[Na]}}]

Lz[Na_, τ_, q_] := szsyTranspose[Na, τ, q].Hy[Na, τ].Totalztotaly[Na, τ, q]

(*Plots of Lz*)

LzPlot[Na_, p_, q_] := Table[Export[StringJoin[ToString[NotebookDirectory[]],
    StringJoin["Fig.example.social.Lz.p=", ToString[p], ".tau=", ToString[τ]],
    ".pdf"], MatrixPlot[Lz[Na, τ, q],
    FrameTicks → {{{{1, 1}, {2, 2}, {3, 3}, {4, 4}, {5, 1}, {6, 2}, {7, 3}, {8, 4}},
        {{1, 1}, {2, 2}, {3, 3}, {4, 4}, {5, 1}, {6, 2}, {7, 3}, {8, 4}}},
       {{{1, 1}, {2, 2}, {3, 3}, {4, 4}, {5, 1}, {6, 2}, {7, 3}, {8, 4}},
        {{1, 1}, {2, 2}, {3, 3}, {4, 4}, {5, 1}, {6, 2}, {7, 3}, {8, 4}}}},
    FrameStyle → Large, PlotLegends → BarLegend[{Automatic, {-50, 50}},
      LabelStyle → Large], ColorFunction → cfScaled,
    ColorFunctionScaling → False]], {τ, {1, 2, 5, 10, 20, Finalτ}}]

(*Do plots of Lz*)
LzPlot[NA, Pp, Qp];

(*Plots of the total selection gradients*)

Totalwtotalz[Na_, τ_, p_, q_] :=
 ArrayFlatten[{{Totalwtotalx[Na, τ, p, q]}, {Totalwtotaly[Na, τ, p, q]}}]
dwdyxPlot[Na_, T_, p_, q_] := {Export[StringJoin[ToString[NotebookDirectory[]],
    StringJoin["Fig.example.social.dwdy.p=", ToString[p]], ".pdf"],
   Show[Table[ListLinePlot[Flatten[Totalwtotaly[Na, τ, p, q]],
      PlotStyle → PlotStyleFunction[τ, T], AxesStyle → Large,
      PlotRange → {-0.4, 0.1}, PlotStyle → {Black, Thickness[0.01]},
      PlotMarkers → {"●", Large}], {τ, {1, 2, 5, 10, 20, Finalτ}}]]],
  Export[StringJoin[ToString[NotebookDirectory[]],
    StringJoin["Fig.example.social.dwdx.p=", ToString[p]], ".pdf"],
   Show[Table[ListLinePlot[Flatten[Totalwtotalx[Na, τ, p, q]],
      PlotStyle → PlotStyleFunction[τ, T], AxesStyle → Large,
      PlotRange → {0, 0.5}, PlotStyle → {Black, Thickness[0.01]},
      PlotMarkers → {"●", Large}], {τ, {1, 2, 5, 10, 20, Finalτ}}]]]}
dwdyxPlot[NA, Finalτ, Pp, Qp];

(*Plots of the partial selection gradients*)

Partialwpartialx[Na_, τ_, p_, q_] :=
 partialwpartialx[Na, p, q] /. Table[y[a] → ysol[τ][[a, 1]], {a, 1, Na}] /.
  Table[x[a] → xsol[τ, a, 1], {a, 1, Na}]
Partialwpartialy[Na_, τ_, p_, q_] :=
 partialwpartialy[Na, p, q] /. Table[y[a] → ysol[τ][[a, 1]], {a, 1, Na}] /.
  Table[x[a] → xsol[τ, a, 1], {a, 1, Na}]
```

```
PartialwpartialyxPlot[Na_, T_, p_, q_] :=
 {Export[StringJoin[ToString[NotebookDirectory[]],
     StringJoin["Fig.example.social.partialwpartialy.p=", ToString[p]], ".pdf"],
    Show[Table[ListLinePlot[Flatten[Partialwpartialy[Na, τ, p, q]],
       PlotStyle → PlotStyleFunction[τ, T], AxesStyle → Large,
       PlotRange → {-0.4, 0.1}, PlotStyle → {Black, Thickness[0.01]},
       PlotMarkers → {"●", Large}], {τ, {1, 2, 5, 10, 20, Finalτ}}]]],
  Export[StringJoin[ToString[NotebookDirectory[]],
     StringJoin["Fig.example.social.partialwpartialx.p=", ToString[p]], ".pdf"],
    Show[Table[ListLinePlot[Flatten[Partialwpartialx[Na, τ, p, q]],
       PlotStyle → PlotStyleFunction[τ, T], AxesStyle → Large,
       PlotRange → {0, 0.5}, PlotStyle → {Black, Thickness[0.01]},
       PlotMarkers → {"●", Large}], {τ, {1, 2, 5, 10, 20, Finalτ}}]]]}

PartialwpartialyxPlot[NA, Finalτ, Pp, Qp];
```
